# A Clock-Driven Neural Network Critical for Arousal

**DOI:** 10.1101/2020.03.12.989921

**Authors:** Benjamin J. Bell, Qiang Liu, Dong Won Kim, Sang Soo Lee, Qili Liu, Ian D. Blum, Annette A. Wang, Joseph L. Bedont, Anna J. Chang, Habon Issa, Jeremiah Y. Cohen, Seth Blackshaw, Mark N. Wu

## Abstract

The daily cycling of sleep and arousal states is among the most prominent biological rhythms under circadian control. While much is known about the core circadian clock^1,2^, how this clock tunes sleep and arousal remains poorly understood^3^. In *Drosophila*, we previously characterized WIDE AWAKE (WAKE), a clock-output molecule that promotes sleep at night^4,5^. Here, we show that the function of WAKE in regulating circadian-dependent neural excitability and arousal is conserved in mice. *mWake^+^* cells are found in the suprachiasmatic nucleus (SCN) and dorsomedial hypothalamus (DMH). *mWake^DMH^* neurons drive wakefulness and exhibit rhythmic spiking, with greater firing during the night vs the day. Loss of mWAKE leads to increased spiking of *mWake^+^* SCN and DMH neurons and prominent behavioural arousal, specifically during the night. Single-cell sequencing, imaging, and patch-clamp experiments reveal that *mWake^DMH^* neurons constitute a glutamatergic/GABAergic population that projects widely, receives neuromodulatory input, and acts on neuromodulatory neurons. Strikingly, broad chemogenetic silencing of *mWake^+^* cells leads to profound loss of behavioural responsiveness and low amplitude, low frequency electroencephalography waveforms. These findings suggest that the genetic mechanisms regulating circadian control of sleep and arousal are conserved across >500 million years of evolution and define a clock-regulated neural network critical for arousal.

## The function of WAKE is conserved in mammals

We previously identified the clock-output molecule WIDE AWAKE (WAKE) from a forward genetic screen in *Drosophila*^4^. WAKE modulates the activity of arousal-promoting clock neurons at night, in order to promote sleep onset and quality^4,5^. The mammalian proteome contains a single ortholog, mWAKE (also named ANKFN1/Nmf9), with 56% sequence similarity and which is enriched in the core region of the master circadian pacemaker suprachiasmatic nucleus (SCN)^4,6^ (Fig. 1a, Extended Data Fig. 1a). To investigate whether the function of WAKE is conserved in mice, we generated a putative null allele of *mWake* (*mWake^(-)^*) using CRISPR/Cas9 (Fig. 1b and see Methods). As expected, *mWake* expression, as assessed by quantitative PCR and *in situ* hybridization (ISH), was markedly reduced in *mWake^(-/-)^* mice, likely due to nonsense-mediated decay (Fig. 1c, 1d). Given *mWake* expression in the SCN, we first examined locomotor circadian rhythms and found that *mWake^(-/-)^* mice exhibit a mild but non-significant decrease in circadian period length (Extended Data Fig. 1b, 1c). These results are similar to findings from fly *wake* mutants and mice bearing the *Nmf9* mutation (a previously identified ENU-generated allele of *mWake*)^4,6^.

**Figure 1.**
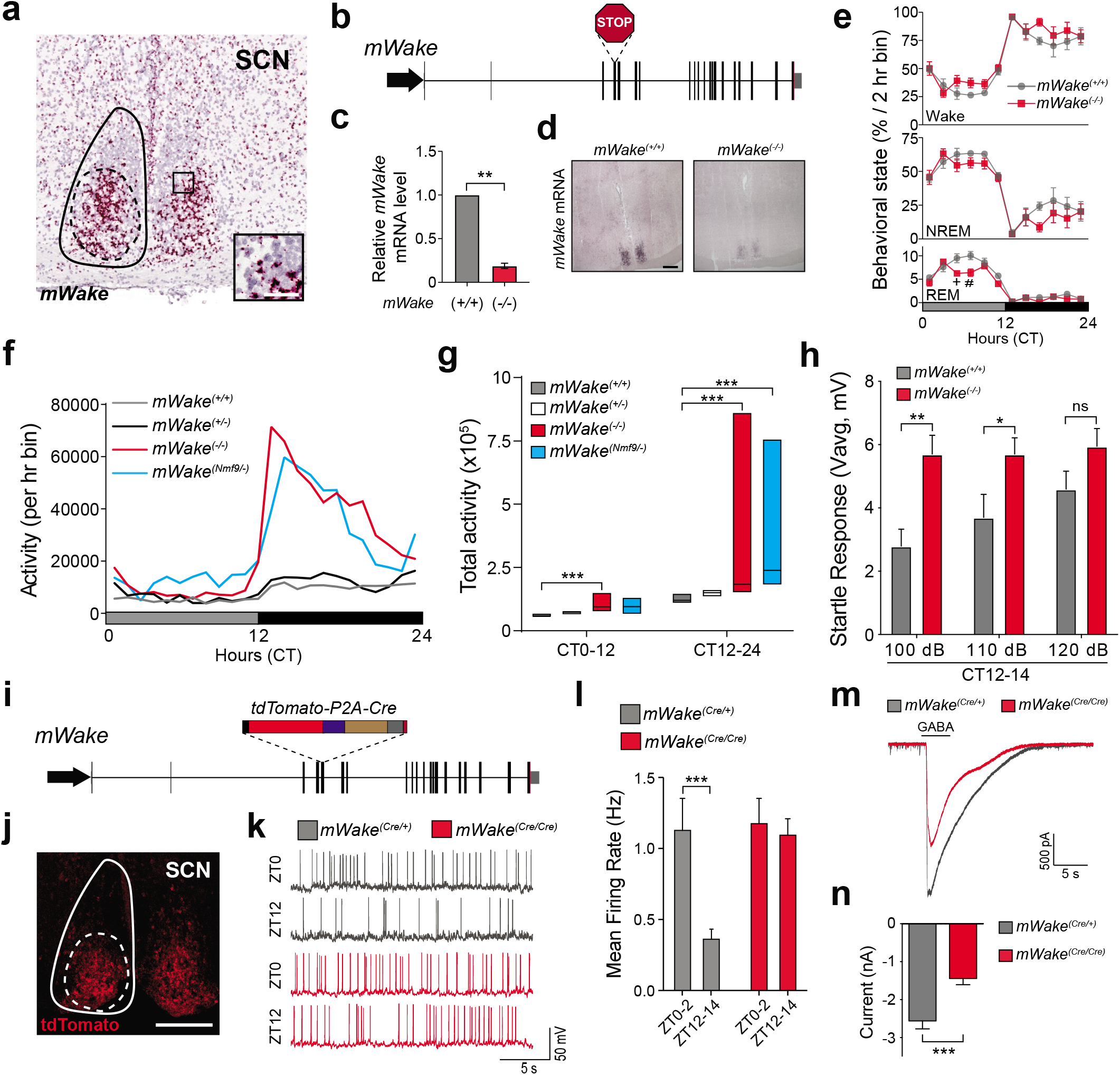
mWAKE inhibits arousal and SCN firing at night. **a**, *mWake* mRNA detected by BaseScope ISH demonstrates expression in the SCN core (solid and dashed lines denote SCN and core region, respectively). Higher magnification inset shows representative *mWake* mRNA expression in cells. Scale bar applies to both images and denotes 200 μm and 50 μm for the entire image and inset, respectively. **b**, Schematic showing genomic structure of the *mWake* locus and CRISPR/Cas9-mediated insertion of an in-frame stop codon in exon 4 in the *mWake^-^* mutation. **c**, Relative mRNA level for *mWake*, determined by qPCR, in *mWake^(-/-)^* vs WT littermate control hypothalami (normalized to 1.0) (n=3 replicates). **d**, Representative images (of 4 replicates each) of ISH using a digoxigenin-labeled *mWake* RNA probe in the SCN of *mWake^(-/-)^* vs WT mice. Scale bar denotes 500 μm. **e**, Vigilance state (% per 2 hr bin) determined by EEG recordings for *mWake^(-/-)^* (n=8) vs WT littermate control (n=6) mice under DD conditions. “^+^” and “^#^” denote *P*<0.01 and *P*<0.001, respectively. **f**, Profile of locomotor activity (defined by beam-breaks) over 24 hrs for *mWake^(+/+)^* (n=19, grey), *mWake^(+/-)^* (n=22, black), *mWake^(-/-)^* (n=19, red), and *mWake^(Nmf9/-)^* (n=9, cyan) mice under DD conditions. **g**, Total locomotor activity (total number of beams broken along X and Y axes) from CT0-12 and CT12-24 from the mice described in (**f**). **h**, Startle response (Vavg) measured in the first 100 ms following a 100, 110, or 120 dB tone for *mWake^(+/+)^* (grey, n=10) vs *mWake^(-/-)^* mice (red, n=10) at CT12-14. The average of 5 responses is shown. **i**, Schematic showing genomic structure of the *mWake* locus and replacement of exon 5 with a *tdTomato-P2A-Cre* cassette in the *mWake^(Cre)^* line. **j**, Native tdTomato fluorescence in the SCN of a *mWake^(Cre/+)^* mouse (solid and dashed lines denote SCN and core region, respectively). Scale bar represents 200 μm. **k**, Representative membrane potential traces from whole-cell patch clamp recordings of *mWake^SCN^* neurons from *mWake^(Cre/+)^* (grey, top) and *mWake^(Cre/Cre)^* (red, bottom) slices at ZT0-2 and ZT12-14. **l**, Spontaneous mean firing rate for *mWake^+^* SCN neurons at ZT0-2 and ZT12-14 from *mWake^(Cre/+)^* (n=21 and n=22) vs *mWake^(Cre/Cre)^* (n=24 and n=20) mice. **m**, Representative traces of voltage-clamp recordings of *mWake^SCN^* neurons of *mWake^(Cre/+)^* vs *mWake^(Cre/Cre)^* at ZT12-14. Timing of GABA (1 mM) application is shown. **n**, GABA-evoked current in *mWake^SCN^* neurons of *mWake^(Cre/+)^* (n=18, grey) vs *mWake^(Cre/Cre)^* (n=20, red) at ZT12-14. For panels (**l**) and (**n**), n represents individual cells from 4 animals for each condition. In this figure and throughout, error bars represent SEM and “*”, “**”, and “***” denote *P*<0.05, *P*<0.01, and *P*<0.001, respectively.

Because we previously demonstrated that WAKE mediates circadian regulation of sleep timing and quality in fruit flies^4,5^, we next assessed sleep in *mWake^(-/-)^* mice via electroencephalography (EEG). Under light:dark (LD) conditions, there was no difference in the amount of wakefulness, non-rapid eye movement (NREM), or REM sleep between *mWake^(-/-)^* mutants and wild-type (WT) littermate controls (Extended Data Fig. 1d). In constant darkness (DD), there is a modest main effect of genotype on wakefulness (*P*<0.05) and NREM sleep (*P*<0.05), and a mild but significant decrease in REM sleep in *mWake^(-/-)^* mutants (Fig. 1e). Although the amount of wakefulness did not appreciably differ in *mWake^(-/-)^* mutants compared to controls, there was a change in the distribution of wakefulness at night; mutants spent more daily time in prolonged wake bouts, with some (~40%) exhibiting dramatically long (>6 hrs) bouts of wakefulness (Extended Data Fig. 1e, 1f).

Because fly WAKE mainly functions at night and mice (as nocturnal animals) are generally awake at that time, we reasoned that arousal, and not sleep, may be primarily affected in *mWake* mutants. Alterations in arousal level can be quantified across different parameters, including sleep/wake behaviour, locomotor activity, and responsiveness to sensory stimuli^7^. Thus, we next examined baseline homecage locomotor activity (Extended Data Fig. 2a). *mWake*^(-/-)^ mutants were markedly hyperactive during the subjective night compared to littermate controls, although a mild but significant increase in locomotor activity was also noted during the subjective day (Fig. 1f, 1g, Supplementary Video 1). To rule out the possibility of 2^nd^ site mutations causing this phenotype, we examined transheterozygous *mWake^(Nmf9/-)^* mutants, which also demonstrated robust locomotor hyperactivity during the night, but not during the day (Fig. 1f, 1g). Similar data were obtained for *mWake^(-/-)^* and *mWake^(Nmf9/-)^* mice under LD conditions (Extended Data Fig. 2b, 2c). Locomotor activity for heterozygous *mWake^(+/-)^* mice was not different from littermate controls, and there was a trend for it being reduced at night, compared to *mWake^(-/-)^* (*P*=0.13) and *mWake^(Nmf9/-)^* (*P*=0.06) (Fig. 1f, 1g). The variability of the pronounced nighttime locomotor activity in *mWake^(-/-)^* and *mWake^(Nmf9/-)^* mutants was driven by intense stereotypic circling behaviour in a subset (~30-40%) of these animals (Supplementary Video 1). Related to this, *mWake* was previously identified as *Nmf9*, and mutations in this gene were noted to cause circling behaviour, which was interpreted to be due to deficits in vestibular function^6^. However, the intense coordinated circling behaviour demonstrated by *mWake* mutants has also been observed in hyperaroused mice^8–10^. In addition, in our swim tests, *mWake^(-/-)^* and *mWake^(Nmf9/-)^* mutants swam vigorously without impairment and never had to be rescued from potential drowning (Supplementary Video 2), which indicates normal vestibular function and contrasts with the findings of Zhang et al. (2015). Thus, our interpretation is that these and other phenotypes described for *mWake^(Nmf9/Nmf9)^* mutants stem from their hyperarousal, although we cannot rule out subtle vestibular dysfunction requiring specialized testing.

In addition to baseline locomotor activity, another measure of arousal is sensory responsiveness^7^. To characterize this phenotype, we evaluated startle response to an acoustic stimulus in *mWake^(-/-)^* mutants during the subjective day and subjective night. *mWake^(-/-)^* mutants exhibited an increased startle response to 100 dB and 110 dB stimuli during subjective night, but not during the subjective day (Fig. 1h, Extended Data Fig. 2d). To examine provoked locomotor arousal, we performed open-field tests and found that *mWake^(-/-)^* mutants were hyperactive both during the day and night and, unlike controls, failed to demonstrate habituation (Extended Data Fig. 2e-h). Taken together, these data suggest that mWAKE mainly acts to suppress arousal at night, but that specific provoked conditions can also reveal underlying hyperarousal in *mWake* mutants during the day.

In *Drosophila*, we previously showed that the large ventrolateral (l-LNv) photorecipient clock neurons in *wake* mutants lose the rhythmicity of their firing at dawn vs dusk. This phenotype results from an increase in spiking frequency specifically at night in the mutants, which is likely due to a reduction in GABA sensitivity in these clock neurons^4^. We thus asked whether loss of mWAKE would cause a similar phenotype in *mWake*^+^ clock neurons from the photorecipient core region of the SCN. To genetically target *mWake^+^* neurons while simultaneously creating a mutant allele, we generated transgenic mice where exon 5 was replaced with a tdTomato-P2A-Cre cassette, which should produce a null allele (*mWake^Cre^*) (Fig. 1i). As predicted, this transgene labels neurons in the core region of the SCN (Fig. 1j), and we further confirmed the overall fidelity of the expression pattern of this transgene, by comparing it to data from whole-brain RNAscope ISH labeling of *mWake*^11^. We also assessed locomotor activity in DD in *mWake^(Cre/Cre^*^)^ animals vs heterozygous controls and found that the homozygous animals phenocopied the nighttime hyperactivity of *mWake^(-/-)^* and *mWake^(Nmf9/-)^* mutants, arguing that *mWake^Cre^* is a bona fide *mWake* mutant allele (Extended Data Fig. 2i, 2j).

Patch-clamp recordings revealed a loss of cycling of spiking frequency in *mWake*^+^ SCN neurons, due to a selective increase in firing rate at night in *mWake^(Cre/Cre^*^)^ mutants (Fig. 1k, 1l, Extended Data Fig. 3a, Supplementary Table 1). The effect of mWAKE on neuronal excitability is likely cell-autonomous, as intrinsic excitability of *mWake^SCN^* neurons in these mutants was increased during the night, but not the day (Extended Data Fig. 3b, 3c). Also analogous to fly *wake* mutants, recordings from *mWake^SCN^* neurons revealed a decrease in GABA-evoked current at night in *mWake^(Cre/Cre)^* mice (Fig. 1m, 1n). This phenotype was also observed in mutants lacking the core clock protein Bmal1^12^, suggesting that mWAKE and BMAL act in the same pathway (Extended Data Fig. 3d, 3e). These findings suggest the basic function of WAKE in clock neurons is shared between flies and mice. However, in contrast to fly WAKE, mWAKE action in the SCN is not crucial for suppressing arousal. Conditional knockout of mWAKE in the ventral forebrain, including the SCN, using the *Six3-Cre* driver^13^ (*Six3^(Cre/+)^>mWake^(flox/-)^* mice) did not alter locomotor activity (Extended Data Fig. 4a-e). Thus, we next sought to identify the brain region(s) where mWAKE acts to regulate arousal.

## mWAKE defines a circadian-dependent arousal circuit in the DMH

Beyond the SCN, mWAKE is expressed in a variety of regions across the brain, such as other hypothalamic sub-regions (including the dorsomedial hypothalamus/DMH, Fig. 2a, 2b, also see Fig. 3a), areas implicated in arousal, the limbic system, sensory processing nuclei, and limited regions in the cortex^6,11^. The DMH has previously been implicated in the circadian regulation of arousal^14–16^, although the specific neurons involved remain unknown. We thus asked whether *mWake^DMH^* neurons mediate circadian-dependent arousal. Interestingly, in controls, these neurons had greater spiking frequency during the night vs the day, which is antiphase to the SCN, but aligned with the active phase of the nocturnal mouse (0.55 ± 0.10 Hz at ZT0-2 vs 1.16 ± 0.15 Hz at ZT12-14 in *mWake^(Cre/+)^*, *P*<0.01). Moreover, this nighttime increase was accentuated in *mWake^(Cre/Cre)^* mutants (Fig. 2c, 2d, Extended Data Fig. 5a, Supplementary Table 1). Similarly, intrinsic excitability of *mWake^DMH^* neurons was greater in *mWake^(Cre/Cre)^* mutants during the night, but not the day (Extended Data Fig. 5b, 5c). These data are consistent with a circadian-dependent role for *mWake^DMH^* neurons in promoting arousal.

**Figure 2.**
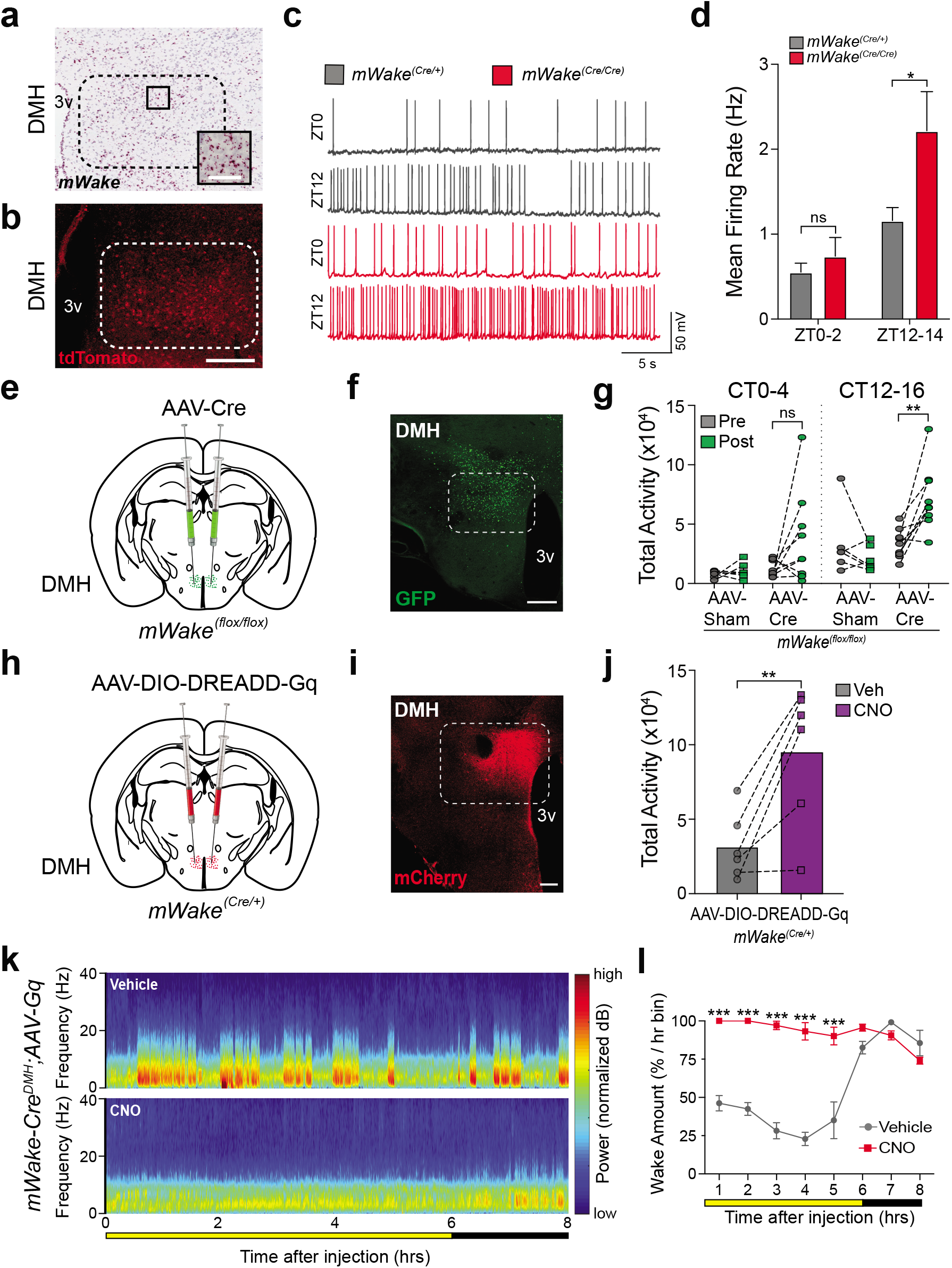
*mWake^DMH^* neurons drive arousal. **a**, *mWake* mRNA detected by Basescope ISH reveals expression in the dorsomedial hypothalamus (DMH). Dashed lines indicate DMH region, and inset shows representative *mWake* mRNA expression in cells. Scale bar applies to both images and denotes 200 μm and 50 μm for the entire image and inset, respectively. **b**, Native tdTomato fluorescence in the DMH of a *mWake^(Cre/+)^* mouse (dashed lines denote DMH region). Scale bar represents 200 μm. **c**, Representative membrane potential traces from whole-cell patch clamp recordings of *mWake^DMH^* neurons in *mWake^(Cre/+)^* (grey, top) and *mWake^(Cre/Cre)^* (red, bottom) slices at ZT0-2 and ZT12-14. **d**, Spontaneous mean firing rate for *mWake^DMH^* neurons at ZT0-2 and ZT12-14 from *mWake^(Cre/+)^* (n=19 and n=14) vs *mWake^(Cre/Cre)^* (n=16 and n=12) mice. n represents individual cells from >4 animals. **e**, Schematic showing bilateral injections of AAV viral vector containing Cre-recombinase and eGFP (AAV-Cre), or eGFP alone (AAV-Sham) into the DMH of *mWake^(flox/flox)^* mice. **f**, Representative image of eGFP fluorescence expression in the DMH following AAV-Cre injection described in (**e**). Scale bar denotes 200 μm. **g**, Total locomotor activity (total number of beams broken along X and Y axis) during CT0-4 vs CT12-16 under DD conditions for *mWake^(flox/flox)^* mice before (“pre”, grey) and after (“post”, green) DMH injection of AAV-Sham (n=6) or AAV-Cre (n=9). **h**, Schematic showing bilateral injections of AAV-DIO-DREADD-Gq into the DMH of *mWake^(Cre/+)^* mice. **i**, Representative image of mCherry fluorescence expression in the DMH of *mWake(Cre/+*) following AAV-DIO-DREADD-Gq injection described in (**h**). Scale bar denotes 200 μm. **j**, Total locomotor activity (total number of beams broken along X and Y axis) of *mWake^(Cre/+)^* mice with bilateral injections of AAV-DIO-DREADD-Gq into the DMH in the 4 hrs following IP injection of vehicle vs CNO (1 mg/kg) at CT 8 (n=6). **k**, Representative short-time Fourier transform spectrograms of 8 hrs of recorded EEG activity, starting after IP injection at ZT6 of vehicle alone (above) or 1 mg/kg CNO (below), from *mWake^(Cre/+)^* mice injected with AAV-DIO-DREADD-Gq bilaterally into the DMH. Power density is represented by the color-scheme and deconvoluted by frequency on the y-axis, over time on the x-axis. **l**, Amount of wakefulness derived from EEG plotted as % time in 1 hr bins following IP injection of vehicle (grey) or 1 mg/kg CNO (red) (n=4) at ZT6. “3v” denotes third ventricle.

**Figure 3.**
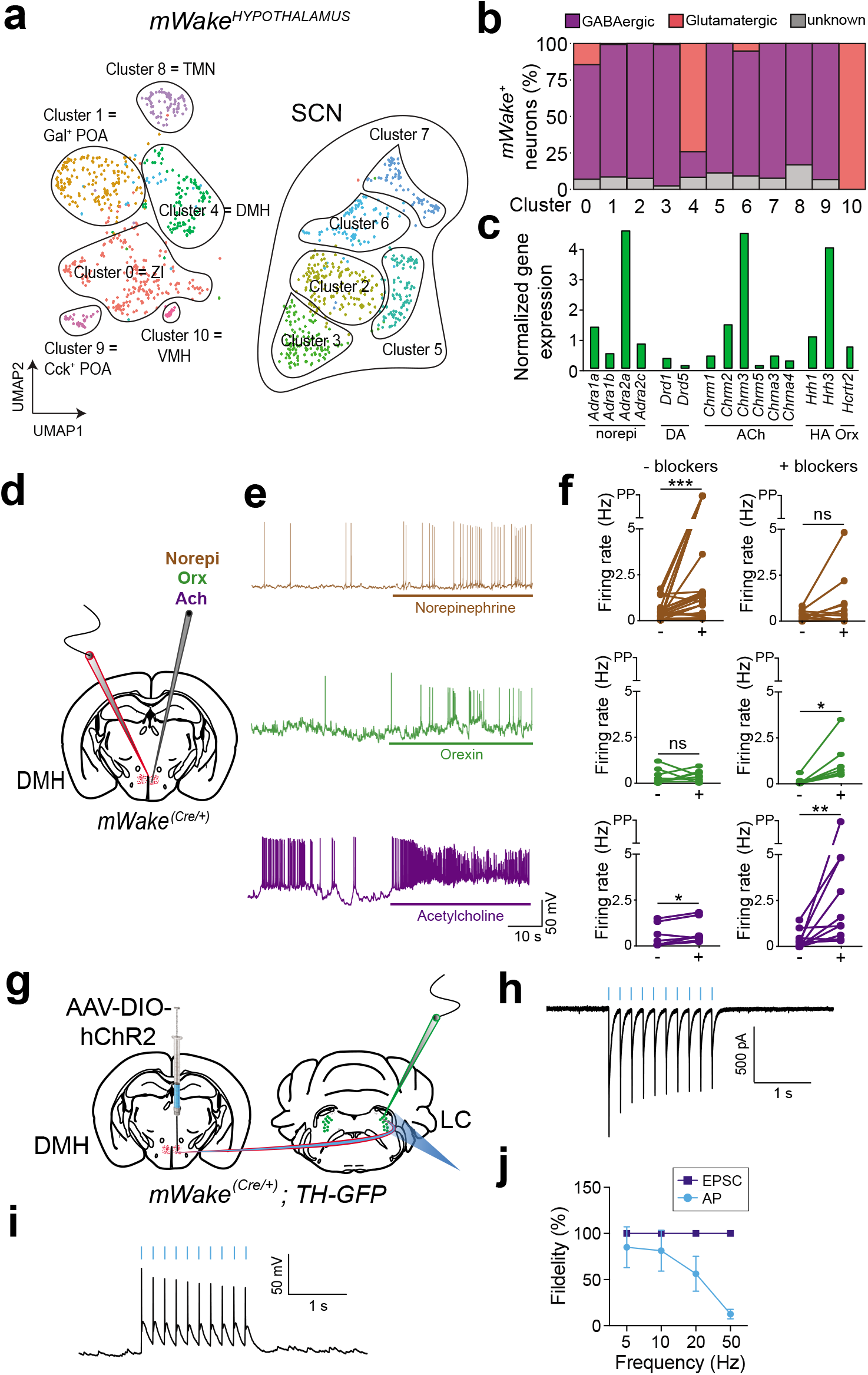
Characterization of *mWake^DMH^* neurons and their interaction with neuromodulatory systems. **a**, UMAP (Uniform Manifold Approximation and Projection) plot showing distribution of *mWake*^+^ neurons across hypothalamic nuclei, as determined by single-cell expression profiling. 11 clusters are defined, for SCN, DMH, POA (pre-optic area), TMN (tuberomammillary nucleus), ZI (zona incerta), and VMH (ventromedial hypothalamus) regions. “Gal^+^” and “Cck^+^” refer to Galanin^+^ and Cholecystokinin^+^. **b**, Bar graph showing proportions of GABAergic and glutamatergic *mWake*^+^ neurons for each scRNA-Seq neuronal cluster. **c**, Bar graph showing distribution of expression for norepinephrine, dopamine, acetylcholine, histamine and orexin receptors identified in DMH *mWake*^+^ neurons (cluster 4). **d**, Schematic showing patch-clamp recordings of *mWake^DMH^* neurons from *mWake^(Cre/+)^* mice, following application of norepinephrine, orexin, or acetylcholine. **e**, Representative membrane potential traces from recordings described in (**d**) following administration of norepinephrine (top, red), orexin (middle, green), or acetylcholine (bottom, blue) in the presence of synaptic blockers at ZT12-14. **f**, Spontaneous mean firing rate of *mWake^DMH^* neurons following application of 100 μM norepinephrine (n=23 and 12 cells), 300 nM orexin-A (n=8 and 7 cells), or 1 mM acetylcholine chloride (n=8 and 10 cells) at ZT12-14 in the absence (“-blockers”) or presence (“+ blockers”) of 20 μM CNQX, 50 μM AP5, and 10 μM picrotoxin. The neurons recorded in **f** were derived from 3-6 animals. **g**, Schematic showing AAV-DIO-hChR2 injected into the DMH of *mWake^(Cre/+)^;TH-GFP* mice unilaterally, and whole-cell patch clamp recordings of *NE^LC^* neurons, following 480 nm blue-light stimulation. **h** and **i**, Representative excitatory post-synaptic currents (EPSCs) (**h**) or action potentials (APs) (**i**) from *NE^LC^* neurons, triggered by optogenetic stimulation of *mWake^DMH^* neuron terminals at 5 Hz. Blue lines indicate timing of blue-light pulses. **j**, Fidelity (%) of *NE^LC^* neuron EPSCs (dark blue) and APs (light blue) triggered by optogenetic stimulation of *mWake^DMH^* neuron terminals. A total of 7 *NE^LC^* cells from 3 independent virally-injected *mWake^(Cre/+)^;TH-GFP* mice were recorded, of which 4 exhibited optogenetically-triggered responses.

To address the function of mWAKE in the DMH, we performed conditional knockout of mWAKE. We generated a floxed allele of *mWake* (*mWake^flox^*) and performed stereotaxic injection of an AAV viral vector expressing Cre-recombinase (AAV-Cre) into the DMH of *mWake^(flox/flox)^* mice (Fig. 2e, 2f, Supplementary Table 2). Reduction of mWAKE in the DMH led to a significant increase in locomotor activity during subjective nighttime, compared to their baseline. During the subjective day, there was a non-significant trend (*P*=0.08) towards an increase in locomotor activity with conditional knockout of *mWake* in the DMH. No differences in locomotor activity were observed in sham-injected controls during the subjective day or night (Fig. 2g). These findings, coupled with the observation that *mWake* mutants demonstrate increased spiking of *mWake^DMH^* neurons at night (Fig. 2c, 2d), suggest that these neurons promote arousal or wakefulness.

To test this possibility, we chemogenetically activated these neurons by injecting an AAV vector carrying Cre-dependent DREADD-hM3Dq (AAV-DIO-DREADD-Gq)^17^ into the DMH of *mWake^(Cre/+)^* mice (Fig. 2h, 2i, Supplementary Table 2). CNO-mediated activation of *mWake^DMH^* neurons resulted in a substantial increase in locomotor activity, compared to vehicle-treated animals (Fig. 2j). Moreover, chemogenetic activation of *mWake^DMH^* neurons markedly increased wakefulness, with concomitant reductions in NREM and REM sleep (Fig. 2k, 2l, Extended Data Fig. 5d, 5e). We next performed chemogenetic inhibition of *mWake^DMH^* neurons, by injecting an AAV vector encoding Cre-dependent DREADD-hM4Di (AAV-DIO-DREADD-Gi) and found no significant effects on locomotor activity or amount of wakefulness or NREM sleep (Extended Data Fig. 5f-i, Supplementary Table 2). However, the amount of REM sleep was reduced following injection of CNO, compared to vehicle alone (Extended Data Fig. 5j). In contrast, CNO alone administered to sham-injected *mWake^(Cre/+)^* mice did not appreciably affect locomotor activity or vigilance state (Extended Data Fig. 5k, 5l). Taken together, these data suggest that *mWake^DMH^* neurons promote arousal and that mWAKE acts to reduce arousal at night by inhibiting the activity of these neurons at that time.

To gain mechanistic insights into how *mWake^DMH^* neurons regulate arousal, we conducted single-cell RNA sequencing (scRNA-Seq) of FACS-sorted tdTomato^+^ cells from the hypothalami of *mWake^(Cre/+)^* mice (Fig. 3a, Extended Data Fig. 6a, 6b, Supplementary Table 3). In addition to neurons, there was a significant population of *mWake*^+^ ependymal cells, which were likely overrepresented due to their ability to survive the dissociation process (Extended Data Fig. 6a). The identity of different neuronal *mWake*^+^ clusters and the spatial location of these clusters were determined by comparison with a hypothalamic scRNA-Seq database and using specific gene markers (Extended Data Fig. 6c)^18^. This analysis revealed 11 *mWake*^+^ clusters in the hypothalamus, with a prominent DMH cluster and 5 SCN-specific clusters (Fig. 3a). Collectively, hypothalamic *mWake*^+^ neurons comprised a heterogeneous group, but were largely GABAergic (most clusters including all SCN clusters) or glutamatergic (DMH and VMH) (Fig. 3b). We confirmed this observation using RNAscope ISH for the SCN and DMH (Extended Data Fig. 6d-g).

Historically, investigations of the neural basis of arousal have focused on neuromodulatory circuits, but there is growing recognition that GABAergic and glutamatergic neurons likely form the core substrate for arousal, which is in turn tuned by neuromodulatory networks^3,19,20^. We thus used our scRNA-Seq dataset to characterize the repertoire of neuromodulatory receptors in *mWake^DMH^* neurons in the DMH (Fig. 3c). *mWake^DMH^* neurons express specific noradrenergic, cholinergic, histaminergic, and orexinergic receptors, and so we conducted rabies virus retrograde tracing (Fig. 3c, Extended Data Fig. 7a, Supplementary Table 2) from these neurons to evaluate potential inputs (Extended Data Fig. 7a). We observed significant retrograde labeling of histidine decarboxylase^+^ (HDC) neurons in the tuberomammillary nucleus (TMN) (Extended Data Fig. 7b). We also found some labeling of orexinergic neurons in the lateral hypothalamus (LH) and cholinergic neurons in the basal forebrain (BF), as well as rare labeling of noradrenergic neurons in the locus coeruleus (LC) (Extended Data Fig. 7c-e).

To address whether *mWake^DMH^* neurons functionally respond to these neuromodulators, we performed whole-cell patch-clamp recordings of these neurons from brain slices of *mWake^(Cre/+)^* mice in the presence or absence of norepinephrine, orexin, acetylcholine, or histamine (Fig. 3d-f, Extended Data Fig. 7f-h, Supplementary Table 1). Application of orexin and acetylcholine resulted in a significant elevation in spontaneous firing rate of *mWake^DMH^* neurons in a cell-autonomous manner (i.e., in the presence of synaptic blockers) (Fig. 3e, 3f). Norepinephrine increased *mWake^DMH^* neuron spiking indirectly, consistent with the relative paucity of noradrenergic neurons labeled by retrograde tracing from *mWake^DMH^* neurons (Extended Data Fig. 7e). In contrast, administration of histamine directly reduced *mWake^DMH^* neuron firing rate (Extended Data Fig. 7f-h), which may reflect a relative enrichment of the inhibitory H3 receptor subtype in these neurons (Fig. 3c).

Neurons that comprise arousal-promoting nuclei can either act as local interneurons or project broadly to influence the activity of multiple brain regions, including the neocortex^21^. To assess which of these categories *mWake^DMH^* neurons belong to, we injected an AAV vector expressing Cre-dependent eGFP (AAV-FLEX-eGFP) into the DMH of *mWake^(Cre/+)^* mice and imaged the GFP^+^ projections (Extended Data Fig. 7i, 7j, Supplementary Table 2). *mWake^DMH^* neurons sent projections throughout the brain, including the BF, caudate/putamen, the corpus callosum, and regions in the brainstem including the LC. Prior work had suggested that the DMH region may mediate circadian timing of arousal by modulating LC firing, although the responsible neurons in the DMH were not identified^15^. We therefore asked whether *mWake^DMH^* neurons can regulate the activity of noradrenergic LC neurons. We injected an AAV vector encoding a Cre-dependent Channelrhodopsin2 (AAV-DIO-hChR2) into the DMH of *mWake^(Cre/+)^;TH-GFP* mice and then performed whole-cell patch-clamp recordings from noradrenergic *NE^LC^* neurons following blue-light stimulation of the terminals of *mWake^DMH^* neurons in the LC (Fig. 3g, Supplementary Table 2). In the majority of cases, optogenetic activation of *mWake^DMH^* triggered noradrenergic *NE^LC^* excitatory post-synaptic currents (EPSCs) and spiking (Fig. 3h-j). In summary, our findings suggest that glutamatergic *mWake^DMH^* neurons are arousal-promoting, project widely, and bidirectionally interact with neuromodulatory networks.

## The *mWake^+^* network is critical for arousal

Whether there is a “core substrate” for arousal is controversial, as silencing of various genetically-defined arousal-promoting neural circuits, either alone or in combination, generally leads to relatively mild phenotypes^22–27^. Recent models suggest that if core arousal networks exist, they may be glutamatergic or GABAergic in nature, rather than the classically-studied monoaminergic or cholinergic systems^19^. Moreover, emerging data suggest that GABA or glutamate-expressing neurons located in or near previously-defined arousal-associated nuclei are important for regulating arousal^27–31^. Interestingly, our data suggest that *mWake^+^* neurons are glutamatergic or GABAergic, and we have recently found that these cells can be found in several regions implicated in arousal (e.g., BF, TMN, vlPAG/DR, LH, PB)^3,11,19,20^.

We thus hypothesized that mWAKE defines a distributed arousal network. Because silencing *mWake^DMH^* neurons led to a mild phenotype (Extended Data Fig. 5h-j), we chose to inhibit the broad *mWake^+^* network. To do this, we crossed transgenic mice expressing Cre-dependent DREADD-hM4Di (*LSL-Gi*)^32^ to *mWake^(Cre/+)^* mice to generate *mWake^(Cre/+)^; LSL-Gi* progeny. We first examined the expression pattern of the DREADD-hM4Di in these mice by immunostaining and found that it largely recapitulated the original expression pattern of the *mWake^(Cre/+)^* mice (Extended Data Fig. 8a). We next characterized behavioural and EEG phenotypes of these mice. Within 15 min of CNO treatment, *mWake^(Cre/+)^;LSL-Gi* mice exhibited reduced spontaneous locomotion and exploratory behavior. After ~90 min, we observed a profound decrease in arousal (reduced or minimal responsiveness to gentle touch or acoustic stimuli, with maintenance of righting reflex) in these mice, which lasted 2-3 hrs (Fig. 4a, 4b; Supplementary Video 3). Strikingly, EEG analyses of these mice revealed a marked shift towards low amplitude, slow frequency waveforms following injection of CNO, but not vehicle control (Fig. 4c-f; Extended Data Fig. 8b, 8c). For all but one animal, both the behavioural and EEG phenotypes were reproducible and reversible, becoming indistinguishable from vehicle-injected animals after 24 hrs. However, one CNO-treated animal died after >24 hrs. These phenotypes were suggestive of a stupor-like state, rather than sleep, and indeed *mWake^(Cre/+)^;LSL-Gi* mice appeared to exhibit a rebound of sleep-like slow-wave activity after the effects of the CNO dissipated (Fig. 4c; Extended Data Fig. 8b, 8c). For comparison, we assessed the behavioural and EEG effects of chemogenetically silencing arousal-promoting HDC^+^ (histamine decarboxylase) neurons by repeating these experiments with *HDC^(Cre/+)^; LSL-Gi* mice. CNO treatment of these mice led to no discernable differences in behavioural responsiveness or EEG spectral power and amplitude (Extended Data Fig. 8d-h). These data suggest that the *mWake*^+^ network is essential for basic arousal.

**Figure 4.**
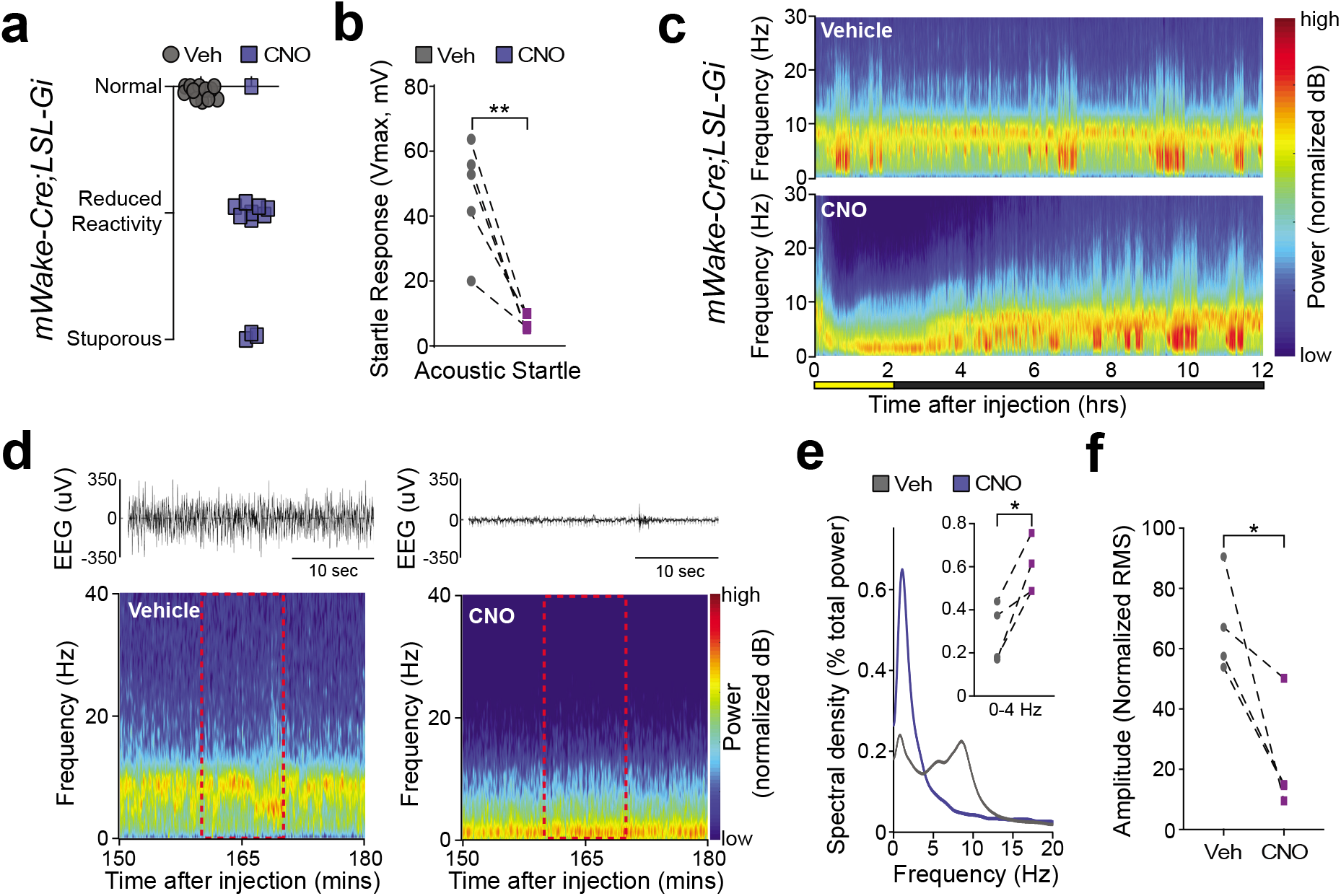
mWAKE defines a network critical for arousal. **a**, Jitter plot showing behavioural classification of arousal level for *mWake^(Cre/+)^; LSL-Gi* mice 90 min following injection of vehicle or 0.3 mg/kg CNO (n=12) at ZT12. **b**, Startle response (Vmax) measured in the first 100 ms following a 120 dB tone for *mWake^(Cre/+)^; LSL-Gi* mice 2 hr following injection of vehicle (grey) or 0.3 mg/kg CNO (magenta) (n=5) at CT12-14. **c**, Representative short-time Fourier transform spectrograms of 12 hrs of recorded EEG activity from *mWake^(Cre/+)^*; *LSL-Gi* mice after IP injection at ZT10 of vehicle alone (above) or 0.3 mg/kg CNO (below). **d**, Representative EEG traces for a 30 s window (above) and short-time Fourier transform spectrograms (below) starting 150 mins after injection of vehicle (left) or 0.3 mg/kg CNO (right) at ZT10. The dashed red box indicates the time window used for spectral and amplitude analysis in (**e**) and (**f**). **e**, Welch’s power spectral density estimates as a percentage of total EEG power across 10 min, averaged across multiple EEG traces. *mWake^(Cre/+)^; LSL-Gi* injected with vehicle (grey) or 0.3 mg/kg CNO (magenta) (n=4). Inset shows a plot of delta-band power as a percentage of total EEG power. **f**, EEG trace amplitude (plotted as normalized root mean square (RMS)) for the animals described in (**e**).

Our studies of WAKE in flies and mice suggest that the basic function of WAKE is to reduce arousal at night, by inhibiting the excitability of *mWake*^+^ arousal-promoting neurons at that time^4,5^. Although it is widely accepted that the circadian clock regulates sleep and arousal, the cellular mechanisms remain unclear. In principle, this process could be driven by direct action of SCN projections on arousal circuits^14,15,33^, by SCN release of diffusible substances^34–37^, or by the activity of local clocks in arousal-promoting nuclei^38^. Our data suggest that mWAKE mediates local clock control of arousal in the DMH, defining the first such neural circuit in this region at the genetic level. Moreover, the observations that mWAKE is expressed in or near several arousal-related regions^11^ and that *mWake^+^* neurons can project broadly (Extended Data Fig. 7j) suggest a new model for clock control of arousal—multi-focal local control, which allows for flexible modulatory input from environmental or internal state factors, yet also facilitates coordinated time-dependent regulation throughout the brain. Although many regions promoting wakefulness have been identified in the mammalian brain^3^, very few have been shown to be critical for fundamental arousal. One prominent example is the parabrachial nucleus (PB); large excitotoxic lesions in the PB lead to a coma-like state, with low amplitude, slow oscillations on EEG, a phenotype similar to that seen with silencing the *mWake*^+^ network^39^. For decades, psychologists have proposed the existence of distinct, but reciprocally connected, substrates for arousal: a “higher-order” modulatory system and a basal network, required for attention and consciousness^40^. Our data suggest the possibility that mWAKE marks a basal arousal network that is tuned by neuromodulatory inputs and under circadian clock control.

## Methods

### Animals

All animal procedures were approved by the Johns Hopkins Institutional Animal Care and Use Committee. All animals were group housed and maintained with standard chow and water available *ad libitum*. Animals were raised in a common animal facility under a 14:10 hr Light:Dark (LD) cycle. Unless otherwise noted, adult male mice (2-4 months) were used in all immunohistochemistry, *in situ* hybridization, and behavioural experiments. All mouse strains were backcrossed to *C57BL/6* at least seven times prior to use in behavioural experiments. Genotyping was performed either by in-house PCR and restriction digest assays, or via Taq-Man based rtPCR probes (Transnetyx). The *mWake^Nmf9^, HDC-Cre, TH-GFP*, and *Six3-Cre* mice were obtained from B. Hamilton (University of California, San Diego), A. Jackson (University of Connecticut), A. McCallion (Johns Hopkins University), and Y. Furuta (Memorial Sloan Kettering Cancer Center), respectively. *Bmall^-^* (stock number: 009100), and *LSL-Gi* (stock number: 026219) mice were obtained from The Jackson Laboratory.

The *mWake* null mutant allele (*mWake^-^*) was generated by CRISPR/Cas9 genome editing (Johns Hopkins Murine Mutagenesis Core), using a targeting guide RNA (gRNA: CGC AGA AGA ATC CTC GCA AT) to the 4^th^ exon of *mWake* and a 136 bp oligonucleotide (AGT GCG GAC TTT CTC TGG CTC CTG TCC GCA GAA GAA TCC TCG GCG GAA TTC AAT GGG CAC GTT GTT GGT CAT GAT GGC GAT GTC CAG GGG TGT CAG CCC TTC GCT GTT CGG TGT) containing two 64 bp homology arms, surrounding an 8 bp insertion (GCGGAATT), which includes an in-frame stop codon and induces downstream frameshifts. Exon 4 is predicted to be in all *mWake* splice isoforms. These constructs were injected into the pronucleus of *C57BL/6J* fertilized zygotes and implanted into pseudopregnant females. Pups were assayed for insertion by Sanger sequencing of *mWake* gDNA, and knockdown confirmed via qPCR and *in situ* hybridization. The conditional *mWake* knockout allele (*mWake^flox^*) was generated via homologous recombination in hybrid (*129/SvEv* x *C57BL/6*) mouse embryonic stem cells (Ingenious Targeting Laboratory). In the stem cells, a construct containing two loxP sites and a Neomycin cassette flanking the 4^th^ exon of *mWake* was integrated into the genomic DNA. These cells were then injected into *C57BL/6J* blastocysts, and offspring with high agouti content were crossed to flipase (FLP)- expressing *C57BL/6J* to remove the Neomycin selective marker. Offspring were then screened via Sanger sequencing to confirm proper insertion of both LoxP loci. A transgenic mouse line expressing both Cre recombinase and tdTomato in *mWake*^+^ cells (*mWake^Cre^*) was generated via homologous recombination in hybrid (*129/SvEv* x *C57BL/6*) mouse embryonic stem cells (Ingenious Targeting Laboratory). The knock-in vector targeted exon 5 of the *mWake* locus and was integrated 21 bp into exon 5, replacing the remainder of the exon with a *tdTomato-P2A-split Cre-Neo-WPRE-BGHpA* cassette, which causes frameshifted nonsense mutations downstream, resulting in an *mWake* loss-of-function allele. Neomycin was excised via crossing to FLP mice, and offspring sequenced to confirm the inclusion of the whole sequence into the *mWake* locus.

### Molecular Biology

To quantify *mWake* transcript in the *mWake^(-/-)^* mutant mice, qPCR was performed. Hypothalami were dissected at ~ZT0, and RNA extracted using Trizol Reagent (Invitrogen). qPCR was performed using a SYBR PCR master mix (Applied Biosystems) and a 7900 Real Time PCR system (Applied Biosystems), with the following primers, which target exon 4: *mWake-F:* 5’-CCC TAA CGG TCA GCT TTC AAG A-3’ and *mWake-R:* 5’-GAC ATG CTC CAT TCC ACT TTG TAC-3’. GAPDH was used as an internal control. Ct value was compared against regression standard curve of the same primers. 3 biological replicates were performed.

### Single Cell Sequencing

Seven week old, male *mWake^(Cre/+)^* mice were processed at ~ZT5 for single-cell RNA-Sequencing (scRNA-Seq). A modified Act-Seq^41^ method was used in conjunction with a previously described dissociation protocol^18^, with supplementation of Actinomycin D during dissociation (45 μM) and after final resuspension (3 μM), following debris removal (Debris Removal Solution (130-109-398, MACS Miltenyi Biotec) in between. 1 mm hypothalamic sections between Bregma 0.02 mm (collecting medial and lateral preoptic area) and Bregma −2.92 mm (beginning of the supramammillary nucleus) were collected, and 2-3 mice pooled per scRNA-Seq library.

Following dissociation, tdTomato^+^ cells were flow-sorted using an Aria IIu Sorter (Becton Dickinson). Between 400 − 1000 cells were flow-sorted per brain. Flow-sorted cells were pelleted and re-suspended in 47.6 μl resuspension media. 1 μl of flow-sorted tdTomato^+^ cells were used to quantify % of tdTomato^+^ cells with a phase-contrast microscope. Only samples containing ~99% flow-sorted tdTomato^+^ cells were processed for scRNA-Seq. The remaining 46.6 μl were used for the 10x Genomics Chromium Single Cell system (10x Genomics, CA, United States) using V3.0 chemistry per manufacturer’s instructions, generating a total of 3 libraries. Libraries were sequenced on Illumina NextSeq 500 with ~150 million reads per library (~200,000 median reads per cell). Sequenced files were processed through the CellRanger pipeline (v 3.1.0, 10x Genomics) using a custom mm10 genome (with *tdTomato-P2A-Cre-WPRE-bGH* sequence). All 3 libraries were aggregated together for downstream analysis.

Seurat V3^42^ was used to perform downstream analysis following the standard pipeline, using cells with more than 200 genes and 1000 UMI counts, removing *mWake-tdTomato*^+^ ependymal cells and non-*mWake*-cells (~1% of the total cluster composed of oligodendrocytes and astrocytes) using known markers genes in the initial clustering^18^. Louvain algorithm was used to generate different clusters, and spatial information (spatial location of different *mWake-tdTomato*^+^ clusters across hypothalamic nuclei) and identity of neuronal clusters were uncovered by referring to a hypothalamus scRNA-Seq database^18^. Region-specific transcription factors expressed in *mWake*^+^ neurons were used to train *mWake*-scRNA-Seq. Spatial information of different *mWake* neuronal populations were further validated by matching to the Allen Brain Atlas ISH data using cocoframer^43^, as well as matching to known *mWake* neuronal distribution across the hypothalamus^32^. The percentage of GABAergic (*Slc32a1*^+^) and Glutamatergic (*Slc17a6*^+^) neurons within each cluster was calculated.

Receptors for norepinephrine, dopamine, acetylcholine, histamine and orexin were identified in the scRNA-Seq dataset, and the normalized gene expression within cluster 4 (DMH) was calculated. scRNA-seq data from this study are accessible through GEO Series accession number GSE146166.

### Behavioural Analysis

Animals were entrained to a 12:12 hr LD cycle for at least 2 weeks before any locomotor or EEG-based behavioural experiments.

#### Homecage locomotor activity

Animals were separated into new individual cages with access to food and water *ad libitum* and allowed to acclimate for 4 days before data collection. Data were recorded over 2 days of 12:12 hr LD and 2 days of constant darkness (DD) cycles. Locomotor activity was recorded and analyzed using the Opto M3 monitoring system with IR beams spaced 0.5 inch apart and Oxymax data-acquisition software (Columbus Instruments). Total activity (the total number of beam breaks along the X and Y axis) was measured in 10 s intervals. Locomotor activity profiles were generated from the 2^nd^ day of LD or the 1^st^ day of DD.

#### Wheel-running activity

Adult male mice (3-5 months) were placed into individual cages with a vertical running wheel (ActiMetrics) and *ad libitum* access to food and water. After 1 week of acclimatization in wheel-cages under 12:12 hr LD conditions, 2 weeks of LD wheel-running activity were recorded, followed by 2 weeks of free-running activity in DD. Wheel-running data were acquired in 1 min bins and analyzed using ClockLab software (ActiMetrics). Period estimates were calculated using data from 12 days of DD.

#### Open Field Test (OFT)

OFT were conducted in 9 x 11 inch polyethylene cages using the Opto M3 beam-break setup described above. Auto-Track software (Columbus Instruments) was used to record the X-Y position, distance, and speed of each mouse at 10 Hz frequency. During the 3 hr test, animals were in constant darkness without bedding, food, or water. Each animal was placed in the arena at the indicated time point, and total activity and distance traveled were summed in 5 min bins as readouts. Behavioural data were analyzed using a custom MATLAB (Mathworks) program. Habituation was calculated by summing the total distance of the first 30 min of the trial and comparing it to the total distance of the final 30 min of each trial.

#### Acoustic Startle

Acoustic startle response was recorded using an SR-LAB Startle Response System (San Diego Instruments) apparatus, which consists of a sound-isolating cabinet containing a pressure-sensitive plate. Mice were placed into a plexiglass tube (I.D. 5 cm) and then enclosed inside the chamber on the pressure-recording plate for the duration of the trial. Mice were acclimated to the test environment, including 50 dB of background white noise, for 5 min before trials began. Each trial consisted of a 20 ms white noise stimulus (100 dB, 110 dB, or 120 dB) presented from a speaker 20 cm above the mouse’s head. The response of the animal in the 100 ms afterwards was recorded as vibration intensity on the pressure platform (in millivolts, mV); Vavg was the total activity averaged over the recorded window, while Vmax was the peak response intensity. All three trial tones were repeated 5 times throughout the experiment, in a pseudorandom order and separated by pseudorandom inter-trial intervals (13-17 s). Trials with significant vibration 100 ms before the tone were excluded from the analysis (<5 instances).

#### Behavioural assessment of reduced arousal

*mWake^(Cre/+)^;LSL-Gi* mice were assessed 90 min after IP injection of vehicle or 0.3 mg/kg CNO. Behaviour was scored by a blinded investigator, and each mouse classified as “Normal,” “Reduced Reactivity,” or “Stuporous,” based on the individual’s spontaneous behaviour and response to gentle handling. “Normal” mice spontaneously explored the environment and briskly responded to stimulation. “Reduced Reactivity” mice were generally immobile, but would try to evade handling. “Stuporous” mice did not exhibit spontaneous locomotion, and had a minimal response to handling. All animals exhibited righting reflex.

### Electroencephalography (EEG)

#### Surgery

8-10 week old male mice were anesthetized to surgical depth with a ketamine/xylazine mixture (100 mg/kg and 10 mg/kg, respectively), and all fur was removed from the top of the head. A skin incision was made along the top of the skull in the rostral-caudal direction, and the scalp was cleaned and connective tissue removed. The 3-channel EEG headmount (Pinnacle Technology) was aligned with the front 3 mm anterior to the bregma and glued to the top of the skull. Four guideholes were hand-drilled, and screws inserted to attach the headmount. EMG wires were then inserted into the left and right neck muscles. After skin closing, the headmount was sealed to the skull using dental cement. All animals recovered from surgery for > 5 days before being affixed to the EEG recording rigs.

#### EEG Recording

Sleep behavioural data were obtained using the Pinnacle Technology EEG/EMG tethered recording system. Following recovery, animals were placed into an 8 in diameter round acrylic cage with lid, provided *ad libitum* food and water, and tethered to a 100x preamplifier. All mice were housed in a 12:12 hr LD cycle and acclimated to the cable tethering for >5 days prior to recording. EEG and EMG channels were sampled at 400 Hz, high-pass filtered at 0.5 Hz for EEG and 10 Hz for EMG, digitized, and then acquired using Sirena software (Pinnacle Technology).

#### Analysis

Mouse sleep was scored visually by one or two trained technicians in Sirenia Sleep software (Pinnacle Technology), using raw EEG/EMG traces in 10 s epochs. Each epoch was scored WAKE, NREM, or REM, and epochs with artifacts were marked for exclusion in further analysis. Animals with severe movement artifacts or poor EEG waveforms were excluded from all behavioural datasets. Sleep or wake bouts were identified as >30 s of continuous sleep or wakefulness, respectively. Spectral analysis was performed using custom MATLAB (Mathworks) programs, and all Fast Fourier Transform spectra used 1024 or 512 point size and the Welch’s power spectral density estimate. Spectrograms were composed with short-time Fourier transforms with a window size of 30 s, 60% overlap, and smoothened by a rolling Hann window.

### Stereotaxic surgeries

8-12 week old male mice were anesthetized to surgical depth with a ketamine/xylazine mixture (100 mg/kg and 10 mg/kg, respectively), and all fur was removed from the top of the head. The mouse was secured into a Stoelting stereotaxis frame, and the microinjector tip was placed on the Bregma and all coordinates zeroed. Small (~0.5 mm) craniotomies were performed to allow for virus injection (50-300 nl at ~25 nl/min). Coordinates, volumes, viruses used and their sources are listed in Supplementary Table 2. Post-injection, animals were allowed to heal and express viral genes for ≥ two weeks (for projection and connectivity studies), and ≥ four weeks (for all behaviour and functional manipulations). If sleep behaviour was measured, EEG headmounts were implanted in a separate surgery. Locations of all viral injections were confirmed by post-hoc immunostaining, and no animals were excluded from our analyses.

### Electrophysiological recordings

Male mice between 5-10 weeks old were deeply anesthetized with isoflurane, and the brains quickly removed and dissected in oxygenated (95% O_2_, 5% CO_2_) ice-cold slicing solution (2.5 mM KCl, 1.25 mM NaH_2_PO_4_, 2 mM MgSO_4_, 2.5 mM CaCl_2_, 248 mM sucrose, 26 mM NaHCO_3_ and 10 mM glucose). Acute coronal brain slices (250 μm) were prepared using a vibratome (VT-1200s, Leica) and then incubated in oxygenated artificial cerebrospinal fluid (ACSF, 124 mM NaCl, 2.5 mM KCl, 1.25 mM NaH_2_PO_4_, 2 mM MgSO_4_, 2.5 mM CaCl_2_, 26 mM NaHCO_3_ and 10 mM glucose, 290-300 mOsm) at 28°C for 30 min and then at room temperature for 1 hr. Slices were then transferred to a recording chamber, continuously perfused with oxygenated ACSF at room temperature and visualized using an upright microscope (BX51WI, Olympus). Labeled cells of interest were visualized using infrared differential interference contrast (IR-DIC) and native fluorescence. Glass electrodes (5-8 MΩ) were filled with the following internal solution (130 mM K-gluconate, 5 mM NaCl, 10 mM C_4_H_8_O_5_PNa_2_, 1 mM MgCl_2_, 0.2 mM EGTA, 10 mM HEPES, 2 mM MgATP and 0.5 mM Na2GTP, pH 7.2-7.3, 300 mOsm). Whole-cell patch clamp recordings were obtained using a Multiclamp 700B amplifier (Molecular Devices). Data were sampled at 20 kHz, low-pass filtered at 2 kHz, and digitized using a Digidata 1440A (Molecular Devices).

For baseline spontaneous and evoked firing rate measurements, recordings were performed under current clamp configuration. Baseline recordings were performed for at least 30 s to measure spontaneous firing rate. To measure evoked firing rate, current injections from −10 to 100 pA were performed for 600 ms. 0.5% biocytin (wt/wt) was added to the internal solution to label the recorded cell. Slices were fixed post-experiment in 4% PFA overnight, and then incubated with Alexa488-conjugated streptavidin (Invitrogen, 1:2000) for 24 hrs, and then imaged on a Zeiss 800 confocal microscope. For measurement of GABA currents, voltage-clamp recordings at −70 mV were performed at ZT12-14. K-gluconate in the patch-pipette was replaced with CeCl_2_ for these recordings, CeOH was used to adjust the pH of the internal solution, and TTX (1 μM) was added to the ACSF to block action potentials. 1 mM GABA was delivered for 5 s using a Picospritzer III (Parker), and GABA-evoked current was recorded. For recordings of *mWake^DMH^* neurons, ~30% cells exhibited spontaneous firing, and so analyses of evoked responses were restricted to this subset of neurons.

For application of norepinephrine, orexin, acetylcholine, and histamine, current-clamp recordings were performed at ZT12-14. Baseline spontaneous activity was recorded for 30 s, and then for another 30 s following continuous application of the neurotransmitter. Compounds were pre-loaded into pulled glass pipettes (3-4 μm I.D. at the tip) and delivered at the recorded cell using a Picospritzer III (Parker). The compounds and concentrations used were as follows: norepinephrine (100 μM), acetylcholine chloride (1 mM), orexin-A (300 nm), and histamine (20 μM). For synaptically isolating recorded cells, ion channel blockers (20 μM CNQX; 50 μM AP5; 10 μM Picrotoxin) were added to the ACSF perfusion. For quantification of firing rates for cells reaching plateau potential, the maximum firing rate observed for any cell following application of the relevant neurotransmitter was used.

For optogenetic experiments, AAV-DIO-hChR2 (Supplementary Table 2) was injected into the DMH of *mWake^(Cre/+)^;TH-GFP* mice. Acute slices were prepared 3 weeks post-viral injection. Current-clamp and voltage-clamp recordings were performed from norepinephrine-expressing cells in the locus coeruleus (*TH^+^*), in the presence of optogenetic activation of ChR2-expressing terminals in the LC. Slices were exposed to blue light (480 nm) from the upper lens for 2 s at different stimulation frequencies (5, 10, 20, and 50 Hz) triggered by the Digidata 1440A.

### Immunohistochemistry

Mice were deeply anesthetized with a ketamine/xylazine mixture then fixed by transcardial perfusion with 4% paraformaldehyde (PFA). Brains were subsequently drop-fixed in 4% PFA for 24-48 hr and transferred into 1x PBS before being sectioned at 40 μm thickness using a vibratome (VT1200S, Leica). Free-floating sections were washed in 1x PBS, blocked for 1 hr in blocking buffer (PBS containing 0.25% Triton-X-100 and 5% normal goat serum or normal donkey serum), then incubated with rabbit anti-HDC (Progen, 16045, 1:800), goat anti-ChAT (Millipore, AB144P, 1:250), rabbit anti-orexin A (Abcam, AB6214, 1:500), chicken anti-TH (Abcam, AB76442, 1:1000), or rat anti-HA (Roche 11867423001, 1:250) in blocking buffer at 4°C overnight. The following day, slices were washed with PBST (PBS and 0.1% Tween-20), then incubated with Alexa 488 anti-rabbit (ThermoFisher, A-11008, 1:2500-1:5000), Alexa 568 anti rabbit (ThermoFisher, A-11011, 1:2500), Alexa 568 anti-goat (ThermoFisher, A11057, 1:2000), Alexa 488 anti-rat (ThermoFisher A-11006, 1:2000), or Alexa 488 anti-chicken (ThermoFisher, A-11039, 1:2000) secondary antibodies for 2.5 hrs in blocking buffer. Brain sections were then washed in PBST, incubated in DAPI (1:2000, Millipore) for 5 min, then washed in PBS. Sections were mounted on slides using VECTASHIELD HardSet Mounting Medium (Vector Laboratories, USA). In all tdTomato images, native fluorescence of tdTomato was visualized. Images were acquired using a Zeiss LSM800 confocal microscope under 10x-63x magnification.

### *In situ* hybridization

ISH for *mWake* was performed as previously described^4^. RNAScope ISH was performed using the RNAscope 2.5 Chromogenic Assay and the BaseScope™ Detection Reagent Kit according to the manufacturer’s instructions (Advanced Cell Diagnostics (ACD))^44^. Target probes were designed to exon 4 of *mWake*. Mouse brains were dissected and fresh frozen in Tissue-Tek O.C.T. (VWR) and cryosectioned at 10 μm. Sections were treated with hydrogen peroxide and protease, before hybridization to the custom *mWake* probe and subsequent amplification. Signal was detected by chromogenic reaction with BaseScope™ Fast RED, and sections counterstained with hematoxylin. Images were acquired on a Keyence BZ-X700 microscope (Keyence) under 10x brightfield illumination. Fluorescent *in situ* hybridization (FISH) was performed with the RNAscope™ Fluorescent Multiplex Assay (ACD), using probes targeting *tdTomato* and *Slc32a1 (VGat*) or *Slc17a6 (VGlut2*) mRNA. Preparation of tissue sections was performed as above, followed by simultaneous hybridization to both probes. Probe binding was indicated by deposition of target-specific fluorophores at each location via TSA Plus Fluorescence kit (PerkinElmer), and sections were then counterstained with DAPI. Sections were imaged on a Zeiss LSM880 confocal at 20x and 63x.

### Designer Receptors Exclusively Activated by Designer Drugs (DREADDs)

DREADD receptors coupled to either Gq or Gi were expressed in a Cre-dependent fashion in *mWake^+^* neurons of *mWake^(Cre/+)^* mice, either locally in the DMH, via stereotaxic injection of a viral vector (AAV-DIO-DREADD-Gq or AAV-DIO-DREADD-Gi) (Supplementary Table 2), or globally via crossing with a transgenic effector mouse line (*B6N;129-CAG-LSL-HA-hM4Di-mCitrine*, “LSL-Gi”). Clozapine *N*-oxide (CNO) (SigmaAldrich) was prepared as a stock solution of 50 mg/ml in DMSO, and then freshly diluted to 0.1 mg/ml in sterile PBS before IP injection. Solution clarity was monitored throughout dosing, and the solution was warmed to 37°C if precipitates were observed. Vehicle control was prepared as sterile saline + 0.01% DMSO. All injections occurred at the same ZT/CT time within each experiment, and all animals were treated with vehicle or CNO each day in a cross-over design, with ≥2 days recovery between experimental recording days. To control for CNO activity on its own, 1 or 3 mg/kg CNO were IP injected into sham-injected *mWake^(Cre/+)^* mice and locomotion and EEG data were assessed.

### Statistical analysis

Statistical analyses were performed in Prism 7 and 8 (Graphpad). For comparisons of two groups of normally distributed data, unpaired Student t-tests were used; if these comparisons were before and after treatment of the same animals or cells, paired t-tests were used instead. For comparisons of two groups of non-normally distributed data, Mann Whitney U tests were performed, with a Holm-Bonferroni correction, if required for multiple comparisons. For multiple comparisons of normally distributed data with 2 factors, 2 way ANOVAs were performed (with repeated measures, if applicable), followed by post-hoc Sidak tests. For multiple comparisons of non-normally distributed data, Kruskall-Wallis tests were performed with post-hoc Dunn’s tests.

## Acknowledgements

We thank A. Meredith and B. McNally for advice on SCN patch-clamp recordings and S. Hattar for assistance with EEG recordings. We thank B. Hamilton, A. Jackson, A. McCallion, and Y. Furuta for sharing mouse strains. We thank the Transcriptomics and Deep Sequencing Core for scRNA sequencing and the Ross Flow Cytometry Core flow sorting. We thank members of the Wu Lab for discussion. This work was supported by NIH grants R01NS094571 and R01NS079584. (M.N.W.) and a NINDS Center grant NS05027 for machine shop work.

## Author Contributions

B.J.B. and M.N.W. conceived the project, with input from J.Y.C. and S.B. B.J.B. performed most or all mouse genetics, behavioural experiments, EEG recordings, expression analyses, viral injections, and rabies tracing. Q.L.* performed all electrophysiology and optogenetic experiments and assisted with EEG analyses. D.W.K. performed scRNA-Seq analysis. S.S.L. performed viral injections and immunostaining. Q.L. assisted with mouse genetics and generation of the *mWake^-^* allele. I.D.B. assisted with EEG analyses. A.A.W. and J.L.B. assisted with expression analyses. A.J.C. assisted with viral injections. H.I. assisted with mouse genetics. B.J.B. and M.N.W. wrote the manuscript, with feedback from all authors.

---

**Extended Data Figure 1.**
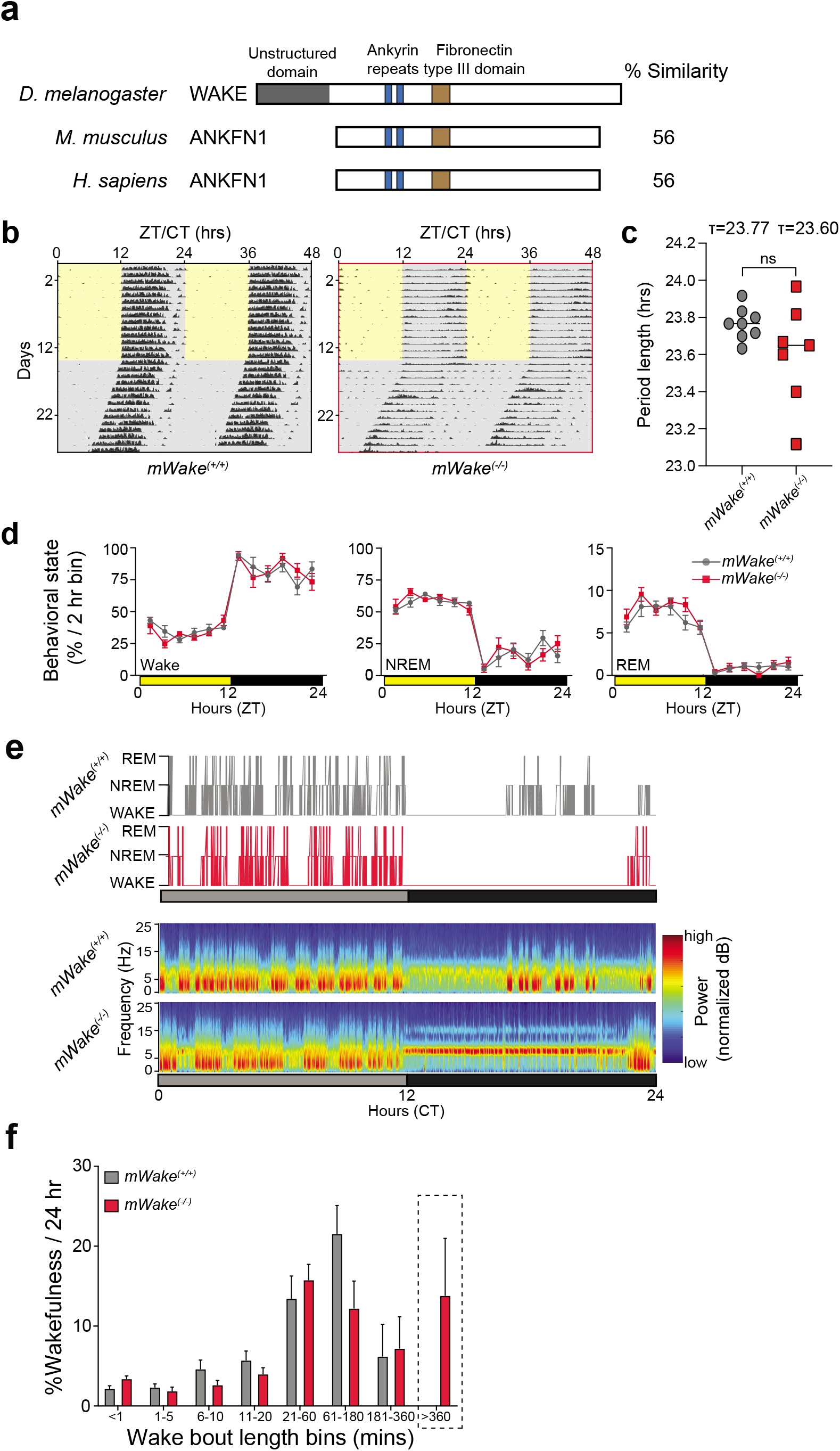
Circadian and EEG-related phenotypes of *mWake* mutant mice. **a**, Schematic showing domain structure of mWAKE in mice and humans, compared to *Drosophila* WAKE. Percentage similarity of mouse and human mWAKE, compared to fly WAKE is shown. **b**, Representative double-plotted actograms of wheel running activity for *mWake^(+/+)^* (left) vs *mWake^(-/-)^* mice (right), covering 14 days of LD then 14 days of free-running in DD. Activity plotted as number of wheel revolutions in 10 min bins. **c**, Locomotor period length (t) during the DD period for *mWake^(+/+)^* (n=8) and *mWake^(-/-)^* (n=8) mice. **d**, Vigilance state (% per 2 hr bin) determined by EEG recordings for *mWake^(-/-)^* (n=8) vs WT littermate control (n=6) mice under LD conditions. The mice in panel (**d**) are the same as in Fig. 1e. **e**, Hypnograms (*top*) and short-time Fourier transform spectrograms (*bottom*) over 24 hrs in DD showing example of prolonged wake bout in *mWake^(-/-)^* mice, compared to *mWake^(+/+)^* control. Power density is represented by the color-scheme and deconvoluted by frequency on the y-axis, over time on the x-axis. **f**, Wakefulness bout length distribution over a 24 hr period expressed as a % of total wakefulness, for *mWake^(+/+)^* vs *mWake^(-/-)^* mice. Dashed line highlights the presence of >6 hr continuous bouts of wakefulness, which are observed in some *mWake^(-/-)^* mutants and never in controls.

**Extended Data Figure 2.**
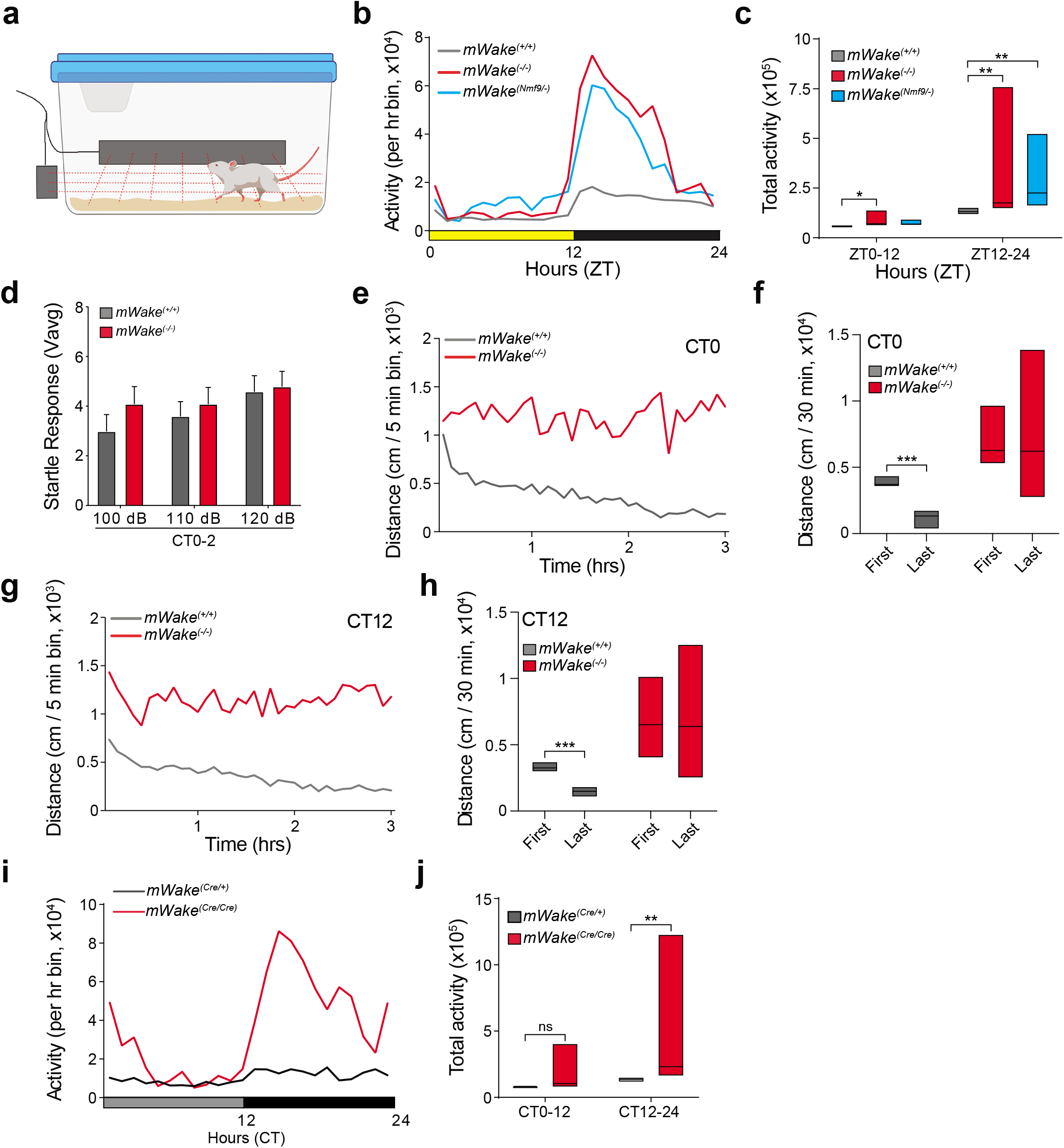
*mWake* mutants are hyperactive at night. **a**, Schematic of the beam-break arena used to measure locomotion, with infrared red beams forming a grid spaced at 0.5 inch intervals across the floor of the cage. **b**, Profile of locomotor activity (defined by beam breaks) over 24 hrs for *mWake^(+/+)^* (n=19, grey), *mWake^(-/-)^* (n=19, red), and *mWake^(Nmf9/-)^* (n=10, cyan) mice under LD conditions. **c**, Total locomotor activity (total number of beams broken along X and Y axes) from the mice in (**b**) at CT0-12 and CT12-24. Note that these data are from the same mice as in Fig. 1f. **d**, Startle response (Vavg) following a 100, 110, or 120 dB tone for *mWake^(+/+)^* (grey, n=10) vs *mWake^(-/-)^* mice (red, n=10) at CT0-2. Note that these data are from the same mice as in Fig. 1h. **e** and **g**, Distance (cm) traveled in five min bins in an open-field test, across 3 hrs of the trial (CT0-CT3, **e**) or (CT12-CT15, **g**) for *mWake^(+/+)^* (n=8, grey) vs *mWake^(-/-)^* (n=8, red) mice. Data were collected from 3 independent trials. **f** and **h**, Habituation of animals shown in (**e**) and (**g**), respectively, as assessed by total distance (cm) traveled in the first 30 mins vs the last 30 mins of the trial. **i**, Locomotor activity profile over 24 hrs for *mWake^(Cre/+)^* (n=9, grey) and *mWake^(Cre/Cre)^* (n=10, red) under DD conditions. **j**, Total locomotor activity (total number of beams broken along X and Y axes) from the mice in (**i**) at CT0-12 vs CT12-24.

**Extended Data Figure 3.**
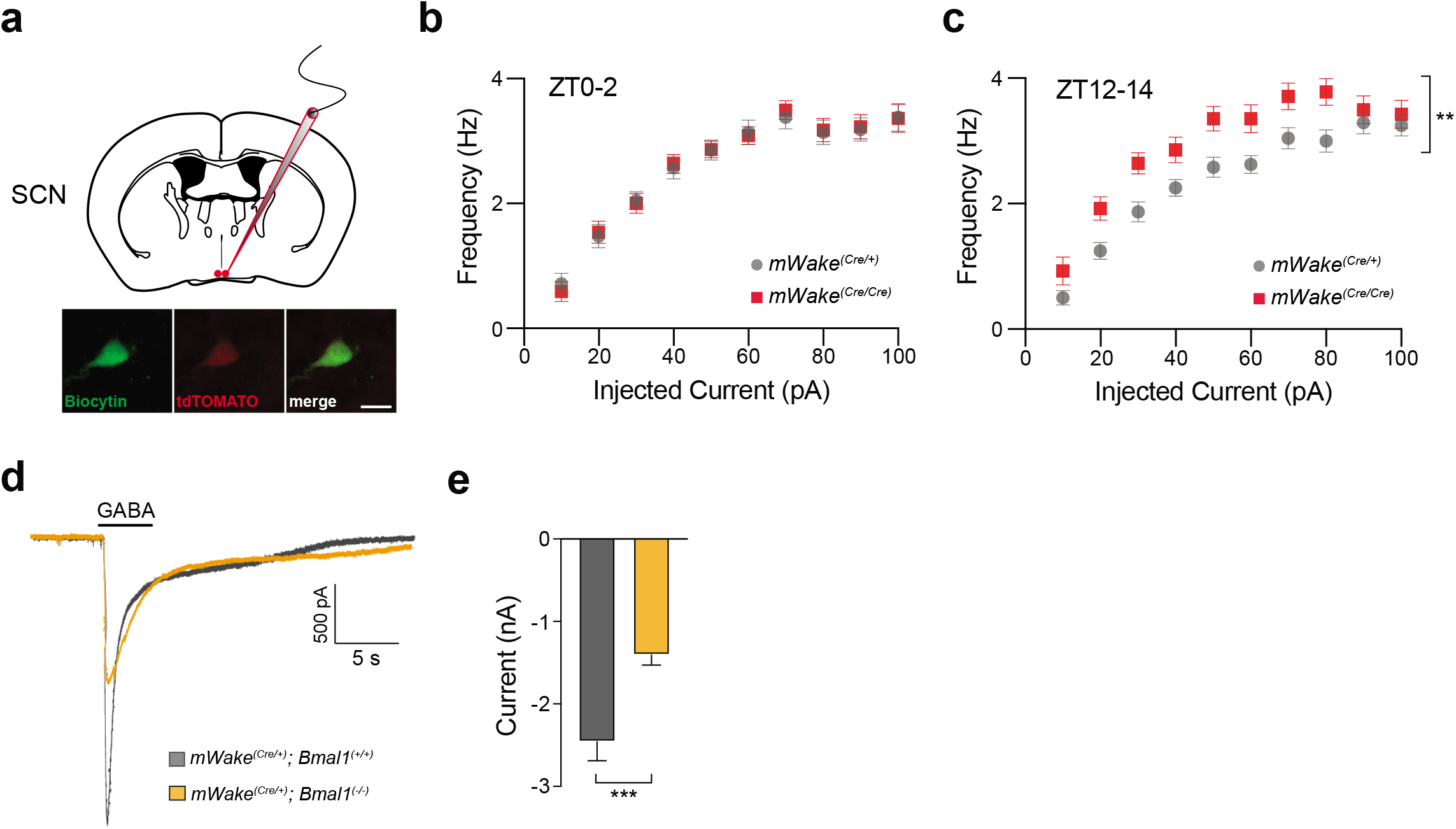
mWAKE inhibits the activity of SCN neurons at night. **a**, Schematic of whole-cell patch-clamp recordings from *mWake*^+^ neurons (tdTomato^+^ neurons in *mWake^(Cre/+)^* or *mWake^(Cre/Cre)^* mice), in the core region of the SCN (top). Representative images of a biocytin- and tdTomato-labeled cell post-recording (bottom). Scale bar denotes 25 μm. **b** and **c**, *f*-*I* curves for *mWake*^+^ SCN neurons from *mWake^(Cre/+)^* vs *mWake^(Cre/Cre)^* mice at ZT0-2 (n=21 and 19) (**b**) or ZT12-14 (n=19 and 13) (**c**). **d**, Representative traces of voltage-clamp recordings of *mWake^+^* SCN neurons of *mWake^(Cre/+)^; Bmal1^(+/+)^* (grey) vs *mWake^(Cre/+)^; Bmal1^(-/-)^* (yellow) at ZT12-14. Timing of GABA (1 mM) application is shown. **e**, GABA-evoked current in *mWake^SCN^* neurons from *mWake^(Cre/+)^; Bmal1^(+/+)^* (n=24) or *mWake^(Cre/+)^; Bmal1^(-/-)^* (n=26) mice at ZT12-14. n represents individual cells from 4 animals.

**Extended Data Figure 4.**
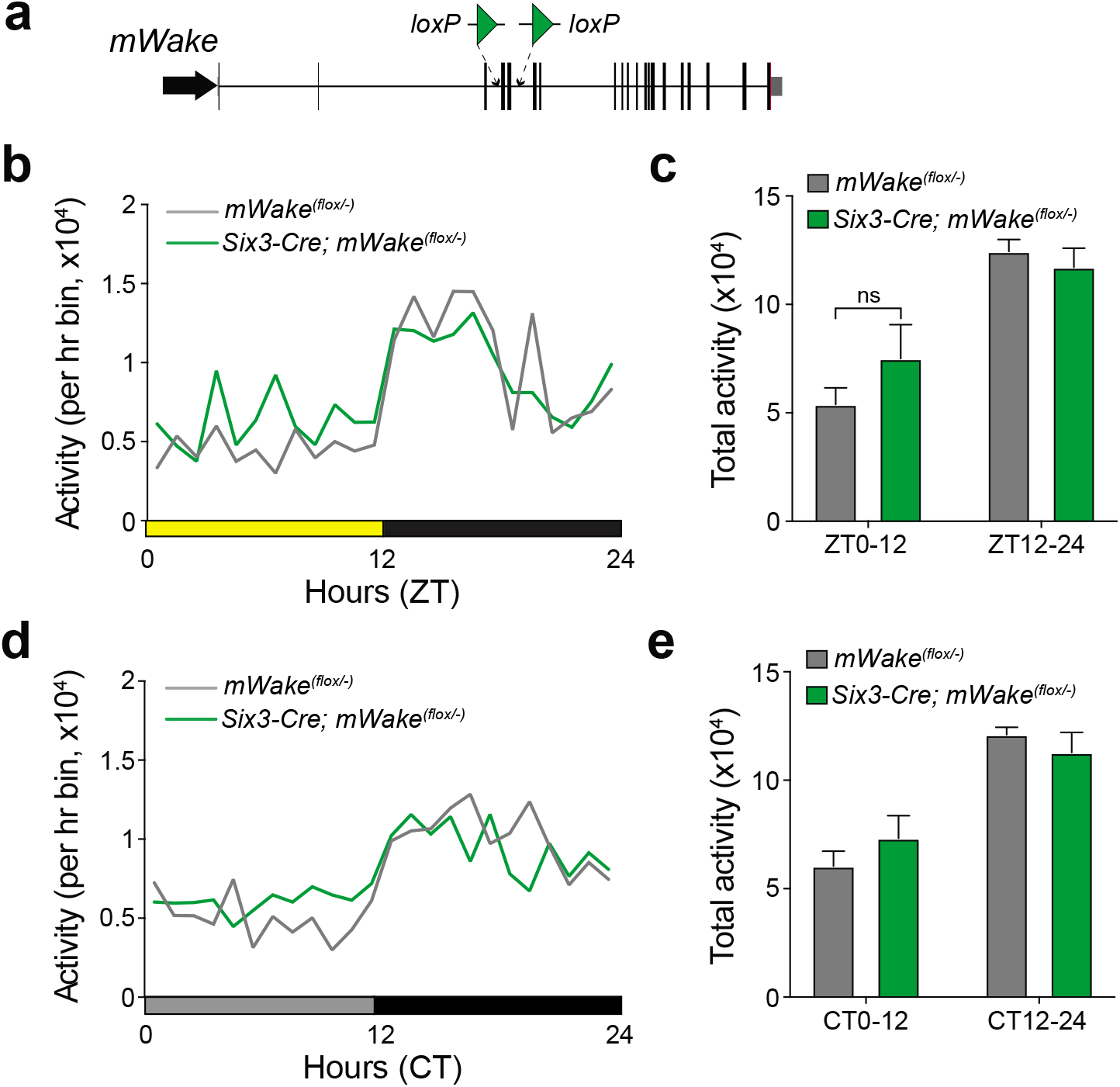
Loss of mWAKE in SCN does not cause hyperactivity. **a**, Schematic of the genomic structure of the *mWake* locus and insertion of *loxP* sites flanking exons 4 and 5 in the *mWake^(flox)^* allele. **b** and **d**, Locomotor activity profile over 24 hrs for *mWake^(flox/-)^* (n=5, grey) and *Six3^(Cre/+)^; mWake^(flox/-)^* (n=6, green) under LD (**b**) or DD (**d**) conditions. **c** and **e**, Total locomotor activity from the mice in (**b**) at ZT0-12 and ZT12-24 or (**d**) at CT0-12 and CT12-24.

**Extended Data Figure 5.**
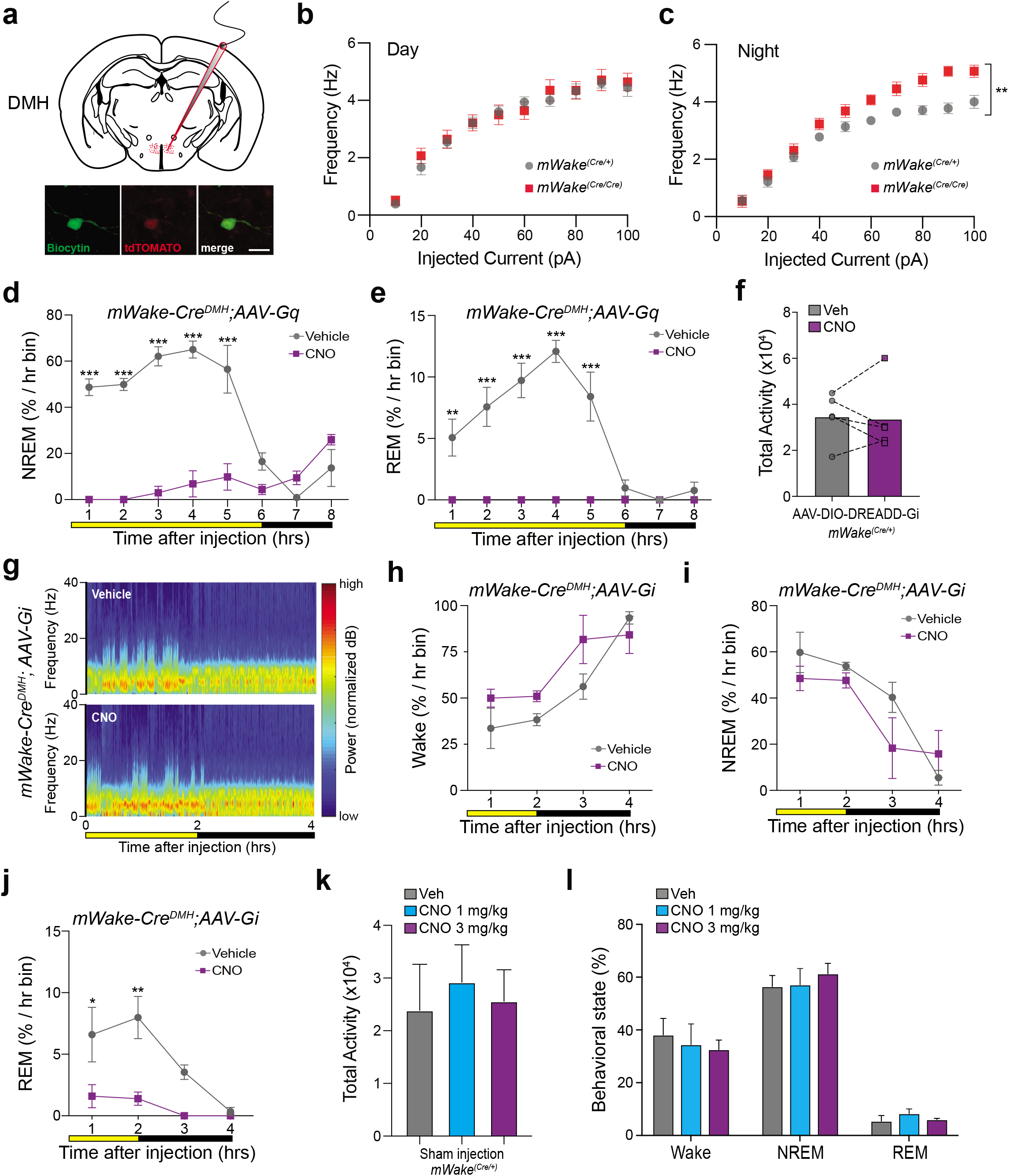
*mWake*^+^ DMH neurons promote arousal. **a**, Schematic of whole-cell patch-clamp recordings from *mWake^+^* neurons (tdTomato^+^ neurons in *mWake^(Cre/+)^* or *mWake^(Cre/Cre)^* mice), in the DMH region (top). Representative images of a biocytin- and tdTomato-labeled cell post-recording (bottom). Scale bar denotes 25 μm. **b** and **c**, *f*-*I* curves for *mWake^DMH^* neurons from *mWake^(Cre/+)^* vs *mWake^(Cre/Cre)^* mice at ZT0-2 (n=11 and 11, Day) (**b**) or ZT12-14 (n=8 and 8, Night) (**c**). n represent individual cells from 4 animals. **d** and **e**, NREM (**d**) and REM sleep (**e**) (% per hr bin) for *mWake^(Cre/+)^* mice with AAV-DIO-DREADD-Gq injected bilaterally into the DMH, following IP injection of vehicle (grey) or 1 mg/kg CNO (magenta) at ZT6. Note these are data are from the same mice as in Fig. 2l. **f**, Total locomotor activity (total number of beams broken along X and Y axes) of *mWake^(Cre/+)^* mice with bilateral injections of AAV-DIO-DREADD-Gi into the DMH in the 4 hrs following IP injection of vehicle (grey) vs CNO (1 mg/kg, magenta) (n=4) at ZT10. **g**, Representative short-time Fourier transform spectrograms of 4 hrs of recorded EEG activity, starting after IP injection of vehicle alone (above) or 3 mg/kg CNO (below) at ZT10, from *mWake^(Cre/+)^* mice injected with AAV-DIO-DREADD-Gi bilaterally into the DMH. **h**-**j**, Wakefulness (**h**), NREM (**i**), and REM (**j**) amount for the mice described in (**g**), plotted as % time in 1 hr bins following IP injection of vehicle (grey) or 3 mg/kg CNO (magenta) (n=4) at ZT10. **k**, total locomotor activity (total number of beams broken along X and Y axes) of *mWake^(Cre/+)^* mice with sham injections into the DMH in the 4 hrs following IP injection of vehicle (grey), CNO (1 mg/kg, cyan), or CNO (3 mg/kg, magenta) (n=3) at ZT6. **l**, Vigilance state (%) of 2 hrs of recorded EEG activity, starting after IP injection of vehicle alone, 1 mg/kg CNO, or 3 mg/kg CNO at ZT6, from the mice described in (**k**).

**Extended Data Figure 6.**
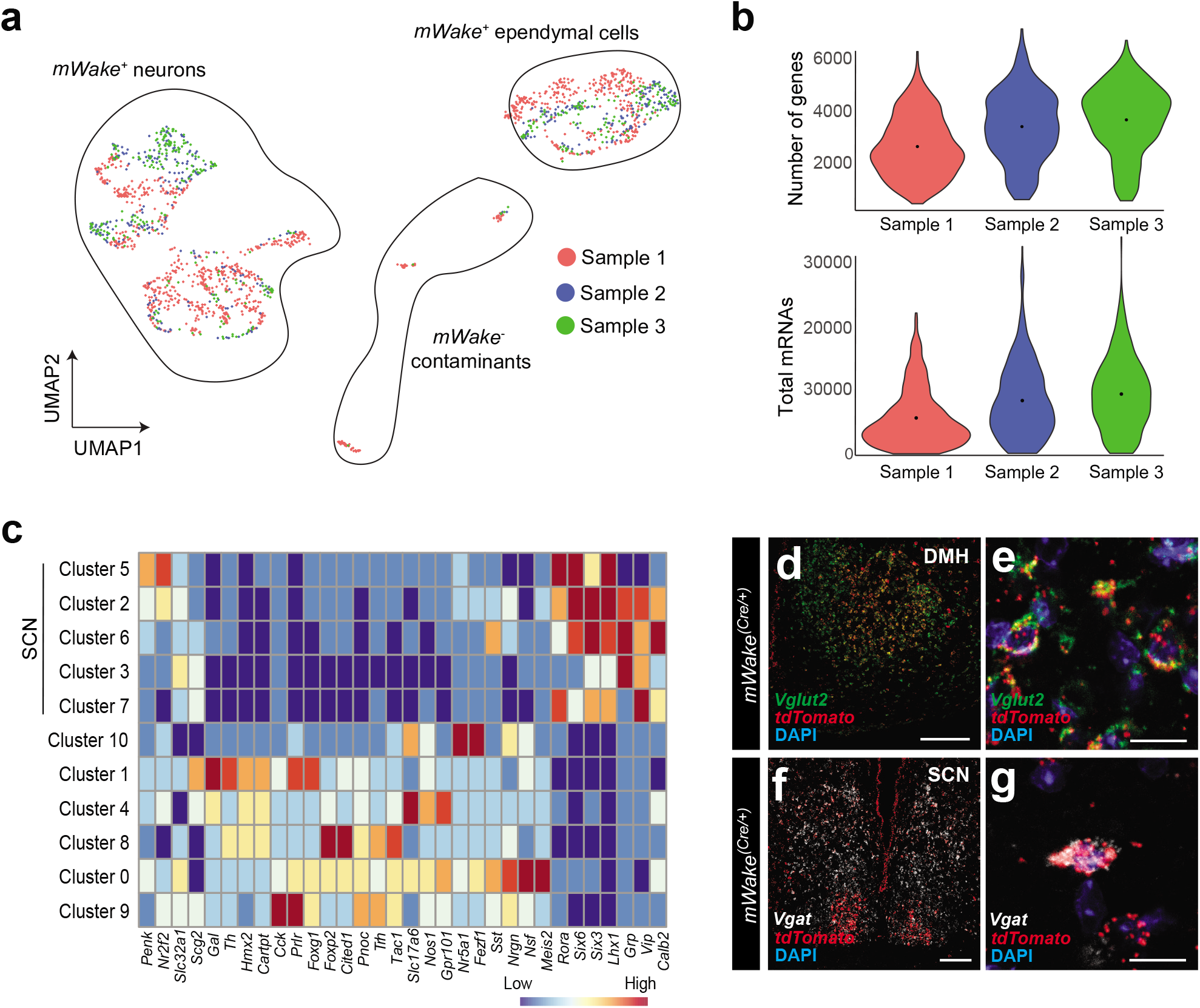
Additional data related to single cell RNA sequencing. **a**, UMAP plot showing distribution of *mWake*^+^ cells in individual scRNA-Seq libraries and distribution of *mWake*^+^ neurons and ependymal cells. **b**, Violin plot showing distribution of number and mean (black dot) of genes (top) and total mRNAs (bottom, calculated by the number of unique molecular identifiers (UMIs)) in individual scRNA-Seq libraries. **c**, Heatmap showing key marker genes that were used to identify spatial location of each *mWake*^+^ neuronal cluster. **d**-**g**, Representative images of FISH experiments using RNAscope probes against *tdTomato*, *Vglut2*, and/or *Vgat* mRNA in the DMH (**d** and **e**) and SCN (**f** and **g**) of *mWake^(Cre/+)^* mice. Nuclei counterstained with DAPI. Scale bars denote 200 μm in **d** and **f** and 20 μm in **e** and **g**.

**Extended Data Figure 7.**
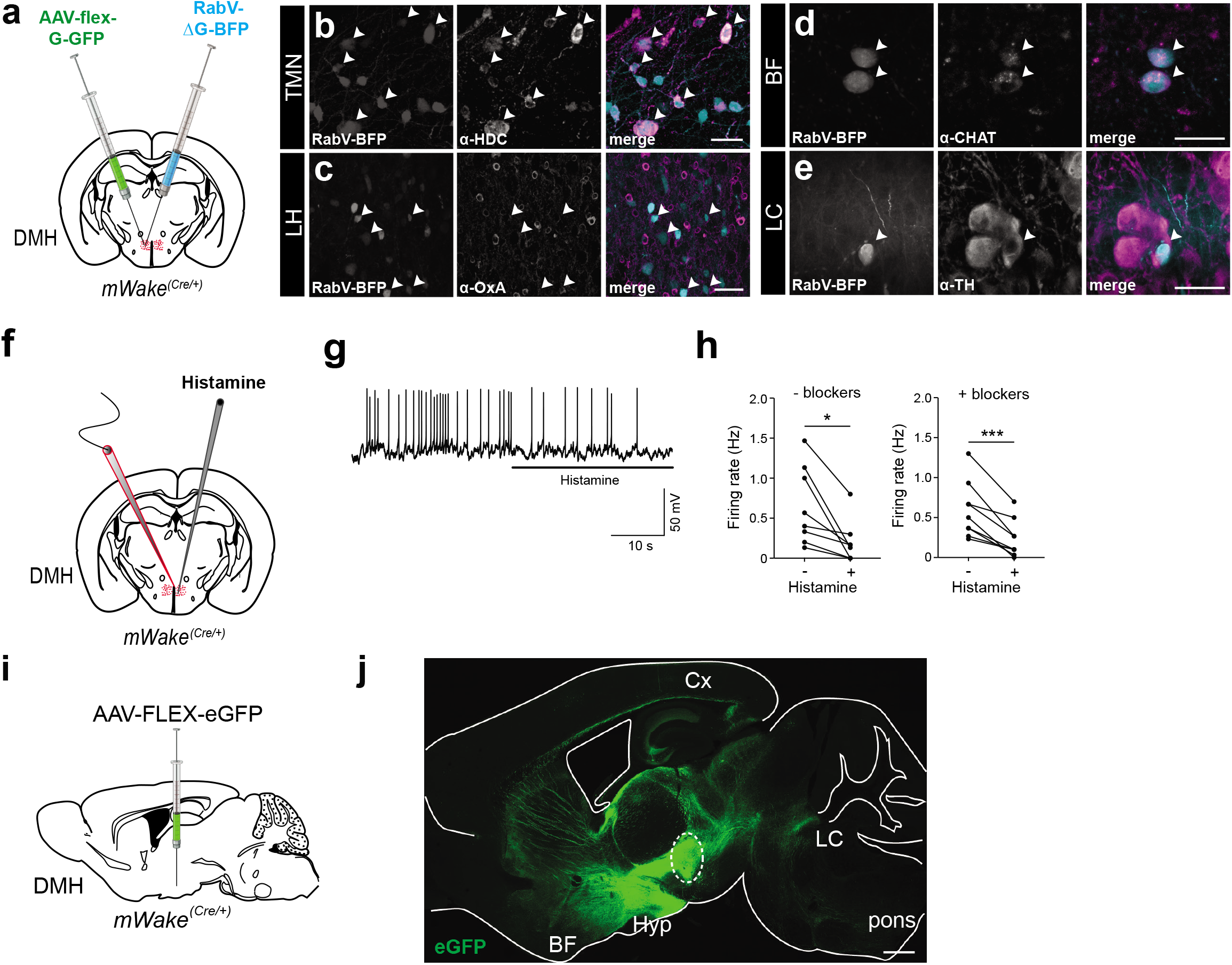
Additional data related to neuromodulatory input to *mWake^DMH^* neurons. **a**, Schematic showing dual injections of the Cre-dependent rabies helper virus (AAV-FLEX-G) and the rabies-ΔG-BFP (RabV-ΔG-BFP) virus into the DMH of *mWake^(Cre/+)^* mice. **b**-**e**, Representative images of BFP fluorescence and anti-histamine decarboxylase (HDC) (**b**), anti-Orexin A (OxA) (**c**), anti-choline acetyltransferase (CHAT) (**d**), or anti-tyrosine hydroxylase (TH) (**e**), as well as merged images from TMN (**b**), lateral hypothalamus (LH) (**c**), BF (**d**), or LC (**e**) regions, from the mice described in (**a**). Scale bar denotes 20 μm in (**b**), 50 μm in (**c**), 20 μm in (**d**), 20 μm in (**e**). **f**, Schematic showing patch-clamp recordings of *mWake^DMH^* neurons from *mWake^(Cre/+)^* mice, following application of histamine. **g**, Representative membrane potential traces from the recordings described in (**f**) in the presence of synaptic blockers at ZT12-14. **h**, Spontaneous mean firing rate of *mWake^DMH^* neurons following application of 20 μM histamine (n=8 and 9 cells) at ZT12-14 in the absence (“-blockers”) or presence (“+ blockers”) of 20 μM CNQX, 50 μM AP5, and 10 μM picrotoxin. The neurons recorded from in (**h**) were derived from 3 animals. **i**, Schematic showing unilateral stereotaxic injection of AAV-viral vector expressing Cre-dependent eGFP (AAV-FLEX-eGFP) into the DMH of *mWake^(Cre/+)^* mice. **j**, Representative sagittal section showing broad projection of GFP fluorescence following injection of AAV-FLEX-eGFP shown in (**e**). Dashed ellipse denotes injection site, and locations of the basal forebrain (BF), cortex (Cx), hypothalamus (Hyp), locus coeruleus (LC), and pons are noted. Scale bar denotes 1 mm.

**Extended Data Figure 8.**
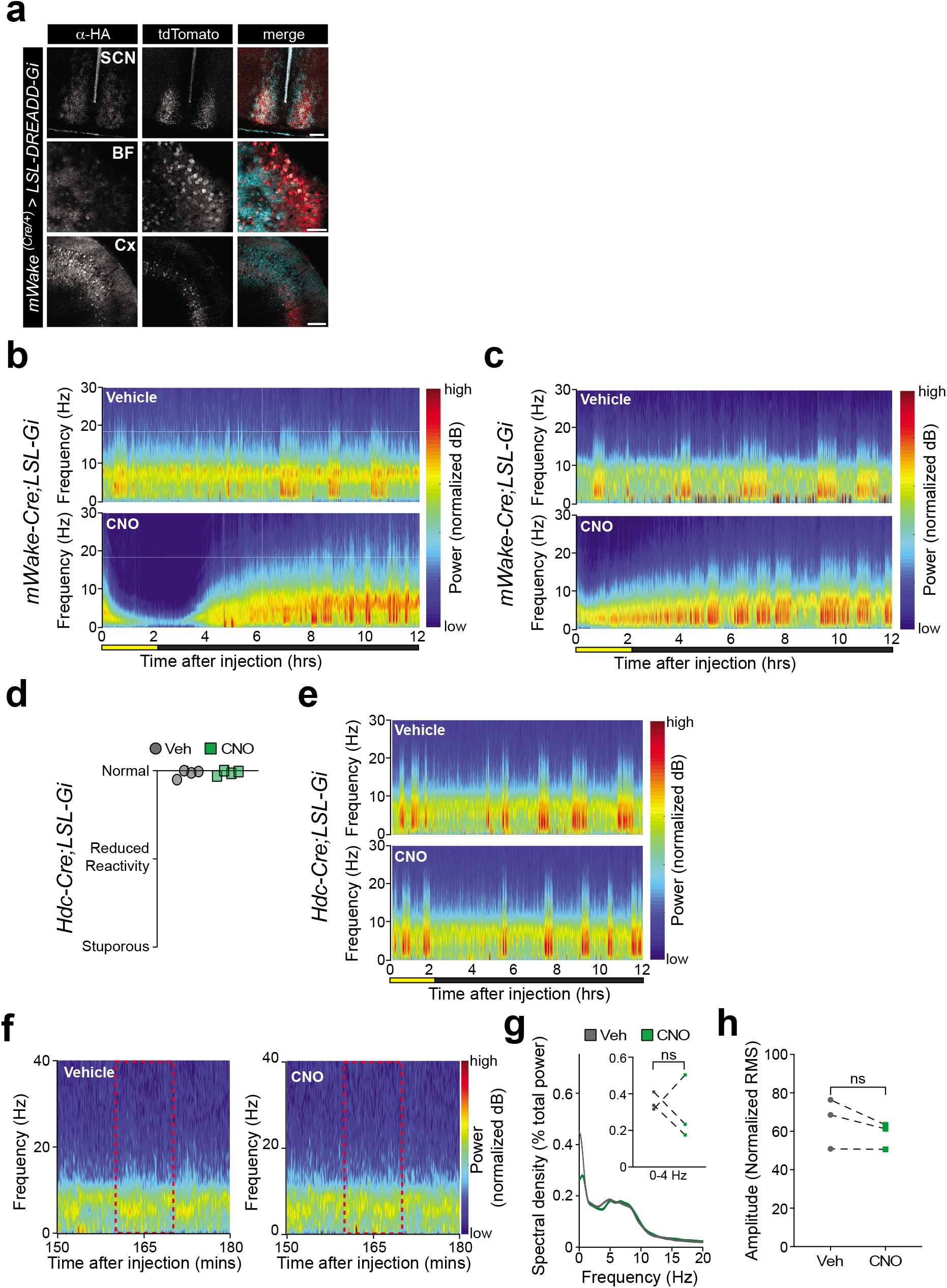
The mWAKE network is critical for arousal. **a**, Representative images of anti-HA labeling of cells and processes expressing LSL-DREADD-Gi, tdTomato fluorescence from *mWake*^+^ cells, or merged images in representative brain regions from *mWake^(Cre/+)^*; *LSL-DREADD-Gi* mice. Sections containing suprachiasmatic nucleus (SCN), basal forebrain (BF), and cortex (Cx) are shown. Scale bar denotes 100 μm for SCN, 20 μm for BF, and 100 μm for Cx panels. **b, c**, Two additional short-time Fourier transform spectrograms of 12 hrs of recorded EEG activity from *mWake^(Cre/+)^*; *LSL-Gi* mice after IP injection at ZT10 of vehicle alone (above) or 0.3 mg/kg CNO (below). These panels illustrate the range of EEG phenotypes observed. **d**, Jitter plot showing behavioural classification of arousal level for *HDC^(Cre/+)^; LSL-Gi* mice following injection of vehicle or 1 mg/kg CNO (n=4). **e**, Representative short-time Fourier transform spectrograms of 12 hrs of recorded EEG activity from *HDC^(Cre/+)^*; *LSL-Gi* mice after IP injection at ZT10 of vehicle alone (above) or 1 mg/kg CNO (below). **f**, Short-time Fourier transform spectrograms for a *HDC^(Cre/+)^;LSL-Gi* mouse starting 150 mins after injection of vehicle (left) or 1 mg/kg CNO (right) at ZT10. The dashed red box indicates the time window used for spectral and amplitude analysis in (**g**) and (**h**). **g**, Welch’s power spectral density estimates as a percentage of total EEG power across 10 min, averaged across multiple EEG traces. *HDC^(Cre/+)^;LSL-Gi* injected with vehicle (grey) or 1 mg/kg CNO (green) (n=3). Inset shows a plot of delta-band power as a percentage of total EEG power. **h**, EEG trace amplitude (plotted as normalized root mean square (RMS)) for the animals described in (**f**).

**Supplementary Table 1 | Additional electrophysiological properties for *mWake^SCN^* and *mWake^DMH^* neurons.**

**Supplementary Table 2 | Stereotaxic coordinates and viruses injected**

**Supplementary Table 3 | List of differentially expressed genes in *mWake^+^* neurons**

**Supplementary Video 1 | *mWake* KO exhibit nighttime hyperactivity and circling.**

**(0:00 – 0:15**), A single *mWake^(-/-)^* mouse exhibiting elevated homecage locomotor activity, compared to heterozygote littermates. (**0:15 – 0:19**), *mWake^(-/-)^* mouse demonstrating rapid (~1.6 Hz) tight circling behavior. (**0:20 – 0:31**), Two *mWake^(-/-)^* mice display broad circling activity (~1.4 Hz). Video speed is not adjusted.

**Supplementary Video 2 | *mWake^(-/-)^* mice are not impaired in the forced-swim assay.**

**(0:00 – 0:40**), Representative trial of forced swim assay for a control (“Wild Type”) mouse. (**0:40 – 1:22**), Representative trial of forced swim assay for a *mWake^(-/-)^* (“*mWake-KO*”) mouse. (**1:22 – 2:08**), Representative trial of forced swim assay of *mWake^(NMF9/-)^* (“*NMF9 imWake-KO*”) mouse. All animals were allowed to swim for > 30 s before being removed.

**Supplementary Video 3 | Chemogenetic inhibition of *mWake^+^* network induces profound reduction in arousal.**

**(0:00 – 0:35**), *mWake^(Cre/+)^;LSL-Gi* mouse (“Vehicle”) 90 min after injection of vehicle solution, classified as “Normal,” exhibiting normal spontaneous locomotion and exploration, as well as avoidance of paper object. (**0:36 – 1:10**), *mWake^(Cre/+)^;LSL-Gi* mouse (“CNO#1”) 90 min after injection of 0.3 mg/kg CNO solution, classified as “Reduced Reactivity,” displaying markedly decreased spontaneous locomotion, with intact righting reflex and reduced avoidance of paper object. (**1:11 – 1:40**), *mWake^(Cre/+)^;LSL-Gi* mouse (“CNO#2”) 90 min after injection of 0.3 mg/kg CNO solution, classified as “Stuporous,” with absent spontaneous locomotion, slow righting reflex, and minimal responsiveness to paper object.

