## Supplementary Table 1 for "A Clock-Driven Neural Network Critical for Arousal"

| Genotype and condition | MFR (Hz) | RMP (mV) | Rin (GΩ) | τ (ms) |
|---|---|---|---|---|
| *mWAKE*$^{Cre/+}$ SCN ZT0-2 | 1.1 ± 0.2 | -50.6 ± 1.6 | 1.8 ± 0.2 | 61.3 ± 1.0 |
| *mWAKE*$^{Cre/Cre}$ SCN ZT0-2 | 1.2 ± 0.2 (*P*=0.86) | -48.7 ± 1.3 (*P*=0.29) | 2.2 ± 0.2 (*P*=0.16) | 62.8 ± 1.4 (*P*=0.37) |
| *mWAKE*$^{Cre/+}$ SCN ZT12-14 | 0.4 ± 0.1 | -53.4 ± 1.6 | 1.6 ± 0.1 | 61.8 ± 1.5 |
| *mWAKE*$^{Cre/Cre}$ SCN ZT12-14 | 1.1 ± 0.1 (*P*<0.001) | -50.7 ± 1.6 (*P*=0.24) | 2.3 ± 0.3 (*P*=0.053) | 61.4 ± 1.1 (*P*=0.87) |
| *mWAKE*$^{Cre/+}$ DMH ZT0-2 | 0.5 ± 1.0 | -54.6 ± 3.2 | 1.4 ± 0.2 | 61.1 ± 2.1 |
| *mWAKE*$^{Cre/Cre}$ DMH ZT0-2 | 0.7 ± 0.2 (*P*=0.37) | -56.4 ± 3.8 (*P*=0.72) | 1.6 ± 0.1 (*P*=0.53) | 64.3 ± 0.2 (*P*=0.21) |
| *mWAKE*$^{Cre/+}$ DMH ZT12-14 | 1.2 ± 0.2 | -51.4 ± 1.5 | 1.6 ± 0.1 | 63.8 ± 0.3 |
| *mWAKE*$^{Cre/Cre}$ DMH ZT12-14 | 2.2 ± 0.5 (*P*<0.05) | -47.6 ± 2.2 (*P*=0.15) | 1.4 ± 0.2 (*P*=0.45) | 63.2 ± 0.4 (*P*=0.24) |

| Neuromodulator and condition | MFR (ACSF) (Hz) | RMP (ACSF) (mV) | MFR (blockers) (Hz) | RMP (blockers) (mV) |
|---|---|---|---|---|
| ACETYLCHOLINE (before) | 0.5 ± 0.3 | -55.0 ± 4.1 | 0.4 ± 0.2 | -56.4 ± 3.3 |
| ACETYLCHOLINE (during) | 0.8 ± 0.3 (*P*<0.05) | -53.3 ± 3.5 (*P*=0.17) | 2.7 ± 0.7 (*P*<0.01) | -44.2 ± 4.5 (*P*<0.01) |
| OREXIN (before) | 0.4 ± 0.2 | -58.2 ± 3.4 | 0.1 ± 0.08 | -60.7 ± 2.4 |
| OREXIN (during) | 0.4 ± 0.1 (*P*=0.87) | -58.2 ± 2.9 (*P*=0.99) | 1.2 ± 0.4 (*P*<0.05) | -53.2 ± 3.2 (*P*=0.07) |
| NOREPINEPHRINE (before) | 0.5 ± 0.1 | -55.3 ± 2.3 | 0.3 ± 0.1 | -57.4 ± 2.0 |
| NOREPINEPHRINE (during) | 1.94±0.3 (*P*<0.001) | -31.2 ± 2.9 (*P*<0.001) | 0.9 ± 0.4 (*P*=0.12) | -59.1 ± 2.4 (*P*=0.36) |
| HISTAMINE (before) | 0.7 ± 0.2 | -49.9 ± 4.2 | 0.6 ± 0.1 | -49.7 ± 2.1 |
| HISTAMINE (during) | 0.2 ± 0.1 (*P*<0.05) | -54.3 ± 5.1 (*P*<0.05) | 0.2 ± 0.1 (*P*<0.001) | -52.9 ± 3.1 (*P*=0.06) |

**Supplementary Table 1 | Additional electrophysiological properties for *mWake*$^{SCN}$ and *mWake*$^{DMH}$ neurons.**

(Top) Mean firing rate (MFR), resting membrane potential (RMP), input resistance (Rin), and membrane time constant (τ) are shown for the displayed genotypes and cells during ZT0-2 and ZT12-14. *P* values are shown for comparisons between ZT0-2 and ZT12-14 for a given genotype.

(Bottom) MFR and RMP shown for mWAKE$^{DMH}$ neurons before or during application of the given neuromodulator, in the absence (ACSF/artificial cerebrospinal fluid) or presence of synaptic blockers, as described in Methods. *P* values are shown for comparisons between before and during application of the neuromodulator.
