## Supplementary Table 2 for "A Clock-Driven Neural Network Critical for Arousal"

**Supplementary Table 2 | Stereotaxic coordinates and viruses injected**

| Location | Coordinates | Virus (Source) | Use | Vol. | Laterality |
|---|---|---|---|---|---|
| **DMH** | AP:-1.55, ML:+/- 0.25, DV:-5.55 | AAV9.CMV.HI.eGFP-Cre.WPRE.SV40 (Penn Vector) "AAV-Cre" | Conditional KO | 300 nl | Bi- |
| | AP:-1.55, ML:+/- 0.25, DV:-5.55 | AAV9.CMV.HI.eGFP-WPRE.SV40 (Penn Vector) "AAV-Sham" | Sham-Control | 300 nl | Bi- |
| | AP:-1.55, ML:+/- 0.25, DV:-5.55 | AAV1.pCAG.FLEX.eGFP-WPRE (Addgene# 51502) "AAV-FLEX-eGFP" | Projection mapping | 50 nl | Uni- |
| | AP:-1.55, ML:+/- 0.25, DV:-5.55 | AAV9.hSyn.DIO.hM3D(Gq)-mCherry (Addgene# 44361) "AAV-DIO-DREADD-Gq" | DREADD activation | 250 nl | Bi- |
| | AP:-1.55, ML:+/- 0.25, DV:-5.55 | AAV8.hSyn.DIO.hM4D(Gi)-mCherry (Addgene# 44362) "AAV-DIO-DREADD-Gi" | DREADD inhibition | 250 nl | Bi- |
| | AP:-1.55, ML:+/- 0.25, DV:-5.55 | AAV8.hSyn.FLEX.TVA.P2A.eGFP.oG (Salk Institute) "AAV-FLEX-G" | Rabies Helper-virus w/ G protein | 250 nl | Uni- |
| | AP:-1.55, ML:+/- 0.25, DV:-5.55 | Rabies.ΔG.BFP (Salk Institute) "RabV-ΔG-BFP" | Transynaptic labeling | 50 nl | Uni- |
| | AP:-1.55, ML:+/- 0.25, DV:-5.55 | AAV9.EF1α.DIO-hChR2(H134R).EYFP-pA (Addgene# 20297) "AAV-DIO-hChR2" | Optogenetic activation | 250 nl | Uni- |
