## Supplementary Table 3 for "A Clock-Driven Neural Network Critical for Arousal"

**Supplementary Table 3 | List of differentially expressed genes in *mWake*+ neurons**

| gene | avg_logFC | pct.1 | pct.2 | p_val_adj | cluster |
|---|---|---|---|---|---|
| Gal | 2.8545 | 0.921 | 0.26 | 7.99E-88 | 1 |
| Vip | 2.3005 | 0.897 | 0.405 | 3.89E-32 | 7 |
| Grp | 1.986 | 0.886 | 0.277 | 6.08E-46 | 3 |
| Nrgn | 1.8691 | 1 | 0.224 | 2.09E-07 | 10 |
| Nptx1 | 1.8491 | 1 | 0.185 | 1.33E-16 | 10 |
| Rgs16 | 1.8278 | 0.859 | 0.32 | 9.82E-42 | 7 |
| Cck | 1.8231 | 0.4 | 0.028 | 2.20E-06 | 9 |
| Thy1 | 1.7703 | 1 | 0.412 | 2.17E-11 | 10 |
| Slc17a6 | 1.7622 | 1 | 0.111 | 3.40E-09 | 10 |
| Ppp1r1a | 1.7463 | 0.628 | 0.391 | 2.16E-20 | 7 |
| Ncdn | 1.7191 | 1 | 0.613 | 3.44E-15 | 10 |
| Cacna2d1 | 1.6592 | 1 | 0.365 | 1.48E-13 | 10 |
| Fos | 1.4947 | 0.897 | 0.395 | 3.04E-26 | 7 |
| Rbfox1 | 1.4945 | 0.941 | 0.199 | 3.44E-10 | 10 |
| Fezf1 | 1.4804 | 0.824 | 0.006 | 1.78E-11 | 10 |
| Ppp3ca | 1.4538 | 1 | 0.79 | 1.51E-13 | 10 |
| Camkv | 1.4483 | 1 | 0.475 | 2.53E-13 | 10 |
| Lmo3 | 1.4159 | 0.882 | 0.142 | 6.12E-09 | 10 |
| Tac1 | 1.4139 | 0.493 | 0.065 | 3.70E-12 | 8 |
| Unc13c | 1.3972 | 0.915 | 0.031 | 5.62E-70 | 8 |
| Gabra1 | 1.3893 | 0.941 | 0.357 | 0.0076 | 10 |
| Gda | 1.3872 | 0.882 | 0.627 | 8.88E-05 | 10 |
| Brinp3 | 1.3742 | 0.941 | 0.405 | 1.50E-05 | 10 |
| Galnt16 | 1.3691 | 0.941 | 0.366 | 1.09E-07 | 10 |
| Cdh13 | 1.3638 | 0.882 | 0.308 | 1.95E-05 | 10 |
| Dusp1 | 1.3565 | 0.821 | 0.249 | 9.75E-32 | 7 |
| App | 1.3507 | 1 | 0.929 | 6.91E-13 | 10 |
| Rxrg | 1.329 | 0.901 | 0.095 | 1.67E-62 | 8 |
| Napb | 1.3135 | 1 | 0.465 | 3.10E-07 | 10 |
| Eno2 | 1.2697 | 0.941 | 0.713 | 9.43E-06 | 10 |
| Kcnq3 | 1.2671 | 0.882 | 0.162 | 4.22E-07 | 10 |
| Nfix | 1.2572 | 0.733 | 0.084 | 3.50E-14 | 9 |
| Cnr1 | 1.2496 | 0.882 | 0.199 | 1.53E-06 | 10 |
| Rgs4 | 1.248 | 0.789 | 0.173 | 3.67E-28 | 8 |
| Cartpt | 1.245 | 0.434 | 0.079 | 9.69E-20 | 1 |
| Jun | 1.2386 | 0.897 | 0.525 | 1.05E-23 | 7 |
| C1ql3 | 1.2306 | 0.436 | 0.146 | 1.08E-13 | 7 |
| Atp2b4 | 1.2303 | 1 | 0.261 | 4.51E-05 | 10 |
| Hpcal1 | 1.2292 | 0.9 | 0.422 | 1.64E-07 | 9 |

| Cluster |
|---|
| #0 Gad2, Meis2, Pcdh17, Nsf, Sst, Clu, Nrgn, Tac1 -> ZI |
| #1 Gal, Cartpt, Scg2, Prlr, Th, Cck, Slc18a2, Foxg1 -> High activation POA |
| #2 Cd24a, Six3, Pkib, Vip, Lhx1os, Lhx1, Six6, Grp -> SCN |
| #3 Grp, Vip, Chst8 -> SCN |
| #4 Slc17a6, Ntng1, Cartpt, Isl1, Gpr101, Nos1, Lmo2 -> high activation -> DMH |
| #5 Lhx1, Six6, Lhx1os, Rora, Nr2f2, Penk -> SCN |
| #6 Grp, Calb2, Lhx1os, Six3 -> SCN |
| #7 Vip, Dusp1, Vgf, Jund, Egr1, Jun -> high activation SCN neurons |
| #8 Tac1, Isl1, Ntng1, Cited1, Thrb, Trh, Hmx2, Foxp2 -> high activation - TMN |
| #9 Cck, Nfix, Prlr, Trh, Gad2, Tcf4, Pnoc, Foxg1 -> POA |
| #10 Nrgn, Nptx1, Thy1, Slc17a6, Fezf1, Gabra1, Nr5a1 -> VMH Glutamatergic neurons |

| | | | | | |
|---|---|---|---|---|---|
| Syt1 | 1.2271 | 0.965 | 0.766 | 1.68E-59 | 0 |
| Ets2 | 1.216 | 0.941 | 0.263 | 8.75E-09 | 10 |
| Arf3 | 1.2151 | 1 | 0.824 | 2.18E-06 | 10 |
| Ppp3r1 | 1.214 | 1 | 0.691 | 9.45E-05 | 10 |
| Sparcl1 | 1.2105 | 0.882 | 0.161 | 4.17E-06 | 10 |
| Crabp1 | 1.2095 | 0.083 | 0.021 | 0.0053 | 4 |
| Rasgrf2 | 1.1876 | 0.941 | 0.688 | 7.37E-05 | 10 |
| Nsf | 1.1781 | 1 | 0.833 | 1.92E-05 | 10 |
| Gadd45g | 1.1692 | 0.821 | 0.491 | 7.75E-17 | 7 |
| Celf2 | 1.1563 | 1 | 0.632 | 0.0003 | 10 |
| Npas4 | 1.1556 | 0.667 | 0.205 | 1.36E-17 | 7 |
| Slc17a6 | 1.1519 | 0.759 | 0.056 | 7.66E-47 | 4 |
| Isl1 | 1.1458 | 0.93 | 0.2 | 5.05E-37 | 8 |
| Vgf | 1.1442 | 0.577 | 0.258 | 3.03E-09 | 7 |
| Hrh3 | 1.1425 | 1 | 0.262 | 4.95E-07 | 10 |
| Rasl11b | 1.1365 | 0.615 | 0.137 | 2.12E-26 | 7 |
| Lhx1 | 1.1291 | 0.99 | 0.575 | 2.68E-45 | 5 |
| Tmem215 | 1.1258 | 0.8 | 0.048 | 1.06E-24 | 9 |
| Hpcal1 | 1.1215 | 0.901 | 0.362 | 1.90E-47 | 1 |
| Hpcal4 | 1.1164 | 0.941 | 0.54 | 0.0015 | 10 |
| Wbscr17 | 1.1148 | 0.882 | 0.172 | 6.22E-06 | 10 |
| Gria1 | 1.109 | 1 | 0.564 | 9.54E-05 | 10 |
| Tmem35 | 1.0985 | 1 | 0.372 | 3.75E-06 | 10 |
| Lynx1 | 1.0978 | 0.941 | 0.155 | 1.11E-05 | 10 |
| Prlr | 1.0964 | 0.967 | 0.197 | 6.70E-15 | 9 |
| Cx3cl1 | 1.0844 | 1 | 0.578 | 3.15E-05 | 10 |
| Camkk2 | 1.0821 | 0.824 | 0.198 | 0.0003 | 10 |
| Ntng1 | 1.0815 | 0.611 | 0.318 | 6.69E-14 | 4 |
| Ogfod1 | 1.0784 | 1 | 0.398 | 0.002 | 10 |
| Grm5 | 1.0736 | 1 | 0.564 | 4.11E-05 | 10 |
| Jund | 1.0716 | 1 | 0.987 | 8.75E-44 | 7 |
| Prkcb | 1.0706 | 0.941 | 0.315 | 2.66E-05 | 10 |
| Gria3 | 1.0693 | 0.882 | 0.192 | 0.0062 | 10 |
| Cbln1 | 1.0659 | 0.941 | 0.033 | 8.73E-14 | 10 |
| Ntng1 | 1.0478 | 0.944 | 0.305 | 2.43E-28 | 8 |
| Nefl | 1.0443 | 1 | 0.23 | 0.0013 | 10 |
| Cyfip2 | 1.0415 | 1 | 0.652 | 0.0023 | 10 |
| Sez6l | 1.0413 | 1 | 0.599 | 1.18E-07 | 10 |
| Cntn5 | 1.0403 | 0.824 | 0.156 | 0.0002 | 10 |
| Cdh4 | 1.0304 | 0.824 | 0.166 | 8.39E-05 | 10 |
| Nell2 | 1.0302 | 0.941 | 0.458 | 0.0018 | 10 |
| Tmem132 | 1.0233 | 0.824 | 0.273 | 0.0008 | 10 |

| | | | | | |
|---|---|---|---|---|---|
| Egr1 | 1.0202 | 0.718 | 0.296 | 2.94E-17 | 7 |
| Cd24a | 1.0192 | 0.945 | 0.419 | 5.59E-62 | 2 |
| Ywhag | 1.0184 | 1 | 0.978 | 2.48E-06 | 10 |
| Cadm3 | 1.0149 | 1 | 0.568 | 0.0005 | 10 |
| Sirpa | 0.9985 | 1 | 0.544 | 9.01E-07 | 10 |
| Igfbp5 | 0.9954 | 0.923 | 0.606 | 1.24E-20 | 7 |
| Tpbg | 0.9953 | 0.925 | 0.443 | 1.94E-50 | 2 |
| Baiap2 | 0.9914 | 1 | 0.375 | 3.04E-06 | 10 |
| Spock1 | 0.9906 | 1 | 0.39 | 0.1169 | 10 |
| Mapk1 | 0.9899 | 0.941 | 0.525 | 0.0063 | 10 |
| Apbb2 | 0.988 | 0.928 | 0.277 | 6.37E-54 | 5 |
| Plcxd2 | 0.9859 | 0.765 | 0.147 | 0.0053 | 10 |
| Prkacb | 0.9824 | 1 | 0.949 | 4.26E-08 | 10 |
| Fgfr1 | 0.98 | 0.765 | 0.258 | 0.0572 | 10 |
| Irs4 | 0.9759 | 0.757 | 0.195 | 2.78E-53 | 1 |
| Sorcs3 | 0.9733 | 0.765 | 0.182 | 0.0117 | 10 |
| Hapln4 | 0.9719 | 0.882 | 0.158 | 0.0003 | 10 |
| Synpr | 0.9711 | 0.992 | 0.683 | 2.98E-40 | 3 |
| Tppp | 0.9703 | 0.941 | 0.358 | 1 | 10 |
| Fstl5 | 0.9703 | 0.783 | 0.186 | 4.34E-47 | 1 |
| Nptx2 | 0.97 | 0.824 | 0.038 | 3.36E-09 | 10 |
| Dnm1 | 0.9689 | 1 | 0.667 | 1 | 10 |
| Sema5a | 0.9688 | 0.706 | 0.164 | 0.6688 | 10 |
| Btg2 | 0.9635 | 0.718 | 0.258 | 3.13E-12 | 7 |
| Syt10 | 0.9631 | 0.99 | 0.613 | 7.84E-43 | 5 |
| H2-Q2 | 0.9602 | 0.471 | 0.074 | 0.0167 | 10 |
| Thrb | 0.9596 | 0.817 | 0.094 | 1.01E-42 | 8 |
| Gpr149 | 0.9564 | 0.647 | 0.014 | 5.24E-09 | 10 |
| Six6 | 0.946 | 0.938 | 0.37 | 1.72E-34 | 5 |
| Cbx6 | 0.9454 | 1 | 0.495 | 8.73E-05 | 10 |
| Synpr | 0.9434 | 0.993 | 0.675 | 2.37E-59 | 2 |
| Atp1a1 | 0.9425 | 1 | 0.59 | 3.02E-06 | 10 |
| Dgkb | 0.9423 | 0.915 | 0.253 | 2.01E-27 | 8 |
| Trh | 0.9395 | 0.233 | 0.038 | 1 | 9 |
| Phyhip | 0.9393 | 0.706 | 0.247 | 1 | 10 |
| Atp6v1a | 0.938 | 1 | 0.804 | 0.0011 | 10 |
| Adcyap1 | 0.9373 | 0.647 | 0.023 | 7.23E-05 | 10 |
| Pcdh17 | 0.9361 | 1 | 0.634 | 2.33E-12 | 9 |
| Scg2 | 0.9294 | 1 | 0.956 | 2.13E-54 | 1 |
| Gad2 | 0.9249 | 1 | 0.814 | 1.04E-12 | 9 |
| Nfib | 0.9236 | 0.7 | 0.051 | 1.84E-15 | 9 |
| Lmo1 | 0.9229 | 0.3 | 0.049 | 1.37E-05 | 9 |

| | | | | | |
|---|---|---|---|---|---|
| Prok2 | 0.9218 | 0.244 | 0.04 | 1.78E-09 | 7 |
| Snca | 0.9208 | 0.882 | 0.315 | 0.1818 | 10 |
| Kif5a | 0.9206 | 1 | 0.715 | 0.0058 | 10 |
| A230065I | 0.9199 | 0.706 | 0.151 | 2.59E-05 | 10 |
| Mgat4c | 0.9174 | 0.824 | 0.215 | 1 | 10 |
| Rora | 0.9165 | 0.859 | 0.419 | 3.85E-22 | 7 |
| Gstm7 | 0.9161 | 0.873 | 0.306 | 6.57E-24 | 8 |
| Fosb | 0.9156 | 0.615 | 0.146 | 1.19E-16 | 7 |
| Tuba4a | 0.9117 | 0.882 | 0.505 | 0.0414 | 10 |
| Cited1 | 0.9098 | 0.634 | 0.131 | 1.39E-17 | 8 |
| Susd4 | 0.9069 | 0.941 | 0.358 | 0.0076 | 10 |
| Enc1 | 0.9059 | 0.824 | 0.261 | 1 | 10 |
| Hpcal1 | 0.9032 | 0.861 | 0.39 | 1.77E-22 | 4 |
| Syt1 | 0.8981 | 1 | 0.796 | 1.60E-10 | 9 |
| Camk2b | 0.8932 | 1 | 0.785 | 0.0001 | 10 |
| Erdr1 | 0.8917 | 1 | 0.895 | 1.92E-19 | 3 |
| Cyb5b | 0.8911 | 0.941 | 0.334 | 0.0018 | 10 |
| Ngb | 0.8902 | 1 | 0.763 | 7.74E-23 | 8 |
| Scd2 | 0.8859 | 1 | 0.802 | 3.47E-07 | 10 |
| Cox6a2 | 0.882 | 0.4 | 0.026 | 1.96E-06 | 9 |
| Faim2 | 0.8815 | 1 | 0.974 | 1.34E-11 | 10 |
| Cdk14 | 0.8805 | 0.882 | 0.3 | 0.8115 | 10 |
| Doc2b | 0.8758 | 0.706 | 0.09 | 4.86E-05 | 10 |
| Prlr | 0.8731 | 0.678 | 0.146 | 1.30E-38 | 1 |
| Adam23 | 0.8729 | 0.882 | 0.344 | 0.0005 | 10 |
| Six3 | 0.8726 | 0.979 | 0.638 | 5.68E-43 | 2 |
| Pde8b | 0.8702 | 0.824 | 0.107 | 0.0032 | 10 |
| Lhx1os | 0.867 | 0.979 | 0.502 | 2.25E-28 | 5 |
| Nnat | 0.8669 | 0.967 | 0.888 | 1 | 9 |
| Unc80 | 0.8664 | 0.941 | 0.661 | 0.0041 | 10 |
| Tpbg | 0.8652 | 0.927 | 0.454 | 7.12E-34 | 3 |
| Nnat | 0.8649 | 0.936 | 0.887 | 4.67E-06 | 7 |
| Arhgap36 | 0.8637 | 0.706 | 0.189 | 0.6525 | 10 |
| Fam234b | 0.8636 | 0.941 | 0.368 | 5.32E-05 | 10 |
| Scamp1 | 0.8611 | 1 | 0.542 | 0.0083 | 10 |
| Cartpt | 0.8608 | 0.259 | 0.113 | 0.2298 | 4 |
| Hbegf | 0.8566 | 0.628 | 0.183 | 3.53E-17 | 7 |
| Map2k1 | 0.8564 | 0.882 | 0.55 | 1 | 10 |
| Psap | 0.8546 | 1 | 0.964 | 3.07E-09 | 10 |
| Gria1 | 0.8541 | 0.967 | 0.56 | 1.67E-09 | 9 |
| Rprm | 0.8535 | 0.836 | 0.349 | 6.78E-32 | 2 |
| Cpne5 | 0.8532 | 0.873 | 0.245 | 1.45E-26 | 8 |

| | | | | | |
|---|---|---|---|---|---|
| Mboat7 | 0.848 | 0.941 | 0.639 | 0.0001 | 10 |
| Pgm2l1 | 0.8456 | 0.941 | 0.468 | 1 | 10 |
| Deptor | 0.8454 | 0.588 | 0.044 | 6.67E-05 | 10 |
| Rock2 | 0.842 | 1 | 0.614 | 0.0015 | 10 |
| Adgrl2 | 0.8417 | 0.882 | 0.159 | 0.0283 | 10 |
| Pip4k2b | 0.8413 | 1 | 0.309 | 0.0236 | 10 |
| Gabrb3 | 0.841 | 1 | 0.672 | 4.01E-05 | 10 |
| Tmsb4x | 0.8404 | 1 | 0.994 | 7.83E-10 | 9 |
| Tspan7 | 0.8395 | 1 | 0.846 | 0.0033 | 10 |
| Impad1 | 0.8395 | 1 | 0.538 | 0.0099 | 10 |
| Grp | 0.8382 | 0.753 | 0.305 | 2.20E-05 | 6 |
| Sox14 | 0.8377 | 0.775 | 0.013 | 9.51E-54 | 8 |
| Ptn | 0.8356 | 0.796 | 0.48 | 7.54E-17 | 4 |
| Ncald | 0.8352 | 0.986 | 0.658 | 8.18E-24 | 8 |
| Tenm4 | 0.835 | 0.959 | 0.487 | 1.36E-36 | 5 |
| Dkk3 | 0.8339 | 0.588 | 0.101 | 0.1754 | 10 |
| Calb1 | 0.8319 | 0.967 | 0.758 | 1.54E-05 | 9 |
| Kcnb2 | 0.8304 | 0.824 | 0.167 | 0.0027 | 10 |
| Tef | 0.8301 | 1 | 0.637 | 0.003 | 10 |
| Thy1 | 0.8287 | 0.775 | 0.344 | 2.83E-31 | 0 |
| Pkib | 0.8266 | 0.856 | 0.432 | 4.58E-19 | 5 |
| Ecel1 | 0.8254 | 0.767 | 0.215 | 7.22E-05 | 9 |
| Nat8l | 0.8224 | 1 | 0.301 | 0.1617 | 10 |
| Kcna2 | 0.8214 | 0.765 | 0.255 | 0.9897 | 10 |
| Crim1 | 0.8213 | 0.7 | 0.195 | 6.01E-08 | 9 |
| Aatk | 0.8188 | 0.941 | 0.599 | 5.37E-05 | 10 |
| Ajap1 | 0.8185 | 0.918 | 0.392 | 2.33E-29 | 5 |
| Rangap1 | 0.818 | 1 | 0.637 | 0.0003 | 10 |
| Atp9a | 0.8169 | 1 | 0.736 | 2.54E-06 | 10 |
| Ptprt | 0.8154 | 0.867 | 0.24 | 8.19E-09 | 9 |
| Snap25 | 0.814 | 1 | 0.915 | 8.96E-23 | 5 |
| Tac1 | 0.8139 | 0.125 | 0.085 | 1 | 0 |
| Syt10 | 0.8123 | 0.949 | 0.622 | 1.43E-15 | 7 |
| Pcnx | 0.8103 | 0.882 | 0.334 | 0.512 | 10 |
| Cbln2 | 0.805 | 0.626 | 0.227 | 4.89E-24 | 3 |
| Arhgap36 | 0.804 | 0.38 | 0.157 | 3.27E-17 | 0 |
| Plec | 0.8036 | 0.706 | 0.158 | 0.3513 | 10 |
| Lamp5 | 0.8018 | 0.533 | 0.085 | 1.56E-06 | 9 |
| Th | 0.8002 | 0.467 | 0.053 | 1.18E-32 | 1 |
| 1500009 | 0.7989 | 0.833 | 0.584 | 0.0004 | 9 |
| Tox3 | 0.7988 | 0.866 | 0.333 | 7.24E-27 | 5 |
| Doc2a | 0.7969 | 0.824 | 0.174 | 0.0004 | 10 |

| | | | | | |
|---|---|---|---|---|---|
| Nlgn1 | 0.7963 | 0.941 | 0.456 | 0.0045 | 10 |
| Hspa4l | 0.795 | 0.882 | 0.522 | 1 | 10 |
| Nckap1 | 0.7947 | 1 | 0.625 | 0.0646 | 10 |
| Syn1 | 0.7925 | 0.941 | 0.609 | 1 | 10 |
| Crtc1 | 0.7909 | 0.882 | 0.268 | 0.0018 | 10 |
| Vstm2l | 0.7907 | 1 | 0.825 | 2.55E-20 | 3 |
| Rora | 0.7903 | 0.876 | 0.409 | 6.86E-23 | 5 |
| Id4 | 0.7883 | 0.979 | 0.593 | 1.15E-42 | 2 |
| Isl1 | 0.7876 | 0.75 | 0.193 | 1.28E-26 | 4 |
| Moxd1 | 0.7862 | 0.633 | 0.013 | 2.35E-19 | 9 |
| Nsg2 | 0.786 | 1 | 0.948 | 0.0906 | 10 |
| Cntnap1 | 0.7845 | 0.824 | 0.199 | 0.0339 | 10 |
| Fam168b | 0.7815 | 0.941 | 0.374 | 1 | 10 |
| Hs6st2 | 0.779 | 0.824 | 0.352 | 3.31E-27 | 4 |
| Fabp5 | 0.7789 | 0.986 | 0.856 | 5.28E-20 | 8 |
| Resp18 | 0.778 | 1 | 0.999 | 7.17E-08 | 3 |
| Cdh22 | 0.7773 | 0.824 | 0.243 | 0.074 | 10 |
| Atp2a2 | 0.7769 | 1 | 0.792 | 0.01 | 10 |
| Ryr2 | 0.7762 | 0.882 | 0.191 | 0.0004 | 10 |
| Aldoc | 0.7754 | 0.941 | 0.291 | 1 | 10 |
| Slc35f1 | 0.7749 | 0.647 | 0.168 | 1 | 10 |
| Fut9 | 0.7746 | 0.882 | 0.347 | 1 | 10 |
| Rrad | 0.7745 | 0.437 | 0.152 | 0.0061 | 8 |
| Ackr1 | 0.7743 | 0.824 | 0.137 | 0.0051 | 10 |
| Cck | 0.7727 | 0.092 | 0.029 | 0.6607 | 1 |
| Lmo3 | 0.7725 | 0.667 | 0.139 | 1.02E-05 | 9 |
| Pfkp | 0.7699 | 0.941 | 0.321 | 0.0116 | 10 |
| Cbx3 | 0.7679 | 1 | 0.818 | 2.47E-37 | 2 |
| Tcf4 | 0.7676 | 0.8 | 0.291 | 1.01E-06 | 9 |
| Ngb | 0.7674 | 0.991 | 0.756 | 6.11E-28 | 4 |
| Stx1a | 0.7667 | 0.765 | 0.327 | 0.3602 | 10 |
| Camk2a | 0.7643 | 1 | 0.711 | 0.0316 | 10 |
| Rasd1 | 0.7634 | 0.454 | 0.061 | 1.32E-25 | 5 |
| Lhx1os | 0.7632 | 0.923 | 0.515 | 3.84E-19 | 7 |
| Rtn3 | 0.7629 | 1 | 0.978 | 5.79E-05 | 10 |
| Scn8a | 0.7628 | 0.765 | 0.343 | 0.9975 | 10 |
| Pnoc | 0.7624 | 0.6 | 0.141 | 0.0054 | 9 |
| Six3 | 0.76 | 0.987 | 0.66 | 7.13E-23 | 7 |
| Zbtb20 | 0.7591 | 0.961 | 0.704 | 7.96E-34 | 1 |
| Nt5dc3 | 0.7575 | 0.882 | 0.326 | 0.0038 | 10 |
| Synj1 | 0.7574 | 1 | 0.651 | 0.0072 | 10 |
| Ids | 0.7565 | 0.941 | 0.537 | 0.8091 | 10 |

| | | | | | |
|---|---|---|---|---|---|
| Slc18a2 | 0.7536 | 0.579 | 0.059 | 1.25E-44 | 1 |
| Slit1 | 0.7533 | 0.765 | 0.261 | 0.0308 | 10 |
| Fam19a1 | 0.7531 | 0.756 | 0.385 | 6.11E-22 | 3 |
| Adgrl3 | 0.7521 | 0.941 | 0.499 | 0.9371 | 10 |
| Tpd52l1 | 0.752 | 0.863 | 0.411 | 5.36E-32 | 2 |
| Gas6 | 0.7495 | 0.7 | 0.281 | 1.16E-05 | 9 |
| Grin1 | 0.7489 | 1 | 0.679 | 0.0225 | 10 |
| Nptn | 0.7479 | 1 | 0.823 | 1 | 10 |
| Dzank1 | 0.7478 | 1 | 0.574 | 1 | 10 |
| Trh | 0.7477 | 0.282 | 0.027 | 1.38E-05 | 8 |
| Car10 | 0.7469 | 0.882 | 0.301 | 1 | 10 |
| Adgrl1 | 0.7464 | 0.941 | 0.676 | 0.1329 | 10 |
| Ogfrl1 | 0.746 | 0.796 | 0.617 | 5.32E-21 | 1 |
| Atmin | 0.7458 | 0.882 | 0.187 | 0.0003 | 10 |
| Sesn3 | 0.7457 | 1 | 0.641 | 0.0139 | 10 |
| C1ql2 | 0.7456 | 0.167 | 0.053 | 1 | 9 |
| Alcam | 0.7452 | 1 | 0.774 | 7.96E-20 | 3 |
| Fam155a | 0.7448 | 0.882 | 0.519 | 0.215 | 10 |
| Eef1a2 | 0.7442 | 1 | 0.878 | 0.0011 | 10 |
| Cnr1 | 0.7436 | 0.704 | 0.156 | 1.42E-31 | 4 |
| Slc12a5 | 0.7426 | 1 | 0.894 | 6.65E-14 | 3 |
| Glrb | 0.7424 | 0.765 | 0.346 | 1 | 10 |
| Grin2b | 0.7409 | 1 | 0.669 | 1 | 10 |
| Crtac1 | 0.7403 | 0.718 | 0.178 | 5.54E-19 | 8 |
| Nova1 | 0.7391 | 0.647 | 0.215 | 1 | 10 |
| Pkib | 0.7374 | 0.89 | 0.405 | 8.72E-26 | 2 |
| Dgcr2 | 0.7366 | 0.941 | 0.434 | 1 | 10 |
| Nts | 0.7361 | 0.235 | 0.025 | 1 | 10 |
| Gabrb1 | 0.7359 | 1 | 0.854 | 0.003 | 10 |
| Tmem163 | 0.7354 | 0.647 | 0.113 | 1 | 10 |
| Cpne7 | 0.7349 | 0.706 | 0.187 | 1 | 10 |
| Btg1 | 0.7345 | 0.987 | 0.762 | 3.75E-19 | 7 |
| Lmo4 | 0.7345 | 0.765 | 0.237 | 1 | 10 |
| Nkx2-1 | 0.7337 | 0.588 | 0.076 | 0.1931 | 10 |
| Bhlhe41 | 0.7334 | 0.733 | 0.262 | 2.89E-17 | 2 |
| Nrgn | 0.7316 | 0.32 | 0.218 | 1.20E-05 | 0 |
| Gabra2 | 0.7308 | 0.824 | 0.601 | 0.5245 | 10 |
| Mkl2 | 0.7299 | 0.941 | 0.493 | 0.4326 | 10 |
| Gpr101 | 0.7273 | 0.491 | 0.071 | 9.95E-22 | 4 |
| Dgkb | 0.7254 | 0.737 | 0.225 | 9.29E-32 | 1 |
| Ddc | 0.7254 | 0.746 | 0.2 | 1.74E-14 | 8 |
| Hspa1a | 0.7253 | 0.295 | 0.183 | 1 | 7 |

| | | | | | |
|---|---|---|---|---|---|
| Serpine2 | 0.7251 | 0.882 | 0.265 | 0.4118 | 10 |
| Atp8a1 | 0.725 | 1 | 0.786 | 0.9635 | 10 |
| Cntn1 | 0.7246 | 0.941 | 0.629 | 1 | 10 |
| Pde1a | 0.7242 | 0.7 | 0.217 | 0.0005 | 9 |
| Gabrg1 | 0.7233 | 0.854 | 0.55 | 6.95E-20 | 3 |
| Efr3b | 0.7224 | 0.765 | 0.196 | 1 | 10 |
| Vat1l | 0.7223 | 0.941 | 0.591 | 1 | 10 |
| Cthrc1 | 0.7217 | 0.815 | 0.292 | 1.23E-32 | 2 |
| Vip | 0.7197 | 0.699 | 0.401 | 1.11E-28 | 2 |
| Nr5a1 | 0.7185 | 0.706 | 0.003 | 5.72E-12 | 10 |
| Greb1 | 0.7184 | 0.633 | 0.051 | 1.87E-11 | 9 |
| Fos | 0.7184 | 0.718 | 0.41 | 6.47E-05 | 8 |
| Lhx1os | 0.7184 | 0.986 | 0.477 | 4.72E-38 | 2 |
| Cntn4 | 0.7177 | 0.69 | 0.107 | 2.69E-24 | 8 |
| Ptprd | 0.7165 | 0.984 | 0.821 | 5.36E-13 | 3 |
| Kcnq2 | 0.7164 | 0.941 | 0.499 | 0.0174 | 10 |
| Prkar1b | 0.7164 | 1 | 0.961 | 0.1704 | 10 |
| Akap17b | 0.7159 | 0.835 | 0.333 | 1.40E-26 | 5 |
| Bmpr2 | 0.7151 | 0.941 | 0.364 | 1 | 10 |
| Akap11 | 0.7141 | 1 | 0.776 | 1 | 10 |
| Camk1g | 0.7138 | 0.588 | 0.129 | 1 | 10 |
| Pcsk2 | 0.7135 | 0.907 | 0.832 | 3.26E-15 | 4 |
| Chga | 0.7118 | 0.821 | 0.728 | 0.0007 | 7 |
| Syt7 | 0.7109 | 0.706 | 0.174 | 1 | 10 |
| Gm2888‹ | 0.7104 | 0.433 | 0.013 | 4.14E-12 | 9 |
| Lin7a | 0.7101 | 0.966 | 0.643 | 2.06E-38 | 2 |
| Cpne4 | 0.7085 | 0.732 | 0.146 | 1.32E-18 | 8 |
| Rcn1 | 0.7073 | 0.767 | 0.316 | 0.0008 | 9 |
| Syn2 | 0.7063 | 1 | 0.956 | 0.576 | 10 |
| Cplx2 | 0.7047 | 0.824 | 0.261 | 1 | 10 |
| Ube2ql1 | 0.7037 | 0.941 | 0.378 | 1 | 10 |
| Junb | 0.7036 | 0.654 | 0.312 | 0.0002 | 7 |
| Rprm | 0.7035 | 0.833 | 0.401 | 0.01 | 9 |
| Atp6ap2 | 0.703 | 0.992 | 0.816 | 3.46E-15 | 3 |
| Itpr1 | 0.7022 | 0.706 | 0.146 | 0.7162 | 10 |
| Got2 | 0.702 | 0.941 | 0.607 | 0.9217 | 10 |
| Lrrc4b | 0.7015 | 0.882 | 0.75 | 1 | 10 |
| Ctla2a | 0.7004 | 0.41 | 0.092 | 1.36E-10 | 7 |
| Mef2d | 0.7003 | 0.941 | 0.334 | 0.1431 | 10 |
| Rpl36a | 0.7003 | 1 | 0.925 | 1.73E-15 | 8 |
| Hist3h2b | 0.7 | 0.887 | 0.676 | 1.23E-16 | 5 |
| Snph | 0.6996 | 0.824 | 0.222 | 0.0423 | 10 |

| | | | | | |
|---|---|---|---|---|---|
| Cmtm4 | 0.6992 | 0.824 | 0.26 | 0.6314 | 10 |
| Gnaq | 0.6977 | 0.941 | 0.549 | 1 | 10 |
| 6-Mar | 0.6967 | 0.941 | 0.674 | 0.0479 | 10 |
| Gm13889 | 0.6955 | 0.732 | 0.343 | 5.91E-06 | 8 |
| Cntnap5a | 0.6951 | 0.887 | 0.275 | 1.60E-21 | 8 |
| Stxbp1 | 0.6944 | 1 | 0.921 | 1 | 10 |
| Abr | 0.6923 | 0.941 | 0.703 | 1 | 10 |
| Gabra5 | 0.692 | 0.706 | 0.246 | 1 | 10 |
| Sez6 | 0.6919 | 0.706 | 0.146 | 1 | 10 |
| Egr4 | 0.6902 | 0.239 | 0.071 | 0.0051 | 8 |
| Snap25 | 0.6895 | 1 | 0.911 | 2.76E-28 | 2 |
| D430041 | 0.6894 | 0.706 | 0.236 | 0.377 | 10 |
| Syp | 0.6894 | 1 | 0.935 | 0.4075 | 10 |
| Clstn1 | 0.6887 | 1 | 0.847 | 0.0038 | 10 |
| Hsph1 | 0.6885 | 1 | 0.843 | 0.0079 | 10 |
| Matk | 0.688 | 1 | 0.742 | 0.094 | 10 |
| Cdk18 | 0.6879 | 0.765 | 0.206 | 0.3448 | 10 |
| Atp6v1b2 | 0.6876 | 1 | 0.859 | 1 | 10 |
| Ppp1r9a | 0.6873 | 0.882 | 0.563 | 1 | 10 |
| Adam11 | 0.6869 | 0.765 | 0.211 | 1 | 10 |
| Ppfia2 | 0.6868 | 0.884 | 0.528 | 2.64E-32 | 2 |
| Epha5 | 0.6861 | 1 | 0.831 | 1.18E-14 | 3 |
| Lhx1 | 0.6843 | 1 | 0.553 | 4.34E-38 | 2 |
| Actr1b | 0.6815 | 0.941 | 0.453 | 1 | 10 |
| Taok1 | 0.6798 | 0.941 | 0.535 | 1 | 10 |
| Acvr1c | 0.6796 | 0.412 | 0.01 | 0.0004 | 10 |
| Ecel1 | 0.679 | 0.704 | 0.198 | 6.81E-09 | 8 |
| Gabra5 | 0.6782 | 0.722 | 0.203 | 2.16E-26 | 4 |
| Six6 | 0.6775 | 0.945 | 0.34 | 5.30E-29 | 2 |
| Kif3c | 0.6771 | 0.824 | 0.306 | 1 | 10 |
| Oprl1 | 0.6767 | 0.941 | 0.475 | 0.0106 | 10 |
| Zcchc12 | 0.6758 | 0.993 | 0.957 | 1.82E-34 | 1 |
| Oprk1 | 0.674 | 0.625 | 0.189 | 3.55E-26 | 1 |
| Pde10a | 0.6737 | 0.676 | 0.275 | 9.66E-09 | 8 |
| Epb41l3 | 0.6736 | 0.866 | 0.518 | 1.42E-21 | 5 |
| Ier2 | 0.673 | 0.423 | 0.084 | 2.59E-10 | 7 |
| Epha4 | 0.6728 | 0.706 | 0.329 | 1 | 10 |
| mt-Co3 | 0.6713 | 1 | 1 | 0.0008 | 10 |
| Dlg4 | 0.6697 | 0.941 | 0.721 | 0.5536 | 10 |
| Rac1 | 0.6696 | 1 | 0.632 | 1 | 10 |
| Pou2f2 | 0.669 | 0.945 | 0.55 | 1.72E-43 | 2 |
| Ptchd2 | 0.668 | 0.647 | 0.118 | 0.0503 | 10 |

| | | | | | |
|---|---|---|---|---|---|
| Cited2 | 0.6674 | 0.676 | 0.472 | 0.0006 | 8 |
| Rgs2 | 0.6665 | 0.765 | 0.193 | 0.6278 | 10 |
| Efhd2 | 0.6665 | 0.882 | 0.351 | 1 | 10 |
| Ptprz1 | 0.6661 | 0.778 | 0.388 | 3.45E-14 | 4 |
| Ppp2r1a | 0.6659 | 0.941 | 0.744 | 1 | 10 |
| Dbp | 0.6641 | 0.91 | 0.632 | 2.13E-13 | 7 |
| Htr1b | 0.6638 | 0.529 | 0.084 | 1 | 10 |
| Ablim2 | 0.6637 | 0.824 | 0.204 | 1 | 10 |
| Plxna1 | 0.6635 | 0.941 | 0.245 | 1 | 10 |
| Celsr2 | 0.6635 | 0.941 | 0.495 | 1 | 10 |
| Tppp3 | 0.6633 | 0.941 | 0.454 | 0.0022 | 10 |
| Chodl | 0.6633 | 0.732 | 0.215 | 3.85E-13 | 8 |
| Cadps2 | 0.6619 | 0.829 | 0.518 | 1.23E-20 | 3 |
| Mtpn | 0.6617 | 0.882 | 0.411 | 1 | 10 |
| Sun1 | 0.6568 | 0.882 | 0.335 | 0.4587 | 10 |
| Rasgrp1 | 0.6566 | 0.814 | 0.234 | 4.49E-22 | 5 |
| Lhx1 | 0.6566 | 1 | 0.582 | 1.34E-17 | 7 |
| Pabpn1 | 0.6562 | 0.984 | 0.859 | 5.43E-09 | 3 |
| Pcp4 | 0.656 | 1 | 0.872 | 1.15E-08 | 8 |
| Uba52 | 0.6547 | 0.993 | 0.856 | 3.06E-29 | 2 |
| Cthrc1 | 0.6543 | 0.593 | 0.331 | 5.04E-22 | 3 |
| Unc5d | 0.6536 | 0.8 | 0.335 | 9.91E-05 | 9 |
| Aak1 | 0.6519 | 0.882 | 0.518 | 1 | 10 |
| Scn2b | 0.6518 | 0.882 | 0.509 | 1 | 10 |
| Cd24a | 0.6516 | 0.897 | 0.457 | 2.02E-11 | 7 |
| Ube2o | 0.6503 | 0.824 | 0.415 | 0.9922 | 10 |
| Ret | 0.6497 | 0.634 | 0.094 | 2.31E-22 | 8 |
| Scamp5 | 0.6492 | 0.941 | 0.485 | 1 | 10 |
| Adgrb3 | 0.6477 | 0.824 | 0.475 | 1 | 10 |
| Stox2 | 0.647 | 0.765 | 0.371 | 1 | 10 |
| Aco2 | 0.6458 | 0.941 | 0.536 | 1 | 10 |
| Fam171b | 0.6457 | 0.933 | 0.81 | 7.22E-07 | 9 |
| Dpp6 | 0.6456 | 1 | 0.842 | 0.0054 | 10 |
| D17Wsu9 | 0.6453 | 0.824 | 0.268 | 0.3065 | 10 |
| Plxna4 | 0.6453 | 0.824 | 0.205 | 1 | 10 |
| Nrip1 | 0.6452 | 0.7 | 0.166 | 3.96E-06 | 9 |
| Psd3 | 0.6448 | 0.867 | 0.433 | 2.50E-06 | 9 |
| Slc25a5 | 0.6435 | 0.711 | 0.422 | 1.37E-11 | 5 |
| Hspa12a | 0.6433 | 0.706 | 0.288 | 0.8908 | 10 |
| Arnt2 | 0.6428 | 0.824 | 0.319 | 1 | 10 |
| Gnb1 | 0.6427 | 1 | 0.93 | 3.12E-29 | 2 |
| Msi2 | 0.6405 | 0.928 | 0.63 | 1.06E-16 | 5 |

| | | | | | |
|---|---|---|---|---|---|
| Nr2f2 | 0.6403 | 0.773 | 0.387 | 4.08E-11 | 5 |
| Bcl11a | 0.6402 | 0.835 | 0.403 | 3.09E-18 | 5 |
| Itm2a | 0.6397 | 0.423 | 0.101 | 9.17E-12 | 8 |
| Map1a | 0.6395 | 0.941 | 0.492 | 1 | 10 |
| Scn1a | 0.6392 | 0.941 | 0.271 | 1 | 10 |
| Spin1 | 0.639 | 0.941 | 0.486 | 1 | 10 |
| Sorcs1 | 0.6384 | 0.882 | 0.283 | 0.4275 | 10 |
| Cnksr2 | 0.6384 | 0.773 | 0.299 | 2.79E-18 | 5 |
| Gm1007€ | 0.6373 | 0.859 | 0.703 | 9.80E-08 | 8 |
| AU04032 | 0.6371 | 0.824 | 0.253 | 1 | 10 |
| Rprml | 0.6364 | 0.647 | 0.073 | 1 | 10 |
| Mmp17 | 0.6358 | 0.765 | 0.183 | 0.3456 | 10 |
| Slc4a10 | 0.6356 | 0.765 | 0.336 | 1 | 10 |
| Vip | 0.6352 | 0.862 | 0.388 | 2.83E-16 | 3 |
| Cd83 | 0.6341 | 0.765 | 0.386 | 1 | 10 |
| Cnih3 | 0.6338 | 0.706 | 0.129 | 0.0126 | 10 |
| Zyg11b | 0.6316 | 0.882 | 0.38 | 1 | 10 |
| Grp | 0.6315 | 0.664 | 0.296 | 1.74E-19 | 2 |
| Sv2a | 0.63 | 1 | 0.924 | 0.1945 | 10 |
| Gpr155 | 0.6299 | 0.588 | 0.105 | 1 | 10 |
| Tmem30a | 0.6298 | 1 | 0.867 | 0.0962 | 10 |
| Tenm2 | 0.6285 | 0.706 | 0.376 | 1 | 10 |
| Lmo2 | 0.6271 | 0.454 | 0.069 | 7.37E-24 | 4 |
| Wnt5a | 0.6265 | 0.471 | 0.055 | 1 | 10 |
| Gabrb1 | 0.6259 | 0.992 | 0.839 | 1.54E-10 | 3 |
| Parva | 0.6259 | 0.706 | 0.107 | 0.588 | 10 |
| Syngr1 | 0.6257 | 0.941 | 0.511 | 1 | 10 |
| Gad2 | 0.6251 | 0.86 | 0.81 | 3.62E-13 | 0 |
| Spock3 | 0.6239 | 0.78 | 0.521 | 1.14E-15 | 3 |
| Atp2b2 | 0.6239 | 1 | 0.723 | 0.4626 | 10 |
| Lmtk3 | 0.6237 | 0.824 | 0.523 | 1 | 10 |
| Clstn3 | 0.6235 | 1 | 0.822 | 0.8324 | 10 |
| Ankrd34b | 0.6233 | 0.647 | 0.13 | 0.8384 | 10 |
| Ptma | 0.623 | 1 | 0.986 | 2.41E-21 | 7 |
| Rgs10 | 0.6223 | 0.593 | 0.272 | 5.32E-12 | 4 |
| Ank3 | 0.6219 | 1 | 0.848 | 1 | 10 |
| Cntn2 | 0.6217 | 0.882 | 0.23 | 1 | 10 |
| Gls | 0.6216 | 0.882 | 0.636 | 1 | 10 |
| Calb2 | 0.6214 | 0.795 | 0.563 | 2.89E-05 | 7 |
| Ldb2 | 0.6211 | 0.882 | 0.238 | 0.0518 | 10 |
| Ache | 0.621 | 0.882 | 0.546 | 0.901 | 10 |
| Ctnna2 | 0.6208 | 0.7 | 0.347 | 0.0003 | 9 |

| | | | | | |
|---|---|---|---|---|---|
| Dcun1d4 | 0.6206 | 0.882 | 0.346 | 1 | 10 |
| Grcc10 | 0.6205 | 0.986 | 0.879 | 2.01E-36 | 2 |
| Dnajb1 | 0.6205 | 0.577 | 0.313 | 0.0069 | 8 |
| Hmgn1 | 0.6197 | 1 | 0.915 | 1.87E-23 | 5 |
| Kcnh7 | 0.6194 | 0.588 | 0.136 | 1 | 10 |
| Hs3st4 | 0.6188 | 0.647 | 0.053 | 0.2086 | 10 |
| Penk | 0.6168 | 0.66 | 0.303 | 2.58E-06 | 5 |
| Mfsd6 | 0.6163 | 0.824 | 0.404 | 1 | 10 |
| Fbxw7 | 0.6157 | 0.824 | 0.379 | 1 | 10 |
| Brinp2 | 0.6143 | 0.667 | 0.314 | 0.0044 | 9 |
| Itfg1 | 0.614 | 0.882 | 0.634 | 1 | 10 |
| Khdrbs3 | 0.6138 | 0.867 | 0.663 | 0.0008 | 9 |
| Syt2 | 0.6136 | 0.588 | 0.015 | 3.46E-05 | 10 |
| Tmsb10 | 0.6136 | 1 | 0.997 | 1.85E-36 | 2 |
| Endod1 | 0.6133 | 0.706 | 0.317 | 1 | 10 |
| Mapk9 | 0.6131 | 0.824 | 0.429 | 1 | 10 |
| Gm1610! | 0.6121 | 0.897 | 0.401 | 2.55E-28 | 2 |
| Pcdh17 | 0.612 | 0.925 | 0.582 | 7.77E-24 | 0 |
| Cpne4 | 0.612 | 0.639 | 0.135 | 2.91E-23 | 4 |
| Cplx2 | 0.6113 | 0.8 | 0.255 | 0.0023 | 9 |
| Anxa5 | 0.6111 | 0.583 | 0.205 | 2.73E-19 | 4 |
| Cab39 | 0.6108 | 0.882 | 0.447 | 1 | 10 |
| Slc7a14 | 0.6106 | 0.882 | 0.296 | 1 | 10 |
| Kcnip1 | 0.6105 | 0.706 | 0.164 | 1 | 10 |
| Fam162a | 0.6104 | 0.732 | 0.496 | 2.77E-11 | 5 |
| Svop | 0.6093 | 0.882 | 0.532 | 1 | 10 |
| Gabrg2 | 0.6092 | 1 | 0.669 | 1 | 10 |
| Nms | 0.6091 | 0.321 | 0.017 | 7.06E-17 | 7 |
| Chrna4 | 0.6083 | 0.588 | 0.021 | 0.0029 | 10 |
| Igfbp5 | 0.6082 | 0.959 | 0.587 | 1.64E-16 | 3 |
| Dlk1 | 0.6082 | 0.93 | 0.692 | 2.00E-10 | 8 |
| Syt4 | 0.6081 | 0.987 | 0.867 | 6.02E-12 | 7 |
| Slc22a17 | 0.6071 | 1 | 0.962 | 0.0002 | 3 |
| Tapt1 | 0.6071 | 0.765 | 0.154 | 0.0252 | 10 |
| Unc5a | 0.6066 | 0.706 | 0.279 | 1 | 10 |
| Slc36a4 | 0.6064 | 0.882 | 0.284 | 1 | 10 |
| Ap2a1 | 0.6058 | 0.941 | 0.446 | 1 | 10 |
| Amigo2 | 0.6055 | 0.647 | 0.232 | 1 | 10 |
| Gmfb | 0.6054 | 0.824 | 0.358 | 1 | 10 |
| Pcbd1 | 0.6046 | 0.93 | 0.707 | 4.65E-10 | 8 |
| Tmem151 | 0.6045 | 0.588 | 0.136 | 0.4311 | 10 |
| Rmst | 0.6042 | 0.713 | 0.445 | 3.01E-11 | 4 |

| | | | | | |
|---|---|---|---|---|---|
| Hk1 | 0.6019 | 0.882 | 0.515 | 1 | 10 |
| Zmat4 | 0.6018 | 0.765 | 0.14 | 1 | 10 |
| Atp6v1g2 | 0.6007 | 1 | 0.909 | 0.0001 | 9 |
| Id2 | 0.6003 | 0.567 | 0.068 | 6.50E-07 | 9 |
| Flot2 | 0.6 | 0.765 | 0.359 | 1 | 10 |
| Calm3 | 0.5995 | 1 | 0.988 | 0.0078 | 10 |
| Slc20a1 | 0.5995 | 0.765 | 0.252 | 1 | 10 |
| Stk32a | 0.5985 | 0.849 | 0.344 | 1.84E-26 | 2 |
| Cdh2 | 0.5979 | 0.984 | 0.75 | 6.87E-14 | 3 |
| Ddx3x | 0.5978 | 0.941 | 0.43 | 1 | 10 |
| Gabra3 | 0.5971 | 0.824 | 0.32 | 1 | 10 |
| Cyp46a1 | 0.5967 | 0.765 | 0.361 | 1 | 10 |
| Chn1 | 0.5964 | 0.941 | 0.673 | 1 | 10 |
| Calm1 | 0.5962 | 1 | 1 | 3.24E-28 | 5 |
| Slc24a2 | 0.5959 | 0.824 | 0.318 | 1 | 10 |
| Pde1a | 0.5943 | 0.824 | 0.221 | 1 | 10 |
| Kcnab1 | 0.5939 | 0.588 | 0.076 | 1 | 10 |
| Nkd1 | 0.5935 | 0.505 | 0.099 | 6.57E-18 | 5 |
| Csrnp3 | 0.5933 | 0.941 | 0.341 | 1 | 10 |
| Fyn | 0.5928 | 0.765 | 0.37 | 1 | 10 |
| Vps26b | 0.5908 | 0.765 | 0.292 | 1 | 10 |
| Rab28 | 0.5907 | 0.925 | 0.512 | 1.83E-22 | 2 |
| Ecel1 | 0.5906 | 0.513 | 0.185 | 1.01E-10 | 1 |
| Foxp2 | 0.5901 | 0.592 | 0.039 | 2.40E-22 | 8 |
| Ngfrap1 | 0.5899 | 1 | 0.992 | 1.81E-22 | 7 |
| Msi2 | 0.5898 | 0.979 | 0.607 | 1.16E-28 | 2 |
| Chst8 | 0.5895 | 0.561 | 0.274 | 6.73E-14 | 3 |
| Pbx1 | 0.5893 | 0.973 | 0.732 | 4.10E-34 | 2 |
| Apc2 | 0.5892 | 0.941 | 0.444 | 0.378 | 10 |
| Adgrb2 | 0.5892 | 0.882 | 0.439 | 1 | 10 |
| Avpi1 | 0.589 | 0.615 | 0.297 | 3.52E-09 | 7 |
| mt-Co2 | 0.589 | 1 | 1 | 0.0033 | 10 |
| Rarb | 0.5886 | 0.577 | 0.042 | 2.28E-28 | 8 |
| Fxyd7 | 0.5884 | 0.9 | 0.658 | 0.0068 | 9 |
| Oxr1 | 0.5879 | 0.667 | 0.363 | 0.0809 | 9 |
| Prkce | 0.5869 | 1 | 0.567 | 1 | 10 |
| Elmo1 | 0.5861 | 0.718 | 0.241 | 9.28E-09 | 8 |
| Myt1l | 0.5859 | 1 | 0.689 | 1 | 10 |
| Ubl3 | 0.5858 | 0.718 | 0.377 | 5.15E-12 | 7 |
| Igsf1 | 0.5858 | 0.647 | 0.1 | 0.1162 | 10 |
| Ncald | 0.5853 | 0.917 | 0.654 | 1.97E-17 | 4 |
| Dlgap1 | 0.5853 | 0.833 | 0.389 | 0.0018 | 9 |

| | | | | | |
|---|---|---|---|---|---|
| Dctn1 | 0.5852 | 1 | 0.714 | 1 | 10 |
| Syt11 | 0.5846 | 1 | 0.941 | 0.3557 | 10 |
| 5031439 | 0.5845 | 0.824 | 0.328 | 1 | 10 |
| Rgs11 | 0.5844 | 0.765 | 0.161 | 0.0523 | 10 |
| Pcdh19 | 0.584 | 0.824 | 0.522 | 1 | 10 |
| Calb2 | 0.5838 | 0.866 | 0.552 | 1.04E-07 | 6 |
| Gpd1l | 0.5835 | 0.765 | 0.149 | 0.1229 | 10 |
| Kcnip4 | 0.5833 | 0.733 | 0.213 | 0.0102 | 9 |
| Cadps2 | 0.5831 | 0.936 | 0.524 | 6.35E-13 | 7 |
| Ppm1l | 0.583 | 0.706 | 0.369 | 1 | 10 |
| Tmem158 | 0.5826 | 0.882 | 0.687 | 2.41E-16 | 1 |
| Cacna1d | 0.5825 | 0.756 | 0.498 | 1.24E-13 | 3 |
| Mfn2 | 0.5822 | 0.941 | 0.5 | 1 | 10 |
| Per3 | 0.582 | 0.863 | 0.545 | 1.10E-27 | 2 |
| Tmem56 | 0.5818 | 0.706 | 0.123 | 1 | 10 |
| Prrc2b | 0.5818 | 0.824 | 0.715 | 1 | 10 |
| mt-Atp6 | 0.5816 | 1 | 1 | 0.0877 | 10 |
| Coro1a | 0.5816 | 0.765 | 0.203 | 0.0134 | 10 |
| Epn2 | 0.5814 | 0.824 | 0.343 | 1 | 10 |
| Arl8b | 0.581 | 0.824 | 0.402 | 1 | 10 |
| Atp2b1 | 0.5806 | 1 | 0.825 | 8.02E-09 | 9 |
| Lef1 | 0.5806 | 0.549 | 0.042 | 5.14E-21 | 8 |
| Ldha | 0.5799 | 0.958 | 0.747 | 3.89E-10 | 8 |
| Malat1 | 0.5798 | 1 | 0.999 | 2.64E-25 | 5 |
| Rftn1 | 0.5795 | 0.825 | 0.349 | 2.55E-19 | 5 |
| Pcdh17 | 0.5792 | 0.934 | 0.598 | 1.31E-22 | 1 |
| Rpusd1 | 0.5782 | 0.765 | 0.23 | 1 | 10 |
| Camta2 | 0.5782 | 0.765 | 0.238 | 1 | 10 |
| Plxna2 | 0.5782 | 0.824 | 0.198 | 1 | 10 |
| Pitpnc1 | 0.5782 | 0.765 | 0.138 | 1 | 10 |
| Tob1 | 0.5781 | 0.808 | 0.386 | 1.77E-14 | 7 |
| C77370 | 0.5778 | 0.941 | 0.357 | 1 | 10 |
| Camk4 | 0.5777 | 0.647 | 0.153 | 1 | 10 |
| Cmip | 0.577 | 1 | 0.944 | 0.2517 | 10 |
| Arhgap6 | 0.5769 | 0.925 | 0.558 | 4.22E-30 | 2 |
| Vps13a | 0.5769 | 0.882 | 0.323 | 1 | 10 |
| Pfn1 | 0.5768 | 0.979 | 0.793 | 3.64E-28 | 2 |
| Pja2 | 0.5768 | 1 | 0.894 | 1 | 10 |
| Lypd1 | 0.5752 | 0.487 | 0.171 | 6.15E-13 | 1 |
| Rab9b | 0.575 | 0.824 | 0.287 | 1 | 10 |
| Zmiz2 | 0.575 | 0.706 | 0.328 | 1 | 10 |
| Camk2d | 0.575 | 0.8 | 0.351 | 0.1181 | 9 |

| | | | | | |
|---|---|---|---|---|---|
| Ptms | 0.5749 | 1 | 0.967 | 2.10E-35 | 2 |
| Usp9x | 0.5745 | 1 | 0.808 | 1 | 10 |
| Rnf112 | 0.5728 | 0.941 | 0.565 | 1 | 10 |
| Grid2 | 0.5727 | 0.765 | 0.228 | 1 | 10 |
| Cds2 | 0.5721 | 0.941 | 0.684 | 1 | 10 |
| Zfhx3 | 0.5721 | 0.986 | 0.693 | 9.85E-33 | 2 |
| Cacna1h | 0.5716 | 0.647 | 0.131 | 1 | 10 |
| Lsamp | 0.5714 | 0.981 | 0.829 | 5.73E-19 | 4 |
| Kpna6 | 0.5712 | 0.882 | 0.221 | 1 | 10 |
| Entpd6 | 0.5705 | 0.824 | 0.414 | 1 | 10 |
| Rab6b | 0.5689 | 0.967 | 0.794 | 5.11E-06 | 9 |
| Zfand5 | 0.5683 | 0.9 | 0.728 | 0.0087 | 9 |
| Klhl13 | 0.5675 | 0.882 | 0.315 | 0.3535 | 10 |
| Tox2 | 0.5671 | 0.789 | 0.424 | 1.70E-10 | 8 |
| Nsf | 0.567 | 0.955 | 0.81 | 8.75E-22 | 0 |
| Dclk1 | 0.5668 | 0.882 | 0.7 | 1 | 10 |
| Ftl1 | 0.5662 | 1 | 0.95 | 2.01E-13 | 7 |
| Zfhx3 | 0.566 | 0.969 | 0.708 | 1.06E-15 | 5 |
| Lhx6 | 0.5654 | 0.533 | 0.003 | 3.35E-19 | 9 |
| Abca2 | 0.5651 | 0.941 | 0.493 | 1 | 10 |
| Megf11 | 0.5647 | 0.647 | 0.095 | 0.7292 | 10 |
| Cacna2d3 | 0.5647 | 0.642 | 0.395 | 1.72E-11 | 3 |
| Fam49a | 0.5646 | 0.529 | 0.134 | 1 | 10 |
| Agap3 | 0.5644 | 1 | 0.662 | 0.7208 | 10 |
| 1700086l | 0.5639 | 0.603 | 0.354 | 0.0002 | 7 |
| Adamts2 | 0.5634 | 0.588 | 0.025 | 0.2979 | 10 |
| Crebl2 | 0.5628 | 0.824 | 0.271 | 1 | 10 |
| Ppfia2 | 0.5625 | 0.87 | 0.538 | 1.39E-18 | 3 |
| Meis2 | 0.5625 | 0.08 | 0 | 1.58E-09 | 0 |
| Tsc22d1 | 0.562 | 1 | 0.848 | 1.07E-20 | 4 |
| Pnkd | 0.5618 | 0.824 | 0.45 | 1 | 10 |
| Mtfp1 | 0.5618 | 0.867 | 0.38 | 0.0002 | 9 |
| Parm1 | 0.5614 | 0.805 | 0.56 | 8.56E-11 | 3 |
| Paqr9 | 0.5601 | 0.824 | 0.303 | 0.215 | 10 |
| Flrt2 | 0.5601 | 0.529 | 0.224 | 1 | 10 |
| Fam46a | 0.5597 | 0.461 | 0.146 | 7.60E-17 | 1 |
| Nudt3 | 0.5585 | 0.941 | 0.583 | 1 | 10 |
| Dynll2 | 0.5584 | 1 | 0.917 | 1 | 10 |
| Cd99l2 | 0.5574 | 0.882 | 0.631 | 1 | 10 |
| Rnf138rt1 | 0.5573 | 0.493 | 0.172 | 3.28E-08 | 8 |
| Epha5 | 0.557 | 1 | 0.848 | 0.1674 | 10 |
| Tmem9b | 0.5568 | 0.941 | 0.405 | 0.5697 | 10 |

| | | | | | |
|---|---|---|---|---|---|
| Chga | 0.5568 | 0.941 | 0.731 | 0.0172 | 10 |
| D430019 | 0.5564 | 0.765 | 0.318 | 1 | 10 |
| Myh10 | 0.5562 | 0.941 | 0.482 | 1 | 10 |
| Tango2 | 0.5557 | 0.824 | 0.407 | 1 | 10 |
| Fyn | 0.5556 | 0.8 | 0.365 | 0.0119 | 9 |
| Sptbn4 | 0.5555 | 0.765 | 0.285 | 1 | 10 |
| Tle4 | 0.5555 | 0.897 | 0.653 | 6.20E-11 | 7 |
| Tmem65 | 0.5554 | 0.765 | 0.153 | 0.5728 | 10 |
| Pvrl3 | 0.5552 | 0.602 | 0.339 | 4.45E-15 | 3 |
| Baiap3 | 0.555 | 1 | 0.903 | 0.0911 | 10 |
| Clu | 0.5548 | 0.868 | 0.562 | 2.02E-17 | 1 |
| mt-Nd5 | 0.5548 | 1 | 0.978 | 0.0897 | 10 |
| Timp2 | 0.5548 | 1 | 0.892 | 1 | 10 |
| Prrg3 | 0.5548 | 0.706 | 0.17 | 1 | 10 |
| Scap | 0.5545 | 0.588 | 0.24 | 1 | 10 |
| Gsta4 | 0.5542 | 0.647 | 0.167 | 1 | 10 |
| Dbp | 0.5536 | 0.849 | 0.622 | 5.12E-14 | 2 |
| Tmem10( | 0.5535 | 0.941 | 0.511 | 1 | 10 |
| Miat | 0.5531 | 0.765 | 0.549 | 1 | 10 |
| Prkca | 0.5528 | 0.722 | 0.297 | 1.63E-17 | 4 |
| Fam81a | 0.5526 | 0.5 | 0.154 | 0.1108 | 9 |
| Impdh1 | 0.5523 | 0.647 | 0.232 | 1 | 10 |
| Tnpo2 | 0.552 | 0.647 | 0.2 | 1 | 10 |
| Ncoa1 | 0.552 | 0.765 | 0.336 | 1 | 10 |
| Nos1ap | 0.552 | 0.8 | 0.36 | 0.0009 | 9 |
| Cplx2 | 0.5513 | 0.53 | 0.213 | 1.65E-22 | 0 |
| Calb1 | 0.5509 | 0.889 | 0.75 | 6.49E-09 | 4 |
| Ramp2 | 0.5508 | 0.433 | 0.05 | 3.15E-06 | 9 |
| Plcb4 | 0.5507 | 0.778 | 0.486 | 4.72E-12 | 4 |
| Mllt11 | 0.5506 | 1 | 0.934 | 2.33E-21 | 2 |
| Cdh4 | 0.5502 | 0.567 | 0.165 | 0.0332 | 9 |
| Ghitm | 0.5498 | 1 | 0.633 | 1 | 10 |
| A830039I | 0.5496 | 0.765 | 0.18 | 1 | 10 |
| Zbtb7a | 0.5496 | 0.765 | 0.408 | 1 | 10 |
| BC04854( | 0.549 | 0.471 | 0.061 | 1 | 10 |
| Add1 | 0.5489 | 0.882 | 0.434 | 1 | 10 |
| Nefl | 0.5489 | 0.704 | 0.211 | 1.01E-09 | 8 |
| Arpp21 | 0.5486 | 0.824 | 0.374 | 1 | 10 |
| Fhad1 | 0.5486 | 0.69 | 0.189 | 3.30E-10 | 8 |
| Samd4 | 0.5485 | 0.588 | 0.151 | 1 | 10 |
| Eps15 | 0.548 | 0.765 | 0.302 | 1 | 10 |
| Kctd4 | 0.5478 | 0.529 | 0.042 | 1 | 10 |

| | | | | | |
|---|---|---|---|---|---|
| Usp31 | 0.5477 | 0.706 | 0.154 | 1 | 10 |
| Flrt3 | 0.5472 | 0.837 | 0.513 | 1.67E-16 | 3 |
| Wbp5 | 0.5471 | 0.987 | 0.941 | 1.19E-14 | 7 |
| Ano3 | 0.5465 | 0.765 | 0.063 | 0.0122 | 10 |
| Nefm | 0.5465 | 0.588 | 0.068 | 0.1744 | 10 |
| Stmn1 | 0.5465 | 1 | 0.999 | 3.34E-20 | 6 |
| Rpl27 | 0.5451 | 1 | 0.906 | 7.75E-23 | 2 |
| Pde4b | 0.5451 | 0.765 | 0.271 | 1 | 10 |
| H3f3a | 0.5443 | 1 | 0.945 | 1.11E-34 | 2 |
| Krt1 | 0.5437 | 0.704 | 0.258 | 1.09E-11 | 8 |
| Pitpnm1 | 0.5436 | 0.824 | 0.446 | 0.9017 | 10 |
| Ak5 | 0.5434 | 0.824 | 0.123 | 1 | 10 |
| Malat1 | 0.5432 | 1 | 0.999 | 3.53E-16 | 7 |
| Camta1 | 0.5427 | 1 | 0.933 | 1 | 10 |
| Kcnab2 | 0.5421 | 0.588 | 0.109 | 1 | 10 |
| Rrad | 0.5416 | 0.361 | 0.149 | 0.0002 | 4 |
| Nfe2l1 | 0.5413 | 0.941 | 0.771 | 1 | 10 |
| Tsnax | 0.5411 | 0.925 | 0.578 | 8.10E-17 | 2 |
| Enpp2 | 0.5411 | 0.421 | 0.083 | 1.25E-22 | 1 |
| Ppp3ca | 0.5405 | 0.933 | 0.79 | 0.2376 | 9 |
| Phpt1 | 0.5404 | 0.918 | 0.544 | 1.76E-19 | 2 |
| 1700086 | 0.5402 | 0.718 | 0.348 | 4.07E-07 | 8 |
| Cadps2 | 0.5401 | 0.925 | 0.496 | 3.06E-27 | 2 |
| Adh5 | 0.54 | 0.567 | 0.348 | 0.5 | 5 |
| Ttll5 | 0.5394 | 0.671 | 0.19 | 4.89E-23 | 2 |
| Kif1a | 0.5387 | 1 | 0.916 | 0.2048 | 10 |
| Pdyn | 0.5383 | 0.588 | 0.042 | 0.1423 | 10 |
| Cntn3 | 0.5375 | 0.471 | 0.049 | 1 | 10 |
| Helz | 0.5367 | 0.882 | 0.423 | 1 | 10 |
| Atg9a | 0.5365 | 0.882 | 0.33 | 0.3311 | 10 |
| Prkce | 0.5365 | 0.733 | 0.569 | 0.0961 | 9 |
| Pik3r1 | 0.5361 | 0.647 | 0.243 | 1 | 10 |
| Map2 | 0.536 | 1 | 0.896 | 1 | 10 |
| Ucp2 | 0.5358 | 0.746 | 0.302 | 5.90E-07 | 8 |
| Rgs10 | 0.5352 | 0.605 | 0.255 | 7.34E-12 | 1 |
| Btg1 | 0.5351 | 0.986 | 0.746 | 7.81E-24 | 2 |
| Ralgapb | 0.5348 | 0.765 | 0.248 | 1 | 10 |
| Rps6ka2 | 0.5348 | 0.706 | 0.238 | 1 | 10 |
| Auts2 | 0.5341 | 0.951 | 0.712 | 4.73E-14 | 3 |
| Lgi3 | 0.5341 | 0.706 | 0.194 | 1 | 10 |
| Syt6 | 0.534 | 0.417 | 0.096 | 1.12E-15 | 4 |
| Slc22a17 | 0.534 | 1 | 0.966 | 9.00E-06 | 10 |

| | | | | | |
|---|---|---|---|---|---|
| Pou2f2 | 0.5336 | 0.859 | 0.582 | 1.24E-09 | 7 |
| Rorb | 0.5333 | 0.918 | 0.647 | 4.55E-08 | 5 |
| Spon1 | 0.5331 | 0.65 | 0.38 | 5.09E-14 | 3 |
| Egr1 | 0.5329 | 0.577 | 0.308 | 0.0256 | 8 |
| Phactr1 | 0.5329 | 0.824 | 0.345 | 1 | 10 |
| L1cam | 0.5321 | 0.941 | 0.5 | 0.6855 | 10 |
| Slc24a5 | 0.5319 | 0.732 | 0.544 | 1.16E-07 | 3 |
| Calr | 0.5318 | 1 | 0.829 | 1 | 10 |
| Kcnk2 | 0.5317 | 0.535 | 0.323 | 5.45E-11 | 0 |
| Zbtb43 | 0.5317 | 0.765 | 0.123 | 0.005 | 10 |
| Sparcl1 | 0.5307 | 0.602 | 0.126 | 2.31E-12 | 4 |
| Spred1 | 0.5307 | 0.824 | 0.271 | 1 | 10 |
| Zdhhc3 | 0.5302 | 0.882 | 0.302 | 1 | 10 |
| Mast3 | 0.5299 | 0.706 | 0.238 | 1 | 10 |
| Elp3 | 0.5295 | 0.824 | 0.295 | 1 | 10 |
| Celf1 | 0.5295 | 0.941 | 0.548 | 1 | 10 |
| Kndc1 | 0.5294 | 0.706 | 0.255 | 1 | 10 |
| Git1 | 0.5291 | 0.706 | 0.225 | 1 | 10 |
| Clptm1 | 0.529 | 0.941 | 0.535 | 1 | 10 |
| Atxn7l3 | 0.5285 | 0.882 | 0.328 | 1 | 10 |
| Cntnap4 | 0.5283 | 0.588 | 0.162 | 1 | 10 |
| Baiap3 | 0.5277 | 1 | 0.893 | 0.001 | 3 |
| Ctsb | 0.5271 | 1 | 0.922 | 0.0743 | 10 |
| Tmsb4x | 0.5271 | 1 | 0.994 | 4.59E-07 | 8 |
| Pnrc1 | 0.5263 | 0.897 | 0.709 | 2.21E-19 | 2 |
| Arhgef17 | 0.526 | 0.824 | 0.466 | 1 | 10 |
| Plk2 | 0.5252 | 0.41 | 0.159 | 5.75E-09 | 7 |
| Ring1 | 0.5251 | 0.748 | 0.615 | 1.85E-06 | 3 |
| Atp6ap1 | 0.5246 | 0.941 | 0.789 | 1 | 10 |
| Slc9a6 | 0.5243 | 0.824 | 0.441 | 1 | 10 |
| mt-Cytb | 0.5243 | 1 | 1 | 1 | 10 |
| Tenm3 | 0.5239 | 0.715 | 0.516 | 2.25E-08 | 3 |
| Sst | 0.5233 | 0.06 | 0.011 | 1 | 0 |
| Hnrnpul2 | 0.5231 | 0.882 | 0.447 | 1 | 10 |
| Ifi27l2a | 0.5224 | 0.235 | 0.007 | 1 | 10 |
| Arxes2 | 0.5222 | 0.882 | 0.861 | 1 | 10 |
| Cst3 | 0.5213 | 1 | 0.988 | 1.66E-20 | 1 |
| Sh3kbp1 | 0.5211 | 0.602 | 0.349 | 6.90E-11 | 4 |
| Nnat | 0.5211 | 0.979 | 0.882 | 6.77E-10 | 5 |
| Phka2 | 0.5205 | 0.629 | 0.194 | 2.52E-15 | 5 |
| Hmgcr | 0.5196 | 0.886 | 0.672 | 2.97E-05 | 3 |
| Cdk17 | 0.5193 | 0.647 | 0.246 | 1 | 10 |

| | | | | | |
|---|---|---|---|---|---|
| Rnf208 | 0.5192 | 1 | 0.696 | 1 | 10 |
| Map3k7 | 0.5192 | 0.588 | 0.19 | 1 | 10 |
| Lmo4 | 0.5187 | 0.634 | 0.219 | 4.87E-08 | 8 |
| Dhcr24 | 0.5179 | 0.824 | 0.318 | 1 | 10 |
| Cbln2 | 0.5175 | 0.658 | 0.213 | 3.22E-17 | 2 |
| Nxph1 | 0.5173 | 0.76 | 0.324 | 1.10E-16 | 2 |
| Atl1 | 0.5172 | 0.941 | 0.432 | 1 | 10 |
| Grik4 | 0.517 | 0.588 | 0.1 | 1 | 10 |
| Kcnip1 | 0.5166 | 0.634 | 0.141 | 3.26E-12 | 8 |
| Atp2b2 | 0.5161 | 0.935 | 0.702 | 5.25E-07 | 3 |
| Elmo1 | 0.5158 | 0.533 | 0.264 | 1 | 9 |
| Npnt | 0.5158 | 0.471 | 0.034 | 0.0659 | 10 |
| Elmo1 | 0.5155 | 0.612 | 0.218 | 3.09E-16 | 1 |
| Per3 | 0.5148 | 0.764 | 0.564 | 1.13E-06 | 3 |
| Arc | 0.5148 | 0.346 | 0.096 | 0.0013 | 7 |
| Lonrf2 | 0.5139 | 0.803 | 0.547 | 3.77E-17 | 1 |
| Cd24a | 0.5132 | 0.821 | 0.447 | 1.65E-18 | 3 |
| Disp2 | 0.5126 | 1 | 0.833 | 1 | 10 |
| Lmtk2 | 0.5121 | 0.706 | 0.113 | 0.057 | 10 |
| Actr1a | 0.5114 | 0.824 | 0.43 | 1 | 10 |
| Cd47 | 0.5107 | 0.984 | 0.86 | 1.35E-09 | 3 |
| Nudt11 | 0.5105 | 0.829 | 0.384 | 8.45E-12 | 2 |
| Syt13 | 0.5104 | 0.941 | 0.666 | 1 | 10 |
| Tspan7 | 0.5093 | 0.99 | 0.817 | 4.67E-20 | 0 |
| Nkd2 | 0.509 | 0.353 | 0.021 | 0.4779 | 10 |
| Atp5o | 0.5089 | 0.9 | 0.872 | 1 | 9 |
| Timp2 | 0.5087 | 0.965 | 0.878 | 4.59E-18 | 0 |
| Pak3 | 0.5087 | 0.933 | 0.827 | 0.0219 | 9 |
| Slc38a10 | 0.5081 | 0.706 | 0.294 | 1 | 10 |
| Hmgb3 | 0.5079 | 0.849 | 0.554 | 1.51E-12 | 2 |
| Pcp4 | 0.5079 | 0.967 | 0.867 | 0.0013 | 1 |
| Ssbp4 | 0.5078 | 1 | 0.864 | 0.0161 | 10 |
| Mrps6 | 0.5075 | 0.773 | 0.437 | 3.52E-12 | 5 |
| Kif5c | 0.507 | 1 | 0.802 | 1.55E-05 | 9 |
| Snhg20 | 0.5069 | 0.959 | 0.814 | 0.0172 | 3 |
| Idh1 | 0.506 | 0.765 | 0.249 | 1 | 10 |
| Ptprj | 0.5057 | 0.588 | 0.117 | 1 | 10 |
| Rasgrf1 | 0.5052 | 0.967 | 0.788 | 1.36E-06 | 3 |
| Rorb | 0.5046 | 0.875 | 0.638 | 5.98E-12 | 1 |
| Junb | 0.5034 | 0.549 | 0.322 | 1 | 8 |
| Klhl5 | 0.5032 | 0.706 | 0.083 | 0.3207 | 10 |
| Kcnc3 | 0.5028 | 0.706 | 0.17 | 0.4129 | 10 |

| | | | | | |
|---|---|---|---|---|---|
| Sema6b | 0.5026 | 0.824 | 0.333 | 1 | 10 |
| Oxct1 | 0.5021 | 0.941 | 0.659 | 1 | 10 |
| Tmem163 | 0.5017 | 0.463 | 0.085 | 8.06E-18 | 4 |
| Tceal8 | 0.5016 | 0.731 | 0.386 | 9.64E-11 | 7 |
| Stk32a | 0.5015 | 0.769 | 0.383 | 4.76E-10 | 7 |
| Lynx1 | 0.5015 | 0.533 | 0.157 | 0.2494 | 9 |
| Tmem132 | 0.5009 | 0.512 | 0.252 | 1.86E-13 | 3 |
| Nppc | 0.5008 | 0.3 | 0.086 | 1 | 9 |
| Map1b | 0.5006 | 1 | 0.978 | 1 | 10 |
| Hlf | 0.5002 | 0.756 | 0.506 | 4.19E-07 | 7 |
| Btg2 | 0.5 | 0.577 | 0.271 | 0.1411 | 8 |
| Hmx2 | 0.5 | 0.48 | 0.085 | 2.67E-24 | 1 |
| Nrxn1 | 0.4993 | 1 | 0.921 | 1 | 10 |
| Ssr1 | 0.4989 | 0.984 | 0.749 | 2.14E-08 | 3 |
| Hmgn2 | 0.4986 | 0.993 | 0.891 | 2.90E-25 | 2 |
| Kit | 0.4985 | 0.577 | 0.279 | 3.44E-13 | 3 |
| Gm36266 | 0.4982 | 0.322 | 0.067 | 2.12E-10 | 2 |
| Nfasc | 0.4977 | 0.882 | 0.504 | 1 | 10 |
| Tacc1 | 0.4976 | 0.824 | 0.283 | 1 | 10 |
| Ylpm1 | 0.4973 | 1 | 0.397 | 1 | 10 |
| Dock3 | 0.4971 | 0.882 | 0.377 | 1 | 10 |
| Ap1s2 | 0.4964 | 0.761 | 0.6 | 0.0804 | 8 |
| Lhx1os | 0.496 | 0.938 | 0.506 | 4.75E-12 | 6 |
| Myrip | 0.4958 | 0.412 | 0.034 | 1 | 10 |
| 1810043 | 0.4957 | 0.773 | 0.404 | 7.98E-11 | 5 |
| Apoc3 | 0.4956 | 0.521 | 0.091 | 7.25E-09 | 8 |
| Dlx1 | 0.4947 | 0.775 | 0.267 | 2.84E-08 | 8 |
| Usp22 | 0.4947 | 1 | 0.837 | 1 | 10 |
| Pbx1 | 0.4944 | 0.984 | 0.736 | 1.36E-12 | 3 |
| Mapk10 | 0.4943 | 1 | 0.608 | 1 | 10 |
| Fam81a | 0.4939 | 0.588 | 0.157 | 1 | 10 |
| Lrrtm3 | 0.4935 | 0.675 | 0.482 | 2.74E-10 | 3 |
| Wfs1 | 0.4935 | 0.572 | 0.277 | 9.19E-14 | 1 |
| Gpr12 | 0.4933 | 0.588 | 0.093 | 0.1411 | 10 |
| Rnf7 | 0.4931 | 0.993 | 0.883 | 2.83E-27 | 2 |
| Zdhhc17 | 0.493 | 0.824 | 0.361 | 1 | 10 |
| Xpr1 | 0.4929 | 0.824 | 0.538 | 1 | 10 |
| Plppr2 | 0.4928 | 0.882 | 0.481 | 0.8437 | 10 |
| Dner | 0.4922 | 0.972 | 0.824 | 3.92E-11 | 8 |
| Ttc14 | 0.492 | 0.911 | 0.719 | 0.0001 | 3 |
| Id4 | 0.4915 | 0.948 | 0.614 | 9.09E-11 | 5 |
| Ncam1 | 0.491 | 1 | 0.946 | 1 | 3 |

| | | | | | |
|---|---|---|---|---|---|
| mt-Atp8 | 0.4909 | 0.992 | 0.957 | 1 | 3 |
| Adcy5 | 0.4907 | 0.824 | 0.328 | 1 | 10 |
| Tenm1 | 0.4907 | 0.595 | 0.337 | 2.29E-12 | 0 |
| Gpat4 | 0.4907 | 0.765 | 0.276 | 1 | 10 |
| Amigo2 | 0.4906 | 0.546 | 0.206 | 1.13E-13 | 4 |
| A230065I | 0.4902 | 0.493 | 0.136 | 0.0001 | 8 |
| Gdpd2 | 0.4901 | 0.706 | 0.092 | 0.5157 | 10 |
| H1fx | 0.4896 | 0.925 | 0.703 | 1.40E-12 | 2 |
| Adar | 0.4894 | 0.941 | 0.465 | 1 | 10 |
| Rbfox3 | 0.4891 | 0.733 | 0.398 | 0.1038 | 9 |
| Edil3 | 0.4888 | 0.797 | 0.64 | 0.0002 | 3 |
| Tusc3 | 0.4887 | 1 | 0.965 | 9.76E-05 | 3 |
| Aldoa | 0.4878 | 1 | 1 | 6.16E-18 | 5 |
| Atp2b4 | 0.4877 | 0.42 | 0.24 | 1.25E-09 | 0 |
| Tead1 | 0.4874 | 0.704 | 0.287 | 1.02E-08 | 8 |
| Plxdc1 | 0.4873 | 0.647 | 0.151 | 1 | 10 |
| Opcml | 0.487 | 0.861 | 0.657 | 1.17E-07 | 4 |
| Scn1a | 0.487 | 0.525 | 0.229 | 3.70E-16 | 0 |
| Stim2 | 0.4867 | 0.567 | 0.228 | 0.2253 | 9 |
| Mtmr3 | 0.4866 | 0.765 | 0.314 | 1 | 10 |
| Gm15261 | 0.4865 | 0.712 | 0.152 | 1.30E-30 | 2 |
| Stat3 | 0.4865 | 0.588 | 0.187 | 1 | 10 |
| Hid1 | 0.4862 | 0.706 | 0.251 | 1 | 10 |
| Gfra2 | 0.4859 | 0.732 | 0.361 | 1.07E-07 | 8 |
| Mgst3 | 0.4856 | 0.732 | 0.369 | 2.41E-07 | 8 |
| Leng8 | 0.4848 | 0.862 | 0.674 | 0.0194 | 3 |
| Gabrg2 | 0.4846 | 0.887 | 0.659 | 2.00E-09 | 8 |
| Tceal3 | 0.4844 | 1 | 0.941 | 5.25E-10 | 8 |
| Sulf2 | 0.484 | 0.563 | 0.127 | 5.30E-14 | 8 |
| Nfia | 0.484 | 0.333 | 0.029 | 0.0027 | 9 |
| Ywhaz | 0.4835 | 1 | 0.972 | 1 | 10 |
| C330006A | 0.4835 | 0.647 | 0.221 | 1 | 10 |
| mt-Nd1 | 0.4834 | 1 | 1 | 0.0034 | 10 |
| Fam19a1 | 0.4833 | 0.815 | 0.367 | 7.74E-24 | 2 |
| Peg10 | 0.4831 | 0.507 | 0.298 | 0.3626 | 2 |
| Aplp2 | 0.4829 | 0.9 | 0.694 | 9.38E-15 | 0 |
| Sulf1 | 0.4825 | 0.592 | 0.151 | 8.82E-09 | 8 |
| Itm2c | 0.4824 | 1 | 0.998 | 0.3886 | 3 |
| Lrrn2 | 0.4823 | 0.824 | 0.476 | 1 | 10 |
| Arf1 | 0.482 | 1 | 0.99 | 1 | 10 |
| Vps35 | 0.4819 | 0.882 | 0.603 | 1 | 10 |
| Gdi1 | 0.4818 | 1 | 0.97 | 0.4878 | 10 |

| | | | | | |
|---|---|---|---|---|---|
| 1110004I | 0.4818 | 0.99 | 0.961 | 1.33E-11 | 5 |
| Coq10b | 0.4817 | 0.628 | 0.268 | 5.59E-08 | 7 |
| Kbtbd11 | 0.4814 | 0.824 | 0.258 | 1 | 10 |
| Vwa5b2 | 0.4814 | 0.765 | 0.23 | 1 | 10 |
| Clu | 0.4811 | 0.86 | 0.547 | 2.45E-13 | 0 |
| Rps28 | 0.4808 | 1 | 0.988 | 2.96E-22 | 2 |
| Hectd3 | 0.4804 | 0.588 | 0.143 | 1 | 10 |
| PISD | 0.4803 | 0.959 | 0.818 | 1 | 3 |
| Ppp1r17 | 0.4802 | 0.69 | 0.282 | 8.37E-06 | 8 |
| Pvrl3 | 0.48 | 0.76 | 0.309 | 7.73E-20 | 2 |
| Ptpro | 0.4799 | 0.732 | 0.273 | 1.74E-08 | 8 |
| Sncb | 0.4797 | 0.967 | 0.916 | 1 | 9 |
| Gaa | 0.4796 | 1 | 0.974 | 0.0081 | 10 |
| Spock2 | 0.4791 | 1 | 0.856 | 1 | 10 |
| Slitrk1 | 0.479 | 0.706 | 0.284 | 1 | 10 |
| Slc4a8 | 0.4788 | 0.882 | 0.281 | 1 | 10 |
| Nos1 | 0.4785 | 0.435 | 0.089 | 7.65E-13 | 4 |
| Map6 | 0.4782 | 0.794 | 0.39 | 7.67E-12 | 5 |
| Dlx1 | 0.4781 | 0.605 | 0.251 | 3.70E-14 | 1 |
| Xpo7 | 0.4779 | 0.824 | 0.249 | 1 | 10 |
| Etv1 | 0.4774 | 0.4 | 0.047 | 0.0018 | 9 |
| Aifm3 | 0.4771 | 0.588 | 0.189 | 1 | 10 |
| Dpysl2 | 0.4771 | 1 | 0.959 | 2.02E-14 | 2 |
| Lrrc4b | 0.477 | 0.919 | 0.732 | 0.0004 | 3 |
| Cadm3 | 0.477 | 0.785 | 0.529 | 3.90E-14 | 0 |
| Chst1 | 0.4767 | 0.992 | 0.872 | 5.41E-08 | 3 |
| Neto2 | 0.4764 | 0.6 | 0.136 | 0.0004 | 9 |
| Cyc1 | 0.4754 | 0.933 | 0.829 | 0.0023 | 9 |
| Nr3c1 | 0.4751 | 0.602 | 0.252 | 1.50E-10 | 4 |
| Tsc22d1 | 0.4749 | 0.972 | 0.855 | 2.63E-08 | 8 |
| Rundc3b | 0.4745 | 0.882 | 0.248 | 1 | 10 |
| Cetn3 | 0.4742 | 0.89 | 0.632 | 2.69E-14 | 2 |
| Hprt | 0.4741 | 0.952 | 0.847 | 5.79E-11 | 2 |
| Gria3 | 0.474 | 0.667 | 0.19 | 0.1133 | 9 |
| Dlx1 | 0.4739 | 0.767 | 0.287 | 0.4066 | 9 |
| Pde1c | 0.4736 | 0.421 | 0.052 | 7.73E-26 | 1 |
| Rpl15 | 0.4734 | 1 | 0.967 | 7.15E-20 | 2 |
| Paip1 | 0.4731 | 0.795 | 0.379 | 6.40E-11 | 2 |
| Podxl2 | 0.4727 | 1 | 0.984 | 0.0396 | 3 |
| Dync1h1 | 0.4725 | 0.882 | 0.595 | 1 | 10 |
| Tmem179 | 0.4721 | 1 | 0.862 | 1 | 10 |
| Stim2 | 0.4717 | 0.765 | 0.229 | 1 | 10 |

| | | | | | |
|---|---|---|---|---|---|
| Camk2d | 0.4713 | 0.555 | 0.321 | 8.04E-15 | 0 |
| Chgb | 0.4712 | 0.944 | 0.847 | 6.63E-09 | 8 |
| Hspa5 | 0.471 | 1 | 0.886 | 1 | 3 |
| Pcdh19 | 0.471 | 0.699 | 0.505 | 1.32E-07 | 3 |
| Sncb | 0.4708 | 0.975 | 0.905 | 3.83E-18 | 0 |
| Jph3 | 0.4707 | 0.765 | 0.272 | 1 | 10 |
| Npas1 | 0.4706 | 0.267 | 0.012 | 0.0056 | 9 |
| Kcna1 | 0.4704 | 0.353 | 0.025 | 1 | 10 |
| Map7 | 0.47 | 0.824 | 0.237 | 1 | 10 |
| Rcn1 | 0.4699 | 0.602 | 0.299 | 1.11E-08 | 4 |
| Aldoc | 0.4698 | 0.505 | 0.257 | 5.47E-14 | 0 |
| Ildr2 | 0.4698 | 0.941 | 0.31 | 1 | 10 |
| Ccdc109b | 0.4693 | 0.397 | 0.097 | 3.16E-10 | 7 |
| Lrp1b | 0.4692 | 0.757 | 0.627 | 2.92E-07 | 1 |
| Auts2 | 0.4692 | 0.979 | 0.702 | 1.26E-31 | 2 |
| Ifi27 | 0.4691 | 0.567 | 0.103 | 0.0506 | 9 |
| Bcl7a | 0.4691 | 0.706 | 0.295 | 1 | 10 |
| Foxg1 | 0.4689 | 0.447 | 0.033 | 8.40E-33 | 1 |
| Lrtm2 | 0.4682 | 0.765 | 0.195 | 1 | 10 |
| Dnajc21 | 0.4681 | 0.633 | 0.362 | 1 | 9 |
| Abhd8 | 0.4679 | 0.986 | 0.859 | 1.52E-19 | 2 |
| Gabrq | 0.4672 | 0.583 | 0.173 | 5.35E-12 | 4 |
| Cirbp | 0.4668 | 1 | 0.97 | 1.26E-19 | 5 |
| Rictor | 0.4667 | 0.706 | 0.201 | 1 | 10 |
| Dock10 | 0.4665 | 0.5 | 0.126 | 0.0012 | 9 |
| Hrk | 0.4663 | 0.706 | 0.164 | 1 | 10 |
| Cpd | 0.4663 | 0.706 | 0.172 | 1 | 10 |
| Pcdh9 | 0.4662 | 0.797 | 0.583 | 9.41E-06 | 3 |
| Nxph1 | 0.4661 | 0.585 | 0.355 | 1.87E-12 | 3 |
| Gpx1 | 0.4656 | 0.885 | 0.543 | 4.52E-09 | 7 |
| Sox1 | 0.4655 | 0.685 | 0.323 | 2.05E-09 | 2 |
| Crip2 | 0.4645 | 0.887 | 0.655 | 1.01E-06 | 8 |
| Pam | 0.4644 | 1 | 0.681 | 1 | 10 |
| Robo2 | 0.4644 | 0.647 | 0.108 | 1 | 10 |
| Tmem47 | 0.4644 | 0.941 | 0.362 | 1 | 10 |
| Vsnl1 | 0.4643 | 0.932 | 0.819 | 1.49E-06 | 2 |
| Calb2 | 0.4643 | 0.719 | 0.558 | 0.0037 | 2 |
| Lrp3 | 0.4642 | 0.765 | 0.341 | 1 | 10 |
| Prpf19 | 0.4642 | 0.824 | 0.481 | 1 | 10 |
| Snx3 | 0.464 | 0.966 | 0.85 | 1.02E-15 | 2 |
| Got1 | 0.4639 | 0.967 | 0.845 | 0.1674 | 9 |
| Dgkk | 0.4637 | 0.479 | 0.11 | 6.65E-06 | 8 |

| | | | | | |
|---|---|---|---|---|---|
| Trp53i11 | 0.4635 | 0.951 | 0.696 | 1.85E-13 | 3 |
| Tbl1xr1 | 0.4634 | 0.588 | 0.152 | 1 | 10 |
| Hmgb3 | 0.463 | 0.845 | 0.568 | 9.68E-10 | 5 |
| Igsf8 | 0.4628 | 1 | 0.817 | 1 | 10 |
| Pdcd4 | 0.4628 | 0.718 | 0.67 | 0.1949 | 7 |
| Wnk2 | 0.4627 | 0.765 | 0.274 | 1 | 10 |
| Dlx6os1 | 0.4626 | 0.533 | 0.119 | 0.0135 | 9 |
| Slc6a17 | 0.4624 | 0.882 | 0.483 | 1 | 10 |
| Scn5a | 0.4623 | 0.588 | 0.056 | 1 | 10 |
| Trpc4ap | 0.4614 | 0.941 | 0.505 | 1 | 10 |
| Nenf | 0.4612 | 1 | 0.99 | 1.03E-15 | 1 |
| 5730409 | 0.4611 | 0.588 | 0.189 | 1 | 10 |
| Sox2 | 0.4608 | 0.856 | 0.551 | 8.43E-10 | 5 |
| Ptprz1 | 0.4608 | 0.625 | 0.394 | 8.97E-08 | 1 |
| Marf1 | 0.4602 | 0.863 | 0.543 | 3.45E-09 | 2 |
| Scn2a1 | 0.46 | 0.959 | 0.873 | 1 | 3 |
| Sidt1 | 0.4598 | 0.588 | 0.068 | 0.4984 | 10 |
| Rreb1 | 0.4598 | 0.412 | 0.058 | 1 | 10 |
| Gm1388 | 0.4596 | 0.586 | 0.333 | 5.38E-05 | 1 |
| Calm1 | 0.4592 | 1 | 1 | 4.10E-27 | 6 |
| Adam22 | 0.459 | 1 | 0.531 | 1 | 10 |
| Gstm6 | 0.4586 | 0.69 | 0.281 | 1.63E-06 | 8 |
| Nrxn2 | 0.4582 | 0.992 | 0.96 | 0.0746 | 3 |
| Gt(ROSA) | 0.4582 | 0.678 | 0.38 | 0.0002 | 2 |
| Tmtc1 | 0.4581 | 0.529 | 0.099 | 1 | 10 |
| Nrsn1 | 0.4581 | 1 | 0.957 | 0.0134 | 9 |
| Rnd3 | 0.4578 | 0.549 | 0.141 | 2.76E-09 | 8 |
| Thsd7b | 0.4578 | 0.561 | 0.335 | 6.28E-11 | 3 |
| Tmem63 | 0.4576 | 0.941 | 0.546 | 1 | 10 |
| Efna5 | 0.4573 | 0.618 | 0.259 | 1.84E-16 | 1 |
| 1110004 | 0.4572 | 1 | 0.958 | 9.88E-29 | 2 |
| Ogt | 0.457 | 0.984 | 0.865 | 0.006 | 3 |
| Herc3 | 0.4566 | 0.658 | 0.289 | 2.28E-17 | 1 |
| Enah | 0.4563 | 0.867 | 0.686 | 0.1782 | 9 |
| Arhgap6 | 0.4563 | 0.885 | 0.585 | 5.66E-08 | 7 |
| Ipo9 | 0.4561 | 0.765 | 0.263 | 1 | 10 |
| Atp6v0a2 | 0.4559 | 0.647 | 0.212 | 1 | 10 |
| Slc32a1 | 0.4558 | 0.976 | 0.827 | 1.38E-05 | 3 |
| 6030419 | 0.4557 | 0.588 | 0.136 | 1 | 10 |
| Mgat5 | 0.4557 | 0.824 | 0.342 | 1 | 10 |
| Arhgdia | 0.4557 | 1 | 0.784 | 1 | 10 |
| Dlk1 | 0.4556 | 0.876 | 0.691 | 4.42E-05 | 5 |

| | | | | | |
|---|---|---|---|---|---|
| Chchd10 | 0.4554 | 0.959 | 0.923 | 3.73E-06 | 5 |
| Camk2n1 | 0.455 | 0.925 | 0.734 | 9.40E-10 | 2 |
| Rasal1 | 0.4548 | 0.529 | 0.101 | 1 | 10 |
| Gnai1 | 0.4546 | 0.9 | 0.738 | 1 | 9 |
| Rps12 | 0.4544 | 0.93 | 0.825 | 0.0006 | 8 |
| Dgkg | 0.4541 | 0.941 | 0.262 | 1 | 10 |
| Ly6h | 0.4541 | 1 | 1 | 7.94E-16 | 3 |
| Chmp5 | 0.454 | 0.87 | 0.478 | 2.61E-11 | 2 |
| Npffr1 | 0.4539 | 0.412 | 0.066 | 1 | 10 |
| Krt10 | 0.4534 | 0.629 | 0.327 | 2.99E-09 | 5 |
| Grina | 0.4533 | 1 | 0.865 | 1 | 10 |
| Ptprs | 0.4531 | 0.984 | 0.832 | 1 | 3 |
| Gem | 0.453 | 0.352 | 0.047 | 1.54E-07 | 8 |
| Bex2 | 0.453 | 1 | 1 | 3.78E-16 | 7 |
| Lrrtm1 | 0.4522 | 0.706 | 0.163 | 1 | 10 |
| App | 0.452 | 0.985 | 0.918 | 1.91E-13 | 0 |
| Crocc | 0.4519 | 0.824 | 0.304 | 1 | 10 |
| Gng3 | 0.4508 | 1 | 0.986 | 4.07E-23 | 2 |
| Akirin1 | 0.4506 | 0.732 | 0.473 | 3.05E-07 | 5 |
| Wnt4 | 0.4505 | 0.367 | 0.081 | 0.0469 | 9 |
| C1ql2 | 0.4503 | 0.412 | 0.051 | 1 | 10 |
| Ddah1 | 0.4502 | 0.6 | 0.162 | 0.0338 | 9 |
| Eif1 | 0.4501 | 1 | 0.998 | 4.33E-14 | 7 |
| Spns1 | 0.4498 | 0.706 | 0.309 | 1 | 10 |
| Tmem25 | 0.4498 | 0.647 | 0.234 | 1 | 10 |
| Rbfox1 | 0.4497 | 0.6 | 0.199 | 1 | 9 |
| Cox5a | 0.4494 | 1 | 0.977 | 1 | 9 |
| Strip2 | 0.4491 | 0.353 | 0.013 | 0.1142 | 10 |
| Chst8 | 0.449 | 0.664 | 0.252 | 8.28E-18 | 2 |
| Pnmal1 | 0.4488 | 1 | 0.885 | 1 | 10 |
| Clcn3 | 0.4488 | 0.992 | 0.862 | 0.1602 | 3 |
| Gm6483 | 0.4485 | 0.911 | 0.691 | 0.0002 | 3 |
| Igsf8 | 0.4481 | 0.976 | 0.8 | 0.1322 | 3 |
| Cntn2 | 0.4477 | 0.559 | 0.19 | 5.08E-17 | 1 |
| Srm | 0.4474 | 0.667 | 0.364 | 1 | 9 |
| Myl6 | 0.4472 | 1 | 0.948 | 2.42E-06 | 6 |
| Trappc13 | 0.4468 | 0.753 | 0.349 | 2.12E-09 | 2 |
| Sox4 | 0.4467 | 0.567 | 0.325 | 1 | 9 |
| Opcml | 0.4466 | 0.915 | 0.66 | 9.79E-09 | 8 |
| Pgap1 | 0.4465 | 0.659 | 0.517 | 0.0007 | 3 |
| Cntn1 | 0.4464 | 0.796 | 0.616 | 8.22E-07 | 4 |
| Gpx4 | 0.4461 | 1 | 0.987 | 2.52E-14 | 7 |

| | | | | | |
|---|---|---|---|---|---|
| Zfr | 0.446 | 1 | 0.874 | 1 | 10 |
| Nkx2-1 | 0.4457 | 0.367 | 0.076 | 1 | 9 |
| Ost4 | 0.4448 | 0.842 | 0.53 | 1.11E-12 | 2 |
| Nptxr | 0.4445 | 0.824 | 0.569 | 1 | 10 |
| Wrnip1 | 0.4443 | 0.529 | 0.256 | 1 | 10 |
| Igfbp5 | 0.4442 | 0.918 | 0.601 | 9.63E-10 | 5 |
| Cntn5 | 0.4438 | 0.467 | 0.158 | 1 | 9 |
| Trp53i11 | 0.4437 | 0.938 | 0.692 | 4.27E-20 | 2 |
| mt-Nd6 | 0.4428 | 0.706 | 0.286 | 1 | 10 |
| Pcdh19 | 0.4426 | 0.821 | 0.504 | 1.31E-07 | 7 |
| Gria3 | 0.4423 | 0.455 | 0.148 | 3.69E-14 | 0 |
| Nov | 0.4423 | 0.361 | 0.103 | 1.91E-08 | 5 |
| Ppfibp1 | 0.4422 | 0.471 | 0.079 | 1 | 10 |
| Tppp3 | 0.4418 | 0.667 | 0.446 | 0.0031 | 7 |
| Tial1 | 0.4416 | 0.87 | 0.654 | 9.27E-07 | 3 |
| Unc5c | 0.4416 | 0.593 | 0.429 | 1.26E-08 | 3 |
| Vat1 | 0.4412 | 1 | 0.889 | 6.92E-09 | 8 |
| Omg | 0.4411 | 0.765 | 0.292 | 1 | 10 |
| Mzt1 | 0.4409 | 0.767 | 0.391 | 1.06E-07 | 2 |
| Rasa3 | 0.4409 | 0.647 | 0.143 | 1 | 10 |
| Enc1 | 0.4409 | 0.667 | 0.259 | 0.3001 | 9 |
| Tiparp | 0.4403 | 0.467 | 0.064 | 0.0029 | 9 |
| Gabra1 | 0.44 | 0.639 | 0.336 | 6.61E-06 | 4 |
| Sgcz | 0.4399 | 0.692 | 0.261 | 6.09E-15 | 2 |
| Tmsb10 | 0.4399 | 1 | 0.997 | 1.15E-11 | 7 |
| Trappc13 | 0.4398 | 0.742 | 0.37 | 5.62E-07 | 5 |
| Klf6 | 0.4396 | 0.487 | 0.209 | 4.23E-05 | 7 |
| Gpx3 | 0.4394 | 0.789 | 0.403 | 3.51E-05 | 8 |
| Cox17 | 0.4393 | 0.973 | 0.849 | 6.14E-15 | 2 |
| Cplx1 | 0.4393 | 0.36 | 0.131 | 3.82E-17 | 0 |
| Sema4f | 0.4389 | 0.706 | 0.223 | 1 | 10 |
| Kit | 0.4388 | 0.781 | 0.242 | 5.79E-25 | 2 |
| Cckbr | 0.4384 | 0.529 | 0.022 | 0.0356 | 10 |
| Pdha1 | 0.4384 | 0.767 | 0.365 | 0.1386 | 9 |
| Klf4 | 0.438 | 0.205 | 0.046 | 6.33E-05 | 7 |
| Ndufa8 | 0.4377 | 0.967 | 0.931 | 0.634 | 9 |
| Slc1a6 | 0.4375 | 0.765 | 0.201 | 1 | 10 |
| Tcf12 | 0.437 | 0.679 | 0.368 | 1.29E-06 | 7 |
| Sult4a1 | 0.4369 | 0.9 | 0.828 | 0.2631 | 9 |
| Slc6a6 | 0.4368 | 0.824 | 0.388 | 1 | 10 |
| Marcks | 0.4368 | 0.993 | 0.939 | 1.59E-17 | 2 |
| Trp53i11 | 0.4366 | 0.907 | 0.706 | 1.28E-08 | 5 |

| | | | | | |
|---|---|---|---|---|---|
| Wdfy3 | 0.4362 | 0.882 | 0.423 | 1 | 10 |
| Map1lc3a | 0.4361 | 1 | 0.969 | 1.32E-10 | 7 |
| Bub3 | 0.436 | 0.849 | 0.706 | 1.87E-10 | 1 |
| Gad1 | 0.436 | 0.795 | 0.757 | 1 | 7 |
| Cdk18 | 0.4359 | 0.6 | 0.204 | 0.4819 | 9 |
| Flot1 | 0.4358 | 1 | 0.554 | 0.4575 | 10 |
| Rps4x | 0.4354 | 0.986 | 0.982 | 0.0033 | 8 |
| Gnao1 | 0.4353 | 1 | 0.935 | 1 | 10 |
| Lrrc45 | 0.4353 | 0.706 | 0.269 | 1 | 10 |
| Olfm3 | 0.4352 | 0.454 | 0.177 | 5.14E-13 | 1 |
| Arpc1a | 0.4352 | 0.967 | 0.9 | 0.4449 | 9 |
| Nkain3 | 0.4351 | 0.461 | 0.179 | 2.28E-10 | 1 |
| Ccnd1 | 0.4348 | 0.417 | 0.068 | 7.20E-17 | 4 |
| Atp2a2 | 0.4346 | 0.92 | 0.768 | 5.55E-11 | 0 |
| Tcf4 | 0.434 | 0.539 | 0.268 | 1.75E-09 | 1 |
| Ddah1 | 0.434 | 0.706 | 0.165 | 1 | 10 |
| Tpm2 | 0.4338 | 0.031 | 0.004 | 1 | 6 |
| Sv2a | 0.4337 | 0.98 | 0.913 | 8.44E-16 | 0 |
| Fgf14 | 0.4337 | 0.68 | 0.312 | 2.74E-09 | 5 |
| Six3 | 0.4335 | 0.99 | 0.654 | 8.34E-09 | 6 |
| Serp2 | 0.4334 | 0.967 | 0.885 | 1 | 9 |
| Gm996 | 0.4333 | 0.647 | 0.203 | 1 | 10 |
| 4933431 | 0.4332 | 0.588 | 0.119 | 1 | 10 |
| Mum1 | 0.4331 | 0.629 | 0.31 | 5.40E-08 | 5 |
| Stmn1 | 0.433 | 1 | 0.999 | 2.16E-22 | 2 |
| Nudc | 0.4328 | 0.972 | 0.861 | 2.59E-06 | 8 |
| Crtac1 | 0.4325 | 0.461 | 0.174 | 5.02E-09 | 1 |
| As3mt | 0.4324 | 0.69 | 0.28 | 2.77E-05 | 8 |
| Sumo1 | 0.4323 | 0.993 | 0.939 | 8.91E-19 | 2 |
| Scg5 | 0.4322 | 1 | 0.975 | 1 | 3 |
| Gm2694 | 0.4314 | 0.529 | 0.03 | 7.45E-05 | 10 |
| Ina | 0.4312 | 0.941 | 0.765 | 1 | 10 |
| Acsl6 | 0.4312 | 0.706 | 0.226 | 1 | 10 |
| Zfp503 | 0.4311 | 0.41 | 0.055 | 1.34E-15 | 7 |
| Snhg11 | 0.4298 | 1 | 0.999 | 3.72E-17 | 3 |
| 6330403 | 0.4298 | 0.487 | 0.11 | 1.55E-18 | 1 |
| Rsrp1 | 0.4298 | 1 | 0.972 | 4.27E-09 | 3 |
| Pcdh20 | 0.4296 | 0.479 | 0.131 | 3.48E-14 | 2 |
| Plppr3 | 0.4294 | 0.882 | 0.598 | 1 | 10 |
| 1700037 | 0.4293 | 0.789 | 0.438 | 9.45E-05 | 8 |
| Tbx3 | 0.4285 | 0.338 | 0.014 | 1.09E-13 | 8 |
| Vamp1 | 0.4285 | 0.588 | 0.113 | 1 | 10 |

| | | | | | |
|---|---|---|---|---|---|
| Shtn1 | 0.4281 | 0.814 | 0.511 | 1.12E-08 | 5 |
| Thrb | 0.4281 | 0.588 | 0.132 | 1 | 10 |
| Thsd7b | 0.4278 | 0.753 | 0.301 | 1.19E-17 | 2 |
| Snca | 0.4278 | 0.767 | 0.311 | 1 | 9 |
| Cst3 | 0.4276 | 1 | 0.988 | 1 | 3 |
| Ccdc25 | 0.4275 | 0.7 | 0.216 | 0.0057 | 9 |
| Tcf25 | 0.4275 | 1 | 0.983 | 1 | 10 |
| Cdkn1c | 0.4273 | 0.342 | 0.09 | 9.00E-09 | 2 |
| Mef2c | 0.4271 | 0.32 | 0.21 | 0.0701 | 0 |
| Lmo4 | 0.4267 | 0.433 | 0.24 | 1 | 9 |
| 2010300( | 0.4266 | 0.588 | 0.098 | 1 | 10 |
| Syt11 | 0.4261 | 1 | 0.929 | 2.63E-15 | 0 |
| Stmn1 | 0.426 | 1 | 0.999 | 2.60E-08 | 7 |
| Thoc7 | 0.4255 | 0.952 | 0.714 | 4.78E-17 | 2 |
| Tmem132 | 0.4255 | 0.412 | 0.063 | 1 | 10 |
| Scn1b | 0.4254 | 0.87 | 0.622 | 5.51E-15 | 0 |
| Ppargc1a | 0.4254 | 0.515 | 0.135 | 5.19E-11 | 5 |
| Nol9 | 0.425 | 0.471 | 0.093 | 1 | 10 |
| Plcl1 | 0.4249 | 0.639 | 0.26 | 5.53E-09 | 5 |
| Nkx2-2 | 0.4249 | 0.412 | 0 | 5.96E-06 | 10 |
| Xk | 0.4247 | 0.529 | 0.067 | 0.9899 | 10 |
| Slc24a3 | 0.424 | 0.824 | 0.388 | 1 | 10 |
| Arhgdia | 0.4236 | 0.967 | 0.782 | 0.1269 | 9 |
| Lhfpl5 | 0.4235 | 0.69 | 0.355 | 2.51E-05 | 8 |
| Pgrmc1 | 0.4233 | 1 | 0.999 | 0.0424 | 3 |
| Lrpprc | 0.4232 | 0.765 | 0.26 | 1 | 10 |
| Ptk2b | 0.4226 | 0.412 | 0.03 | 1 | 10 |
| Nars | 0.4226 | 0.933 | 0.798 | 0.0601 | 9 |
| Pnoc | 0.4218 | 0.257 | 0.138 | 1 | 1 |
| Rorb | 0.4218 | 0.806 | 0.656 | 0.0001 | 4 |
| Ptpro | 0.4217 | 0.612 | 0.253 | 2.47E-12 | 1 |
| Rpl9 | 0.4214 | 1 | 0.986 | 1.16E-24 | 2 |
| Cystm1 | 0.4214 | 0.897 | 0.685 | 3.53E-07 | 7 |
| Tomm6 | 0.4213 | 0.795 | 0.355 | 6.56E-10 | 2 |
| Fat3 | 0.4211 | 0.765 | 0.196 | 1 | 10 |
| Atf6b | 0.421 | 0.588 | 0.161 | 1 | 10 |
| Tmem35 | 0.421 | 0.375 | 0.383 | 1 | 1 |
| Vamp2 | 0.4208 | 1 | 0.971 | 3.38E-10 | 2 |
| Npy2r | 0.4208 | 0.342 | 0.044 | 2.62E-17 | 1 |
| Cd24a | 0.4207 | 0.845 | 0.454 | 4.33E-05 | 6 |
| Sh3gl2 | 0.4207 | 0.795 | 0.44 | 1.58E-05 | 2 |
| Tekt1 | 0.4206 | 0.479 | 0.153 | 1.67E-05 | 8 |

| | | | | | |
|---|---|---|---|---|---|
| mt-Nd4 | 0.4203 | 1 | 1 | 1 | 10 |
| Cd47 | 0.4196 | 0.979 | 0.858 | 2.23E-15 | 2 |
| Gabra1 | 0.4196 | 0.555 | 0.324 | 3.44E-08 | 0 |
| Chl1 | 0.4196 | 0.718 | 0.459 | 0.0003 | 8 |
| Gng3 | 0.4195 | 1 | 0.986 | 1.05E-14 | 6 |
| Pcsk1n | 0.4194 | 1 | 1 | 5.60E-09 | 8 |
| Akr1a1 | 0.4192 | 0.972 | 0.936 | 8.38E-07 | 8 |
| Adgrl2 | 0.4189 | 0.6 | 0.158 | 0.14 | 9 |
| Cebpg | 0.4188 | 0.59 | 0.272 | 4.90E-07 | 7 |
| Spock3 | 0.4188 | 0.863 | 0.503 | 4.64E-15 | 2 |
| H3f3b | 0.4186 | 1 | 0.998 | 1.41E-08 | 7 |
| Rpl32 | 0.4186 | 0.986 | 0.968 | 0.0062 | 8 |
| Mir124-2 | 0.4186 | 0.699 | 0.255 | 4.06E-10 | 2 |
| Nat8l | 0.4185 | 0.5 | 0.271 | 8.88E-12 | 0 |
| Ndufa5 | 0.4185 | 1 | 0.97 | 0.3948 | 9 |
| Gt(ROSA)2 | 0.4184 | 0.487 | 0.414 | 1 | 7 |
| Enpp5 | 0.418 | 0.862 | 0.68 | 0.0073 | 3 |
| Phactr1 | 0.418 | 0.667 | 0.343 | 0.5475 | 9 |
| Gria2 | 0.4178 | 1 | 0.954 | 1 | 10 |
| Wipf2 | 0.4178 | 0.647 | 0.2 | 1 | 10 |
| Gabrq | 0.4175 | 0.507 | 0.166 | 3.09E-12 | 1 |
| Tox | 0.4173 | 0.583 | 0.3 | 4.78E-08 | 4 |
| Arid5b | 0.4172 | 0.633 | 0.373 | 1 | 9 |
| Zfp467 | 0.4172 | 0.753 | 0.374 | 3.78E-11 | 2 |
| Rit2 | 0.4168 | 0.979 | 0.874 | 1.28E-15 | 2 |
| Cpne6 | 0.4166 | 0.533 | 0.206 | 0.8822 | 9 |
| Sertad4 | 0.4162 | 0.582 | 0.157 | 8.18E-16 | 2 |
| Jak1 | 0.4159 | 0.882 | 0.491 | 1 | 10 |
| Aplp1 | 0.4156 | 1 | 0.981 | 1 | 3 |
| Iqsec3 | 0.4155 | 0.769 | 0.476 | 2.60E-05 | 7 |
| Tle4 | 0.4153 | 0.829 | 0.651 | 2.57E-07 | 3 |
| Cycs | 0.4151 | 0.967 | 0.926 | 1 | 9 |
| Rab11fip5 | 0.4151 | 0.588 | 0.139 | 1 | 10 |
| Arntl | 0.4151 | 0.567 | 0.2 | 6.93E-10 | 5 |
| Acap3 | 0.4148 | 0.588 | 0.186 | 1 | 10 |
| Jun | 0.4146 | 0.69 | 0.542 | 1 | 8 |
| Slc24a5 | 0.4146 | 0.897 | 0.515 | 9.09E-14 | 2 |
| Luc7l2 | 0.4145 | 0.967 | 0.857 | 1 | 3 |
| Rfwd2 | 0.4145 | 0.765 | 0.283 | 1 | 10 |
| Aldoc | 0.4143 | 0.704 | 0.274 | 0.002 | 8 |
| Necab1 | 0.4136 | 0.343 | 0.065 | 2.59E-12 | 4 |
| Ildr2 | 0.4131 | 0.592 | 0.302 | 0.0003 | 8 |

| | | | | | |
|---|---|---|---|---|---|
| Tecpr1 | 0.4129 | 0.941 | 0.357 | 1 | 10 |
| Pnp | 0.4128 | 0.59 | 0.299 | 3.81E-05 | 7 |
| Apoc3 | 0.4124 | 0.408 | 0.072 | 1.34E-17 | 1 |
| Sdc2 | 0.4124 | 0.706 | 0.472 | 1 | 10 |
| Cpne5 | 0.4123 | 0.583 | 0.253 | 8.41E-08 | 4 |
| Gng3 | 0.4118 | 1 | 0.986 | 2.24E-09 | 5 |
| 2210016I | 0.4117 | 0.634 | 0.276 | 4.01E-06 | 8 |
| Hmx3 | 0.4116 | 0.362 | 0.071 | 4.48E-17 | 1 |
| Fkbp1a | 0.4115 | 1 | 0.974 | 2.32E-11 | 2 |
| Gstm1 | 0.4115 | 0.493 | 0.136 | 3.27E-06 | 8 |
| Magi1 | 0.4114 | 0.941 | 0.521 | 1 | 10 |
| Eef1a1 | 0.411 | 1 | 1 | 8.00E-05 | 8 |
| Chchd10 | 0.4109 | 1 | 0.924 | 1 | 9 |
| Myef2 | 0.4105 | 0.904 | 0.597 | 2.77E-10 | 2 |
| Hnrnpf | 0.4105 | 0.705 | 0.266 | 4.57E-10 | 2 |
| Fus | 0.4105 | 0.967 | 0.872 | 1 | 3 |
| Rnf123 | 0.4105 | 0.588 | 0.11 | 1 | 10 |
| Usp29 | 0.4103 | 1 | 0.899 | 0.7385 | 3 |
| Pbx1 | 0.41 | 0.936 | 0.75 | 2.04E-07 | 7 |
| 1700001I | 0.41 | 0.781 | 0.344 | 1.71E-13 | 2 |
| Ccnd2 | 0.4098 | 0.352 | 0.052 | 2.02E-11 | 4 |
| Commd8 | 0.4098 | 0.801 | 0.433 | 1.88E-11 | 2 |
| Bbip1 | 0.4098 | 0.815 | 0.538 | 6.90E-07 | 2 |
| Ykt6 | 0.4098 | 0.706 | 0.24 | 1 | 10 |
| Fam3c | 0.4097 | 0.633 | 0.376 | 1 | 9 |
| Set | 0.4095 | 1 | 0.979 | 1.53E-19 | 2 |
| Hdac3 | 0.4095 | 0.765 | 0.498 | 1 | 10 |
| Tmem178 | 0.4094 | 0.382 | 0.169 | 2.89E-10 | 3 |
| Arglu1 | 0.4093 | 0.979 | 0.832 | 1.82E-15 | 2 |
| Thy1 | 0.4092 | 0.816 | 0.359 | 2.90E-13 | 1 |
| Sub1 | 0.4092 | 1 | 0.987 | 3.12E-20 | 2 |
| Tcf12 | 0.409 | 0.753 | 0.335 | 4.09E-16 | 2 |
| Nrxn3 | 0.4088 | 1 | 0.896 | 1.12E-16 | 1 |
| Itm2b | 0.4087 | 1 | 0.999 | 1 | 3 |
| Rab18 | 0.4085 | 0.918 | 0.695 | 1.87E-12 | 2 |
| Tada2b | 0.4081 | 0.529 | 0.113 | 1 | 10 |
| Tagln3 | 0.4077 | 0.986 | 0.944 | 6.04E-05 | 8 |
| Spint2 | 0.4077 | 0.954 | 0.912 | 5.66E-12 | 1 |
| Irs4 | 0.4075 | 0.69 | 0.243 | 0.0013 | 8 |
| Irf2bpl | 0.4073 | 0.654 | 0.414 | 1.34E-05 | 7 |
| Nlgn2 | 0.4072 | 0.882 | 0.529 | 1 | 10 |
| Grm1 | 0.4062 | 0.59 | 0.252 | 1.80E-06 | 7 |

| | | | | | |
|---|---|---|---|---|---|
| Gabra2 | 0.4062 | 0.764 | 0.584 | 8.85E-05 | 3 |
| St8sia4 | 0.4061 | 0.521 | 0.214 | 5.10E-05 | 8 |
| Acat1 | 0.4058 | 0.784 | 0.53 | 2.21E-05 | 5 |
| Gabre | 0.4055 | 0.487 | 0.102 | 1.19E-15 | 1 |
| Nxph1 | 0.4052 | 0.732 | 0.347 | 5.25E-05 | 5 |
| Unc5c | 0.405 | 0.788 | 0.396 | 1.31E-14 | 2 |
| Pdia3 | 0.4049 | 0.984 | 0.843 | 1 | 3 |
| Ldha | 0.4049 | 0.9 | 0.757 | 1 | 9 |
| Atp2b1 | 0.4045 | 0.915 | 0.811 | 2.13E-05 | 0 |
| Pcsk1 | 0.4041 | 0.308 | 0.101 | 0.0001 | 7 |
| Plcb4 | 0.4041 | 0.697 | 0.485 | 1.22E-08 | 1 |
| Stard4 | 0.4038 | 0.529 | 0.086 | 1 | 10 |
| Tgfa | 0.4038 | 0.63 | 0.186 | 4.43E-19 | 2 |
| Cyfip2 | 0.4037 | 0.85 | 0.616 | 1.33E-12 | 0 |
| Wnt5a | 0.4034 | 0.4 | 0.052 | 0.0021 | 9 |
| Grm5 | 0.4034 | 0.74 | 0.533 | 1.35E-06 | 0 |
| Tmx4 | 0.4032 | 1 | 0.982 | 1.29E-08 | 4 |
| Rps19 | 0.4031 | 1 | 0.988 | 4.63E-20 | 6 |
| Idnk | 0.4031 | 0.633 | 0.291 | 1 | 9 |
| Gramd1a | 0.4029 | 0.824 | 0.424 | 1 | 10 |
| Nudt10 | 0.4029 | 0.753 | 0.36 | 7.44E-06 | 2 |
| Sulf1 | 0.4028 | 0.481 | 0.146 | 1.17E-08 | 4 |
| Slc2a13 | 0.4028 | 0.41 | 0.212 | 0.0006 | 7 |
| Slc37a3 | 0.4027 | 0.529 | 0.156 | 1 | 10 |
| Rhoa | 0.4027 | 0.897 | 0.641 | 9.08E-12 | 2 |
| Atp5g2 | 0.4025 | 1 | 0.968 | 6.65E-20 | 2 |
| Arl6 | 0.4025 | 0.76 | 0.366 | 1.76E-05 | 2 |
| Vbp1 | 0.4025 | 0.842 | 0.528 | 8.07E-08 | 2 |
| Nts | 0.4024 | 0.072 | 0.022 | 1 | 1 |
| Gck | 0.4023 | 0.676 | 0.237 | 1.71E-07 | 8 |
| Flrt3 | 0.4022 | 0.795 | 0.53 | 0.0004 | 7 |
| Slc38a2 | 0.4019 | 0.472 | 0.382 | 0.0007 | 3 |
| Fam107a | 0.4019 | 0.529 | 0.05 | 1 | 10 |
| Acot7 | 0.4018 | 0.941 | 0.822 | 1 | 10 |
| Sgcz | 0.4018 | 0.488 | 0.296 | 5.33E-08 | 3 |
| Izumo4 | 0.4016 | 0.74 | 0.585 | 0.0002 | 3 |
| Mpped2 | 0.4015 | 0.692 | 0.362 | 7.53E-06 | 7 |
| Atp5g3 | 0.4012 | 1 | 0.977 | 0.0646 | 9 |
| Tmem255 | 0.4011 | 0.763 | 0.523 | 1.86E-09 | 1 |
| Arpc5 | 0.4011 | 0.887 | 0.748 | 0.0004 | 8 |
| Dip2c | 0.401 | 0.706 | 0.362 | 1 | 10 |
| 6330403 | 0.4007 | 1 | 0.999 | 3.31E-07 | 8 |

| | | | | | |
|---|---|---|---|---|---|
| Pcnp | 0.4007 | 0.918 | 0.676 | 4.75E-08 | 2 |
| Cetn2 | 0.4002 | 0.915 | 0.742 | 3.17E-05 | 8 |
| Ppia | 0.3999 | 1 | 1 | 5.47E-11 | 2 |
| Napb | 0.3997 | 0.667 | 0.467 | 1 | 9 |
| Isca1 | 0.3993 | 0.89 | 0.631 | 1.89E-07 | 2 |
| Zfp865 | 0.3993 | 0.588 | 0.123 | 1 | 10 |
| Plxdc2 | 0.3989 | 0.4 | 0.123 | 1 | 9 |
| Glul | 0.3985 | 0.588 | 0.236 | 5.06E-07 | 5 |
| Rpl35a | 0.3985 | 1 | 0.991 | 1 | 8 |
| Wsb1 | 0.3984 | 0.577 | 0.49 | 0.0114 | 3 |
| Rpl3 | 0.398 | 1 | 1 | 0.0358 | 8 |
| Gnal | 0.3979 | 0.533 | 0.24 | 1 | 9 |
| Gamt | 0.3977 | 0.647 | 0.125 | 1 | 10 |
| Nrp1 | 0.3977 | 0.504 | 0.34 | 9.09E-06 | 3 |
| Gm1007[illegible] | 0.3975 | 0.833 | 0.71 | 1 | 9 |
| Erbb4 | 0.3974 | 0.433 | 0.113 | 1 | 9 |
| C77370 | 0.3972 | 0.62 | 0.338 | 7.09E-08 | 4 |
| Fam19a1 | 0.397 | 0.825 | 0.387 | 1.98E-08 | 5 |
| Rit2 | 0.397 | 0.969 | 0.88 | 1.98E-08 | 5 |
| Clstn1 | 0.3968 | 0.95 | 0.827 | 2.47E-09 | 0 |
| Fam189a[illegible] | 0.3968 | 0.471 | 0.06 | 1 | 10 |
| Fam126a | 0.3964 | 0.588 | 0.085 | 1 | 10 |
| Timp3 | 0.3964 | 0.333 | 0.143 | 0.0013 | 7 |
| Rps2 | 0.3962 | 1 | 0.991 | 3.44E-24 | 2 |
| Atp6v1g2 | 0.3962 | 1 | 0.91 | 1 | 10 |
| Rell2 | 0.3961 | 0.767 | 0.445 | 0.3027 | 9 |
| Atp5k | 0.396 | 0.967 | 0.965 | 1 | 9 |
| Myo1b | 0.3959 | 0.471 | 0.056 | 1 | 10 |
| Rps29 | 0.3958 | 1 | 0.997 | 5.90E-09 | 7 |
| Ralyl | 0.3957 | 0.788 | 0.501 | 4.47E-05 | 2 |
| Slc25a5 | 0.3955 | 0.667 | 0.43 | 0.0163 | 7 |
| Prune2 | 0.3954 | 0.8 | 0.448 | 1 | 9 |
| Kcnq3 | 0.3953 | 0.5 | 0.163 | 1 | 9 |
| Sparcl1 | 0.3953 | 0.32 | 0.139 | 3.87E-05 | 0 |
| Arl4a | 0.3949 | 0.763 | 0.535 | 1.63E-06 | 5 |
| Sdcbp | 0.3947 | 0.808 | 0.419 | 0.0024 | 2 |
| Fam19a5 | 0.3947 | 1 | 0.644 | 1 | 10 |
| Pgd | 0.3946 | 0.647 | 0.212 | 1 | 10 |
| Glra1 | 0.3945 | 0.412 | 0.028 | 1 | 10 |
| Ntrk3 | 0.3945 | 0.741 | 0.501 | 1.82E-05 | 4 |
| Gap43 | 0.3944 | 1 | 0.99 | 2.14E-06 | 8 |
| Slc2a6 | 0.3943 | 0.529 | 0.158 | 1 | 10 |

| | | | | | |
|---|---|---|---|---|---|
| Grin1 | 0.3938 | 0.855 | 0.646 | 1.95E-10 | 0 |
| Pdia6 | 0.3936 | 0.976 | 0.816 | 1 | 3 |
| Ptma | 0.3934 | 1 | 0.985 | 1.11E-20 | 2 |
| Ntm | 0.3932 | 0.855 | 0.804 | 1.09E-05 | 1 |
| Ptp4a1 | 0.3928 | 0.564 | 0.382 | 0.0007 | 7 |
| Ncdn | 0.3927 | 0.8 | 0.579 | 1.58E-11 | 0 |
| Ap4s1 | 0.3927 | 0.767 | 0.524 | 1 | 9 |
| Cntn1 | 0.3924 | 0.845 | 0.588 | 3.73E-10 | 0 |
| B630019I | 0.3918 | 0.859 | 0.814 | 0.0029 | 7 |
| Cx3cl1 | 0.3917 | 0.78 | 0.542 | 1.43E-08 | 0 |
| Pou2f2 | 0.3917 | 0.862 | 0.569 | 1.27E-09 | 3 |
| Rps3a1 | 0.3917 | 1 | 0.978 | 0.315 | 8 |
| Cspg5 | 0.3916 | 0.963 | 0.895 | 4.76E-07 | 4 |
| Id2 | 0.3915 | 0.412 | 0.076 | 1 | 10 |
| Lrp1b | 0.3915 | 0.878 | 0.615 | 0.1944 | 3 |
| Htr2c | 0.3914 | 0.493 | 0.262 | 7.54E-06 | 1 |
| Gng3 | 0.3912 | 1 | 0.987 | 1.08E-08 | 7 |
| Saraf | 0.391 | 0.993 | 0.925 | 5.11E-14 | 1 |
| Adra2c | 0.391 | 0.471 | 0.031 | 0.213 | 10 |
| Abcc8 | 0.391 | 0.412 | 0.033 | 0.5242 | 10 |
| Zwint | 0.3906 | 1 | 0.996 | 1.09E-07 | 2 |
| Ctsb | 0.3906 | 0.985 | 0.91 | 1.51E-12 | 0 |
| Scd2 | 0.3904 | 0.959 | 0.786 | 1 | 3 |
| Vwa5b1 | 0.3902 | 0.423 | 0.106 | 4.90E-08 | 8 |
| Vkorc1l1 | 0.3901 | 0.824 | 0.301 | 1 | 10 |
| Msi2 | 0.3899 | 0.87 | 0.63 | 3.99E-10 | 3 |
| Churc1 | 0.3899 | 0.89 | 0.576 | 1.71E-10 | 2 |
| Sox5 | 0.3896 | 0.767 | 0.359 | 3.28E-13 | 2 |
| Vim | 0.3894 | 0.296 | 0.034 | 3.93E-06 | 8 |
| Insm2 | 0.3893 | 0.342 | 0.023 | 6.97E-25 | 1 |
| Magel2 | 0.3892 | 0.504 | 0.425 | 0.0099 | 3 |
| Rpl34 | 0.3883 | 1 | 0.971 | 0.0085 | 8 |
| Ndufs6 | 0.3882 | 0.933 | 0.883 | 1 | 9 |
| Pde1a | 0.3881 | 0.45 | 0.182 | 1.01E-09 | 0 |
| Wdr6 | 0.3881 | 1 | 0.933 | 1 | 10 |
| Tspan7 | 0.3881 | 0.967 | 0.829 | 5.67E-12 | 1 |
| Higd1a | 0.3879 | 0.986 | 0.752 | 2.93E-13 | 2 |
| Aifm3 | 0.3878 | 0.507 | 0.174 | 4.02E-05 | 8 |
| Rps27 | 0.3875 | 1 | 0.992 | 0.289 | 8 |
| Epha10 | 0.3875 | 0.471 | 0.065 | 1 | 10 |
| Lypd1 | 0.3874 | 0.535 | 0.192 | 0.0856 | 8 |
| Plcb1 | 0.3871 | 0.48 | 0.397 | 0.0001 | 3 |

| | | | | | |
|---|---|---|---|---|---|
| Fkbp2 | 0.387 | 0.985 | 0.917 | 1.74E-11 | 0 |
| Rnf112 | 0.3868 | 0.683 | 0.557 | 0.0234 | 3 |
| Rps18 | 0.3868 | 0.986 | 0.958 | 4.56E-17 | 2 |
| Gfm1 | 0.3867 | 0.529 | 0.109 | 1 | 10 |
| Atp5l | 0.3864 | 1 | 0.974 | 1 | 9 |
| Stx12 | 0.3862 | 0.87 | 0.608 | 5.27E-10 | 2 |
| Rps19bp1 | 0.3862 | 0.7 | 0.416 | 1 | 9 |
| Gas7 | 0.3862 | 0.733 | 0.319 | 1 | 9 |
| Six3 | 0.3861 | 0.856 | 0.666 | 0.0106 | 5 |
| Lgi1 | 0.3859 | 0.533 | 0.246 | 1 | 9 |
| Nkiras1 | 0.3855 | 0.833 | 0.621 | 1 | 9 |
| Arhgap6 | 0.3854 | 0.878 | 0.572 | 8.19E-11 | 3 |
| Oaz1 | 0.3851 | 1 | 0.998 | 3.60E-11 | 8 |
| Sumo2 | 0.3849 | 1 | 0.992 | 4.15E-24 | 2 |
| Cacna1d | 0.3849 | 0.836 | 0.48 | 6.25E-16 | 2 |
| Srsf5 | 0.3845 | 0.979 | 0.897 | 1.61E-11 | 2 |
| Inpp5k | 0.3845 | 0.647 | 0.169 | 1 | 10 |
| Id4 | 0.3844 | 0.885 | 0.625 | 9.37E-05 | 7 |
| Ptpn5 | 0.3843 | 0.667 | 0.34 | 1 | 9 |
| Nsg2 | 0.3843 | 1 | 0.943 | 1 | 3 |
| Fam196a | 0.3841 | 0.582 | 0.131 | 2.22E-16 | 2 |
| Tmem179 | 0.384 | 0.967 | 0.861 | 0.1106 | 9 |
| Rph3a | 0.3838 | 0.856 | 0.565 | 7.72E-06 | 5 |
| Slitrk4 | 0.3838 | 0.38 | 0.129 | 4.61E-15 | 0 |
| Atp6v0b | 0.3837 | 1 | 1 | 1 | 3 |
| Atp5d | 0.3837 | 1 | 0.961 | 1 | 9 |
| Reep2 | 0.3836 | 0.941 | 0.686 | 1 | 10 |
| Gstm6 | 0.3835 | 0.556 | 0.28 | 1.29E-07 | 4 |
| Cox5b | 0.3834 | 1 | 0.989 | 1 | 9 |
| Dnajb1 | 0.3834 | 0.435 | 0.318 | 1 | 4 |
| Prnp | 0.3834 | 0.993 | 0.997 | 6.00E-12 | 1 |
| Fxyd7 | 0.3833 | 0.852 | 0.645 | 6.84E-06 | 4 |
| Lman2 | 0.3832 | 0.882 | 0.334 | 1 | 10 |
| Fam19a2 | 0.3832 | 0.278 | 0.066 | 2.43E-07 | 4 |
| Dopey2 | 0.3831 | 0.588 | 0.19 | 1 | 10 |
| Cplx1 | 0.383 | 0.4 | 0.165 | 1 | 9 |
| Cry1 | 0.383 | 0.557 | 0.166 | 5.04E-09 | 5 |
| Negr1 | 0.3829 | 0.941 | 0.82 | 1.28E-10 | 1 |
| Det1 | 0.3829 | 0.454 | 0.076 | 4.89E-13 | 5 |
| Rgs17 | 0.3828 | 0.845 | 0.68 | 0.0076 | 8 |
| Fgf13 | 0.3827 | 0.567 | 0.266 | 1 | 9 |
| Cd164 | 0.3826 | 0.495 | 0.191 | 1.25E-06 | 5 |

| | | | | | |
|---|---|---|---|---|---|
| Rps19 | 0.3825 | 1 | 0.988 | 1.79E-08 | 7 |
| Ak5 | 0.3824 | 0.533 | 0.123 | 0.0314 | 9 |
| Pde1a | 0.3821 | 0.472 | 0.204 | 0.0002 | 4 |
| Alcam | 0.3819 | 0.966 | 0.774 | 1.04E-26 | 2 |
| Grin2b | 0.3817 | 0.885 | 0.628 | 8.75E-12 | 0 |
| Pld5 | 0.3817 | 0.343 | 0.076 | 2.65E-11 | 4 |
| Chchd7 | 0.3816 | 0.648 | 0.454 | 1 | 8 |
| Atpif1 | 0.3813 | 1 | 0.992 | 2.25E-05 | 8 |
| Map3k5 | 0.3809 | 0.536 | 0.192 | 5.81E-09 | 5 |
| Rps9 | 0.3805 | 1 | 0.972 | 3.28E-08 | 7 |
| Prkaca | 0.3805 | 1 | 0.885 | 1 | 10 |
| Rtn4 | 0.3805 | 0.991 | 0.944 | 3.09E-07 | 4 |
| Rbm39 | 0.3799 | 0.992 | 0.982 | 0.024 | 3 |
| Sccpdh | 0.3798 | 0.711 | 0.518 | 8.10E-07 | 1 |
| Atp1b1 | 0.3797 | 1 | 0.998 | 1.40E-11 | 0 |
| Chmp5 | 0.3796 | 0.763 | 0.507 | 0.0001 | 5 |
| Srsf2 | 0.3793 | 1 | 0.923 | 0.0001 | 3 |
| Rps21 | 0.3789 | 1 | 0.984 | 1.15E-14 | 6 |
| Wbp5 | 0.3788 | 0.958 | 0.944 | 0.0091 | 8 |
| Tox3 | 0.3788 | 0.671 | 0.335 | 1.34E-05 | 2 |
| Rgs7 | 0.3788 | 0.5 | 0.197 | 1 | 9 |
| Rps13 | 0.3787 | 1 | 0.977 | 1.60E-11 | 7 |
| Atp13a2 | 0.3786 | 0.882 | 0.469 | 1 | 10 |
| Rab40c | 0.3784 | 0.588 | 0.172 | 1 | 10 |
| Spint2 | 0.3782 | 1 | 0.908 | 0.0574 | 3 |
| 36324510 | 0.3782 | 0.415 | 0.272 | 5.87E-05 | 3 |
| Rnf114 | 0.3781 | 0.647 | 0.147 | 1 | 10 |
| Iqsec3 | 0.378 | 0.634 | 0.48 | 5.67E-05 | 3 |
| Chrm1 | 0.3779 | 0.353 | 0.037 | 1 | 10 |
| AI480526 | 0.3778 | 0.447 | 0.328 | 0.001 | 3 |
| Hnrnpk | 0.3776 | 0.993 | 0.952 | 2.45E-10 | 2 |
| Arhgdig | 0.3772 | 0.947 | 0.807 | 8.92E-12 | 1 |
| Fam183b | 0.377 | 0.648 | 0.284 | 0.0063 | 8 |
| Dusp22 | 0.377 | 0.471 | 0.117 | 1 | 10 |
| Esyt3 | 0.377 | 0.412 | 0.095 | 1 | 10 |
| Mt3 | 0.3767 | 1 | 0.961 | 0.0008 | 8 |
| Atp6v1g2 | 0.3764 | 0.955 | 0.902 | 2.29E-08 | 0 |
| Gabrg3 | 0.3762 | 0.657 | 0.372 | 2.32E-05 | 4 |
| Prex1 | 0.3761 | 0.515 | 0.141 | 3.24E-10 | 5 |
| Nt5dc3 | 0.376 | 0.463 | 0.318 | 4.80E-06 | 3 |
| Cbx3 | 0.3759 | 0.876 | 0.839 | 1 | 5 |
| Ptprt | 0.3757 | 0.588 | 0.225 | 0.0003 | 5 |

| | | | | | |
|---|---|---|---|---|---|
| Gmppa | 0.3754 | 0.588 | 0.181 | 1 | 10 |
| Mafg | 0.375 | 0.824 | 0.361 | 1 | 10 |
| Pdzrn4 | 0.3749 | 0.593 | 0.249 | 2.22E-07 | 4 |
| Adcy3 | 0.3748 | 0.487 | 0.329 | 0.1117 | 7 |
| Ocrl | 0.3747 | 0.794 | 0.496 | 7.32E-06 | 5 |
| Marcksl1 | 0.3744 | 0.986 | 0.942 | 0.0012 | 8 |
| Slc36a1 | 0.3741 | 0.529 | 0.085 | 1 | 10 |
| Fabp3 | 0.3738 | 0.833 | 0.656 | 0.0099 | 7 |
| Ywhab | 0.3736 | 1 | 0.984 | 1 | 10 |
| Ndufv2 | 0.3732 | 0.933 | 0.938 | 1 | 9 |
| Ldha | 0.3729 | 0.926 | 0.743 | 1.09E-07 | 4 |
| Rps16 | 0.3728 | 1 | 0.984 | 0.0125 | 8 |
| H3f3b | 0.3728 | 1 | 0.998 | 8.51E-25 | 2 |
| Kl | 0.3727 | 0.366 | 0.03 | 3.09E-11 | 8 |
| Gabra5 | 0.3723 | 0.667 | 0.242 | 1 | 9 |
| Romo1 | 0.3723 | 1 | 0.959 | 2.42E-06 | 7 |
| Cycs | 0.3721 | 0.959 | 0.924 | 2.14E-05 | 5 |
| Met | 0.372 | 0.25 | 0.011 | 2.05E-17 | 4 |
| Psap | 0.372 | 0.975 | 0.962 | 9.58E-09 | 0 |
| Zfp423 | 0.372 | 0.486 | 0.131 | 8.20E-12 | 2 |
| Tbl2 | 0.3716 | 0.471 | 0.077 | 1 | 10 |
| Arxes2 | 0.3711 | 0.984 | 0.846 | 0.0045 | 3 |
| Cdk14 | 0.3711 | 0.648 | 0.286 | 0.0012 | 8 |
| Sash1 | 0.3707 | 0.577 | 0.25 | 9.14E-06 | 5 |
| Lrp1b | 0.3706 | 0.8 | 0.64 | 1 | 9 |
| Lamp5 | 0.3705 | 0.316 | 0.063 | 2.19E-09 | 1 |
| Tecr | 0.3704 | 1 | 0.992 | 8.47E-12 | 0 |
| Uqcrh | 0.37 | 0.972 | 0.948 | 0.0277 | 8 |
| Efnb3 | 0.3699 | 0.797 | 0.692 | 1 | 3 |
| Rpl37a | 0.3699 | 1 | 0.998 | 1.36E-08 | 7 |
| Impact | 0.3695 | 1 | 0.984 | 2.57E-08 | 2 |
| Tgfa | 0.3694 | 0.439 | 0.22 | 1.83E-11 | 3 |
| Sirpa | 0.3691 | 0.745 | 0.509 | 1.17E-09 | 0 |
| Rap1gap | 0.3691 | 0.684 | 0.373 | 2.04E-09 | 1 |
| Zdhhc14 | 0.369 | 0.5 | 0.137 | 0.2025 | 9 |
| Mfng | 0.3685 | 0.309 | 0.042 | 3.06E-17 | 1 |
| Serpina3r | 0.3685 | 0.333 | 0.03 | 0.0039 | 9 |
| l7Rn6 | 0.3683 | 0.795 | 0.556 | 0.0005 | 7 |
| Dbn1 | 0.3681 | 0.927 | 0.77 | 0.2019 | 3 |
| Pdxk | 0.368 | 0.767 | 0.485 | 1 | 9 |
| Galnt16 | 0.3678 | 0.565 | 0.333 | 1.28E-07 | 0 |
| C1ql3 | 0.3677 | 0.676 | 0.132 | 0.0003 | 8 |

| | | | | | |
|---|---|---|---|---|---|
| Mgat4c | 0.3677 | 0.44 | 0.177 | 1.54E-09 | 0 |
| Ctbp2 | 0.3676 | 0.563 | 0.24 | 8.87E-05 | 8 |
| Rps19 | 0.3675 | 1 | 0.989 | 0.2302 | 8 |
| Ecel1 | 0.3675 | 0.36 | 0.201 | 0.0149 | 0 |
| Ptprm | 0.3674 | 0.454 | 0.122 | 2.94E-08 | 5 |
| Dpp9 | 0.3672 | 0.529 | 0.178 | 1 | 10 |
| Gnas | 0.3672 | 1 | 1 | 1.11E-13 | 1 |
| Morn2 | 0.3671 | 0.718 | 0.374 | 0.0049 | 8 |
| Rpl11 | 0.367 | 1 | 0.994 | 0.0035 | 8 |
| Prox1 | 0.3669 | 0.333 | 0.042 | 0.1701 | 9 |
| Ppm1g | 0.3669 | 0.882 | 0.563 | 1 | 10 |
| Zbtb20 | 0.3668 | 0.958 | 0.724 | 4.40E-05 | 8 |
| Nfix | 0.3666 | 0.215 | 0.077 | 5.78E-06 | 0 |
| Lsm14b | 0.3664 | 0.588 | 0.219 | 1 | 10 |
| Mpped2 | 0.366 | 0.712 | 0.336 | 2.25E-10 | 2 |
| Stk32c | 0.366 | 0.763 | 0.505 | 0.0002 | 5 |
| Clstn3 | 0.3659 | 0.935 | 0.801 | 4.64E-09 | 0 |
| Ucp2 | 0.3659 | 0.539 | 0.297 | 7.04E-05 | 1 |
| Ubb | 0.3658 | 1 | 1 | 1 | 8 |
| Tceal3 | 0.3656 | 0.9 | 0.946 | 1 | 9 |
| Tsc22d1 | 0.3652 | 1 | 0.859 | 1 | 9 |
| Srsf3 | 0.3651 | 0.979 | 0.962 | 1.15E-08 | 5 |
| Gm9866 | 0.365 | 0.412 | 0.113 | 1 | 10 |
| Atp5k | 0.3649 | 1 | 0.963 | 0.0001 | 8 |
| Ptms | 0.3648 | 1 | 0.969 | 2.26E-07 | 7 |
| Atp1a1 | 0.3643 | 0.759 | 0.579 | 0.0046 | 4 |
| Rab28 | 0.3642 | 0.763 | 0.547 | 0.0162 | 5 |
| Rnf182 | 0.3642 | 0.471 | 0.054 | 1 | 10 |
| Fhad1 | 0.3641 | 0.434 | 0.187 | 3.97E-07 | 1 |
| Dlgap1 | 0.364 | 0.59 | 0.387 | 0.0036 | 7 |
| 1110004 | 0.364 | 1 | 0.96 | 2.44E-12 | 6 |
| Ctsz | 0.3639 | 0.349 | 0.107 | 8.15E-09 | 1 |
| Trappc6b | 0.3639 | 0.973 | 0.861 | 1.54E-09 | 2 |
| Shtn1 | 0.3637 | 0.808 | 0.496 | 4.33E-11 | 2 |
| Atp9a | 0.3637 | 0.885 | 0.708 | 7.15E-08 | 0 |
| Mpc2 | 0.3637 | 0.933 | 0.927 | 1 | 9 |
| Elmo2 | 0.3635 | 0.647 | 0.19 | 1 | 10 |
| Mpped1 | 0.3632 | 0.4 | 0.097 | 1 | 9 |
| Ptprn | 0.3632 | 0.941 | 0.895 | 1 | 10 |
| Tpgs2 | 0.3632 | 0.701 | 0.415 | 3.31E-05 | 5 |
| Me3 | 0.3629 | 0.324 | 0.066 | 2.17E-11 | 4 |
| Atp2b1 | 0.3628 | 0.941 | 0.828 | 1 | 10 |

| | | | | | |
|---|---|---|---|---|---|
| Ubc | 0.3626 | 1 | 0.905 | 0.0018 | 7 |
| Whrn | 0.3625 | 0.529 | 0.079 | 1 | 10 |
| Tia1 | 0.3625 | 0.935 | 0.72 | 0.8447 | 3 |
| Mt3 | 0.3625 | 1 | 0.96 | 1.88E-12 | 6 |
| Coro2a | 0.3624 | 0.471 | 0.06 | 1 | 10 |
| Ephb1 | 0.3624 | 0.488 | 0.347 | 1.19E-06 | 3 |
| Eef1a1 | 0.3621 | 1 | 1 | 1.38E-07 | 7 |
| Tceal1 | 0.362 | 0.774 | 0.458 | 0.0066 | 2 |
| Arf4 | 0.3617 | 0.897 | 0.67 | 4.04E-08 | 2 |
| Cmpk1 | 0.3616 | 0.788 | 0.515 | 0.0441 | 2 |
| Gap43 | 0.3614 | 1 | 0.99 | 8.60E-11 | 6 |
| Gstm5 | 0.3614 | 0.9 | 0.763 | 1 | 9 |
| Hmgb1 | 0.3614 | 0.986 | 0.965 | 3.05E-11 | 2 |
| Clu | 0.3612 | 0.833 | 0.579 | 0.0002 | 4 |
| Maz | 0.3612 | 0.801 | 0.504 | 4.22E-07 | 2 |
| Cd83 | 0.3609 | 0.54 | 0.359 | 4.39E-06 | 0 |
| Serpine2 | 0.3608 | 0.382 | 0.261 | 0.3876 | 3 |
| Mrpl33 | 0.3607 | 0.859 | 0.707 | 0.1756 | 8 |
| Rragc | 0.3607 | 0.699 | 0.292 | 3.17E-05 | 2 |
| Rpl41 | 0.3606 | 1 | 1 | 5.48E-09 | 7 |
| Zfp703 | 0.3606 | 0.562 | 0.15 | 5.33E-15 | 2 |
| 1810037I | 0.3603 | 0.925 | 0.749 | 1.47E-09 | 2 |
| Nos1ap | 0.3602 | 0.69 | 0.35 | 0.0004 | 8 |
| Ndufa10 | 0.3601 | 0.9 | 0.824 | 1 | 9 |
| Brinp2 | 0.3601 | 0.586 | 0.282 | 7.58E-09 | 1 |
| Rora | 0.3601 | 0.801 | 0.397 | 1.08E-08 | 2 |
| Scn2a1 | 0.3601 | 1 | 0.881 | 1 | 10 |
| Mir124a-: | 0.36 | 0.545 | 0.396 | 0.003 | 3 |
| Pbxip1 | 0.3598 | 0.407 | 0.139 | 2.00E-06 | 4 |
| Tbc1d2b | 0.3597 | 0.235 | 0.025 | 1 | 10 |
| Clk1 | 0.3596 | 0.854 | 0.687 | 0.0033 | 3 |
| Adgrl3 | 0.3595 | 0.7 | 0.464 | 1.53E-07 | 0 |
| Zer1 | 0.3595 | 0.824 | 0.541 | 1 | 10 |
| Vstm2b | 0.3594 | 0.537 | 0.347 | 0.0015 | 4 |
| Kctd13 | 0.3592 | 0.808 | 0.41 | 0.0003 | 2 |
| Tmem255 | 0.3591 | 0.778 | 0.532 | 3.67E-05 | 4 |
| Snhg9 | 0.3591 | 0.753 | 0.419 | 3.81E-08 | 2 |
| Hmgn2 | 0.3591 | 0.949 | 0.901 | 0.0004 | 7 |
| Rhbdl3 | 0.359 | 0.353 | 0.064 | 1 | 10 |
| Cntn6 | 0.359 | 0.437 | 0.13 | 1.65E-06 | 8 |
| Tox2 | 0.3588 | 0.704 | 0.419 | 9.75E-06 | 4 |
| Matk | 0.3583 | 0.927 | 0.724 | 1 | 3 |

| | | | | | |
|---|---|---|---|---|---|
| Nog | 0.358 | 0.6 | 0.174 | 0.5773 | 9 |
| Fgf12 | 0.3578 | 0.6 | 0.335 | 1 | 9 |
| Rps27a | 0.3569 | 1 | 0.994 | 1.58E-06 | 7 |
| Ephb1 | 0.3568 | 0.74 | 0.306 | 3.04E-12 | 2 |
| Tle4 | 0.3566 | 0.863 | 0.641 | 1.42E-12 | 2 |
| Ctxn1 | 0.3566 | 0.972 | 0.9 | 0.0028 | 8 |
| Bhlhe41 | 0.3565 | 0.5 | 0.31 | 0.2545 | 7 |
| Resp18 | 0.3565 | 1 | 0.999 | 9.07E-05 | 1 |
| Celf5 | 0.3565 | 0.952 | 0.806 | 4.15E-16 | 2 |
| Psme1 | 0.3564 | 0.817 | 0.583 | 0.0265 | 8 |
| Mast4 | 0.3563 | 0.691 | 0.431 | 2.75E-07 | 1 |
| St13 | 0.3563 | 0.932 | 0.789 | 6.66E-07 | 2 |
| Cox7c | 0.3563 | 1 | 0.99 | 1.30E-07 | 5 |
| Syt9 | 0.356 | 0.529 | 0.093 | 1 | 10 |
| Atp1b3 | 0.3559 | 0.633 | 0.405 | 1 | 9 |
| Grina | 0.3556 | 0.93 | 0.853 | 2.95E-07 | 0 |
| Grin3a | 0.3553 | 0.472 | 0.323 | 0.2872 | 3 |
| Gria1 | 0.3553 | 0.82 | 0.517 | 5.35E-09 | 0 |
| 0610009 | 0.3552 | 0.701 | 0.48 | 0.0015 | 5 |
| Rprm | 0.3552 | 0.649 | 0.39 | 0.0004 | 6 |
| Dzank1 | 0.3551 | 0.74 | 0.546 | 6.32E-07 | 0 |
| Dach1 | 0.3551 | 0.282 | 0.03 | 6.85E-08 | 8 |
| Snca | 0.355 | 0.634 | 0.302 | 0.4284 | 8 |
| Ccdc184 | 0.3549 | 0.474 | 0.318 | 0.0059 | 7 |
| Klhl9 | 0.3548 | 0.719 | 0.361 | 0.0125 | 2 |
| Hbegf | 0.3546 | 0.473 | 0.176 | 2.15E-07 | 2 |
| BC00396 | 0.3546 | 0.667 | 0.426 | 1 | 9 |
| mt-Atp8 | 0.3545 | 1 | 0.96 | 1 | 10 |
| H3f3a | 0.3544 | 1 | 0.947 | 7.44E-06 | 5 |
| Pnck | 0.3544 | 1 | 0.925 | 1 | 10 |
| Rgs4 | 0.3544 | 0.454 | 0.186 | 0.0016 | 4 |
| Car8 | 0.3544 | 0.333 | 0.045 | 6.67E-16 | 4 |
| Zfp706 | 0.3543 | 0.959 | 0.888 | 2.65E-07 | 2 |
| Nup205 | 0.354 | 0.529 | 0.134 | 1 | 10 |
| Ppp1r2 | 0.3538 | 0.877 | 0.696 | 1.24E-06 | 2 |
| Trpc5 | 0.3536 | 0.491 | 0.264 | 0.0023 | 4 |
| Rps27a | 0.3532 | 1 | 0.994 | 2.53E-19 | 6 |
| Ckb | 0.353 | 0.99 | 0.991 | 6.24E-07 | 5 |
| Pnmal2 | 0.3528 | 1 | 0.993 | 1 | 10 |
| Snrpn | 0.3526 | 0.993 | 0.994 | 7.90E-12 | 2 |
| Frrs1l | 0.3523 | 0.785 | 0.569 | 4.26E-09 | 0 |
| Gm3038 | 0.3523 | 0.412 | 0.054 | 1 | 10 |

| | | | | | |
|---|---|---|---|---|---|
| Pole4 | 0.3522 | 0.596 | 0.238 | 9.51E-10 | 2 |
| Eno2 | 0.3519 | 0.835 | 0.691 | 1.20E-07 | 0 |
| Klhl2 | 0.3518 | 0.664 | 0.238 | 0.0006 | 2 |
| Synpo | 0.3518 | 0.471 | 0.033 | 1 | 10 |
| Sytl4 | 0.3517 | 0.257 | 0.034 | 2.64E-15 | 1 |
| Syp | 0.3515 | 0.97 | 0.928 | 1.14E-05 | 0 |
| Cyfip1 | 0.3513 | 0.529 | 0.162 | 1 | 10 |
| Elavl2 | 0.3513 | 0.7 | 0.307 | 1 | 9 |
| Amd1 | 0.3512 | 0.815 | 0.438 | 1.49E-07 | 2 |
| Prdx3 | 0.3511 | 0.733 | 0.429 | 0.0001 | 2 |
| Kcnk2 | 0.351 | 0.62 | 0.333 | 0.0019 | 4 |
| Pclo | 0.351 | 0.895 | 0.74 | 2.96E-06 | 0 |
| Cystm1 | 0.351 | 0.904 | 0.669 | 1.83E-08 | 2 |
| Ajap1 | 0.3505 | 0.496 | 0.431 | 0.0006 | 3 |
| Serp2 | 0.3505 | 0.963 | 0.879 | 2.37E-07 | 4 |
| Atp8a1 | 0.3505 | 0.907 | 0.776 | 6.96E-05 | 4 |
| Tcp1 | 0.3505 | 0.944 | 0.724 | 0.0004 | 8 |
| Grin2a | 0.3505 | 0.567 | 0.133 | 0.2631 | 9 |
| AI593442 | 0.3503 | 0.692 | 0.268 | 3.22E-07 | 2 |
| Map1b | 0.3501 | 1 | 0.976 | 9.80E-08 | 4 |
| Pi16 | 0.3497 | 0.493 | 0.158 | 4.94E-05 | 8 |
| Fbxo44 | 0.3493 | 0.845 | 0.548 | 5.07E-07 | 5 |
| Lhfpl3 | 0.3492 | 0.461 | 0.165 | 1.28E-10 | 1 |
| Dnajc19 | 0.3491 | 0.9 | 0.646 | 1 | 9 |
| Ndufa4 | 0.349 | 1 | 0.995 | 0.0212 | 9 |
| Actb | 0.349 | 1 | 0.998 | 1.10E-08 | 2 |
| Eif1 | 0.3489 | 1 | 0.998 | 3.62E-24 | 2 |
| Gm15261 | 0.3487 | 0.374 | 0.207 | 4.51E-06 | 3 |
| Stmn4 | 0.3486 | 0.907 | 0.776 | 0.0008 | 4 |
| Mrpl17 | 0.3486 | 0.808 | 0.65 | 0.0118 | 7 |
| C1ql2 | 0.3486 | 0.204 | 0.033 | 1.01E-05 | 1 |
| Kcnab1 | 0.3486 | 0.4 | 0.075 | 0.9622 | 9 |
| Glra2 | 0.3483 | 0.275 | 0.065 | 5.03E-14 | 0 |
| Rad23a | 0.348 | 0.767 | 0.395 | 5.45E-06 | 2 |
| Map1lc3a | 0.348 | 1 | 0.967 | 3.19E-12 | 2 |
| Mpp6 | 0.3479 | 0.546 | 0.245 | 4.18E-05 | 5 |
| Cpne2 | 0.3477 | 0.733 | 0.445 | 1 | 9 |
| Gm10076 | 0.3476 | 0.821 | 0.705 | 1 | 7 |
| Tmed9 | 0.3475 | 0.911 | 0.826 | 1 | 3 |
| Tshz2 | 0.3475 | 0.882 | 0.514 | 1 | 10 |
| Rnf181 | 0.3474 | 0.678 | 0.317 | 0.0001 | 2 |
| Fgfr1 | 0.3474 | 0.368 | 0.249 | 0.0041 | 1 |

| | | | | | |
|---|---|---|---|---|---|
| Tbca | 0.3473 | 0.944 | 0.881 | 0.005 | 8 |
| Phlda1 | 0.3471 | 0.467 | 0.198 | 1 | 9 |
| Thsd7b | 0.3471 | 0.649 | 0.333 | 0.0008 | 5 |
| Rexo2 | 0.347 | 0.603 | 0.467 | 0.4194 | 7 |
| Rps24 | 0.3469 | 1 | 0.997 | 1.75E-19 | 6 |
| Rps3 | 0.3469 | 0.986 | 0.981 | 0.13 | 8 |
| Sphkap | 0.3466 | 0.5 | 0.187 | 1 | 9 |
| Pomt1 | 0.3463 | 0.529 | 0.098 | 1 | 10 |
| Cux1 | 0.3461 | 0.733 | 0.391 | 5.80E-08 | 2 |
| Lin7a | 0.3461 | 0.918 | 0.663 | 0.0002 | 6 |
| Abhd17b | 0.3461 | 0.562 | 0.236 | 0.0002 | 2 |
| Cox8a | 0.3458 | 1 | 0.998 | 4.48E-05 | 8 |
| Negr1 | 0.3457 | 0.967 | 0.833 | 0.5962 | 9 |
| Ywhae | 0.3455 | 1 | 1 | 5.07E-13 | 2 |
| 6330403 | 0.3454 | 1 | 0.999 | 1 | 9 |
| Cnih2 | 0.3452 | 0.967 | 0.976 | 0.1633 | 9 |
| Frat2 | 0.3452 | 0.282 | 0.187 | 1 | 8 |
| Ap1s2 | 0.3451 | 0.825 | 0.59 | 0.0008 | 5 |
| Rpl37a | 0.3451 | 1 | 0.998 | 9.72E-15 | 6 |
| Nudt4 | 0.345 | 0.6 | 0.214 | 1 | 9 |
| Ift22 | 0.345 | 0.767 | 0.639 | 1 | 9 |
| Flrt3 | 0.3449 | 0.814 | 0.523 | 3.77E-05 | 5 |
| Faim2 | 0.3447 | 1 | 0.968 | 1.44E-09 | 0 |
| Dynll1 | 0.3446 | 1 | 0.977 | 2.66E-13 | 6 |
| Med19 | 0.3443 | 0.667 | 0.339 | 1 | 9 |
| Mest | 0.3443 | 0.706 | 0.281 | 1 | 10 |
| Anp32a | 0.344 | 0.907 | 0.818 | 4.15E-05 | 5 |
| Phpt1 | 0.344 | 0.773 | 0.575 | 0.0389 | 5 |
| Aplp1 | 0.3439 | 1 | 0.983 | 0.4458 | 10 |
| Maged1 | 0.3434 | 1 | 0.987 | 0.0557 | 10 |
| Tead1 | 0.3433 | 0.602 | 0.283 | 8.69E-07 | 4 |
| Ndufa2 | 0.3432 | 1 | 0.933 | 1.55E-16 | 6 |
| Ogfod1 | 0.3431 | 0.55 | 0.376 | 2.55E-07 | 0 |
| Nmu | 0.343 | 0.176 | 0.002 | 1 | 10 |
| Plcl1 | 0.3429 | 0.651 | 0.239 | 5.17E-09 | 2 |
| Suclg1 | 0.3426 | 0.833 | 0.769 | 1 | 9 |
| Celf6 | 0.3426 | 0.904 | 0.698 | 1.81E-07 | 2 |
| Ndufs7 | 0.3424 | 1 | 0.952 | 0.0022 | 9 |
| 1700023 | 0.3424 | 0.649 | 0.367 | 0.0014 | 5 |
| Plxna2 | 0.342 | 0.454 | 0.169 | 4.93E-10 | 1 |
| Rpl37 | 0.3419 | 1 | 0.996 | 0.9344 | 8 |
| Hap1 | 0.3419 | 1 | 0.996 | 1 | 3 |

| | | | | | |
|---|---|---|---|---|---|
| Rnf208 | 0.3418 | 0.967 | 0.693 | 0.6522 | 9 |
| Pcdh10 | 0.3416 | 0.918 | 0.709 | 6.38E-05 | 5 |
| Fgf8 | 0.3416 | 0.295 | 0.045 | 0.0057 | 7 |
| Rps15 | 0.3414 | 0.928 | 0.821 | 1.04E-05 | 6 |
| Fam222a | 0.3413 | 0.333 | 0.079 | 0.808 | 9 |
| Rps26 | 0.3412 | 0.979 | 0.936 | 1.67E-09 | 6 |
| Tram1l1 | 0.3411 | 0.814 | 0.685 | 1.20E-05 | 5 |
| Galnt16 | 0.3409 | 0.567 | 0.369 | 1 | 9 |
| Ly6e | 0.3409 | 0.671 | 0.432 | 2.34E-06 | 1 |
| Tiprl | 0.3408 | 0.767 | 0.548 | 0.0821 | 2 |
| Hmcn2 | 0.3403 | 0.353 | 0.004 | 0.3418 | 10 |
| Rps4x | 0.34 | 1 | 0.98 | 3.33E-16 | 6 |
| Pnn | 0.3399 | 0.894 | 0.775 | 1 | 3 |
| Scgn | 0.3399 | 0.233 | 0.028 | 1 | 9 |
| Arhgdig | 0.3398 | 0.926 | 0.815 | 0.0002 | 4 |
| Ssbp2 | 0.3397 | 0.878 | 0.73 | 0.1265 | 3 |
| Hypk | 0.3395 | 0.966 | 0.887 | 1.05E-08 | 2 |
| Ache | 0.3395 | 0.73 | 0.513 | 1.15E-06 | 0 |
| Nrn1 | 0.3394 | 0.222 | 0.018 | 1.27E-06 | 4 |
| Dtd1 | 0.3391 | 0.945 | 0.842 | 1.13E-10 | 2 |
| Tmem132 | 0.339 | 0.568 | 0.237 | 3.58E-07 | 2 |
| Atp5j | 0.339 | 1 | 0.981 | 1 | 9 |
| Nefl | 0.3388 | 0.445 | 0.198 | 6.44E-10 | 0 |
| Pxn | 0.3387 | 0.353 | 0.042 | 1 | 10 |
| Wipf3 | 0.3387 | 0.294 | 0.044 | 1 | 10 |
| Map2k4 | 0.3385 | 0.705 | 0.491 | 5.20E-08 | 0 |
| Ankrd9 | 0.3383 | 0.533 | 0.159 | 0.7098 | 9 |
| Mir124-2 | 0.3383 | 0.447 | 0.296 | 7.06E-09 | 3 |
| Qk | 0.3383 | 0.433 | 0.163 | 1 | 9 |
| Atp2b1 | 0.3381 | 1 | 0.808 | 1 | 3 |
| Nap1l5 | 0.3381 | 1 | 0.999 | 4.77E-08 | 1 |
| Kctd8 | 0.338 | 0.473 | 0.19 | 1.86E-06 | 2 |
| Csnk2b | 0.338 | 0.767 | 0.621 | 1 | 9 |
| Pgk1 | 0.3379 | 0.774 | 0.435 | 0.0004 | 2 |
| Mageh1 | 0.3377 | 0.806 | 0.591 | 0.0002 | 4 |
| Dgkg | 0.3377 | 0.6 | 0.264 | 1 | 9 |
| Zscan26 | 0.3377 | 0.664 | 0.345 | 0.1154 | 2 |
| Rpl13 | 0.3376 | 1 | 0.996 | 0.0101 | 8 |
| Sumo3 | 0.3375 | 0.958 | 0.835 | 0.001 | 8 |
| Tcaf1 | 0.3373 | 0.976 | 0.873 | 0.4409 | 3 |
| Tcf12 | 0.3373 | 0.496 | 0.377 | 7.38E-07 | 3 |
| Negr1 | 0.3373 | 0.92 | 0.818 | 4.73E-06 | 0 |

| | | | | | |
|---|---|---|---|---|---|
| Fhad1 | 0.3371 | 0.567 | 0.211 | 1 | 9 |
| Tcf4 | 0.3371 | 0.481 | 0.286 | 0.0207 | 4 |
| Kdm6b | 0.337 | 0.651 | 0.286 | 6.48E-07 | 2 |
| Sephs2 | 0.3369 | 0.61 | 0.247 | 0.0003 | 2 |
| Adra2a | 0.3363 | 0.352 | 0.052 | 5.28E-15 | 4 |
| Tmem179 | 0.3362 | 0.96 | 0.843 | 3.86E-09 | 0 |
| Cox6b1 | 0.3362 | 1 | 0.973 | 0.0254 | 8 |
| Glul | 0.3361 | 0.575 | 0.22 | 0.0007 | 2 |
| Ttc39c | 0.3361 | 0.529 | 0.152 | 1 | 10 |
| Selk | 0.336 | 1 | 0.99 | 1 | 3 |
| C2cd4b | 0.3358 | 0.103 | 0.026 | 1 | 7 |
| Sox3 | 0.3357 | 0.487 | 0.156 | 4.24E-06 | 7 |
| Gda | 0.3353 | 0.831 | 0.617 | 0.0664 | 8 |
| Spon1 | 0.3351 | 0.74 | 0.36 | 4.08E-09 | 2 |
| Ddc | 0.3351 | 0.528 | 0.204 | 3.69E-05 | 4 |
| Smad3 | 0.3349 | 0.353 | 0.01 | 1 | 10 |
| Fam171b | 0.3349 | 0.921 | 0.796 | 1.04E-07 | 1 |
| Pgm2l1 | 0.3348 | 0.61 | 0.446 | 1.99E-05 | 0 |
| Gpr150 | 0.3347 | 0.412 | 0.077 | 1 | 10 |
| Six6 | 0.3345 | 0.732 | 0.389 | 3.09E-06 | 6 |
| Myeov2 | 0.3345 | 0.99 | 0.888 | 7.32E-12 | 6 |
| Bre | 0.3344 | 0.633 | 0.259 | 1 | 9 |
| Hs3st5 | 0.3343 | 0.342 | 0.037 | 3.41E-21 | 1 |
| Sub1 | 0.3341 | 1 | 0.988 | 3.40E-06 | 7 |
| Brinp3 | 0.334 | 0.545 | 0.384 | 0.0006 | 0 |
| Fam204a | 0.3338 | 0.671 | 0.33 | 0.0009 | 2 |
| Gm11808 | 0.3337 | 0.616 | 0.272 | 0.0003 | 2 |
| Hs2st1 | 0.3337 | 0.633 | 0.365 | 1 | 9 |
| Dlx2 | 0.3337 | 0.493 | 0.191 | 0.0009 | 8 |
| Rps27a | 0.3336 | 1 | 0.994 | 1 | 8 |
| Ywhaq | 0.3335 | 1 | 0.985 | 5.87E-07 | 7 |
| Gnai1 | 0.3334 | 0.833 | 0.733 | 0.0026 | 4 |
| Dgki | 0.3333 | 0.589 | 0.192 | 2.60E-13 | 2 |
| Ftl1 | 0.3332 | 1 | 0.95 | 0.001 | 8 |
| Vat1 | 0.3329 | 0.981 | 0.887 | 1.22E-06 | 4 |
| Tuba1a | 0.3329 | 0.99 | 0.979 | 0.0841 | 6 |
| Gabra3 | 0.3327 | 0.592 | 0.286 | 5.60E-09 | 1 |
| Tnrc6c | 0.3327 | 0.618 | 0.575 | 1 | 3 |
| Drd1 | 0.3327 | 0.321 | 0.049 | 7.44E-08 | 7 |
| Ndn | 0.3326 | 0.958 | 0.864 | 0.0005 | 8 |
| Pcnxl3 | 0.3326 | 0.647 | 0.177 | 1 | 10 |
| Nr2f2 | 0.3326 | 0.616 | 0.392 | 1 | 2 |

| | | | | | |
|---|---|---|---|---|---|
| Rpl41 | 0.3325 | 1 | 1 | 0.013 | 8 |
| Mrap2 | 0.3322 | 0.602 | 0.35 | 0.0008 | 4 |
| Uqcc2 | 0.3322 | 1 | 0.916 | 1.61E-12 | 6 |
| Igfbpl1 | 0.3321 | 0.333 | 0.145 | 0.0003 | 3 |
| Lynx1 | 0.332 | 0.355 | 0.126 | 1.54E-09 | 0 |
| Fam213b | 0.3318 | 0.767 | 0.522 | 1 | 9 |
| Gpr88 | 0.3317 | 0.185 | 0.06 | 0.0076 | 4 |
| Rpl37a | 0.3317 | 1 | 0.998 | 0.0131 | 8 |
| Ddc | 0.3315 | 0.454 | 0.201 | 0.0001 | 1 |
| Lingo1 | 0.3315 | 0.731 | 0.518 | 0.0301 | 7 |
| Prr13 | 0.3314 | 0.873 | 0.656 | 0.0079 | 8 |
| Araf | 0.3313 | 0.986 | 0.912 | 1.78E-08 | 2 |
| Whrn | 0.3312 | 0.333 | 0.079 | 1 | 9 |
| Kif5c | 0.3312 | 0.898 | 0.797 | 0.0016 | 4 |
| Cpe | 0.3311 | 1 | 0.964 | 1 | 3 |
| AW55198 | 0.3309 | 0.882 | 0.806 | 7.52E-07 | 1 |
| Gabrg3 | 0.3309 | 0.632 | 0.363 | 1.96E-06 | 1 |
| Mrfap1 | 0.3308 | 1 | 0.993 | 0.001 | 8 |
| Lmo3 | 0.3306 | 0.479 | 0.131 | 0.0149 | 8 |
| 1110008I | 0.3306 | 0.767 | 0.48 | 1 | 9 |
| Nedd8 | 0.3303 | 1 | 0.968 | 0.0304 | 8 |
| Ndn | 0.3303 | 1 | 0.857 | 1 | 6 |
| Cntn5 | 0.3302 | 0.34 | 0.128 | 1.05E-06 | 0 |
| Pcdh11x | 0.33 | 0.368 | 0.143 | 1.31E-05 | 1 |
| Lrrc4c | 0.3299 | 0.628 | 0.373 | 0.0012 | 7 |
| Rpsa | 0.3299 | 1 | 0.998 | 9.27E-06 | 8 |
| Ppp1r17 | 0.3296 | 0.491 | 0.289 | 0.0038 | 4 |
| Pcbp3 | 0.3296 | 0.533 | 0.275 | 1 | 9 |
| Tmem243 | 0.3295 | 0.534 | 0.244 | 1.67E-06 | 2 |
| Prkaa2 | 0.3295 | 0.397 | 0.168 | 3.50E-05 | 7 |
| Atp5o | 0.3295 | 0.915 | 0.87 | 1 | 8 |
| Il1rap | 0.3294 | 0.329 | 0.068 | 6.81E-14 | 1 |
| Tuba1a | 0.3294 | 1 | 0.979 | 0.738 | 7 |
| Grik3 | 0.3294 | 0.401 | 0.13 | 1.36E-10 | 1 |
| Gaa | 0.3293 | 0.987 | 0.972 | 4.18E-08 | 1 |
| Adgrb2 | 0.3293 | 0.585 | 0.429 | 0.063 | 3 |
| Plcxd3 | 0.329 | 0.726 | 0.365 | 4.12E-06 | 2 |
| Unc5b | 0.329 | 0.394 | 0.067 | 9.17E-09 | 8 |
| Ampd2 | 0.329 | 0.618 | 0.517 | 0.1549 | 3 |
| Vsnl1 | 0.329 | 0.845 | 0.833 | 1 | 5 |
| Nrxn3 | 0.3289 | 1 | 0.904 | 4.10E-07 | 8 |
| Eif4g2 | 0.3288 | 0.979 | 0.883 | 2.73E-05 | 5 |

| | | | | | |
|---|---|---|---|---|---|
| Galnt13 | 0.3288 | 0.536 | 0.254 | 0.0076 | 5 |
| Tmem145 | 0.3287 | 0.545 | 0.47 | 1 | 3 |
| Unc5c | 0.3287 | 0.701 | 0.423 | 0.0041 | 5 |
| Fgf1 | 0.3286 | 0.235 | 0.016 | 1 | 10 |
| Strap | 0.3286 | 0.863 | 0.643 | 0.0017 | 2 |
| Nsg2 | 0.3285 | 0.975 | 0.943 | 0.0145 | 0 |
| Atp1a3 | 0.3283 | 0.97 | 0.94 | 0.0017 | 0 |
| 2900055 | 0.3283 | 0.435 | 0.25 | 0.0374 | 4 |
| Cnnm1 | 0.3282 | 0.5 | 0.253 | 1 | 9 |
| Prkcb | 0.3282 | 0.515 | 0.283 | 5.38E-10 | 0 |
| Flrt3 | 0.3282 | 0.815 | 0.509 | 1.59E-10 | 2 |
| Basp1 | 0.328 | 1 | 0.999 | 1 | 9 |
| Grik1 | 0.328 | 0.333 | 0.097 | 1 | 9 |
| Phyh | 0.3279 | 0.704 | 0.477 | 1 | 8 |
| Vgf | 0.3275 | 0.649 | 0.246 | 0.0012 | 5 |
| Auts2 | 0.3274 | 0.962 | 0.721 | 3.08E-05 | 7 |
| Dlx6 | 0.3274 | 0.568 | 0.22 | 3.72E-06 | 2 |
| Pim3 | 0.3273 | 0.633 | 0.305 | 1 | 9 |
| Snhg9 | 0.3273 | 0.537 | 0.454 | 2.95E-05 | 3 |
| Rnf41 | 0.3273 | 0.644 | 0.303 | 0.0083 | 2 |
| Atp1a3 | 0.3273 | 1 | 0.945 | 1 | 10 |
| 2700094 | 0.3272 | 0.907 | 0.656 | 1.05E-09 | 6 |
| Rpl35 | 0.3272 | 1 | 0.984 | 1.21E-19 | 2 |
| Dmd | 0.3272 | 0.537 | 0.285 | 0.0003 | 4 |
| mt-Nd3 | 0.327 | 1 | 0.926 | 1 | 10 |
| Ddx5 | 0.3269 | 0.992 | 0.984 | 1 | 3 |
| Ube2d1 | 0.3269 | 0.692 | 0.299 | 1.13E-05 | 2 |
| Ret | 0.3268 | 0.352 | 0.105 | 1.91E-06 | 4 |
| Mapk3 | 0.3267 | 0.712 | 0.381 | 0.0015 | 2 |
| Eid1 | 0.3267 | 0.986 | 0.961 | 0.1348 | 8 |
| Rpl30 | 0.3267 | 1 | 0.988 | 5.95E-16 | 6 |
| Gpm6a | 0.3266 | 0.98 | 0.963 | 1.27E-06 | 0 |
| Cit | 0.3266 | 0.755 | 0.533 | 5.25E-07 | 0 |
| Fxyd2 | 0.3264 | 0.37 | 0.162 | 1.75E-05 | 4 |
| Camkv | 0.3262 | 0.64 | 0.449 | 2.79E-06 | 0 |
| Uqcr10 | 0.326 | 1 | 0.958 | 9.29E-06 | 5 |
| Ube2e1 | 0.3258 | 0.932 | 0.738 | 6.45E-08 | 2 |
| Rap1gds1 | 0.3257 | 0.726 | 0.394 | 0.0031 | 2 |
| Rps20 | 0.3255 | 1 | 0.977 | 3.53E-05 | 7 |
| Timm10 | 0.3254 | 0.692 | 0.364 | 0.002 | 2 |
| Rps20 | 0.3252 | 1 | 0.977 | 1 | 8 |
| Acat1 | 0.3251 | 0.774 | 0.519 | 0.0716 | 2 |

| | | | | | |
|---|---|---|---|---|---|
| Sap30l | 0.3251 | 0.535 | 0.295 | 0.0463 | 8 |
| Smad2 | 0.3248 | 0.637 | 0.234 | 8.40E-08 | 2 |
| 2700094I | 0.3247 | 0.897 | 0.644 | 1.13E-08 | 2 |
| Arl4a | 0.3245 | 0.756 | 0.54 | 0.0193 | 7 |
| Per2 | 0.3243 | 0.372 | 0.096 | 3.95E-08 | 7 |
| Nrp2 | 0.3239 | 0.732 | 0.464 | 0.002 | 5 |
| Cntnap4 | 0.3236 | 0.401 | 0.132 | 5.11E-09 | 1 |
| Ssu72 | 0.3235 | 0.91 | 0.757 | 0.0318 | 7 |
| Raly | 0.3233 | 0.676 | 0.447 | 0.5979 | 8 |
| Snx2 | 0.3233 | 0.596 | 0.303 | 0.0038 | 2 |
| 2700029I | 0.3231 | 0.795 | 0.497 | 8.78E-05 | 2 |
| Npy1r | 0.323 | 0.536 | 0.218 | 0.0001 | 5 |
| Cpne7 | 0.323 | 0.441 | 0.156 | 3.43E-07 | 1 |
| Ptpn5 | 0.3229 | 0.525 | 0.31 | 3.95E-06 | 0 |
| Gnas | 0.3228 | 1 | 1 | 1.23E-10 | 0 |
| Rpl39 | 0.3228 | 1 | 0.974 | 1 | 8 |
| Nudt19 | 0.3227 | 0.722 | 0.427 | 0.0008 | 5 |
| Pdzrn4 | 0.3227 | 0.592 | 0.261 | 0.0038 | 8 |
| Ctxn2 | 0.3226 | 0.767 | 0.53 | 0.0004 | 2 |
| 2610301I | 0.3225 | 0.733 | 0.428 | 0.0024 | 2 |
| Pcdh18 | 0.3224 | 0.267 | 0.046 | 0.2478 | 9 |
| Hlf | 0.3224 | 0.788 | 0.484 | 1.64E-05 | 2 |
| Fndc5 | 0.3221 | 0.4 | 0.222 | 3.53E-07 | 0 |
| 2010005I | 0.322 | 0.397 | 0.15 | 0.006 | 7 |
| Ppp1r15a | 0.322 | 0.436 | 0.128 | 7.90E-07 | 7 |
| Fam159b | 0.3218 | 0.233 | 0.051 | 1 | 9 |
| Th | 0.3218 | 0.282 | 0.097 | 1 | 8 |
| Brinp3 | 0.3216 | 0.7 | 0.405 | 1 | 9 |
| Sec11c | 0.3216 | 0.967 | 0.858 | 0.3127 | 3 |
| Matk | 0.3216 | 0.89 | 0.715 | 8.39E-07 | 0 |
| Fkbp3 | 0.3215 | 1 | 0.967 | 0.4645 | 8 |
| Tsc22d3 | 0.3215 | 0.733 | 0.385 | 0.0001 | 2 |
| Tmem15C | 0.3213 | 0.654 | 0.372 | 0.0006 | 7 |
| Cd200 | 0.3213 | 1 | 0.851 | 1 | 3 |
| Csmd1 | 0.3212 | 0.526 | 0.263 | 0.0033 | 5 |
| Snca | 0.3212 | 0.44 | 0.298 | 5.19E-06 | 0 |
| Naca | 0.3211 | 1 | 0.984 | 4.35E-15 | 6 |
| Prr7 | 0.321 | 0.933 | 0.709 | 1 | 9 |
| Ankrd34b | 0.321 | 0.352 | 0.123 | 0.012 | 8 |
| Cd63 | 0.3207 | 0.691 | 0.428 | 7.18E-06 | 1 |
| Ar | 0.3206 | 0.7 | 0.403 | 1 | 9 |
| Smim14 | 0.3206 | 0.925 | 0.732 | 1.32E-08 | 2 |

| | | | | | |
|---|---|---|---|---|---|
| Fgf9 | 0.3204 | 0.333 | 0.086 | 1 | 9 |
| Gldn | 0.3204 | 0.367 | 0.031 | 0.0035 | 9 |
| Bloc1s1 | 0.3204 | 0.616 | 0.29 | 0.002 | 2 |
| Rps18 | 0.3201 | 0.987 | 0.96 | 2.55E-06 | 7 |
| Jph4 | 0.3201 | 0.862 | 0.737 | 1 | 3 |
| Esd | 0.3199 | 0.867 | 0.765 | 1 | 9 |
| Ube2e2 | 0.3199 | 0.933 | 0.879 | 1 | 9 |
| Chl1 | 0.3198 | 0.833 | 0.466 | 1 | 9 |
| Kcmf1 | 0.3196 | 0.733 | 0.402 | 1 | 9 |
| Romo1 | 0.3195 | 1 | 0.958 | 1.31E-12 | 6 |
| Dnajb1 | 0.3194 | 0.462 | 0.32 | 1 | 7 |
| Msrb2 | 0.3193 | 0.306 | 0.16 | 0.0042 | 4 |
| Egfl7 | 0.3193 | 0.546 | 0.309 | 6.85E-06 | 1 |
| Rpl35a | 0.3193 | 1 | 0.991 | 9.26E-18 | 6 |
| Zfp608 | 0.3192 | 0.555 | 0.194 | 2.89E-09 | 2 |
| Gm26917 | 0.3189 | 0.583 | 0.397 | 0.0056 | 4 |
| Fam150b | 0.3188 | 0.22 | 0.073 | 7.84E-07 | 3 |
| Zfhx3 | 0.3188 | 0.935 | 0.706 | 1.30E-05 | 3 |
| Gabbr1 | 0.3186 | 0.99 | 0.946 | 1.89E-07 | 0 |
| Gm28050 | 0.3185 | 0.534 | 0.203 | 2.18E-07 | 2 |
| Dusp14 | 0.3183 | 0.433 | 0.119 | 1 | 9 |
| Nell2 | 0.3177 | 0.618 | 0.442 | 0.0065 | 1 |
| Lin7c | 0.3174 | 0.719 | 0.324 | 2.51E-05 | 2 |
| Disp2 | 0.3174 | 0.92 | 0.817 | 0.0012 | 0 |
| Rpl23 | 0.3173 | 0.986 | 0.987 | 0.0774 | 8 |
| Gabrg1 | 0.3172 | 0.815 | 0.549 | 1.10E-09 | 2 |
| Hpca | 0.3172 | 0.577 | 0.284 | 0.5014 | 8 |
| Gpx3 | 0.3172 | 0.685 | 0.4 | 0.0006 | 4 |
| Fjx1 | 0.3171 | 0.342 | 0.103 | 1.81E-09 | 1 |
| Nsmf | 0.3171 | 0.649 | 0.348 | 0.0003 | 5 |
| Scp2 | 0.317 | 0.795 | 0.527 | 1.33E-05 | 2 |
| Hspa8 | 0.317 | 1 | 0.997 | 0.2493 | 8 |
| Gsg1l | 0.3169 | 0.438 | 0.151 | 5.87E-07 | 2 |
| Lsamp | 0.3168 | 0.933 | 0.841 | 1 | 9 |
| Stt3b | 0.3167 | 0.549 | 0.266 | 0.0029 | 8 |
| Tmem237 | 0.3164 | 0.648 | 0.442 | 0.0019 | 4 |
| Magi3 | 0.3163 | 0.5 | 0.2 | 1.73E-05 | 2 |
| Nfyb | 0.3162 | 0.61 | 0.276 | 9.77E-05 | 2 |
| Ube2n | 0.3162 | 0.918 | 0.774 | 0.0052 | 2 |
| Lnp | 0.316 | 0.561 | 0.52 | 1 | 3 |
| Gskip | 0.3158 | 0.567 | 0.305 | 0.0076 | 5 |
| Socs2 | 0.3158 | 0.433 | 0.16 | 1 | 9 |

| | | | | | |
|---|---|---|---|---|---|
| Odc1 | 0.3157 | 0.651 | 0.56 | 0.0426 | 1 |
| Grin3a | 0.3155 | 0.567 | 0.333 | 1 | 9 |
| Bend6 | 0.3155 | 0.767 | 0.416 | 1 | 9 |
| Syngr1 | 0.3154 | 0.67 | 0.484 | 2.38E-06 | 0 |
| Serf2 | 0.3154 | 1 | 0.955 | 6.27E-07 | 6 |
| Rps5 | 0.3154 | 1 | 0.99 | 0.0011 | 8 |
| Ccdc106 | 0.315 | 0.667 | 0.373 | 1 | 9 |
| Ntm | 0.3148 | 0.885 | 0.794 | 0.0214 | 0 |
| mt-Nd4l | 0.3147 | 1 | 1 | 0.005 | 10 |
| 5031425I | 0.3147 | 0.541 | 0.252 | 8.30E-06 | 2 |
| Rps24 | 0.3146 | 1 | 0.997 | 4.78E-05 | 7 |
| Sarnp | 0.3144 | 0.788 | 0.567 | 0.0042 | 2 |
| B630019I | 0.3144 | 0.932 | 0.8 | 3.51E-05 | 2 |
| Bcl7b | 0.314 | 0.722 | 0.442 | 0.0056 | 5 |
| Rps7 | 0.314 | 1 | 0.979 | 1 | 8 |
| Ywhah | 0.3139 | 1 | 0.999 | 2.72E-06 | 7 |
| Pnmal2 | 0.3138 | 1 | 0.992 | 0.0362 | 3 |
| Myef2 | 0.3138 | 0.78 | 0.619 | 0.019 | 3 |
| Avpi1 | 0.3136 | 0.536 | 0.298 | 0.1826 | 5 |
| Pdzrn4 | 0.3135 | 0.493 | 0.249 | 7.78E-05 | 1 |
| Rap1gap | 0.3134 | 0.55 | 0.386 | 8.11E-05 | 0 |
| Myt1l | 0.3133 | 0.85 | 0.659 | 1.01E-06 | 0 |
| Rpl29 | 0.3133 | 0.986 | 0.933 | 9.32E-09 | 2 |
| H1f0 | 0.3133 | 0.589 | 0.259 | 0.0022 | 2 |
| Ciapin1 | 0.3133 | 0.649 | 0.369 | 0.0045 | 5 |
| Sap18 | 0.3133 | 0.966 | 0.767 | 1.33E-07 | 2 |
| Lrpap1 | 0.3131 | 0.895 | 0.884 | 0.0002 | 1 |
| Rps29 | 0.3131 | 1 | 0.997 | 6.35E-10 | 6 |
| Ttc3 | 0.313 | 1 | 1 | 5.79E-09 | 4 |
| Gng4 | 0.313 | 0.6 | 0.25 | 1 | 9 |
| 1700023I | 0.3129 | 0.671 | 0.349 | 0.0006 | 2 |
| Swi5 | 0.3129 | 1 | 0.933 | 3.43E-14 | 6 |
| Stmn2 | 0.3128 | 0.991 | 0.999 | 6.29E-06 | 4 |
| Rps8 | 0.3128 | 1 | 0.998 | 1.82E-09 | 6 |
| Pls3 | 0.3128 | 0.398 | 0.232 | 0.0108 | 4 |
| Rtn3 | 0.3127 | 1 | 0.974 | 2.28E-06 | 0 |
| Camk2a | 0.3127 | 0.8 | 0.713 | 1 | 9 |
| Mrpl50 | 0.3126 | 0.685 | 0.322 | 0.0073 | 2 |
| Ahi1 | 0.3125 | 1 | 1 | 1.59E-09 | 4 |
| Ndufb8 | 0.3125 | 1 | 0.972 | 1 | 9 |
| Uqcrb | 0.3124 | 1 | 0.976 | 1 | 9 |
| 1500009I | 0.3124 | 0.691 | 0.575 | 0.0266 | 1 |

| | | | | | |
|---|---|---|---|---|---|
| Pnrc1 | 0.3122 | 0.907 | 0.717 | 0.0006 | 5 |
| Pmvk | 0.3121 | 0.69 | 0.444 | 1 | 8 |
| Ptges3 | 0.3119 | 0.973 | 0.913 | 2.14E-06 | 2 |
| Hmgn2 | 0.3118 | 0.99 | 0.896 | 8.61E-05 | 5 |
| Mxra7 | 0.3118 | 0.467 | 0.194 | 1 | 9 |
| Mrpl15 | 0.3116 | 0.633 | 0.365 | 1 | 9 |
| Arxes1 | 0.3116 | 0.933 | 0.676 | 0.2624 | 9 |
| Crtac1 | 0.3114 | 0.567 | 0.203 | 1 | 9 |
| Ptms | 0.3113 | 1 | 0.969 | 0.0004 | 5 |
| Zbtb20 | 0.3113 | 0.898 | 0.722 | 0.0015 | 4 |
| Bola3 | 0.3112 | 0.692 | 0.415 | 8.83E-05 | 7 |
| Ndufa11 | 0.3112 | 1 | 0.973 | 1 | 9 |
| Prkra | 0.3111 | 0.606 | 0.355 | 1 | 8 |
| Ugp2 | 0.3111 | 0.562 | 0.221 | 0.0021 | 2 |
| Phlda1 | 0.3109 | 0.397 | 0.191 | 0.0121 | 7 |
| Gstm1 | 0.3109 | 0.5 | 0.15 | 1 | 9 |
| Cst6 | 0.3109 | 0.366 | 0.107 | 0.0005 | 8 |
| Plxna4 | 0.3107 | 0.454 | 0.189 | 2.96E-05 | 4 |
| Nol7 | 0.3107 | 0.742 | 0.613 | 0.0186 | 5 |
| Csrnp3 | 0.3106 | 0.495 | 0.319 | 6.46E-06 | 0 |
| Cadps2 | 0.3106 | 0.824 | 0.548 | 1 | 10 |
| Eif3m | 0.3105 | 0.746 | 0.503 | 1 | 8 |
| Ndufc1 | 0.3105 | 1 | 0.936 | 4.44E-09 | 6 |
| Pmepa1 | 0.3105 | 0.546 | 0.369 | 0.0021 | 1 |
| Creb3l1 | 0.3103 | 0.205 | 0.042 | 0.0014 | 7 |
| Cox7a2 | 0.3103 | 0.986 | 0.98 | 0.2291 | 8 |
| Opcml | 0.3102 | 0.855 | 0.638 | 0.0001 | 0 |
| Fam13a | 0.3101 | 0.388 | 0.185 | 2.53E-05 | 1 |
| Rps15a | 0.3101 | 0.993 | 0.98 | 1.29E-10 | 2 |
| Fxyd2 | 0.3101 | 0.423 | 0.166 | 0.3009 | 8 |
| Grin2b | 0.31 | 0.796 | 0.661 | 0.0427 | 4 |
| Arpp21 | 0.3098 | 0.692 | 0.334 | 2.66E-05 | 2 |
| Dynll1 | 0.3098 | 0.986 | 0.979 | 0.7628 | 8 |
| Dync1i1 | 0.3098 | 0.7 | 0.462 | 1 | 9 |
| Rprml | 0.3098 | 0.367 | 0.073 | 1 | 9 |
| Polr1d | 0.3097 | 0.915 | 0.809 | 0.9639 | 8 |
| Mcu | 0.3096 | 0.473 | 0.149 | 2.14E-07 | 2 |
| Pfdn5 | 0.3096 | 0.986 | 0.961 | 1 | 8 |
| Pls3 | 0.3096 | 0.535 | 0.229 | 0.0872 | 8 |
| 36324510 | 0.309 | 0.575 | 0.245 | 5.48E-11 | 2 |
| Vopp1 | 0.309 | 0.514 | 0.201 | 1.67E-05 | 2 |
| Rit2 | 0.3089 | 0.962 | 0.882 | 0.0039 | 7 |

| | | | | | |
|---|---|---|---|---|---|
| Rpl30 | 0.3087 | 1 | 0.988 | 0.0001 | 7 |
| Tekt2 | 0.3086 | 0.296 | 0.064 | 3.87E-10 | 1 |
| Agap2 | 0.3084 | 0.647 | 0.193 | 1 | 10 |
| Sez6l | 0.3083 | 0.691 | 0.592 | 0.0004 | 1 |
| Tmed2 | 0.3083 | 0.959 | 0.861 | 0.0111 | 3 |
| Fam159b | 0.3082 | 0.222 | 0.039 | 7.10E-07 | 4 |
| Serp2 | 0.3082 | 0.986 | 0.881 | 0.065 | 8 |
| Prim2 | 0.308 | 0.412 | 0.069 | 1 | 10 |
| Rap1gds1 | 0.308 | 0.722 | 0.41 | 0.02 | 5 |
| Rpl21 | 0.308 | 1 | 0.997 | 0.0001 | 7 |
| Naa60 | 0.3079 | 0.706 | 0.267 | 1 | 10 |
| Gfra1 | 0.3078 | 0.27 | 0.071 | 3.03E-11 | 0 |
| Zic1 | 0.3078 | 0.449 | 0.181 | 0.003 | 7 |
| Kcnip1 | 0.3077 | 0.5 | 0.163 | 1 | 9 |
| Fxyd6 | 0.3076 | 1 | 0.99 | 1.62E-08 | 4 |
| Tppp3 | 0.3075 | 0.704 | 0.445 | 1 | 8 |
| Cdc26 | 0.3075 | 0.753 | 0.404 | 0.0007 | 2 |
| Prdx6 | 0.3074 | 0.704 | 0.612 | 0.0678 | 1 |
| Yipf4 | 0.3074 | 0.774 | 0.473 | 0.0078 | 2 |
| Rsph1 | 0.3073 | 0.505 | 0.181 | 1.05E-05 | 5 |
| Borcs7 | 0.3072 | 0.733 | 0.492 | 1 | 9 |
| Marcks | 0.3072 | 0.979 | 0.943 | 0.0007 | 5 |
| Cpne8 | 0.3072 | 0.367 | 0.159 | 1 | 9 |
| Bloc1s5 | 0.3069 | 0.493 | 0.262 | 0.264 | 8 |
| Orai3 | 0.3069 | 0.495 | 0.225 | 0.0001 | 5 |
| Spock1 | 0.3068 | 0.667 | 0.371 | 0.0018 | 4 |
| Tmem25( | 0.3066 | 0.795 | 0.664 | 0.3793 | 7 |
| Psma7 | 0.3066 | 0.99 | 0.931 | 4.81E-07 | 6 |
| Idnk | 0.3065 | 0.634 | 0.278 | 0.0225 | 8 |
| 2010005I | 0.3065 | 0.299 | 0.155 | 1 | 5 |
| Oxr1 | 0.3064 | 0.574 | 0.349 | 0.0014 | 4 |
| Ndufa13 | 0.3064 | 0.967 | 0.983 | 1 | 9 |
| Bzw2 | 0.3064 | 0.521 | 0.291 | 0.1222 | 8 |
| Pfdn1 | 0.3063 | 0.979 | 0.905 | 3.85E-10 | 2 |
| Prkacb | 0.3062 | 0.98 | 0.943 | 1.59E-07 | 0 |
| Abhd12 | 0.3061 | 0.89 | 0.723 | 3.68E-07 | 0 |
| Il1rapl1 | 0.3059 | 0.512 | 0.371 | 0.4652 | 3 |
| Pnoc | 0.3059 | 0.507 | 0.13 | 0.165 | 8 |
| Gnb4 | 0.3058 | 0.563 | 0.213 | 0.0231 | 8 |
| Fbxo27 | 0.3057 | 0.353 | 0.035 | 1 | 10 |
| Bex2 | 0.3055 | 1 | 1 | 0.1491 | 8 |
| Fkbp3 | 0.3054 | 1 | 0.966 | 0.0004 | 7 |

| | | | | | |
|---|---|---|---|---|---|
| Smim8 | 0.3054 | 0.678 | 0.318 | 3.81E-05 | 2 |
| Prkab2 | 0.3053 | 0.471 | 0.108 | 1 | 10 |
| Arx | 0.3053 | 0.4 | 0.127 | 1 | 9 |
| Snrpg | 0.3053 | 0.932 | 0.79 | 3.45E-07 | 2 |
| Higd1a | 0.3052 | 0.835 | 0.778 | 0.1604 | 5 |
| Thumpd1 | 0.3052 | 0.596 | 0.31 | 0.005 | 2 |
| Qpct | 0.3052 | 0.496 | 0.415 | 0.3953 | 3 |
| Ptpn5 | 0.3052 | 0.572 | 0.313 | 1.41E-05 | 1 |
| Polr2l | 0.305 | 0.781 | 0.597 | 0.0076 | 2 |
| Tceb2 | 0.3048 | 1 | 0.992 | 6.16E-06 | 7 |
| Hsp90ab1 | 0.3048 | 1 | 1 | 1.10E-05 | 8 |
| Rpl13 | 0.3047 | 1 | 0.996 | 6.92E-12 | 6 |
| Mdh1 | 0.3047 | 1 | 0.983 | 0.0001 | 5 |
| Hmx2 | 0.3047 | 0.407 | 0.11 | 1.38E-06 | 4 |
| Inpp5f | 0.3046 | 0.759 | 0.683 | 0.1233 | 4 |
| Vamp4 | 0.3046 | 0.699 | 0.428 | 1 | 2 |
| Galntl6 | 0.3045 | 0.5 | 0.126 | 1 | 9 |
| Slc25a33 | 0.3044 | 0.685 | 0.344 | 0.0027 | 2 |
| Spock2 | 0.3043 | 1 | 0.84 | 1 | 3 |
| Ubl5 | 0.3043 | 1 | 0.972 | 6.60E-13 | 6 |
| Cox14 | 0.3042 | 0.9 | 0.723 | 1 | 9 |
| 1700001 | 0.3042 | 0.679 | 0.38 | 0.0002 | 7 |
| Khdrbs1 | 0.3042 | 0.836 | 0.625 | 0.0059 | 2 |
| Slc7a14 | 0.3041 | 0.521 | 0.29 | 0.0804 | 8 |
| Dgkk | 0.304 | 0.342 | 0.1 | 1.76E-06 | 1 |
| Smim10l | 0.304 | 0.959 | 0.859 | 0.0002 | 2 |
| Sepw1 | 0.3039 | 1 | 0.995 | 7.63E-14 | 6 |
| H2afz | 0.3038 | 0.991 | 0.978 | 0.0031 | 4 |
| Rpl31 | 0.3038 | 1 | 0.936 | 4.76E-09 | 6 |
| Zfand6 | 0.3037 | 0.68 | 0.427 | 0.0285 | 5 |
| Fam173a | 0.3037 | 0.984 | 0.908 | 1 | 3 |
| Pgk1 | 0.3035 | 0.732 | 0.455 | 0.0349 | 5 |
| Borcs5 | 0.3034 | 0.633 | 0.277 | 1 | 9 |
| Cox8a | 0.3034 | 1 | 0.998 | 8.22E-13 | 6 |
| Dner | 0.3034 | 0.976 | 0.816 | 1 | 3 |
| Dpp6 | 0.3033 | 0.905 | 0.831 | 0.0019 | 0 |
| Bend5 | 0.3033 | 0.521 | 0.194 | 0.008 | 8 |
| Zcchc18 | 0.3032 | 1 | 0.999 | 1 | 3 |
| Gm2694 | 0.3031 | 0.294 | 0.039 | 1 | 10 |
| Galnt18 | 0.3029 | 0.37 | 0.157 | 7.01E-05 | 4 |
| Sgip1 | 0.3029 | 0.915 | 0.756 | 3.67E-08 | 0 |
| Btf3 | 0.3029 | 1 | 0.969 | 0.1176 | 8 |

| | | | | | |
|---|---|---|---|---|---|
| Caln1 | 0.3028 | 0.467 | 0.144 | 1 | 9 |
| Ehd3 | 0.3027 | 0.5 | 0.273 | 1 | 9 |
| Magi1 | 0.3025 | 0.712 | 0.499 | 4.64E-05 | 2 |
| Higd1a | 0.3025 | 0.91 | 0.773 | 0.0015 | 7 |
| Cirbp | 0.3024 | 1 | 0.97 | 3.01E-06 | 6 |
| Syngr3 | 0.3023 | 0.907 | 0.908 | 0.0424 | 4 |
| Nrxn2 | 0.3022 | 1 | 0.963 | 1 | 10 |
| Ephb1 | 0.3022 | 0.641 | 0.342 | 0.0026 | 7 |
| 0610010I | 0.3022 | 0.596 | 0.248 | 0.0001 | 2 |
| Gucy1a3 | 0.3022 | 0.367 | 0.129 | 1 | 9 |
| Hdac9 | 0.302 | 0.473 | 0.181 | 3.84E-06 | 2 |
| Rpl37 | 0.302 | 1 | 0.996 | 2.85E-08 | 6 |
| Adgra1 | 0.3018 | 0.657 | 0.385 | 0.0017 | 4 |
| Sipa1l1 | 0.3018 | 0.533 | 0.183 | 1 | 9 |
| Paip2 | 0.3018 | 0.973 | 0.831 | 2.13E-06 | 2 |
| Adcyap1r | 0.3016 | 0.515 | 0.354 | 6.78E-05 | 0 |
| Phyhipl | 0.3016 | 0.849 | 0.626 | 0.0001 | 2 |
| Slc6a1 | 0.3016 | 0.74 | 0.638 | 1 | 3 |
| Rplp1 | 0.3016 | 1 | 0.991 | 8.91E-11 | 6 |
| Ndufc1 | 0.3015 | 1 | 0.94 | 1 | 9 |
| Cntnap2 | 0.3015 | 0.632 | 0.471 | 0.0045 | 1 |
| Arglu1 | 0.3015 | 0.951 | 0.839 | 0.0077 | 3 |
| Vat1l | 0.3013 | 0.795 | 0.581 | 0.0065 | 7 |
| Uqcr11 | 0.3013 | 1 | 0.968 | 1.55E-14 | 6 |
| Commd1 | 0.3012 | 0.685 | 0.361 | 0.0011 | 2 |
| Rrn3 | 0.3011 | 0.664 | 0.314 | 0.0085 | 2 |
| Nudt16 | 0.3011 | 0.465 | 0.167 | 0.0141 | 8 |
| 5330434 | 0.3011 | 0.852 | 0.718 | 0.0114 | 4 |
| Golga7b | 0.3009 | 0.733 | 0.434 | 1 | 9 |
| Napa | 0.3008 | 0.9 | 0.789 | 1 | 9 |
| Cry1 | 0.3008 | 0.432 | 0.165 | 0.0138 | 2 |
| Btg1 | 0.3007 | 0.907 | 0.765 | 0.0351 | 5 |
| Nfkbiz | 0.3006 | 0.256 | 0.03 | 2.11E-09 | 7 |
| Mgat1 | 0.3006 | 0.633 | 0.286 | 1 | 9 |
| Slc25a44 | 0.3005 | 0.633 | 0.311 | 1 | 9 |
| Scgn | 0.3005 | 0.197 | 0.023 | 0.0004 | 8 |
| Plk3 | 0.3005 | 0.412 | 0.078 | 1 | 10 |
| Slc12a2 | 0.3004 | 0.372 | 0.165 | 0.019 | 7 |
| Tmx2 | 0.3004 | 0.95 | 0.876 | 7.77E-06 | 0 |
| Klhdc8b | 0.3002 | 0.654 | 0.53 | 1 | 7 |
| Ppp1r9a | 0.3 | 0.745 | 0.529 | 3.59E-05 | 0 |
| Uqcrb | 0.3 | 0.99 | 0.976 | 0.0003 | 5 |

| | | | | | |
|---|---|---|---|---|---|
| Irs2 | 0.3 | 0.6 | 0.28 | 1 | 9 |
| Apba2 | 0.3 | 0.567 | 0.278 | 1 | 9 |
| Isca2 | 0.3 | 0.742 | 0.539 | 0.0125 | 5 |
| Krt73 | 0.2998 | 0.211 | 0.002 | 1.47E-13 | 8 |
| Caly | 0.2995 | 1 | 0.999 | 4.43E-13 | 1 |
| Ndufa12 | 0.2993 | 1 | 0.886 | 1 | 9 |
| Psme2 | 0.2992 | 0.718 | 0.431 | 0.6104 | 8 |
| Rps9 | 0.2992 | 1 | 0.972 | 2.77E-05 | 5 |
| Cstb | 0.2991 | 0.667 | 0.329 | 1 | 9 |
| Blcap | 0.2991 | 0.943 | 0.814 | 9.91E-05 | 3 |
| Napb | 0.2991 | 0.6 | 0.445 | 1.68E-05 | 0 |
| Rpl9 | 0.299 | 1 | 0.987 | 1 | 8 |
| Lmo2 | 0.2989 | 0.324 | 0.092 | 0.7399 | 8 |
| Arf3 | 0.2988 | 0.935 | 0.803 | 0.0009 | 0 |
| Rps11 | 0.2988 | 1 | 0.994 | 1 | 8 |
| Qrfpr | 0.2987 | 0.294 | 0.032 | 1 | 10 |
| Fam96a | 0.2987 | 0.685 | 0.367 | 7.66E-05 | 2 |
| Lin7a | 0.2987 | 0.866 | 0.668 | 0.0093 | 5 |
| Camk2g | 0.2986 | 0.704 | 0.49 | 6.89E-05 | 1 |
| Tubb2b | 0.2984 | 0.831 | 0.585 | 0.2399 | 8 |
| Resp18 | 0.2983 | 1 | 0.999 | 1.39E-05 | 4 |
| St8sia6 | 0.2982 | 0.308 | 0.086 | 1.18E-05 | 7 |
| Pin1 | 0.2982 | 0.859 | 0.694 | 0.955 | 8 |
| Tpgs1 | 0.2981 | 0.833 | 0.443 | 1 | 9 |
| Rsl24d1 | 0.2981 | 0.719 | 0.398 | 0.0045 | 2 |
| Minos1 | 0.2981 | 0.967 | 0.912 | 1 | 9 |
| Pcp4 | 0.2979 | 0.99 | 0.87 | 2.54E-05 | 6 |
| Amt | 0.2979 | 0.418 | 0.132 | 2.77E-05 | 2 |
| Arhgap6 | 0.2978 | 0.866 | 0.581 | 0.0002 | 5 |
| Vps8 | 0.2977 | 0.719 | 0.44 | 1 | 2 |
| Celf2 | 0.2975 | 0.725 | 0.618 | 0.0275 | 0 |
| Nptn | 0.2975 | 0.951 | 0.81 | 1 | 3 |
| Cbln4 | 0.2974 | 0.479 | 0.184 | 0.011 | 2 |
| Nop10 | 0.2971 | 1 | 0.947 | 7.68E-08 | 6 |
| Kcnc1 | 0.2971 | 0.68 | 0.518 | 0.0008 | 0 |
| Ndufb9 | 0.297 | 1 | 0.958 | 0.0002 | 7 |
| Tmem132 | 0.2969 | 0.598 | 0.25 | 0.0016 | 5 |
| Rpl27a | 0.2968 | 1 | 0.994 | 6.69E-05 | 7 |
| Gria3 | 0.2968 | 0.441 | 0.165 | 4.89E-05 | 1 |
| Csgalnact | 0.2967 | 0.533 | 0.15 | 1 | 9 |
| Mmd2 | 0.2967 | 0.343 | 0.112 | 1.29E-06 | 4 |
| Med30 | 0.2967 | 0.685 | 0.367 | 0.0071 | 2 |

| | | | | | |
|---|---|---|---|---|---|
| Gm21092 | 0.2967 | 0.675 | 0.632 | 1 | 3 |
| Mrpl54 | 0.2965 | 0.897 | 0.651 | 1.62E-06 | 2 |
| Ass1 | 0.2964 | 0.451 | 0.176 | 0.0769 | 8 |
| Tubb3 | 0.2963 | 0.986 | 0.945 | 0.1246 | 8 |
| Scn2b | 0.2963 | 0.585 | 0.499 | 0.0913 | 0 |
| Anxa5 | 0.2961 | 0.592 | 0.218 | 0.3038 | 8 |
| Htra1 | 0.296 | 0.267 | 0.012 | 0.0015 | 9 |
| Gm9866 | 0.296 | 0.4 | 0.109 | 1 | 9 |
| Bnip3 | 0.296 | 0.637 | 0.312 | 0.4871 | 2 |
| Ing1 | 0.296 | 0.664 | 0.349 | 6.50E-05 | 2 |
| Psmb4 | 0.2959 | 0.93 | 0.784 | 0.3903 | 8 |
| Timm8b | 0.2959 | 1 | 0.935 | 3.70E-11 | 6 |
| Kcnip4 | 0.2958 | 0.365 | 0.197 | 1.83E-05 | 0 |
| Erbb4 | 0.2957 | 0.315 | 0.101 | 0.001 | 4 |
| Igfbpl1 | 0.2957 | 0.432 | 0.125 | 5.95E-13 | 2 |
| Kif2a | 0.2955 | 0.8 | 0.493 | 1 | 9 |
| Gm16105 | 0.2954 | 0.545 | 0.456 | 1.09E-05 | 3 |
| B4galt6 | 0.2954 | 0.711 | 0.574 | 0.0006 | 1 |
| Mrto4 | 0.2954 | 0.767 | 0.411 | 1 | 9 |
| Tspan13 | 0.2954 | 0.934 | 0.822 | 8.52E-07 | 1 |
| Ndufv3 | 0.2954 | 0.959 | 0.936 | 1.99E-05 | 5 |
| Ndfip2 | 0.2953 | 0.618 | 0.566 | 0.0003 | 3 |
| Mrps22 | 0.2952 | 0.433 | 0.189 | 1 | 9 |
| Myo10 | 0.2952 | 0.568 | 0.271 | 2.94E-06 | 2 |
| Mxra7 | 0.2951 | 0.395 | 0.171 | 2.95E-05 | 1 |
| Arpp21 | 0.295 | 0.512 | 0.364 | 6.20E-06 | 3 |
| Chst8 | 0.2949 | 0.647 | 0.3 | 1 | 10 |
| 6330403/ | 0.2948 | 0.451 | 0.141 | 0.0169 | 8 |
| Rpl24 | 0.2948 | 1 | 0.998 | 0.0257 | 8 |
| Emd | 0.2946 | 0.577 | 0.324 | 0.0022 | 7 |
| Scg2 | 0.2946 | 1 | 0.959 | 1 | 7 |
| Stx7 | 0.2945 | 0.884 | 0.661 | 0.0001 | 2 |
| Insig1 | 0.2944 | 0.603 | 0.312 | 0.0964 | 2 |
| Stx1b | 0.2944 | 0.595 | 0.439 | 1.18E-06 | 0 |
| Slc25a33 | 0.2943 | 0.641 | 0.37 | 9.42E-05 | 7 |
| Vstm2a | 0.2942 | 0.444 | 0.248 | 0.0354 | 4 |
| Sox6 | 0.2942 | 0.2 | 0.007 | 0.369 | 9 |
| Dynlrb1 | 0.2941 | 1 | 0.942 | 3.12E-09 | 6 |
| Npy5r | 0.2941 | 0.464 | 0.164 | 0.0003 | 5 |
| Mrpl18 | 0.294 | 0.732 | 0.525 | 1.02E-05 | 6 |
| Gabarapl2 | 0.294 | 1 | 0.987 | 2.24E-06 | 7 |
| Rplp2 | 0.294 | 1 | 0.985 | 0.0004 | 7 |

| | | | | | |
|---|---|---|---|---|---|
| Ptprs | 0.294 | 0.93 | 0.831 | 0.0002 | 0 |
| Gabre | 0.2938 | 0.306 | 0.138 | 0.1299 | 4 |
| A230065I | 0.2937 | 0.38 | 0.136 | 0.0016 | 4 |
| Serpini1 | 0.2936 | 0.355 | 0.147 | 6.55E-09 | 0 |
| Spin1 | 0.2936 | 0.726 | 0.458 | 1 | 2 |
| Tmed3 | 0.2935 | 0.737 | 0.612 | 0.0009 | 1 |
| Rps24 | 0.2935 | 1 | 0.997 | 1 | 8 |
| Fam96a | 0.2935 | 0.67 | 0.384 | 0.0118 | 5 |
| Psd | 0.2935 | 0.911 | 0.795 | 1 | 3 |
| Nptn | 0.2934 | 0.925 | 0.804 | 5.63E-06 | 0 |
| Rgs17 | 0.2933 | 0.822 | 0.67 | 0.0004 | 1 |
| Fkbp4 | 0.2933 | 0.845 | 0.744 | 1 | 8 |
| Vapa | 0.2932 | 0.979 | 0.957 | 2.86E-05 | 2 |
| Rad23a | 0.2932 | 0.629 | 0.426 | 1 | 5 |
| Nedd4 | 0.2932 | 1 | 0.936 | 1 | 3 |
| Dctn3 | 0.2932 | 1 | 0.893 | 1 | 9 |
| Npy6r | 0.2931 | 0.392 | 0.092 | 9.60E-06 | 5 |
| Rpl10 | 0.2931 | 0.972 | 0.937 | 1 | 8 |
| Rps14 | 0.2931 | 0.986 | 0.926 | 0.458 | 8 |
| Tma7 | 0.293 | 1 | 0.965 | 6.97E-11 | 6 |
| Reln | 0.293 | 0.2 | 0.023 | 8.06E-13 | 0 |
| Cfl1 | 0.2929 | 1 | 0.99 | 0.0225 | 8 |
| Fyttd1 | 0.2929 | 0.87 | 0.654 | 0.0097 | 2 |
| 2210016I | 0.2926 | 0.933 | 0.89 | 1 | 9 |
| Pop5 | 0.2925 | 0.817 | 0.72 | 1 | 8 |
| Dusp4 | 0.2924 | 0.244 | 0.021 | 1.12E-09 | 7 |
| Gtf3c6 | 0.2924 | 0.567 | 0.227 | 1 | 9 |
| Hspe1 | 0.2923 | 0.987 | 0.96 | 0.0143 | 7 |
| 2900011( | 0.2921 | 0.952 | 0.882 | 1.89E-07 | 2 |
| Kif1a | 0.2921 | 0.95 | 0.91 | 0.0019 | 0 |
| Anp32e | 0.2921 | 0.608 | 0.292 | 0.0043 | 5 |
| Tmem59l | 0.292 | 0.935 | 0.92 | 0.1821 | 4 |
| Tomm34 | 0.292 | 0.746 | 0.51 | 0.6939 | 8 |
| Nrsn1 | 0.2919 | 0.98 | 0.954 | 9.60E-06 | 1 |
| Gucy1b3 | 0.2919 | 0.7 | 0.37 | 1 | 9 |
| Pik3r1 | 0.2919 | 0.567 | 0.241 | 1 | 9 |
| Atp5g2 | 0.2919 | 0.979 | 0.972 | 0.0012 | 5 |
| Susd4 | 0.2919 | 0.572 | 0.334 | 0.0001 | 1 |
| Cited1 | 0.2918 | 0.257 | 0.148 | 1 | 1 |
| Sez6 | 0.2918 | 0.285 | 0.126 | 1.36E-05 | 0 |
| Zcrb1 | 0.2918 | 0.962 | 0.796 | 0.0051 | 7 |
| Epm2aip1 | 0.2917 | 0.81 | 0.612 | 3.52E-06 | 0 |

| | | | | | |
|---|---|---|---|---|---|
| Nenf | 0.2917 | 1 | 0.99 | 1 | 3 |
| Ywhaq | 0.2916 | 1 | 0.984 | 2.83E-08 | 2 |
| Plekhm2 | 0.2914 | 0.433 | 0.173 | 0.002 | 5 |
| Sdhaf4 | 0.2912 | 0.904 | 0.676 | 3.92E-07 | 2 |
| Sec11a | 0.2912 | 0.775 | 0.562 | 0.4289 | 8 |
| Nt5m | 0.2912 | 0.691 | 0.446 | 0.0921 | 5 |
| Mycbp2 | 0.2911 | 0.963 | 0.884 | 0.0001 | 4 |
| 1110004 | 0.2911 | 0.551 | 0.399 | 0.0732 | 7 |
| Enc1 | 0.291 | 0.38 | 0.246 | 0.0053 | 0 |
| Cxxc4 | 0.291 | 0.815 | 0.659 | 0.0006 | 2 |
| Rbfox1 | 0.2909 | 0.38 | 0.173 | 1.53E-07 | 0 |
| Polr2d | 0.2909 | 0.678 | 0.328 | 3.04E-06 | 2 |
| Hadhb | 0.2908 | 0.536 | 0.298 | 0.0852 | 5 |
| Rab9 | 0.2908 | 0.637 | 0.332 | 0.12 | 2 |
| Lrfn5 | 0.2908 | 0.658 | 0.447 | 0.0004 | 1 |
| A730017 | 0.2907 | 1 | 0.928 | 0.008 | 8 |
| Mfsd6 | 0.2907 | 0.602 | 0.39 | 0.0087 | 4 |
| Scn9a | 0.2907 | 0.75 | 0.579 | 0.0507 | 4 |
| Gstm7 | 0.2907 | 0.481 | 0.327 | 0.6319 | 4 |
| Prdx6 | 0.2906 | 0.845 | 0.61 | 0.4166 | 8 |
| Ctsd | 0.2904 | 0.76 | 0.642 | 0.0127 | 0 |
| Sdhaf1 | 0.2904 | 0.598 | 0.35 | 0.0692 | 5 |
| Cpne8 | 0.2901 | 0.408 | 0.148 | 0.0522 | 8 |
| Tbkbp1 | 0.2901 | 0.471 | 0.119 | 1 | 10 |
| Ppp3ca | 0.29 | 0.921 | 0.774 | 0.0002 | 1 |
| Galr1 | 0.29 | 0.276 | 0.035 | 1.93E-13 | 1 |
| Tmem178 | 0.2899 | 0.633 | 0.343 | 1 | 9 |
| Pdpk1 | 0.2899 | 0.6 | 0.359 | 1 | 9 |
| Edil3 | 0.2899 | 0.75 | 0.647 | 1 | 4 |
| Syf2 | 0.2898 | 0.705 | 0.466 | 0.0081 | 7 |
| Nrxn1 | 0.2896 | 0.954 | 0.919 | 0.01 | 4 |
| Dpp10 | 0.2894 | 0.395 | 0.277 | 0.1616 | 0 |
| Pcbd1 | 0.2894 | 0.822 | 0.705 | 0.0048 | 1 |
| Ngfrap1 | 0.2893 | 0.986 | 0.993 | 0.9036 | 8 |
| 1-Mar | 0.2892 | 0.733 | 0.42 | 1 | 9 |
| Ybx1 | 0.289 | 0.808 | 0.714 | 1 | 7 |
| Olfm1 | 0.289 | 0.944 | 0.842 | 0.0236 | 8 |
| Nr3c1 | 0.289 | 0.606 | 0.264 | 0.135 | 8 |
| Lef1 | 0.2888 | 0.296 | 0.05 | 6.70E-08 | 4 |
| Trnp1 | 0.2888 | 0.5 | 0.233 | 1.09E-06 | 1 |
| Syt4 | 0.2886 | 0.984 | 0.862 | 1 | 3 |
| Zfand6 | 0.2886 | 0.74 | 0.405 | 0.0002 | 2 |

| | | | | | |
|---|---|---|---|---|---|
| Ptprd | 0.2885 | 0.972 | 0.83 | 7.00E-07 | 8 |
| Coro1b | 0.2885 | 0.529 | 0.126 | 1 | 10 |
| Snx7 | 0.2883 | 0.296 | 0.133 | 1 | 8 |
| Kiss1 | 0.2883 | 0.079 | 0.001 | 7.72E-07 | 1 |
| Pafah1b3 | 0.2882 | 0.925 | 0.728 | 7.06E-10 | 2 |
| Pgrmc1 | 0.2882 | 1 | 0.999 | 6.71E-15 | 5 |
| Rps3a1 | 0.2881 | 1 | 0.977 | 4.66E-14 | 6 |
| Adgrl2 | 0.288 | 0.3 | 0.141 | 1.19E-06 | 0 |
| Mpc1 | 0.2879 | 0.91 | 0.802 | 0.2197 | 7 |
| Ppp1r3d | 0.2879 | 0.294 | 0.015 | 1 | 10 |
| Fam214b | 0.2878 | 0.495 | 0.243 | 0.0044 | 5 |
| Atp5l | 0.2876 | 0.986 | 0.974 | 1 | 8 |
| 1110008I | 0.2876 | 0.784 | 0.609 | 0.0363 | 5 |
| Ntng1 | 0.2876 | 0.724 | 0.286 | 3.98E-07 | 1 |
| H2afz | 0.2876 | 1 | 0.979 | 1 | 9 |
| Nop56 | 0.2876 | 0.776 | 0.651 | 0.0023 | 1 |
| Nrip1 | 0.2876 | 0.349 | 0.154 | 0.0002 | 1 |
| Mid1 | 0.2875 | 0.353 | 0.054 | 1 | 10 |
| Rpl13a | 0.2874 | 0.986 | 0.945 | 0.0333 | 8 |
| Uqcrq | 0.2874 | 1 | 0.967 | 5.86E-05 | 7 |
| Rpl36 | 0.2873 | 1 | 0.992 | 1.31E-12 | 2 |
| Lyrm4 | 0.2873 | 0.61 | 0.309 | 0.0723 | 2 |
| Rhob | 0.2873 | 0.767 | 0.609 | 0.8454 | 2 |
| Grm8 | 0.2872 | 0.367 | 0.11 | 1 | 9 |
| Fuom | 0.2871 | 0.68 | 0.532 | 0.1702 | 5 |
| Calm2 | 0.2869 | 1 | 1 | 1 | 9 |
| Tmcc3 | 0.2869 | 0.526 | 0.28 | 2.47E-05 | 1 |
| Rprm | 0.2869 | 0.577 | 0.393 | 2.48E-07 | 3 |
| Ntsr1 | 0.2868 | 0.278 | 0.03 | 6.58E-13 | 4 |
| Rpl17 | 0.2867 | 0.986 | 0.915 | 0.5187 | 8 |
| Mrps23 | 0.2866 | 0.767 | 0.49 | 1 | 9 |
| Fbxw4 | 0.2865 | 0.452 | 0.163 | 2.64E-07 | 2 |
| Parm1 | 0.2865 | 0.836 | 0.55 | 6.33E-13 | 2 |
| Aig1 | 0.2864 | 0.943 | 0.796 | 1 | 3 |
| Rpl22l1 | 0.2864 | 1 | 0.997 | 5.84E-14 | 2 |
| Atpif1 | 0.2863 | 1 | 0.993 | 1 | 9 |
| Tmem50a | 0.2863 | 0.928 | 0.815 | 1.94E-07 | 1 |
| Malat1 | 0.2862 | 1 | 0.999 | 1.20E-14 | 2 |
| Adgrg2 | 0.2862 | 0.392 | 0.055 | 4.65E-09 | 5 |
| Extl1 | 0.2861 | 0.353 | 0.016 | 1 | 10 |
| Rbbp4 | 0.2861 | 0.897 | 0.744 | 0.0009 | 2 |
| Fam213b | 0.286 | 0.662 | 0.52 | 1 | 8 |

| | | | | | |
|---|---|---|---|---|---|
| Syt7 | 0.2859 | 0.395 | 0.149 | 1.76E-05 | 1 |
| Mphosph | 0.2858 | 0.7 | 0.459 | 1 | 9 |
| Armcx3 | 0.2855 | 0.538 | 0.35 | 0.036 | 7 |
| Ppp2r2b | 0.2855 | 0.651 | 0.473 | 0.0013 | 1 |
| Rheb | 0.2854 | 0.979 | 0.9 | 0.007 | 2 |
| Pfn2 | 0.2854 | 1 | 0.981 | 0.0005 | 7 |
| Dlx6os1 | 0.2851 | 0.329 | 0.099 | 3.03E-07 | 1 |
| Rwdd2a | 0.2851 | 0.671 | 0.448 | 0.0019 | 1 |
| Rps10 | 0.285 | 1 | 0.983 | 1 | 8 |
| Frs3 | 0.2849 | 0.514 | 0.227 | 0.1594 | 2 |
| Rpl30 | 0.2848 | 1 | 0.989 | 1 | 8 |
| Them4 | 0.2847 | 0.667 | 0.403 | 1 | 9 |
| Rogdi | 0.2846 | 0.859 | 0.737 | 0.189 | 8 |
| Arhgef15 | 0.2846 | 0.437 | 0.169 | 0.1467 | 8 |
| Eef1e1 | 0.2846 | 0.788 | 0.521 | 0.1015 | 2 |
| Fau | 0.2844 | 1 | 0.997 | 1 | 8 |
| Dnm1 | 0.2844 | 0.813 | 0.655 | 0.6797 | 3 |
| Dst | 0.2842 | 0.805 | 0.603 | 0.0001 | 0 |
| Ralgapa1 | 0.284 | 0.561 | 0.506 | 1 | 3 |
| Nell1 | 0.284 | 0.431 | 0.281 | 0.1457 | 3 |
| Fam213a | 0.284 | 0.75 | 0.553 | 0.0042 | 4 |
| Rabggtb | 0.2839 | 0.623 | 0.345 | 1 | 2 |
| B3gat2 | 0.2839 | 0.753 | 0.416 | 3.65E-05 | 2 |
| Rec8 | 0.2838 | 0.352 | 0.091 | 0.0014 | 8 |
| Ahi1 | 0.2838 | 1 | 1 | 0.0126 | 9 |
| Tmem59 | 0.2837 | 0.984 | 0.917 | 1 | 3 |
| Phf5a | 0.2837 | 0.667 | 0.389 | 1 | 9 |
| Cggbp1 | 0.2835 | 0.671 | 0.396 | 0.2899 | 2 |
| Sstr3 | 0.2834 | 0.353 | 0.016 | 1 | 10 |
| Reps1 | 0.2833 | 0.521 | 0.225 | 0.0002 | 2 |
| Thada | 0.2833 | 0.353 | 0.057 | 1 | 10 |
| Plppr1 | 0.2832 | 0.407 | 0.249 | 0.0689 | 4 |
| H3f3b | 0.2832 | 1 | 0.998 | 9.03E-06 | 5 |
| Rpl13 | 0.2832 | 1 | 0.996 | 0.0006 | 7 |
| Ttc3 | 0.2832 | 1 | 1 | 1.09E-09 | 0 |
| Adamtsl2 | 0.2832 | 0.25 | 0.046 | 3.56E-08 | 4 |
| Mir124-2 | 0.2831 | 0.588 | 0.287 | 0.1645 | 5 |
| Ppp1r14c | 0.2831 | 0.589 | 0.307 | 0.0049 | 2 |
| Zfp207 | 0.283 | 0.753 | 0.422 | 0.0005 | 2 |
| Atp6v1g1 | 0.283 | 0.962 | 0.941 | 0.0094 | 7 |
| Pbx3 | 0.283 | 0.239 | 0.033 | 8.25E-05 | 8 |
| Snrnp27 | 0.2829 | 0.915 | 0.735 | 0.0259 | 8 |

| | | | | | |
|---|---|---|---|---|---|
| Pcdh15 | 0.2828 | 0.303 | 0.094 | 6.14E-06 | 1 |
| Cdkn1c | 0.2828 | 0.179 | 0.116 | 0.7104 | 3 |
| Unc5d | 0.2827 | 0.481 | 0.333 | 1 | 4 |
| Tpt1 | 0.2826 | 1 | 0.986 | 2.76E-06 | 5 |
| Fis1 | 0.2826 | 1 | 0.927 | 5.58E-06 | 5 |
| Trpm3 | 0.2823 | 0.317 | 0.179 | 1.30E-06 | 3 |
| Reep1 | 0.2822 | 0.63 | 0.446 | 0.1167 | 4 |
| Ipo5 | 0.2822 | 0.567 | 0.308 | 1 | 9 |
| Slc38a1 | 0.2822 | 0.829 | 0.662 | 1 | 3 |
| 2610001. | 0.2821 | 0.438 | 0.158 | 0.0105 | 2 |
| Rap1a | 0.282 | 0.692 | 0.361 | 0.0035 | 2 |
| Hspb11 | 0.2819 | 0.582 | 0.247 | 0.0003 | 2 |
| Usp9x | 0.2819 | 0.943 | 0.794 | 1 | 3 |
| Rps19 | 0.2819 | 1 | 0.988 | 7.53E-13 | 2 |
| Syt10 | 0.2818 | 0.894 | 0.614 | 0.0002 | 3 |
| Slc25a36 | 0.2817 | 0.747 | 0.39 | 8.82E-05 | 2 |
| Mfap1b | 0.2817 | 0.658 | 0.316 | 0.4315 | 2 |
| Cntnap5a | 0.2816 | 0.546 | 0.289 | 0.0847 | 4 |
| Efcab6 | 0.2816 | 0.235 | 0.016 | 1 | 10 |
| Calm1 | 0.2816 | 1 | 1 | 3.19E-05 | 7 |
| Tmie | 0.2816 | 0.493 | 0.3 | 0.0011 | 1 |
| Fgfr1op2 | 0.2815 | 0.726 | 0.383 | 0.0684 | 2 |
| Cnot6 | 0.2814 | 0.568 | 0.293 | 0.0535 | 2 |
| Shisa9 | 0.2814 | 0.472 | 0.199 | 0.0001 | 4 |
| Gm1007€ | 0.2814 | 0.938 | 0.692 | 0.0009 | 6 |
| Rpl10a | 0.2813 | 0.972 | 0.972 | 0.0403 | 8 |
| Ddx24 | 0.2812 | 0.9 | 0.94 | 1 | 9 |
| Hint1 | 0.2812 | 1 | 0.987 | 0.0003 | 7 |
| Rabac1 | 0.2812 | 0.99 | 0.958 | 2.82E-05 | 0 |
| Pex5l | 0.2811 | 0.451 | 0.176 | 0.125 | 8 |
| Spcs1 | 0.281 | 0.991 | 0.952 | 0.0009 | 4 |
| Kcnf1 | 0.2808 | 0.315 | 0.086 | 7.43E-07 | 4 |
| Tmem63c | 0.2807 | 0.353 | 0.056 | 1 | 10 |
| Erc2 | 0.2807 | 0.598 | 0.331 | 0.1064 | 5 |
| Spop | 0.2806 | 0.89 | 0.756 | 0.0121 | 2 |
| Epb41l4a | 0.2803 | 0.433 | 0.16 | 1 | 9 |
| Hras | 0.2803 | 0.967 | 0.924 | 1 | 9 |
| Ubald1 | 0.2802 | 0.667 | 0.396 | 1 | 9 |
| Oaz1 | 0.2802 | 1 | 0.998 | 3.49E-07 | 6 |
| Cadm2 | 0.2801 | 0.43 | 0.248 | 7.02E-06 | 0 |
| Nkx2-4 | 0.28 | 0.183 | 0.015 | 1.92E-05 | 8 |
| Slitrk2 | 0.2799 | 0.338 | 0.105 | 0.0361 | 8 |

| | | | | | |
|---|---|---|---|---|---|
| Hey1 | 0.2798 | 0.412 | 0.093 | 1 | 10 |
| Kitl | 0.2797 | 0.233 | 0.052 | 1 | 9 |
| Cdc37 | 0.2797 | 0.859 | 0.637 | 1 | 8 |
| Fam184b | 0.2795 | 0.25 | 0.054 | 2.44E-06 | 4 |
| Nrp2 | 0.2795 | 0.667 | 0.482 | 1 | 9 |
| Msi1 | 0.2794 | 0.452 | 0.146 | 7.29E-06 | 2 |
| Thrsp | 0.2794 | 0.267 | 0.028 | 1 | 9 |
| Kdsr | 0.2793 | 0.692 | 0.32 | 1.38E-05 | 2 |
| Cox6b1 | 0.2793 | 0.99 | 0.974 | 1.43E-10 | 6 |
| 5330417 | 0.2793 | 0.493 | 0.328 | 0.0026 | 1 |
| Rps25 | 0.2792 | 0.973 | 0.919 | 0.0025 | 2 |
| Kif1b | 0.2792 | 0.955 | 0.86 | 0.0001 | 0 |
| Lrrtm3 | 0.2792 | 0.76 | 0.465 | 6.43E-09 | 2 |
| Rpl23a | 0.2791 | 0.781 | 0.461 | 0.2078 | 2 |
| Pcdh19 | 0.2791 | 0.788 | 0.487 | 3.09E-05 | 2 |
| Hsd17b1 | 0.279 | 0.577 | 0.336 | 1 | 8 |
| Saraf | 0.279 | 0.972 | 0.93 | 0.0038 | 4 |
| Gpx1 | 0.2789 | 0.774 | 0.535 | 0.0003 | 2 |
| Rps4x | 0.2789 | 1 | 0.981 | 0.0122 | 7 |
| Fam150b | 0.2789 | 0.336 | 0.052 | 2.96E-13 | 2 |
| 5031439 | 0.2788 | 0.733 | 0.324 | 1 | 9 |
| Dnm1 | 0.2788 | 0.83 | 0.638 | 2.44E-05 | 0 |
| Cox8a | 0.2788 | 1 | 0.998 | 1 | 9 |
| Nsfl1c | 0.2788 | 0.923 | 0.719 | 0.0112 | 7 |
| Satb1 | 0.2787 | 0.384 | 0.143 | 7.59E-07 | 2 |
| Rpl22l1 | 0.2787 | 1 | 0.997 | 1 | 8 |
| Sema6a | 0.2786 | 0.535 | 0.186 | 0.002 | 8 |
| Ndufb5 | 0.2786 | 1 | 0.98 | 1 | 9 |
| Poldip3 | 0.2783 | 0.719 | 0.382 | 0.0024 | 2 |
| Malat1 | 0.2782 | 1 | 0.999 | 1.77E-06 | 3 |
| Pfdn5 | 0.278 | 1 | 0.96 | 0.0062 | 7 |
| Lsamp | 0.2779 | 0.915 | 0.828 | 0.0677 | 0 |
| Wbp11 | 0.2779 | 0.794 | 0.709 | 0.129 | 5 |
| Hspa5 | 0.2779 | 0.947 | 0.89 | 4.87E-06 | 1 |
| Rpl21 | 0.2778 | 1 | 0.997 | 1.60E-11 | 6 |
| Pxylp1 | 0.2778 | 0.394 | 0.158 | 0.0379 | 8 |
| Trim8 | 0.2777 | 0.623 | 0.284 | 3.23E-05 | 2 |
| Ppp1ca | 0.2777 | 0.915 | 0.822 | 0.4419 | 8 |
| Shisa9 | 0.2776 | 0.395 | 0.199 | 0.0002 | 1 |
| Smdt1 | 0.2775 | 0.972 | 0.94 | 0.297 | 8 |
| Chchd1 | 0.2773 | 0.907 | 0.716 | 1.20E-05 | 6 |
| Tenm3 | 0.2773 | 0.782 | 0.52 | 0.1046 | 7 |

| | | | | | |
|---|---|---|---|---|---|
| Lancl3 | 0.2772 | 0.059 | 0.022 | 1 | 10 |
| Alkbh5 | 0.2771 | 0.589 | 0.296 | 1 | 2 |
| Mrpl32 | 0.2771 | 0.733 | 0.468 | 1 | 9 |
| Dctn3 | 0.2771 | 0.944 | 0.892 | 1 | 8 |
| Mcmbp | 0.277 | 0.514 | 0.215 | 0.0005 | 2 |
| Grid2 | 0.277 | 0.467 | 0.23 | 1 | 9 |
| Lgals8 | 0.2769 | 0.563 | 0.302 | 0.6605 | 8 |
| Prdx1 | 0.2767 | 0.969 | 0.961 | 0.0053 | 5 |
| Mea1 | 0.2767 | 0.69 | 0.449 | 1 | 8 |
| Trp53i11 | 0.2766 | 0.885 | 0.712 | 0.0839 | 7 |
| Sart3 | 0.2766 | 0.533 | 0.24 | 1 | 9 |
| Stmn1 | 0.2766 | 1 | 0.999 | 0.0471 | 5 |
| Scg5 | 0.2765 | 0.98 | 0.977 | 1 | 0 |
| Pex5l | 0.2765 | 0.388 | 0.162 | 7.25E-05 | 1 |
| Ppp2r2b | 0.2765 | 0.718 | 0.482 | 1 | 8 |
| Glrx3 | 0.2763 | 0.753 | 0.587 | 0.0913 | 5 |
| Yeats4 | 0.2763 | 0.835 | 0.633 | 0.0014 | 5 |
| Cox7c | 0.2763 | 1 | 0.99 | 0.0003 | 7 |
| Dstn | 0.2763 | 0.959 | 0.883 | 0.1614 | 6 |
| Insm1 | 0.2762 | 0.309 | 0.117 | 0.025 | 5 |
| Uqcr11 | 0.2762 | 1 | 0.969 | 1 | 8 |
| Arhgap21 | 0.2762 | 0.519 | 0.272 | 0.0012 | 4 |
| Fis1 | 0.2761 | 1 | 0.927 | 7.00E-08 | 6 |
| Ppp2cb | 0.2761 | 0.836 | 0.627 | 0.5905 | 2 |
| Pde6d | 0.2761 | 0.719 | 0.377 | 0.0088 | 2 |
| Rgs7bp | 0.2761 | 0.4 | 0.121 | 1 | 9 |
| Serinc1 | 0.276 | 0.995 | 0.961 | 8.09E-05 | 0 |
| Atxn10 | 0.276 | 0.958 | 0.914 | 1 | 8 |
| Smdt1 | 0.276 | 1 | 0.936 | 5.39E-08 | 6 |
| Zcchc12 | 0.276 | 0.986 | 0.96 | 1 | 8 |
| Sepw1 | 0.2759 | 1 | 0.995 | 0.0009 | 7 |
| Gm2703 | 0.2759 | 0.335 | 0.146 | 1.30E-06 | 0 |
| Gpx1 | 0.2758 | 0.794 | 0.545 | 4.26E-05 | 6 |
| S100a1 | 0.2758 | 0.067 | 0.008 | 1 | 9 |
| Cnot8 | 0.2758 | 0.467 | 0.2 | 1 | 9 |
| Svip | 0.2758 | 0.664 | 0.361 | 0.1618 | 2 |
| Syt7 | 0.2758 | 0.355 | 0.145 | 1.90E-07 | 0 |
| Cyb5a | 0.2758 | 0.746 | 0.581 | 1 | 8 |
| Sdccag3 | 0.2757 | 0.447 | 0.394 | 1 | 3 |
| Chrm3 | 0.2757 | 0.315 | 0.099 | 5.67E-05 | 4 |
| RP24-23 | 0.2756 | 0.472 | 0.409 | 0.4467 | 3 |
| Igsf21 | 0.2756 | 0.352 | 0.149 | 0.0008 | 4 |

| | | | | | |
|---|---|---|---|---|---|
| Rnaset2a | 0.2755 | 0.244 | 0.153 | 1 | 7 |
| Tceb2 | 0.2755 | 1 | 0.992 | 6.52E-09 | 6 |
| Arpc1a | 0.2753 | 0.93 | 0.9 | 1 | 8 |
| Sdf2l1 | 0.2753 | 0.487 | 0.266 | 0.0002 | 1 |
| Dbpht2 | 0.2752 | 0.467 | 0.299 | 0.2488 | 1 |
| Arl6ip1 | 0.275 | 1 | 0.993 | 0.0003 | 4 |
| Ndufa7 | 0.275 | 1 | 0.949 | 7.98E-08 | 6 |
| Hibadh | 0.2749 | 0.651 | 0.302 | 0.0344 | 2 |
| Fez1 | 0.2747 | 0.911 | 0.818 | 0.0004 | 2 |
| Rpl18a | 0.2747 | 1 | 0.99 | 0.2703 | 8 |
| Myo10 | 0.2747 | 0.526 | 0.29 | 0.1231 | 5 |
| Snrpf | 0.2747 | 0.945 | 0.808 | 0.0003 | 2 |
| Park7 | 0.2746 | 1 | 0.941 | 0.0002 | 7 |
| Map1a | 0.2746 | 0.645 | 0.467 | 0.0007 | 0 |
| Pkia | 0.2746 | 0.774 | 0.54 | 1 | 2 |
| Gas7 | 0.2746 | 0.513 | 0.301 | 0.0327 | 1 |
| Atp5e | 0.2746 | 1 | 0.967 | 4.56E-14 | 6 |
| Ipo9 | 0.2745 | 0.533 | 0.264 | 1 | 9 |
| Hcrtr1 | 0.2745 | 0.267 | 0.029 | 0.3265 | 9 |
| Rpl8 | 0.2745 | 1 | 0.998 | 9.19E-08 | 6 |
| Glrb | 0.2744 | 0.475 | 0.325 | 0.0002 | 0 |
| Tef | 0.2742 | 0.836 | 0.614 | 0.0007 | 2 |
| Uqcr11 | 0.2742 | 1 | 0.97 | 1 | 9 |
| Tenm1 | 0.2741 | 0.733 | 0.374 | 1 | 9 |
| Apopt1 | 0.2741 | 0.692 | 0.365 | 0.002 | 2 |
| Smim11 | 0.2738 | 0.685 | 0.367 | 0.0033 | 2 |
| Dnm3 | 0.2737 | 0.82 | 0.699 | 0.0002 | 0 |
| Eno1 | 0.2737 | 0.803 | 0.714 | 1 | 8 |
| Tmem91 | 0.2735 | 0.855 | 0.719 | 0.0006 | 0 |
| Rps6 | 0.2734 | 1 | 0.934 | 1.67E-09 | 6 |
| Ndufv3 | 0.2733 | 0.99 | 0.933 | 0.0001 | 6 |
| Rab3c | 0.2733 | 0.979 | 0.863 | 1.61E-05 | 2 |
| Cox14 | 0.2733 | 0.928 | 0.708 | 1.06E-08 | 6 |
| Sod3 | 0.2732 | 0.154 | 0.005 | 3.23E-07 | 7 |
| Nudt18 | 0.2732 | 0.567 | 0.281 | 1 | 9 |
| Atxn7l3b | 0.2732 | 1 | 0.954 | 1 | 9 |
| Got2 | 0.273 | 0.745 | 0.583 | 5.93E-05 | 0 |
| Fgf13 | 0.273 | 0.521 | 0.237 | 1 | 2 |
| Otub1 | 0.2729 | 0.89 | 0.741 | 0.0034 | 2 |
| Adgrb1 | 0.2728 | 0.605 | 0.431 | 0.0052 | 0 |
| Lsm6 | 0.2728 | 0.897 | 0.735 | 0.2034 | 7 |
| Ldhb | 0.2727 | 0.99 | 0.891 | 0.0104 | 5 |

| | | | | | |
|---|---|---|---|---|---|
| Mcts1 | 0.2726 | 0.718 | 0.519 | 1 | 8 |
| Prdx2 | 0.2726 | 1 | 0.985 | 2.89E-05 | 5 |
| Sft2d1 | 0.2726 | 0.61 | 0.297 | 0.073 | 2 |
| Rorb | 0.2726 | 0.915 | 0.654 | 0.657 | 8 |
| Nav2 | 0.2725 | 0.472 | 0.224 | 0.0013 | 4 |
| l7Rn6 | 0.2725 | 0.774 | 0.543 | 0.0014 | 2 |
| Herc3 | 0.2724 | 0.633 | 0.331 | 1 | 9 |
| Ndfip2 | 0.2723 | 0.856 | 0.529 | 0.0143 | 2 |
| Ran | 0.2723 | 1 | 0.985 | 0.0001 | 5 |
| Nrip3 | 0.2722 | 0.282 | 0.17 | 0.5219 | 7 |
| Tmem20( | 0.2722 | 0.353 | 0.064 | 1 | 10 |
| Tmeff2 | 0.2722 | 0.41 | 0.209 | 1.01E-06 | 0 |
| Syn1 | 0.2722 | 0.77 | 0.58 | 8.55E-06 | 0 |
| Fstl5 | 0.2722 | 0.481 | 0.244 | 0.7257 | 4 |
| Ndrg2 | 0.272 | 0.437 | 0.155 | 0.4605 | 8 |
| Sdhaf4 | 0.2717 | 0.835 | 0.694 | 0.0422 | 5 |
| Park7 | 0.2716 | 0.979 | 0.942 | 3.14E-08 | 6 |
| Snrpd2 | 0.2716 | 0.944 | 0.9 | 1 | 8 |
| Ndufa8 | 0.2715 | 0.958 | 0.93 | 1 | 8 |
| Cox6a1 | 0.2715 | 0.9 | 0.725 | 1 | 9 |
| Rps15a | 0.2714 | 1 | 0.981 | 1 | 8 |
| Sgk1 | 0.2714 | 0.268 | 0.05 | 0.0004 | 8 |
| Cacna1e | 0.2714 | 0.675 | 0.493 | 0.0048 | 0 |
| Kif21a | 0.2713 | 0.87 | 0.784 | 0.8114 | 4 |
| Arhgap6 | 0.2713 | 0.887 | 0.579 | 0.3521 | 6 |
| Skil | 0.2713 | 0.548 | 0.252 | 0.5428 | 2 |
| Mir124a- | 0.2712 | 0.649 | 0.389 | 0.0286 | 5 |
| Lamtor2 | 0.2712 | 0.979 | 0.883 | 1.24E-07 | 6 |
| Etnk1 | 0.2711 | 0.943 | 0.784 | 1 | 3 |
| Vdac2 | 0.2711 | 0.967 | 0.939 | 1 | 9 |
| Usf1 | 0.2709 | 0.61 | 0.286 | 0.0839 | 2 |
| Rcan2 | 0.2709 | 0.5 | 0.433 | 0.0019 | 0 |
| Dnm3 | 0.2709 | 0.867 | 0.716 | 1 | 9 |
| Add3 | 0.2708 | 0.588 | 0.353 | 0.2187 | 5 |
| Hn1 | 0.2707 | 0.944 | 0.859 | 0.196 | 8 |
| Glrx5 | 0.2707 | 0.91 | 0.77 | 0.1546 | 7 |
| Pcdh20 | 0.2707 | 0.268 | 0.165 | 0.0008 | 3 |
| Llph | 0.2707 | 0.726 | 0.43 | 0.0056 | 2 |
| Mrps33 | 0.2706 | 0.969 | 0.906 | 0.0025 | 5 |
| Ncam2 | 0.2705 | 0.4 | 0.211 | 2.39E-05 | 0 |
| Dnpep | 0.2704 | 0.5 | 0.229 | 1 | 9 |
| Pde4b | 0.2704 | 0.505 | 0.257 | 1 | 5 |

| | | | | | |
|---|---|---|---|---|---|
| Arl3 | 0.2703 | 0.907 | 0.808 | 0.0024 | 6 |
| Sf3b6 | 0.2703 | 0.959 | 0.878 | 0.0018 | 5 |
| 9130221I | 0.2701 | 0.353 | 0.033 | 1 | 10 |
| Tmem263 | 0.2701 | 0.534 | 0.251 | 0.455 | 2 |
| Ppib | 0.2701 | 0.71 | 0.594 | 0.063 | 0 |
| Rpl21 | 0.2701 | 1 | 0.997 | 1 | 8 |
| Shfm1 | 0.27 | 0.993 | 0.934 | 1.34E-08 | 2 |
| Fndc9 | 0.27 | 0.408 | 0.194 | 0.0001 | 1 |
| Tmsb4x | 0.27 | 1 | 0.994 | 2.29E-10 | 6 |
| Fahd2a | 0.2699 | 0.507 | 0.219 | 0.0658 | 8 |
| Mllt11 | 0.2699 | 0.948 | 0.942 | 1 | 5 |
| Idi1 | 0.2697 | 0.555 | 0.267 | 1 | 2 |
| Dcaf17 | 0.2697 | 0.282 | 0.218 | 1 | 7 |
| Magel2 | 0.2697 | 0.712 | 0.392 | 8.68E-08 | 2 |
| Kdelr2 | 0.2697 | 0.671 | 0.301 | 0.0293 | 2 |
| Dst | 0.2697 | 0.743 | 0.623 | 0.005 | 1 |
| Taf10 | 0.2696 | 0.769 | 0.648 | 1 | 7 |
| Ap1s2 | 0.2696 | 0.822 | 0.579 | 1 | 2 |
| Mrpl34 | 0.2695 | 0.722 | 0.58 | 0.7829 | 5 |
| Rps3 | 0.2695 | 1 | 0.978 | 1.33E-07 | 2 |
| Meg3 | 0.2695 | 1 | 0.999 | 0.0014 | 5 |
| Mbnl2 | 0.2694 | 0.833 | 0.713 | 0.4434 | 4 |
| Rab3c | 0.2694 | 0.992 | 0.864 | 0.0246 | 3 |
| Trib2 | 0.2694 | 0.394 | 0.132 | 0.1131 | 8 |
| Btg2 | 0.2692 | 0.441 | 0.267 | 1 | 1 |
| Atp1a1 | 0.2692 | 0.743 | 0.573 | 0.0003 | 1 |
| Lrrtm1 | 0.2692 | 0.343 | 0.153 | 0.0145 | 4 |
| Snrpb | 0.2689 | 0.907 | 0.729 | 0.0015 | 6 |
| P3h3 | 0.2689 | 0.358 | 0.287 | 0.2093 | 3 |
| Fau | 0.2689 | 1 | 0.997 | 1.51E-08 | 2 |
| Spryd7 | 0.2688 | 0.644 | 0.327 | 1 | 2 |
| Hmgn5 | 0.2688 | 0.425 | 0.172 | 0.0141 | 2 |
| H1fx | 0.2687 | 0.866 | 0.719 | 0.4697 | 5 |
| Atp6v0e2 | 0.2687 | 0.96 | 0.919 | 0.0003 | 0 |
| Ccdc74a | 0.2686 | 0.521 | 0.228 | 0.1991 | 8 |
| Fam189a1 | 0.2686 | 0.515 | 0.226 | 0.0008 | 5 |
| Alcam | 0.2686 | 0.949 | 0.788 | 0.0437 | 7 |
| Rplp0 | 0.2685 | 0.972 | 0.932 | 0.894 | 8 |
| D8Ertd73 | 0.2684 | 0.936 | 0.875 | 0.7748 | 7 |
| Srp19 | 0.2682 | 0.933 | 0.721 | 1 | 9 |
| Rpl35 | 0.2681 | 1 | 0.984 | 2.11E-07 | 6 |
| Gria4 | 0.2681 | 0.585 | 0.445 | 0.0061 | 0 |

| | | | | | |
|---|---|---|---|---|---|
| Trpc5 | 0.2679 | 0.539 | 0.246 | 3.01E-05 | 1 |
| Ybx1 | 0.2678 | 0.842 | 0.702 | 0.0001 | 2 |
| Sox1 | 0.2678 | 0.603 | 0.353 | 0.0071 | 7 |
| Dap3 | 0.2677 | 0.718 | 0.515 | 1 | 8 |
| Nkx2-1 | 0.2677 | 0.257 | 0.057 | 4.52E-06 | 1 |
| Rnpc3 | 0.2676 | 0.447 | 0.382 | 1 | 3 |
| Sdhaf1 | 0.2676 | 0.596 | 0.338 | 0.0773 | 2 |
| Tmem13( | 0.2676 | 1 | 0.971 | 3.42E-06 | 1 |
| Pgrmc1 | 0.2676 | 1 | 0.999 | 3.37E-06 | 8 |
| Vamp2 | 0.2675 | 0.99 | 0.974 | 1 | 5 |
| Smarca5 | 0.2675 | 0.9 | 0.776 | 1 | 9 |
| AI854517 | 0.2675 | 0.367 | 0.041 | 0.0364 | 9 |
| Phb | 0.2675 | 0.867 | 0.643 | 1 | 9 |
| Sash1 | 0.2675 | 0.527 | 0.242 | 0.0599 | 2 |
| Pvrl1 | 0.2674 | 0.467 | 0.126 | 1 | 9 |
| Synj1 | 0.2674 | 0.81 | 0.622 | 6.29E-05 | 0 |
| Rps26 | 0.2674 | 0.986 | 0.933 | 8.30E-06 | 2 |
| Chtop | 0.2674 | 0.712 | 0.446 | 1 | 2 |
| Shisa5 | 0.2672 | 0.563 | 0.257 | 0.0865 | 8 |
| Bex1 | 0.2672 | 1 | 0.976 | 0.0153 | 7 |
| Lsm6 | 0.2672 | 0.863 | 0.729 | 0.0006 | 2 |
| Atp6v0e2 | 0.2671 | 1 | 0.926 | 1 | 10 |
| Uqcrc1 | 0.2671 | 0.867 | 0.739 | 1 | 9 |
| Tpt1 | 0.2671 | 1 | 0.987 | 0.0505 | 7 |
| Dmtn | 0.267 | 0.753 | 0.455 | 0.5119 | 2 |
| Mrpl54 | 0.267 | 0.821 | 0.672 | 1 | 7 |
| Celf5 | 0.2669 | 0.935 | 0.811 | 1 | 3 |
| Yme1l1 | 0.2669 | 0.585 | 0.544 | 1 | 3 |
| Chgb | 0.2669 | 0.93 | 0.837 | 0.025 | 0 |
| Rpl18a | 0.2669 | 1 | 0.989 | 0.0382 | 7 |
| Otud6b | 0.2668 | 0.678 | 0.338 | 0.0254 | 2 |
| Syt10 | 0.2668 | 0.904 | 0.606 | 3.01E-13 | 2 |
| Cnksr2 | 0.2668 | 0.596 | 0.302 | 0.0169 | 2 |
| Rps7 | 0.2668 | 1 | 0.978 | 9.30E-12 | 6 |
| 4930447( | 0.2668 | 0.485 | 0.244 | 0.0593 | 5 |
| Golga7b | 0.2667 | 0.606 | 0.431 | 1 | 8 |
| Tpm1 | 0.2667 | 0.619 | 0.539 | 1 | 6 |
| Hist1h2a| | 0.2667 | 0.346 | 0.116 | 0.007 | 7 |
| Gsta4 | 0.2666 | 0.361 | 0.157 | 1 | 5 |
| Nefl | 0.2664 | 0.435 | 0.222 | 0.7007 | 4 |
| Tra2b | 0.2664 | 0.679 | 0.512 | 0.7725 | 7 |
| Prmt2 | 0.2663 | 0.991 | 0.909 | 0.0001 | 4 |

| | | | | | |
|---|---|---|---|---|---|
| Bccip | 0.2663 | 0.596 | 0.319 | 1 | 2 |
| Nrcam | 0.2663 | 0.535 | 0.392 | 0.0141 | 0 |
| Mrpl20 | 0.2662 | 0.967 | 0.876 | 1 | 9 |
| Caln1 | 0.2662 | 0.355 | 0.121 | 2.70E-06 | 1 |
| Sorcs3 | 0.2662 | 0.388 | 0.16 | 0.0011 | 1 |
| Rpl23 | 0.2661 | 1 | 0.986 | 0.0047 | 7 |
| Rps9 | 0.2661 | 1 | 0.97 | 3.52E-10 | 2 |
| Map1b | 0.2661 | 0.987 | 0.977 | 0.0005 | 1 |
| Npy5r | 0.266 | 0.285 | 0.179 | 0.0002 | 3 |
| Hist3h2ba | 0.266 | 0.801 | 0.678 | 0.0147 | 2 |
| Kdsr | 0.266 | 0.577 | 0.348 | 0.8466 | 5 |
| Cd81 | 0.266 | 1 | 0.967 | 2.30E-08 | 1 |
| Sdf2 | 0.266 | 0.919 | 0.777 | 1 | 3 |
| Akap11 | 0.2659 | 0.89 | 0.755 | 0.0004 | 0 |
| Tspan4 | 0.2658 | 0.394 | 0.128 | 0.0184 | 8 |
| Zcchc10 | 0.2658 | 0.5 | 0.209 | 0.0039 | 2 |
| Tmbim6 | 0.2657 | 0.927 | 0.787 | 1 | 3 |
| Egln1 | 0.2655 | 0.644 | 0.318 | 0.0063 | 2 |
| Mad1l1 | 0.2655 | 0.353 | 0.076 | 1 | 10 |
| Nt5c | 0.2654 | 0.649 | 0.517 | 0.0037 | 6 |
| Gtf2a1 | 0.2654 | 0.432 | 0.153 | 0.004 | 2 |
| Rps11 | 0.2653 | 1 | 0.994 | 1.73E-07 | 6 |
| Vdac1 | 0.2652 | 0.973 | 0.946 | 0.005 | 2 |
| mt-Nd2 | 0.2652 | 1 | 0.999 | 1 | 10 |
| Gatad1 | 0.2651 | 0.651 | 0.376 | 1 | 2 |
| Slc1a3 | 0.265 | 0.282 | 0.074 | 0.0087 | 7 |
| Syne1 | 0.265 | 0.648 | 0.458 | 0.624 | 4 |
| Rps29 | 0.265 | 1 | 0.997 | 4.25E-15 | 2 |
| Pspc1 | 0.2649 | 0.567 | 0.297 | 1 | 9 |
| Rps27 | 0.2649 | 1 | 0.992 | 4.56E-11 | 6 |
| Gpc5 | 0.2647 | 0.23 | 0.049 | 9.61E-08 | 1 |
| Fkbp2 | 0.2647 | 0.967 | 0.923 | 6.00E-05 | 1 |
| Atp5g2 | 0.2647 | 1 | 0.97 | 0.001 | 7 |
| Hdgf | 0.2647 | 0.884 | 0.787 | 1.74E-05 | 2 |
| Creg2 | 0.2647 | 0.407 | 0.201 | 0.0129 | 4 |
| Lcorl | 0.2646 | 0.467 | 0.237 | 0.0001 | 1 |
| Acyp2 | 0.2646 | 0.603 | 0.318 | 0.5205 | 2 |
| Slc25a14 | 0.2646 | 0.77 | 0.611 | 0.0005 | 1 |
| Ppia | 0.2645 | 1 | 1 | 2.61E-19 | 6 |
| Hnrnpa3 | 0.2645 | 0.973 | 0.861 | 0.0002 | 2 |
| 2010107I | 0.2644 | 0.958 | 0.927 | 1 | 8 |
| Smek2 | 0.2642 | 0.493 | 0.234 | 0.9368 | 2 |

| | | | | | |
|---|---|---|---|---|---|
| C330007[illegible] | 0.2642 | 0.4 | 0.165 | 1 | 9 |
| Ube2ql1 | 0.2642 | 0.7 | 0.378 | 1 | 9 |
| Ostm1 | 0.2641 | 0.555 | 0.243 | 0.0011 | 2 |
| Snrpd2 | 0.264 | 0.99 | 0.894 | 1.86E-08 | 6 |
| Spock1 | 0.2639 | 0.485 | 0.381 | 1 | 0 |
| Rhoc | 0.2639 | 0.243 | 0.046 | 4.66E-08 | 1 |
| Scn5a | 0.2639 | 0.3 | 0.058 | 1 | 9 |
| Kifc2 | 0.2638 | 0.515 | 0.37 | 0.0054 | 0 |
| Slc25a4 | 0.2638 | 1 | 0.999 | 0.0067 | 7 |
| Tgfa | 0.2636 | 0.495 | 0.22 | 0.0116 | 5 |
| Kcnip4 | 0.2635 | 0.37 | 0.212 | 0.8876 | 4 |
| Elavl2 | 0.2635 | 0.42 | 0.295 | 0.0002 | 0 |
| Rab4a | 0.2633 | 0.718 | 0.51 | 1 | 8 |
| Slc9a6 | 0.2633 | 0.52 | 0.438 | 1 | 3 |
| Foxo1 | 0.2633 | 0.418 | 0.132 | 1.10E-05 | 2 |
| Acadm | 0.2632 | 0.479 | 0.232 | 0.4583 | 8 |
| Sub1 | 0.2632 | 1 | 0.987 | 0.0003 | 5 |
| Rpl8 | 0.2632 | 1 | 0.998 | 0.0899 | 8 |
| Rps21 | 0.2631 | 1 | 0.985 | 0.0176 | 7 |
| Trappc2 | 0.2631 | 0.603 | 0.294 | 1 | 2 |
| Fam19a5 | 0.2631 | 0.78 | 0.621 | 0.0002 | 0 |
| Zfp428 | 0.263 | 0.788 | 0.579 | 0.0036 | 2 |
| Rps8 | 0.2629 | 0.986 | 0.999 | 1 | 8 |
| Zfhx3 | 0.2629 | 0.949 | 0.715 | 0.0196 | 7 |
| Ddx24 | 0.2629 | 0.963 | 0.937 | 0.0049 | 4 |
| Abhd18 | 0.2629 | 0.596 | 0.312 | 0.0001 | 2 |
| Tub | 0.2628 | 0.612 | 0.449 | 0.0053 | 1 |
| Rab27b | 0.2627 | 0.692 | 0.353 | 0.6086 | 2 |
| Gpr50 | 0.2627 | 0.105 | 0.009 | 0.0029 | 1 |
| Pcgf5 | 0.2626 | 0.404 | 0.182 | 0.5757 | 2 |
| Rpl6 | 0.2626 | 1 | 0.998 | 8.66E-12 | 6 |
| Gria2 | 0.2626 | 0.98 | 0.949 | 0.0209 | 0 |
| Pcsk4 | 0.2625 | 0.355 | 0.208 | 0.0422 | 1 |
| Tasp1 | 0.2625 | 0.452 | 0.176 | 0.4354 | 2 |
| Eif1a | 0.2625 | 0.747 | 0.538 | 1 | 2 |
| Plpp2 | 0.2624 | 0.362 | 0.116 | 4.57E-08 | 1 |
| Mrps6 | 0.2624 | 0.76 | 0.422 | 0.0042 | 2 |
| Zfp428 | 0.2623 | 0.804 | 0.587 | 5.54E-05 | 6 |
| Tmem13[illegible] | 0.2623 | 0.4 | 0.256 | 0.0088 | 0 |
| Cisd1 | 0.2623 | 0.967 | 0.963 | 1 | 9 |
| Snx4 | 0.2623 | 0.555 | 0.253 | 0.2285 | 2 |
| Jmjd4 | 0.2623 | 0.294 | 0.041 | 1 | 10 |

| | | | | | |
|---|---|---|---|---|---|
| Wtap | 0.2622 | 0.644 | 0.402 | 1 | 2 |
| 4932438/ | 0.2622 | 0.713 | 0.522 | 0.0711 | 4 |
| Sult4a1 | 0.2621 | 0.92 | 0.811 | 5.56E-07 | 0 |
| Syt11 | 0.2621 | 0.991 | 0.937 | 0.0024 | 4 |
| Fam184b | 0.2619 | 0.296 | 0.058 | 0.0017 | 8 |
| Dlx5 | 0.2618 | 0.563 | 0.299 | 1 | 8 |
| Naa38 | 0.2618 | 0.814 | 0.594 | 4.03E-06 | 6 |
| Gsk3b | 0.2617 | 0.986 | 0.965 | 0.0001 | 2 |
| Bmyc | 0.2617 | 0.967 | 0.851 | 1 | 9 |
| Dut | 0.2617 | 0.733 | 0.459 | 0.0247 | 2 |
| Rpl4 | 0.2617 | 0.93 | 0.89 | 1 | 8 |
| Fam43a | 0.2617 | 0.258 | 0.059 | 0.0004 | 5 |
| Sf3b6 | 0.2617 | 0.979 | 0.871 | 4.48E-05 | 2 |
| mt-Nd3 | 0.2617 | 1 | 0.926 | 0.0753 | 9 |
| Bad | 0.2616 | 0.718 | 0.408 | 0.8412 | 8 |
| Kcnh1 | 0.2616 | 0.438 | 0.15 | 0.0005 | 2 |
| Vax1 | 0.2616 | 0.351 | 0.116 | 0.0015 | 5 |
| Nmral1 | 0.2615 | 0.366 | 0.139 | 0.189 | 8 |
| Fam173a | 0.2615 | 0.954 | 0.912 | 0.0384 | 4 |
| Pbx2 | 0.2614 | 0.575 | 0.281 | 0.009 | 2 |
| Irak2 | 0.2614 | 0.353 | 0.037 | 1 | 10 |
| Tpm3 | 0.2613 | 0.911 | 0.768 | 0.0003 | 2 |
| Cdkn1c | 0.2613 | 0.216 | 0.114 | 1 | 6 |
| Rnf5 | 0.2613 | 0.637 | 0.356 | 0.8057 | 2 |
| Atox1 | 0.2613 | 0.817 | 0.635 | 1 | 8 |
| Aff2 | 0.2612 | 0.4 | 0.104 | 1 | 9 |
| Ankrd54 | 0.2612 | 0.535 | 0.29 | 1 | 8 |
| Ociad1 | 0.2612 | 0.955 | 0.899 | 3.00E-05 | 0 |
| Fabp5 | 0.2612 | 0.944 | 0.856 | 0.0835 | 4 |
| Pcdh11x | 0.2611 | 0.285 | 0.149 | 0.0628 | 0 |
| Aplp2 | 0.2611 | 0.769 | 0.727 | 1 | 4 |
| Lamtor3 | 0.2611 | 0.616 | 0.314 | 0.2265 | 2 |
| Ccdc43 | 0.261 | 0.493 | 0.194 | 0.0065 | 2 |
| Ptp4a2 | 0.2609 | 0.784 | 0.619 | 0.0063 | 6 |
| Rpl35a | 0.2608 | 1 | 0.991 | 7.34E-10 | 2 |
| Inpp4a | 0.2607 | 0.829 | 0.593 | 1 | 2 |
| Fam174a | 0.2607 | 0.935 | 0.848 | 1 | 3 |
| Cpne2 | 0.2607 | 0.512 | 0.446 | 0.4336 | 3 |
| Sugt1 | 0.2607 | 0.746 | 0.446 | 1 | 8 |
| Ucp2 | 0.2607 | 0.537 | 0.308 | 0.2034 | 4 |
| Agpat4 | 0.2606 | 0.657 | 0.475 | 0.1917 | 4 |
| Rab4a | 0.2606 | 0.767 | 0.517 | 1 | 9 |

| | | | | | |
|---|---|---|---|---|---|
| Fundc1 | 0.2606 | 0.603 | 0.327 | 0.5487 | 2 |
| Nptxr | 0.2605 | 0.691 | 0.554 | 0.0089 | 1 |
| Nuak1 | 0.2605 | 0.3 | 0.068 | 1 | 9 |
| Macf1 | 0.2604 | 0.87 | 0.799 | 0.0678 | 4 |
| Sat1 | 0.2604 | 0.295 | 0.159 | 1 | 7 |
| Acsl1 | 0.2603 | 0.5 | 0.209 | 1 | 9 |
| Pfdn2 | 0.2603 | 1 | 0.97 | 0.014 | 7 |
| Tmtc4 | 0.2603 | 0.423 | 0.182 | 0.0086 | 7 |
| Thoc3 | 0.2602 | 0.7 | 0.379 | 1 | 9 |
| Ptchd1 | 0.2602 | 0.397 | 0.14 | 9.42E-05 | 2 |
| Cend1 | 0.2601 | 0.82 | 0.742 | 0.0003 | 0 |
| Tmem47 | 0.2601 | 0.538 | 0.358 | 1 | 7 |
| Dchs2 | 0.2601 | 0.309 | 0.078 | 0.0002 | 5 |
| Arx | 0.2601 | 0.356 | 0.101 | 0.0005 | 2 |
| Grid2 | 0.2601 | 0.42 | 0.196 | 4.13E-06 | 0 |
| Ptp4a2 | 0.2599 | 0.774 | 0.613 | 0.0172 | 2 |
| Hmgcs1 | 0.2599 | 0.659 | 0.598 | 1 | 3 |
| Tnr | 0.2598 | 0.454 | 0.229 | 0.0193 | 4 |
| Rps25 | 0.2598 | 0.99 | 0.92 | 3.39E-08 | 6 |
| Atraid | 0.2598 | 0.954 | 0.894 | 0.0102 | 4 |
| Traf6 | 0.2596 | 0.294 | 0.041 | 1 | 10 |
| Lingo3 | 0.2596 | 0.412 | 0.076 | 1 | 10 |
| Lhx1 | 0.2596 | 0.948 | 0.579 | 0.0077 | 6 |
| Zyg11b | 0.2594 | 0.5 | 0.363 | 0.001 | 0 |
| Fam103a1 | 0.2594 | 0.904 | 0.745 | 0.0777 | 2 |
| Pycr2 | 0.2592 | 0.637 | 0.311 | 0.069 | 2 |
| Fbxo9 | 0.2591 | 0.726 | 0.474 | 1 | 2 |
| 1700086[illegible] | 0.2591 | 0.52 | 0.349 | 1 | 1 |
| Plekhb2 | 0.259 | 0.845 | 0.766 | 1 | 5 |
| Adam23 | 0.259 | 0.447 | 0.34 | 0.0723 | 3 |
| 36324510[illegible] | 0.259 | 0.538 | 0.269 | 0.7769 | 7 |
| Anp32e | 0.2588 | 0.474 | 0.307 | 0.0443 | 7 |
| Lzts2 | 0.2588 | 0.367 | 0.118 | 1 | 9 |
| Il1rap | 0.2588 | 0.3 | 0.098 | 1 | 9 |
| Rbms3 | 0.2588 | 0.3 | 0.076 | 1 | 9 |
| Rpl22l1 | 0.2588 | 1 | 0.997 | 0.0188 | 7 |
| Zfp330 | 0.2587 | 0.514 | 0.216 | 0.0837 | 2 |
| Atp6v1f | 0.2587 | 1 | 0.969 | 1 | 8 |
| Rpl36 | 0.2587 | 1 | 0.992 | 5.10E-13 | 6 |
| Scamp5 | 0.2586 | 0.62 | 0.465 | 0.0122 | 0 |
| Cds2 | 0.2586 | 0.775 | 0.669 | 0.0942 | 0 |
| AK01087[illegible] | 0.2586 | 0.562 | 0.239 | 0.0295 | 2 |

| | | | | | |
|---|---|---|---|---|---|
| Capn10 | 0.2586 | 0.634 | 0.344 | 1 | 8 |
| Eif1 | 0.2586 | 1 | 0.998 | 2.01E-05 | 5 |
| Fbxw7 | 0.2586 | 0.54 | 0.353 | 3.56E-05 | 0 |
| Dnajc21 | 0.2585 | 0.559 | 0.339 | 0.0019 | 1 |
| Epha6 | 0.2585 | 0.322 | 0.066 | 1.00E-08 | 2 |
| Lamtor5 | 0.2585 | 0.856 | 0.595 | 2.22E-05 | 6 |
| Zcchc9 | 0.2585 | 0.549 | 0.338 | 1 | 8 |
| Tenm3 | 0.2585 | 0.753 | 0.506 | 2.90E-08 | 2 |
| Adck4 | 0.2585 | 0.382 | 0.319 | 1 | 3 |
| Txndc15 | 0.2584 | 0.917 | 0.853 | 0.0473 | 4 |
| Dst | 0.2584 | 0.787 | 0.623 | 0.1034 | 4 |
| Lyrm5 | 0.2584 | 0.412 | 0.19 | 0.0724 | 5 |
| Pnpla8 | 0.2583 | 0.658 | 0.396 | 1 | 2 |
| Vstm2l | 0.2583 | 0.966 | 0.826 | 2.00E-18 | 2 |
| Cacna2d2 | 0.2583 | 0.803 | 0.531 | 0.0118 | 8 |
| Kalrn | 0.2582 | 0.461 | 0.23 | 1.78E-05 | 1 |
| Ndufs4 | 0.2582 | 0.948 | 0.828 | 3.02E-05 | 6 |
| Cnnm1 | 0.2581 | 0.493 | 0.223 | 2.90E-06 | 1 |
| Tekt1 | 0.258 | 0.408 | 0.137 | 2.00E-06 | 1 |
| mt-Nd4l | 0.258 | 1 | 1 | 0.0075 | 4 |
| Ndufb9 | 0.2579 | 0.99 | 0.958 | 0.0208 | 5 |
| Ap1ar | 0.2579 | 0.658 | 0.346 | 1 | 2 |
| Dgki | 0.2579 | 0.485 | 0.221 | 0.0095 | 5 |
| Prdx2 | 0.2578 | 1 | 0.985 | 0.0002 | 6 |
| Ell2 | 0.2578 | 0.32 | 0.052 | 1.51E-07 | 5 |
| Slc25a39 | 0.2578 | 0.756 | 0.583 | 1 | 7 |
| Maf | 0.2577 | 0.217 | 0.031 | 3.49E-10 | 1 |
| Prr3 | 0.2575 | 0.651 | 0.34 | 0.1215 | 2 |
| Cops7a | 0.2575 | 0.726 | 0.43 | 1 | 2 |
| Zhx3 | 0.2574 | 0.423 | 0.218 | 0.3134 | 5 |
| Klhl13 | 0.2574 | 0.61 | 0.281 | 0.0194 | 2 |
| 3110040[illegible] | 0.2574 | 0.562 | 0.29 | 0.3575 | 2 |
| Sepw1 | 0.2573 | 1 | 0.995 | 1 | 8 |
| Rpl41 | 0.2573 | 1 | 1 | 0.0004 | 5 |
| Foxd2 | 0.2572 | 0.239 | 0.001 | 1.02E-14 | 8 |
| C1d | 0.2571 | 0.973 | 0.934 | 3.28E-06 | 2 |
| Hipk2 | 0.257 | 0.534 | 0.231 | 0.0003 | 2 |
| Bcap29 | 0.257 | 0.51 | 0.375 | 0.0037 | 0 |
| Gnpda2 | 0.257 | 0.526 | 0.337 | 1 | 5 |
| Fam178a | 0.257 | 0.589 | 0.323 | 0.8279 | 2 |
| 1810043[illegible] | 0.2569 | 0.8 | 0.456 | 1 | 9 |
| Npy5r | 0.2569 | 0.459 | 0.15 | 6.53E-06 | 2 |

| | | | | | |
|---|---|---|---|---|---|
| Sox11 | 0.2568 | 0.648 | 0.479 | 1 | 8 |
| Ociad2 | 0.2568 | 0.454 | 0.269 | 0.0659 | 4 |
| Uhrf2 | 0.2568 | 0.487 | 0.395 | 1 | 7 |
| Kmt2d | 0.2567 | 0.541 | 0.234 | 0.0665 | 2 |
| Slc24a3 | 0.2567 | 0.5 | 0.372 | 0.2457 | 0 |
| Rps24 | 0.2567 | 1 | 0.997 | 3.09E-12 | 2 |
| Akr1a1 | 0.2566 | 1 | 0.932 | 6.12E-08 | 6 |
| Rrad | 0.2565 | 0.282 | 0.161 | 1 | 7 |
| Gm9780 | 0.2565 | 0.218 | 0.026 | 2.57E-05 | 7 |
| Luzp1 | 0.2564 | 0.433 | 0.133 | 1 | 9 |
| Xpa | 0.2564 | 0.567 | 0.335 | 0.1888 | 5 |
| Pdcd5 | 0.2563 | 0.915 | 0.893 | 1 | 8 |
| Pcgf2 | 0.2562 | 0.5 | 0.238 | 0.0339 | 2 |
| Btbd1 | 0.2562 | 0.705 | 0.376 | 1 | 2 |
| Fam57a | 0.2561 | 0.269 | 0.084 | 0.0208 | 7 |
| Stxbp1 | 0.256 | 0.97 | 0.912 | 0.001 | 0 |
| Pcbp1 | 0.256 | 0.859 | 0.823 | 1 | 7 |
| Acyp2 | 0.2559 | 0.538 | 0.341 | 0.0192 | 7 |
| Lsm3 | 0.2558 | 0.705 | 0.387 | 0.0032 | 2 |
| Pepd | 0.2556 | 0.541 | 0.307 | 0.011 | 2 |
| Sema5a | 0.2556 | 0.324 | 0.156 | 0.6211 | 4 |
| Isca2 | 0.2556 | 0.76 | 0.526 | 0.0855 | 2 |
| Nono | 0.2555 | 0.897 | 0.675 | 4.87E-08 | 2 |
| Tox | 0.2555 | 0.536 | 0.307 | 1 | 5 |
| Nos1ap | 0.2555 | 0.566 | 0.341 | 0.0035 | 1 |
| Vta1 | 0.2555 | 0.637 | 0.359 | 0.2685 | 2 |
| Tmem55a | 0.2554 | 0.486 | 0.219 | 1 | 2 |
| Ltbp3 | 0.2553 | 0.441 | 0.212 | 7.67E-05 | 1 |
| Sepw1 | 0.2553 | 0.99 | 0.996 | 0.0119 | 5 |
| Mrpl42 | 0.2552 | 0.704 | 0.559 | 1 | 8 |
| Irf2bp2 | 0.2552 | 0.449 | 0.297 | 0.5 | 7 |
| Ndrg4 | 0.2552 | 0.918 | 0.883 | 1 | 5 |
| Rnf7 | 0.2551 | 0.969 | 0.89 | 0.0431 | 5 |
| Lsm4 | 0.2549 | 0.897 | 0.742 | 0.0011 | 5 |
| Slc35f4 | 0.2548 | 0.296 | 0.069 | 0.0041 | 8 |
| Grhpr | 0.2548 | 0.633 | 0.264 | 1 | 9 |
| Prkacb | 0.2548 | 0.967 | 0.949 | 1 | 9 |
| Gstz1 | 0.2548 | 0.549 | 0.3 | 1 | 8 |
| Polr2l | 0.2547 | 0.784 | 0.606 | 0.5339 | 5 |
| Scn1a | 0.2547 | 0.633 | 0.272 | 1 | 9 |
| Rrp7a | 0.2547 | 0.753 | 0.596 | 0.0119 | 6 |
| Nt5c | 0.2546 | 0.74 | 0.496 | 0.0071 | 2 |

| | | | | | |
|---|---|---|---|---|---|
| Atp5j2 | 0.2546 | 1 | 0.99 | 1 | 8 |
| Car10 | 0.2545 | 0.493 | 0.283 | 1 | 2 |
| Adgrl1 | 0.2544 | 0.796 | 0.662 | 0.0003 | 1 |
| Mrps18c | 0.2543 | 0.592 | 0.41 | 1 | 8 |
| Plcxd3 | 0.2541 | 0.553 | 0.39 | 0.6909 | 1 |
| Nrn1l | 0.254 | 0.179 | 0.04 | 0.0009 | 7 |
| Atp6v0e2 | 0.254 | 1 | 0.925 | 1 | 9 |
| Wfs1 | 0.2539 | 0.592 | 0.299 | 1 | 8 |
| Rpl27a | 0.2539 | 1 | 0.994 | 1 | 8 |
| Kansl2 | 0.2538 | 0.589 | 0.289 | 0.585 | 2 |
| Sgtb | 0.2537 | 0.485 | 0.222 | 0.0455 | 5 |
| Sv2c | 0.2537 | 0.268 | 0.104 | 1 | 8 |
| Rpl18 | 0.2537 | 1 | 0.981 | 0.1025 | 8 |
| D030056 | 0.2536 | 0.548 | 0.217 | 0.2115 | 2 |
| Cox7b | 0.2536 | 0.959 | 0.911 | 0.2324 | 5 |
| Kcnk2 | 0.2536 | 0.586 | 0.326 | 0.0065 | 1 |
| Rpl35 | 0.2536 | 1 | 0.985 | 0.0058 | 7 |
| Ajap1 | 0.2535 | 0.74 | 0.393 | 0.0059 | 2 |
| N4bp2l1 | 0.2534 | 0.74 | 0.414 | 0.0079 | 2 |
| Lgi3 | 0.2534 | 0.36 | 0.168 | 8.31E-07 | 0 |
| Celf3 | 0.2534 | 0.808 | 0.721 | 1 | 7 |
| H3f3a | 0.2533 | 1 | 0.948 | 0.0065 | 7 |
| Hn1 | 0.2533 | 0.959 | 0.855 | 0.0016 | 6 |
| Shd | 0.2532 | 0.527 | 0.244 | 0.0346 | 2 |
| Sox4 | 0.2532 | 0.536 | 0.312 | 1 | 5 |
| Rps13 | 0.2532 | 1 | 0.977 | 1.28E-09 | 6 |
| Prdx2 | 0.2532 | 1 | 0.986 | 0.5599 | 8 |
| Vgll4 | 0.2531 | 0.296 | 0.08 | 0.0021 | 8 |
| 1810043 | 0.253 | 0.718 | 0.448 | 1 | 8 |
| Rwdd3 | 0.253 | 0.4 | 0.1 | 1 | 9 |
| Rln1 | 0.2528 | 0.244 | 0.057 | 0.0003 | 7 |
| Chl1 | 0.2526 | 0.655 | 0.436 | 0.0556 | 0 |
| Ubtf | 0.2525 | 0.61 | 0.31 | 0.8083 | 2 |
| Ssbp1 | 0.2525 | 0.722 | 0.462 | 0.6849 | 5 |
| Gm3626 | 0.2525 | 0.179 | 0.09 | 1 | 3 |
| Brinp1 | 0.2524 | 0.659 | 0.6 | 1 | 3 |
| Tubb5 | 0.2524 | 0.993 | 0.975 | 8.27E-05 | 2 |
| Smim10l | 0.2523 | 0.918 | 0.868 | 0.5429 | 5 |
| Rps27 | 0.2523 | 1 | 0.992 | 7.82E-09 | 2 |
| Efhd2 | 0.2523 | 0.553 | 0.329 | 0.0039 | 1 |
| Phyhip | 0.2522 | 0.37 | 0.229 | 0.0004 | 0 |
| Gigyf1 | 0.2521 | 0.63 | 0.325 | 0.1689 | 2 |

| | | | | | |
|---|---|---|---|---|---|
| Btf3l4 | 0.252 | 0.808 | 0.587 | 0.6274 | 2 |
| Fkbp3 | 0.252 | 1 | 0.966 | 1.76E-08 | 6 |
| Rpl10 | 0.252 | 0.986 | 0.932 | 0.0004 | 2 |
| Cdkn2d | 0.252 | 0.742 | 0.554 | 1 | 5 |
| Cd200 | 0.2519 | 0.907 | 0.864 | 0.2637 | 4 |
| Cap1 | 0.2519 | 0.473 | 0.197 | 0.1557 | 2 |
| Ptpre | 0.2518 | 0.493 | 0.226 | 0.2779 | 8 |
| Usp51 | 0.2516 | 0.31 | 0.134 | 1 | 8 |
| Gstp1 | 0.2516 | 0.952 | 0.916 | 0.009 | 2 |
| Cdk14 | 0.2516 | 0.405 | 0.288 | 0.0409 | 0 |
| Eif5a | 0.2515 | 1 | 0.988 | 0.0007 | 7 |
| Rpl38 | 0.2515 | 1 | 0.997 | 2.22E-08 | 6 |
| N6amt2 | 0.2515 | 0.648 | 0.37 | 1 | 8 |
| Gas7 | 0.2514 | 0.634 | 0.309 | 1 | 8 |
| Ydjc | 0.2514 | 0.513 | 0.336 | 0.0151 | 1 |
| Arxes1 | 0.2514 | 0.81 | 0.655 | 0.0023 | 0 |
| H2afv | 0.2514 | 0.856 | 0.73 | 0.0465 | 5 |
| Pomp | 0.2514 | 0.944 | 0.904 | 1 | 8 |
| Bex1 | 0.2513 | 1 | 0.976 | 1.51E-08 | 6 |
| Wasf3 | 0.2512 | 0.533 | 0.251 | 1 | 9 |
| Stx6 | 0.2512 | 0.596 | 0.323 | 0.4525 | 2 |
| Camk1d | 0.2512 | 0.425 | 0.136 | 0.001 | 2 |
| G0s2 | 0.2512 | 0.315 | 0.095 | 0.0001 | 4 |
| Dfna5 | 0.2511 | 0.562 | 0.254 | 0.0463 | 2 |
| Flywch2 | 0.251 | 0.608 | 0.373 | 1 | 5 |
| Nmnat2 | 0.251 | 0.606 | 0.419 | 1 | 8 |
| Mrpl17 | 0.251 | 0.856 | 0.632 | 0.0036 | 2 |
| Ndufa11 | 0.251 | 0.99 | 0.973 | 0.0007 | 5 |
| Mrps18b | 0.2509 | 0.557 | 0.308 | 0.1934 | 5 |
| Rps23 | 0.2509 | 1 | 0.969 | 0.0239 | 7 |
| Gpld1 | 0.2509 | 0.421 | 0.241 | 0.0048 | 1 |
| Glod4 | 0.2508 | 0.66 | 0.399 | 0.2266 | 5 |
| Sptssa | 0.2507 | 0.747 | 0.455 | 0.0288 | 2 |
| Atxn7l3b | 0.2506 | 1 | 0.951 | 0.0111 | 6 |
| Ppp3ca | 0.2505 | 0.895 | 0.771 | 0.0099 | 0 |
| Alyref | 0.2503 | 0.671 | 0.417 | 1 | 2 |
| Ociad2 | 0.2503 | 0.521 | 0.271 | 1 | 8 |
| Col25a1 | 0.2503 | 0.29 | 0.181 | 0.0545 | 0 |
| Rpl6 | 0.2501 | 1 | 0.998 | 1 | 8 |
| Rps26 | 0.2501 | 0.987 | 0.937 | 0.041 | 7 |
| Thra | 0.2501 | 0.99 | 0.986 | 0.0225 | 6 |
| Atp5g1 | 0.25 | 1 | 0.988 | 1.78E-08 | 6 |

| | | | | | |
|---|---|---|---|---|---|
| Taf10 | 0.25 | 0.849 | 0.628 | 0.0293 | 2 |
| Hmgcs1 | 0.25 | 0.712 | 0.589 | 1 | 2 |
| Lin7b | -0.25 | 0.192 | 0.362 | 1 | 7 |
| Trpc4 | -0.25 | 0.013 | 0.209 | 1.19E-09 | 1 |
| Slc25a12 | -0.25 | 0.165 | 0.401 | 1 | 6 |
| AI48052€ | -0.25 | 0.183 | 0.352 | 1 | 8 |
| Scaf11 | -0.25 | 0.176 | 0.38 | 1 | 10 |
| Kcnip4 | -0.2501 | 0.081 | 0.245 | 1 | 3 |
| Cops4 | -0.2501 | 0.39 | 0.634 | 1 | 3 |
| Sumo2 | -0.2502 | 0.985 | 0.995 | 0.0005 | 0 |
| Tmed4 | -0.2503 | 0.691 | 0.779 | 0.096 | 6 |
| Vps72 | -0.2503 | 0.118 | 0.337 | 1 | 10 |
| Ghitm | -0.2504 | 0.467 | 0.666 | 0.0026 | 1 |
| Kif21a | -0.2504 | 0.679 | 0.801 | 1 | 7 |
| Gm3626€ | -0.2505 | 0 | 0.103 | 1 | 9 |
| Th | -0.2505 | 0.021 | 0.117 | 0.7414 | 5 |
| Prkag2 | -0.2505 | 0.065 | 0.315 | 1 | 3 |
| Trnp1 | -0.2505 | 0.049 | 0.296 | 1 | 3 |
| Prkca | -0.2506 | 0.216 | 0.349 | 1 | 5 |
| Atraid | -0.2506 | 0.846 | 0.904 | 1 | 7 |
| Plxna4 | -0.2507 | 0.026 | 0.229 | 0.0486 | 7 |
| Ppp3r1 | -0.2507 | 0.538 | 0.707 | 1 | 7 |
| 6330403, | -0.2509 | 0 | 0.181 | 0.0003 | 3 |
| Pdcd4 | -0.2509 | 0.479 | 0.686 | 1 | 8 |
| Cib2 | -0.2511 | 0.577 | 0.733 | 0.4735 | 5 |
| Paip1 | -0.2511 | 0.295 | 0.464 | 0.0156 | 0 |
| Bloc1s5 | -0.2512 | 0.059 | 0.28 | 1 | 10 |
| Slc18a2 | -0.2512 | 0 | 0.133 | 1 | 9 |
| Fut9 | -0.2512 | 0.175 | 0.372 | 1 | 6 |
| Srebf2 | -0.2513 | 0.211 | 0.38 | 1 | 8 |
| Elmod1 | -0.2514 | 0.648 | 0.727 | 1 | 8 |
| Npy5r | -0.2514 | 0.019 | 0.209 | 3.03E-05 | 4 |
| Shisa5 | -0.2515 | 0.103 | 0.289 | 0.9877 | 7 |
| Wrb | -0.2515 | 0.564 | 0.697 | 1 | 7 |
| Prpsap1 | -0.2515 | 0.176 | 0.434 | 1 | 10 |
| Brd9 | -0.2515 | 0.235 | 0.446 | 1 | 10 |
| Gstm5 | -0.2516 | 0.615 | 0.778 | 0.396 | 7 |
| Kifap3 | -0.2516 | 0.795 | 0.914 | 0.2892 | 2 |
| Sdf2 | -0.2517 | 0.718 | 0.798 | 1 | 7 |
| Tmie | -0.2517 | 0.165 | 0.341 | 0.0007 | 5 |
| Nfix | -0.2518 | 0.016 | 0.112 | 1 | 3 |
| Rnpc3 | -0.2518 | 0.256 | 0.399 | 1 | 7 |

| | | | | | |
|---|---|---|---|---|---|
| Cntfr | -0.252 | 0.155 | 0.359 | 0.0125 | 5 |
| Kcnip1 | -0.2521 | 0 | 0.189 | 2.95E-08 | 5 |
| Tmed2 | -0.2521 | 0.887 | 0.871 | 1 | 6 |
| Dlx2 | -0.2521 | 0 | 0.213 | 1 | 10 |
| Unc13c | -0.2522 | 0 | 0.096 | 0.0766 | 4 |
| B4gat1 | -0.2522 | 0.603 | 0.682 | 1 | 7 |
| Sema5a | -0.2522 | 0.052 | 0.184 | 0.0261 | 5 |
| Ppp2ca | -0.2523 | 0.943 | 0.942 | 1 | 3 |
| Myt1l | -0.2523 | 0.526 | 0.706 | 1 | 7 |
| Timm10 | -0.2523 | 0.176 | 0.41 | 1 | 10 |
| Gabra3 | -0.2524 | 0.212 | 0.345 | 5.92E-07 | 2 |
| Praf2 | -0.2524 | 0.487 | 0.662 | 0.009 | 1 |
| Smim11 | -0.2525 | 0.23 | 0.447 | 8.18E-05 | 0 |
| Leo1 | -0.2525 | 0.059 | 0.278 | 1 | 10 |
| Gm21092 | -0.2526 | 0.5 | 0.646 | 1 | 7 |
| Cltb | -0.2526 | 0.74 | 0.848 | 0.9079 | 3 |
| Impad1 | -0.2527 | 0.333 | 0.561 | 1 | 7 |
| Dync2h1 | -0.2527 | 0.256 | 0.427 | 1 | 7 |
| Gabre | -0.2528 | 0 | 0.157 | 1 | 10 |
| Nr2f2 | -0.2528 | 0.325 | 0.442 | 1 | 0 |
| Nav2 | -0.253 | 0.065 | 0.27 | 0.1814 | 3 |
| 1-Mar | -0.253 | 0.282 | 0.438 | 1 | 8 |
| Rps13 | -0.253 | 0.96 | 0.983 | 0.0083 | 0 |
| Car11 | -0.2531 | 0.169 | 0.341 | 1 | 8 |
| Klhl13 | -0.2531 | 0.155 | 0.335 | 0.3182 | 8 |
| Gabrb3 | -0.2532 | 0.575 | 0.692 | 0.0408 | 2 |
| Tmem245 | -0.2532 | 0.179 | 0.358 | 1 | 7 |
| Tmem9 | -0.2532 | 0.568 | 0.709 | 1 | 2 |
| Plcl1 | -0.2533 | 0.125 | 0.33 | 4.71E-05 | 0 |
| Adgrb3 | -0.2533 | 0.346 | 0.49 | 1 | 7 |
| Zcchc17 | -0.2534 | 0.732 | 0.817 | 1 | 3 |
| Snx21 | -0.2535 | 0.176 | 0.397 | 1 | 10 |
| Prdx6 | -0.2535 | 0.415 | 0.651 | 1 | 3 |
| Orc6 | -0.2535 | 0.118 | 0.34 | 1 | 10 |
| Gpr101 | -0.2536 | 0 | 0.122 | 0.0042 | 5 |
| Rxrg | -0.2537 | 0.041 | 0.157 | 1 | 6 |
| Anp32b | -0.2537 | 0.57 | 0.728 | 0.0022 | 0 |
| 6430548 | -0.2538 | 0.228 | 0.462 | 1 | 3 |
| Slc24a3 | -0.2539 | 0.278 | 0.406 | 1 | 5 |
| Ptges3 | -0.254 | 0.8 | 0.924 | 1 | 9 |
| B3galt1 | -0.254 | 0.083 | 0.279 | 0.0011 | 4 |
| Stt3b | -0.254 | 0.062 | 0.305 | 0.1436 | 6 |

| | | | | | |
|---|---|---|---|---|---|
| Dgki | -0.2542 | 0.056 | 0.264 | 8.84E-05 | 4 |
| Pbx4 | -0.2543 | 0.089 | 0.316 | 1 | 3 |
| Zfp428 | -0.2544 | 0.54 | 0.62 | 1 | 0 |
| Ttll5 | -0.2545 | 0.099 | 0.263 | 1 | 8 |
| Tecr | -0.2545 | 1 | 0.993 | 1 | 7 |
| Frmd4a | -0.2545 | 0.167 | 0.392 | 1 | 7 |
| Rpl21 | -0.2546 | 1 | 0.997 | 0.0379 | 0 |
| Pfdn5 | -0.2547 | 0.925 | 0.971 | 0.0129 | 0 |
| Vezt | -0.2548 | 0.244 | 0.439 | 1 | 7 |
| Gfra2 | -0.2549 | 0.256 | 0.394 | 1 | 7 |
| Cep83os | -0.2549 | 0 | 0.235 | 1 | 10 |
| Atp6v1h | -0.255 | 0.333 | 0.543 | 1 | 9 |
| Pithd1 | -0.2551 | 0.385 | 0.597 | 1 | 7 |
| Pcdh7 | -0.2551 | 0.351 | 0.489 | 1 | 5 |
| Coq2 | -0.2551 | 0.367 | 0.566 | 1 | 9 |
| Shisa5 | -0.2551 | 0.093 | 0.294 | 0.0004 | 5 |
| Shisa5 | -0.2551 | 0.103 | 0.293 | 1 | 6 |
| Kcna2 | -0.2552 | 0.07 | 0.276 | 0.1823 | 8 |
| Fkbp2 | -0.2552 | 0.928 | 0.93 | 0.09 | 6 |
| Pdia4 | -0.2553 | 0.186 | 0.402 | 1 | 6 |
| Scn1b | -0.2553 | 0.598 | 0.673 | 1 | 6 |
| Cspp1 | -0.2554 | 0.346 | 0.525 | 1 | 7 |
| Tcf4 | -0.2554 | 0.253 | 0.312 | 0.017 | 2 |
| Timm10b | -0.2554 | 0.39 | 0.659 | 1 | 3 |
| Tm9sf3 | -0.2555 | 0.505 | 0.624 | 0.5618 | 6 |
| 22100130 | -0.2555 | 0.684 | 0.834 | 0.0002 | 1 |
| Shisa9 | -0.2556 | 0.033 | 0.23 | 1 | 9 |
| RP23-407 | -0.2556 | 0.353 | 0.552 | 1 | 10 |
| Fabp5 | -0.2557 | 0.744 | 0.873 | 1 | 7 |
| Cystm1 | -0.2557 | 0.615 | 0.718 | 0.2566 | 0 |
| Adgra1 | -0.2558 | 0.231 | 0.425 | 1 | 7 |
| 1810041 | -0.2558 | 0.059 | 0.244 | 0.8141 | 10 |
| Gabra4 | -0.2559 | 0.026 | 0.23 | 0.0193 | 7 |
| Zcchc12 | -0.2559 | 0.962 | 0.962 | 1 | 7 |
| Casc4 | -0.2559 | 0.577 | 0.73 | 1 | 7 |
| Lrrc4c | -0.2561 | 0.185 | 0.412 | 0.0167 | 4 |
| Eif1 | -0.2561 | 0.995 | 0.999 | 0.0083 | 0 |
| Rab3a | -0.2561 | 1 | 0.999 | 0.1109 | 3 |
| Tmem87b | -0.2561 | 0.093 | 0.318 | 0.804 | 6 |
| Wfs1 | -0.2561 | 0.175 | 0.331 | 1 | 6 |
| Limd2 | -0.2562 | 0.118 | 0.353 | 1 | 10 |
| Ndufs4 | -0.2563 | 0.748 | 0.849 | 1 | 3 |

| | | | | | |
|---|---|---|---|---|---|
| Rgs17 | -0.2565 | 0.637 | 0.699 | 0.0385 | 2 |
| Ahi1 | -0.2565 | 1 | 1 | 1 | 7 |
| Srsf2 | -0.2566 | 0.873 | 0.935 | 1 | 8 |
| Sltm | -0.2568 | 0.22 | 0.521 | 1 | 3 |
| Tmem176 | -0.2568 | 0.224 | 0.418 | 0.0128 | 1 |
| Uqcr10 | -0.2569 | 0.908 | 0.97 | 1.55E-05 | 1 |
| Mycbp2 | -0.257 | 0.846 | 0.895 | 1 | 7 |
| Lrp11 | -0.257 | 0.34 | 0.495 | 1 | 6 |
| Bub3 | -0.2571 | 0.567 | 0.741 | 1 | 5 |
| Adam11 | -0.2571 | 0.038 | 0.233 | 0.139 | 7 |
| Slc22a17 | -0.2571 | 0.987 | 0.964 | 1 | 7 |
| Oxr1 | -0.2572 | 0.165 | 0.39 | 1 | 6 |
| Lynx1 | -0.2572 | 0.021 | 0.181 | 1.40E-05 | 5 |
| Hpca | -0.2572 | 0.128 | 0.316 | 1 | 7 |
| Eml2 | -0.2572 | 0.321 | 0.521 | 1 | 7 |
| Ttll5 | -0.2572 | 0.105 | 0.285 | 0.0001 | 0 |
| Stip1 | -0.2573 | 0.496 | 0.709 | 1 | 3 |
| Oxct1 | -0.2573 | 0.538 | 0.672 | 1 | 7 |
| Gad1 | -0.2575 | 0.732 | 0.762 | 1 | 6 |
| Lgi3 | -0.2575 | 0.013 | 0.216 | 0.0026 | 7 |
| Pkig | -0.2575 | 0.415 | 0.656 | 1 | 3 |
| Opcml | -0.2575 | 0.61 | 0.685 | 1.54E-05 | 3 |
| Slc38a2 | -0.2576 | 0.186 | 0.411 | 1 | 6 |
| Nr3c1 | -0.2576 | 0.113 | 0.302 | 1 | 6 |
| Itfg1 | -0.2577 | 0.546 | 0.647 | 0.6059 | 6 |
| Vat1 | -0.2577 | 0.856 | 0.9 | 1 | 5 |
| Tpd52l1 | -0.2578 | 0.394 | 0.475 | 1 | 8 |
| Rpl7a | -0.2578 | 0.8 | 0.918 | 0.0004 | 0 |
| Slc30a9 | -0.2579 | 0.529 | 0.639 | 1 | 10 |
| Zfand6 | -0.2579 | 0.3 | 0.481 | 0.0056 | 0 |
| Mrpl50 | -0.258 | 0.176 | 0.372 | 1 | 10 |
| Serinc3 | -0.258 | 0.183 | 0.378 | 0.4454 | 8 |
| Pcmtd1 | -0.2581 | 0.254 | 0.498 | 0.8096 | 8 |
| Tmem91 | -0.2582 | 0.634 | 0.751 | 1 | 8 |
| Gclm | -0.2582 | 0.114 | 0.381 | 1 | 3 |
| Celf4 | -0.2582 | 0.897 | 0.984 | 0.6458 | 7 |
| Nt5dc3 | -0.2582 | 0.194 | 0.349 | 0.3675 | 4 |
| Hbegf | -0.2583 | 0.13 | 0.224 | 1 | 4 |
| Marf1 | -0.2583 | 0.3 | 0.592 | 1 | 9 |
| Usp9x | -0.2585 | 0.722 | 0.819 | 0.0042 | 6 |
| Sec11a | -0.2585 | 0.462 | 0.584 | 1 | 7 |
| Trim32 | -0.2586 | 0.205 | 0.378 | 1 | 7 |

| | | | | | |
|---|---|---|---|---|---|
| Reep2 | -0.2587 | 0.582 | 0.706 | 1 | 2 |
| Srp54b | -0.2588 | 0.561 | 0.728 | 1 | 3 |
| Nedd4l | -0.2589 | 0.077 | 0.305 | 0.1803 | 7 |
| Cend1 | -0.2589 | 0.618 | 0.778 | 0.0011 | 1 |
| Plcl1 | -0.259 | 0.1 | 0.298 | 1 | 9 |
| Epb41l4b | -0.259 | 0.052 | 0.25 | 1.00E-05 | 5 |
| Cdkn1c | -0.259 | 0.014 | 0.131 | 1 | 8 |
| Atp5h | -0.2591 | 0.954 | 0.989 | 8.20E-08 | 1 |
| Sap30l | -0.2591 | 0.081 | 0.338 | 1 | 3 |
| Zbtb20 | -0.2591 | 0.639 | 0.749 | 0.0135 | 6 |
| Gm36266 | -0.2591 | 0 | 0.107 | 0.6833 | 8 |
| Gabre | -0.2592 | 0.01 | 0.168 | 0.0008 | 5 |
| Nfix | -0.2592 | 0 | 0.11 | 0.7178 | 7 |
| Ildr2 | -0.2593 | 0.124 | 0.339 | 1 | 6 |
| Gabbr2 | -0.2594 | 0.206 | 0.405 | 1 | 6 |
| Gpr101 | -0.2594 | 0 | 0.126 | 0.0001 | 3 |
| Tmx1 | -0.2594 | 0.346 | 0.524 | 1 | 7 |
| Crim1 | -0.2595 | 0.028 | 0.22 | 0.0142 | 8 |
| Tox | -0.2595 | 0.13 | 0.351 | 1 | 3 |
| Gm15261 | -0.2598 | 0.056 | 0.243 | 0.0038 | 4 |
| Igfbpl1 | -0.2599 | 0 | 0.177 | 0.0055 | 8 |
| Akap17b | -0.2599 | 0.245 | 0.405 | 0.0135 | 0 |
| Rpl28 | -0.2599 | 0.976 | 0.97 | 1 | 3 |
| Med28 | -0.26 | 0.553 | 0.739 | 1 | 3 |
| Usp22 | -0.26 | 0.705 | 0.849 | 1 | 7 |
| Cd99l2 | -0.2601 | 0.493 | 0.656 | 0.0109 | 2 |
| Cntn2 | -0.2602 | 0.072 | 0.256 | 0.0005 | 5 |
| Grid2 | -0.2602 | 0.072 | 0.251 | 0.0007 | 5 |
| AI480526 | -0.2602 | 0.206 | 0.354 | 1 | 6 |
| Nav2 | -0.2602 | 0.051 | 0.262 | 0.2204 | 7 |
| Sox3 | -0.2604 | 0 | 0.196 | 1.12E-07 | 5 |
| Tmem106 | -0.2604 | 0.359 | 0.529 | 1 | 7 |
| Wbscr17 | -0.2604 | 0.013 | 0.195 | 0.019 | 7 |
| Kctd8 | -0.2605 | 0.046 | 0.255 | 5.07E-08 | 1 |
| Gnas | -0.2606 | 1 | 1 | 0.0008 | 2 |
| Herc1 | -0.2606 | 0.479 | 0.675 | 1 | 8 |
| Amd1 | -0.2607 | 0.333 | 0.491 | 1 | 9 |
| Tnik | -0.2608 | 0.536 | 0.689 | 1 | 5 |
| Unc13c | -0.2609 | 0 | 0.1 | 1.20E-05 | 2 |
| Atp6v0a1 | -0.261 | 0.856 | 0.843 | 0.0002 | 6 |
| Mrps6 | -0.2611 | 0.335 | 0.495 | 0.0457 | 0 |
| Tubb2b | -0.2611 | 0.407 | 0.624 | 1 | 3 |

| | | | | | |
|---|---|---|---|---|---|
| Dnajc12 | -0.2612 | 0.2 | 0.406 | 1 | 9 |
| Yme1l1 | -0.2613 | 0.412 | 0.562 | 1 | 6 |
| Sdhaf4 | -0.2613 | 0.507 | 0.719 | 0.1881 | 8 |
| Kif21a | -0.2613 | 0.712 | 0.805 | 0.0427 | 2 |
| Tmem47 | -0.2613 | 0.197 | 0.383 | 1 | 8 |
| Actb | -0.2613 | 0.995 | 0.999 | 0.0903 | 0 |
| Camkv | -0.2614 | 0.438 | 0.49 | 0.0025 | 2 |
| Cntn2 | -0.2614 | 0.064 | 0.254 | 0.5186 | 7 |
| C1galt1c1 | -0.2615 | 0.1 | 0.309 | 1 | 9 |
| Slc24a5 | -0.2616 | 0.401 | 0.59 | 0.1329 | 1 |
| Mgat4c | -0.2616 | 0.113 | 0.232 | 1 | 8 |
| Tomm20 | -0.2617 | 0.986 | 0.985 | 1 | 2 |
| Snhg9 | -0.2617 | 0.324 | 0.472 | 1 | 8 |
| Trappc3 | -0.2618 | 0.309 | 0.63 | 0.758 | 3 |
| Pvrl3 | -0.262 | 0.25 | 0.381 | 1 | 4 |
| Fam196a | -0.2621 | 0.007 | 0.218 | 5.30E-12 | 1 |
| Mdh2 | -0.2621 | 0.951 | 0.918 | 1 | 3 |
| Cd99l2 | -0.2621 | 0.449 | 0.648 | 1 | 7 |
| Slc18a2 | -0.2622 | 0.014 | 0.147 | 5.49E-07 | 2 |
| Aatk | -0.2626 | 0.534 | 0.615 | 0.5238 | 2 |
| Cacna2d3 | -0.2627 | 0.235 | 0.462 | 1.92E-05 | 0 |
| Rps8 | -0.2627 | 1 | 0.998 | 0.0003 | 0 |
| Syngr1 | -0.2628 | 0.33 | 0.535 | 1 | 6 |
| Npy5r | -0.2628 | 0 | 0.203 | 8.83E-05 | 8 |
| Cyp51 | -0.2628 | 0.247 | 0.404 | 1 | 5 |
| Thsd7b | -0.2629 | 0.204 | 0.385 | 0.0287 | 1 |
| Slc18a2 | -0.2629 | 0.008 | 0.145 | 0.0015 | 3 |
| Zfp950 | -0.2629 | 0.372 | 0.536 | 1 | 7 |
| Tsc22d1 | -0.2629 | 0.782 | 0.868 | 1 | 7 |
| Pnpla8 | -0.2629 | 0.227 | 0.449 | 1 | 6 |
| RP24-236 | -0.2629 | 0.118 | 0.326 | 1 | 10 |
| Matk | -0.2629 | 0.651 | 0.761 | 1 | 2 |
| Hap1 | -0.263 | 0.99 | 0.997 | 0.0166 | 5 |
| Mgat5 | -0.263 | 0.154 | 0.364 | 1 | 7 |
| Ctnnd2 | -0.263 | 0.179 | 0.371 | 1 | 7 |
| Med30 | -0.2631 | 0.235 | 0.411 | 1 | 10 |
| Tcf12 | -0.2631 | 0.25 | 0.42 | 0.0042 | 0 |
| Mrpl16 | -0.2631 | 0.033 | 0.291 | 1 | 3 |
| Spats2l | -0.2631 | 0.039 | 0.215 | 3.22E-06 | 1 |
| Map2k1 | -0.2632 | 0.412 | 0.568 | 0.0063 | 5 |
| Clptm1 | -0.2632 | 0.466 | 0.553 | 0.1496 | 2 |
| Map1a | -0.2632 | 0.384 | 0.516 | 4.85E-06 | 2 |

| | | | | | |
|---|---|---|---|---|---|
| Rpl38 | -0.2633 | 0.995 | 0.998 | 0.0124 | 0 |
| As3mt | -0.2633 | 0.145 | 0.341 | 0.0009 | 0 |
| Plppr4 | -0.2634 | 0.436 | 0.566 | 1 | 7 |
| Cox17 | -0.2635 | 0.76 | 0.888 | 0.0023 | 0 |
| Ctla2a | -0.2635 | 0.005 | 0.138 | 4.37E-07 | 0 |
| Ubc | -0.2635 | 0.825 | 0.92 | 1 | 5 |
| Calb1 | -0.2635 | 0.65 | 0.788 | 0.0743 | 0 |
| Herc3 | -0.2635 | 0.154 | 0.353 | 1 | 7 |
| Tmed9 | -0.2637 | 0.691 | 0.849 | 1 | 5 |
| Fgf13 | -0.2637 | 0.113 | 0.285 | 0.0272 | 8 |
| Asic2 | -0.2638 | 0.216 | 0.428 | 0.0032 | 5 |
| Olfm1 | -0.2638 | 0.747 | 0.863 | 1 | 2 |
| 1110004I | -0.2639 | 0.972 | 0.962 | 1 | 4 |
| Naca | -0.264 | 0.96 | 0.991 | 0.0021 | 0 |
| Tpd52l1 | -0.264 | 0.375 | 0.485 | 0.1623 | 1 |
| Fam134b | -0.264 | 0.235 | 0.434 | 1 | 10 |
| Pcsk1n | -0.2641 | 1 | 1 | 1 | 7 |
| Kpna3 | -0.2642 | 0.205 | 0.413 | 1 | 7 |
| Ttll5 | -0.2642 | 0.083 | 0.271 | 0.0105 | 4 |
| Fez1 | -0.2642 | 0.697 | 0.851 | 7.98E-05 | 1 |
| 2900055. | -0.2643 | 0.085 | 0.281 | 1 | 8 |
| Rps16 | -0.2643 | 0.975 | 0.987 | 0.0351 | 0 |
| Mapk8ip2 | -0.2643 | 0.551 | 0.688 | 1 | 7 |
| Cfap36 | -0.2643 | 0.553 | 0.74 | 1 | 3 |
| Taz | -0.2644 | 0.487 | 0.617 | 1 | 7 |
| Adcy2 | -0.2644 | 0.067 | 0.313 | 1 | 9 |
| Lpgat1 | -0.2644 | 0.282 | 0.502 | 0.4048 | 8 |
| Cpne6 | -0.2645 | 0.026 | 0.229 | 0.0163 | 7 |
| Arhgap20 | -0.2645 | 0.167 | 0.404 | 1 | 7 |
| Rps12 | -0.2645 | 0.77 | 0.845 | 0.1826 | 0 |
| Cdkn1c | -0.2647 | 0 | 0.127 | 1 | 9 |
| Crtac1 | -0.2647 | 0.064 | 0.224 | 1 | 7 |
| Adgrb2 | -0.2647 | 0.299 | 0.46 | 0.0932 | 5 |
| Adgrb1 | -0.2648 | 0.269 | 0.476 | 1 | 7 |
| Gm16105 | -0.2649 | 0.299 | 0.481 | 1 | 6 |
| Grid2 | -0.2649 | 0.075 | 0.26 | 1.43E-10 | 2 |
| Gabrg3 | -0.2649 | 0.256 | 0.41 | 1 | 7 |
| Nudt2 | -0.2651 | 0.118 | 0.382 | 1 | 10 |
| Tnrc6c | -0.2651 | 0.408 | 0.592 | 1 | 8 |
| Gabra3 | -0.2651 | 0.155 | 0.344 | 1 | 6 |
| Araf | -0.2653 | 0.897 | 0.924 | 1 | 6 |
| R3hdm2 | -0.2653 | 0.22 | 0.508 | 1 | 3 |

| | | | | | |
|---|---|---|---|---|---|
| Zwint | -0.2654 | 0.987 | 0.998 | 2.89E-07 | 1 |
| Tmem178 | -0.2654 | 0 | 0.195 | 1 | 10 |
| Araf | -0.2654 | 0.873 | 0.925 | 1 | 8 |
| Rpl27a | -0.2655 | 0.99 | 0.996 | 0.0014 | 0 |
| Pkia | -0.2656 | 0.437 | 0.579 | 1 | 8 |
| Bend6 | -0.2656 | 0.256 | 0.438 | 1 | 7 |
| Mcfd2 | -0.2656 | 0.588 | 0.793 | 0.1787 | 5 |
| Adam23 | -0.2656 | 0.175 | 0.369 | 1 | 6 |
| Slc18a2 | -0.2657 | 0 | 0.142 | 0.0003 | 5 |
| Plppr3 | -0.2657 | 0.464 | 0.615 | 1 | 5 |
| Sf3b6 | -0.2659 | 0.815 | 0.9 | 0.0029 | 0 |
| Tma7 | -0.2659 | 0.959 | 0.969 | 1 | 3 |
| Ssbp4 | -0.2662 | 0.753 | 0.877 | 0.3662 | 5 |
| Chodl | -0.2662 | 0.113 | 0.26 | 1 | 6 |
| Sncb | -0.2663 | 0.846 | 0.923 | 1 | 7 |
| Scp2 | -0.2663 | 0.414 | 0.585 | 0.0009 | 1 |
| Fam234b | -0.2664 | 0.196 | 0.394 | 1 | 6 |
| Lrrn3 | -0.2664 | 0.062 | 0.294 | 3.43E-06 | 5 |
| Anxa5 | -0.2665 | 0.116 | 0.26 | 6.95E-08 | 2 |
| Cntnap2 | -0.2666 | 0.333 | 0.504 | 1 | 7 |
| Tuba4a | -0.2667 | 0.276 | 0.54 | 1 | 3 |
| Cnnm1 | -0.2668 | 0.062 | 0.279 | 1.14E-06 | 5 |
| Rpl15 | -0.2668 | 0.95 | 0.976 | 0.0016 | 0 |
| Cited2 | -0.2668 | 0.38 | 0.508 | 1 | 0 |
| Tmsb10 | -0.2669 | 1 | 0.997 | 0.0004 | 1 |
| Frrs1l | -0.2669 | 0.514 | 0.622 | 5.61E-06 | 2 |
| Enc1 | -0.2669 | 0.106 | 0.29 | 1 | 3 |
| Ric8 | -0.2671 | 0.059 | 0.305 | 1 | 10 |
| Mob4 | -0.2671 | 0.353 | 0.633 | 1 | 10 |
| Auts2 | -0.2671 | 0.675 | 0.752 | 0.0392 | 0 |
| Gde1 | -0.2671 | 0.808 | 0.884 | 1 | 2 |
| Stx12 | -0.2672 | 0.423 | 0.67 | 1 | 3 |
| Grid2 | -0.2673 | 0.057 | 0.258 | 0.007 | 3 |
| Nfix | -0.2674 | 0.007 | 0.116 | 4.84E-06 | 2 |
| Brinp2 | -0.2675 | 0.175 | 0.338 | 1 | 6 |
| Rab26 | -0.2675 | 0.128 | 0.35 | 0.8666 | 7 |
| Cox7c | -0.2676 | 0.967 | 0.995 | 1.13E-09 | 1 |
| Marf1 | -0.2676 | 0.471 | 0.586 | 1 | 10 |
| Dpp10 | -0.2677 | 0.141 | 0.309 | 1 | 8 |
| Nsmce2 | -0.2677 | 0.059 | 0.297 | 1 | 10 |
| Trpc5 | -0.2679 | 0.185 | 0.301 | 0.005 | 2 |
| Atp1a3 | -0.268 | 0.948 | 0.945 | 1.47E-05 | 6 |

| | | | | | |
|---|---|---|---|---|---|
| Fam69a | -0.268 | 0.113 | 0.36 | 0.8907 | 6 |
| Th | -0.268 | 0 | 0.117 | 0.1637 | 7 |
| Cdh4 | -0.2681 | 0.01 | 0.192 | 3.81E-06 | 5 |
| Stmn3 | -0.2681 | 1 | 1 | 1 | 7 |
| Efna5 | -0.2681 | 0.115 | 0.322 | 0.7157 | 7 |
| Cdc42bpa | -0.2682 | 0.3 | 0.513 | 1 | 9 |
| AI480526 | -0.2682 | 0.179 | 0.354 | 1 | 7 |
| Gnas | -0.2682 | 1 | 1 | 1 | 7 |
| Rab4a | -0.2682 | 0.252 | 0.557 | 1 | 3 |
| Faap20 | -0.2685 | 0.098 | 0.377 | 1 | 3 |
| Phka2 | -0.2686 | 0.033 | 0.237 | 1 | 9 |
| Lynx1 | -0.2686 | 0.021 | 0.189 | 4.56E-16 | 2 |
| Crip2 | -0.2687 | 0.52 | 0.688 | 1 | 3 |
| Trim2 | -0.2687 | 0.436 | 0.621 | 1 | 7 |
| Suv420h1 | -0.2689 | 0.033 | 0.297 | 0.905 | 9 |
| Epb41l3 | -0.269 | 0.408 | 0.57 | 0.0358 | 1 |
| Dzank1 | -0.269 | 0.485 | 0.59 | 0.7919 | 5 |
| Fubp1 | -0.2691 | 0.467 | 0.73 | 1 | 9 |
| Mrto4 | -0.2692 | 0.171 | 0.452 | 1 | 3 |
| Ccdc127 | -0.2692 | 0.233 | 0.458 | 1 | 9 |
| Nlgn1 | -0.2692 | 0.309 | 0.478 | 0.0498 | 5 |
| Ptn | -0.2693 | 0.495 | 0.512 | 1 | 6 |
| Tead1 | -0.2694 | 0.128 | 0.328 | 1 | 7 |
| Nop10 | -0.2695 | 0.935 | 0.954 | 1 | 3 |
| Gpld1 | -0.2696 | 0.067 | 0.271 | 1 | 9 |
| Creg2 | -0.2697 | 0.021 | 0.24 | 1.32E-08 | 5 |
| Rpl22l1 | -0.2698 | 0.995 | 0.998 | 0.0009 | 0 |
| Ufsp2 | -0.2698 | 0.176 | 0.395 | 1 | 10 |
| Ldb2 | -0.2699 | 0.033 | 0.253 | 1 | 9 |
| Plekhb2 | -0.27 | 0.7 | 0.775 | 1 | 9 |
| Ank2 | -0.27 | 0.784 | 0.845 | 0.0247 | 6 |
| Pnrc1 | -0.2701 | 0.63 | 0.745 | 1 | 4 |
| Mmd | -0.2702 | 0.155 | 0.359 | 1 | 6 |
| Chchd6 | -0.2703 | 0.732 | 0.809 | 1 | 3 |
| 3632451( | -0.2703 | 0.145 | 0.319 | 0.0001 | 0 |
| Alg2 | -0.2704 | 0.538 | 0.76 | 0.1544 | 7 |
| Galnt11 | -0.2705 | 0.165 | 0.397 | 1 | 6 |
| Cds2 | -0.2705 | 0.535 | 0.698 | 1 | 8 |
| Rpl29 | -0.2706 | 0.885 | 0.952 | 0.0001 | 0 |
| Jund | -0.2706 | 0.976 | 0.989 | 1 | 3 |
| Slc12a5 | -0.2706 | 0.855 | 0.913 | 1 | 1 |
| Ntrk2 | -0.2708 | 0.541 | 0.68 | 0.0185 | 2 |

| | | | | | |
|---|---|---|---|---|---|
| H3f3a | -0.2708 | 0.934 | 0.954 | 0.0988 | 1 |
| Apex1 | -0.2709 | 0.295 | 0.502 | 0.0009 | 0 |
| Phyhip | -0.271 | 0.077 | 0.267 | 0.802 | 7 |
| Srsf4 | -0.271 | 0.118 | 0.349 | 1 | 10 |
| Gabbr2 | -0.271 | 0.211 | 0.4 | 1 | 8 |
| Atp6v0e2 | -0.2711 | 0.859 | 0.932 | 1 | 7 |
| Mrpl41 | -0.2711 | 0.813 | 0.88 | 1 | 3 |
| Unc5c | -0.2712 | 0.295 | 0.48 | 0.0013 | 0 |
| Sppl3 | -0.2712 | 0.216 | 0.449 | 1 | 6 |
| Flot2 | -0.2712 | 0.144 | 0.386 | 1 | 6 |
| Ttc19 | -0.2713 | 0.588 | 0.737 | 1 | 5 |
| Cnr1 | -0.2715 | 0.041 | 0.225 | 1 | 6 |
| Ndufaf4 | -0.2716 | 0.176 | 0.396 | 1 | 10 |
| Fubp1 | -0.2716 | 0.606 | 0.731 | 1 | 8 |
| Etnk1 | -0.2716 | 0.753 | 0.806 | 0.0354 | 6 |
| Pclo | -0.2717 | 0.563 | 0.781 | 1 | 8 |
| Srsf1 | -0.2717 | 0.479 | 0.711 | 1 | 8 |
| Nptx1 | -0.2719 | 0.062 | 0.21 | 0.0163 | 5 |
| Nell2 | -0.272 | 0.299 | 0.481 | 1 | 6 |
| Ank3 | -0.2721 | 0.69 | 0.862 | 1 | 8 |
| Tmem147 | -0.2721 | 0.788 | 0.841 | 1 | 2 |
| Setx | -0.2722 | 0.205 | 0.385 | 1 | 7 |
| Dclk1 | -0.2722 | 0.596 | 0.718 | 0.0917 | 2 |
| Rab3b | -0.2722 | 0.134 | 0.338 | 1 | 6 |
| Uba52 | -0.2722 | 0.785 | 0.893 | 0.0024 | 0 |
| Zfp938 | -0.2722 | 0.059 | 0.348 | 1 | 10 |
| Spock1 | -0.2723 | 0.384 | 0.402 | 0.0286 | 2 |
| Enah | -0.2723 | 0.551 | 0.701 | 1 | 7 |
| Hdac9 | -0.2724 | 0 | 0.222 | 1 | 10 |
| Rnf112 | -0.2724 | 0.398 | 0.59 | 0.16 | 4 |
| Ccdc59 | -0.2725 | 0.179 | 0.468 | 1 | 3 |
| Dnm1 | -0.2725 | 0.551 | 0.681 | 1 | 7 |
| Rims3 | -0.2725 | 0.437 | 0.599 | 1 | 8 |
| Chn1 | -0.2725 | 0.528 | 0.696 | 1 | 3 |
| Syngr3 | -0.2727 | 0.897 | 0.909 | 1 | 7 |
| Selk | -0.2727 | 0.99 | 0.991 | 3.83E-05 | 6 |
| Gm1580( | -0.2728 | 0.269 | 0.491 | 1 | 7 |
| Ogfrl1 | -0.2732 | 0.568 | 0.653 | 1 | 2 |
| Osbpl9 | -0.2732 | 0.2 | 0.416 | 1 | 9 |
| Rps26 | -0.2732 | 0.895 | 0.95 | 0.0135 | 0 |
| Cadps | -0.2733 | 0.392 | 0.597 | 0.0077 | 5 |
| Tmeff2 | -0.2734 | 0.051 | 0.259 | 0.0761 | 7 |

| | | | | | |
|---|---|---|---|---|---|
| Ccnt2 | -0.2735 | 0.436 | 0.596 | 1 | 7 |
| Trappc13 | -0.2735 | 0.267 | 0.406 | 1 | 9 |
| Lrp1b | -0.2736 | 0.583 | 0.651 | 1 | 4 |
| Gas7 | -0.2736 | 0.155 | 0.346 | 0.0011 | 5 |
| Ar | -0.2736 | 0.213 | 0.432 | 0.0318 | 4 |
| Ptma | -0.2736 | 0.987 | 0.987 | 0.5012 | 1 |
| Tspan7 | -0.2736 | 0.794 | 0.853 | 0.0009 | 6 |
| Psmd10 | -0.2736 | 0.059 | 0.309 | 1 | 10 |
| Syt4 | -0.2736 | 0.876 | 0.876 | 0.1207 | 6 |
| Gpm6a | -0.2738 | 0.979 | 0.965 | 1 | 5 |
| 2010005I | -0.2738 | 0 | 0.172 | 1 | 9 |
| Tceal5 | -0.2738 | 0.569 | 0.749 | 1 | 3 |
| Pam | -0.2738 | 0.572 | 0.704 | 1 | 1 |
| Pnck | -0.2738 | 0.87 | 0.934 | 1 | 2 |
| Tmem130 | -0.2739 | 0.974 | 0.975 | 1 | 7 |
| Efna5 | -0.2741 | 0.103 | 0.327 | 1 | 6 |
| Arpc5 | -0.2741 | 0.626 | 0.773 | 1 | 3 |
| Phka2 | -0.2741 | 0.026 | 0.247 | 0.0132 | 7 |
| Fam204a | -0.2742 | 0.167 | 0.38 | 1 | 9 |
| Ahsa1 | -0.2742 | 0.398 | 0.676 | 1 | 3 |
| Gnb1 | -0.2742 | 0.789 | 0.949 | 0.0736 | 8 |
| Gabre | -0.2744 | 0.007 | 0.177 | 1.28E-09 | 2 |
| Scn2b | -0.2744 | 0.394 | 0.523 | 1 | 8 |
| Gpr83 | -0.2745 | 0 | 0.19 | 0.0007 | 8 |
| Lynx1 | -0.2746 | 0 | 0.18 | 0.0053 | 7 |
| Plxna2 | -0.2747 | 0.01 | 0.226 | 7.41E-09 | 5 |
| Calm1 | -0.2748 | 1 | 1 | 0.0322 | 0 |
| Chodl | -0.2749 | 0.102 | 0.263 | 0.2989 | 4 |
| Grm7 | -0.2749 | 0.013 | 0.236 | 0.001 | 7 |
| Camk2b | -0.275 | 0.718 | 0.793 | 1 | 7 |
| Smpd3 | -0.275 | 0.216 | 0.445 | 1 | 6 |
| Gng2 | -0.275 | 0.732 | 0.831 | 1 | 8 |
| Herc1 | -0.275 | 0.474 | 0.676 | 1 | 7 |
| Adcy2 | -0.275 | 0.141 | 0.319 | 1 | 7 |
| Efhd2 | -0.2751 | 0.205 | 0.371 | 1 | 7 |
| Copz1 | -0.2752 | 0.268 | 0.56 | 1 | 3 |
| Inafm1 | -0.2753 | 0.651 | 0.737 | 1 | 2 |
| Arf3 | -0.2753 | 0.767 | 0.836 | 0.008 | 2 |
| Atp2b4 | -0.2753 | 0.184 | 0.286 | 0.7693 | 1 |
| Tead1 | -0.2754 | 0.124 | 0.332 | 1 | 6 |
| Cd63 | -0.2754 | 0.329 | 0.484 | 0.05 | 2 |
| Ccl27a | -0.2754 | 0.496 | 0.731 | 0.9811 | 3 |

| | | | | | |
|---|---|---|---|---|---|
| Kif5a | -0.2754 | 0.664 | 0.728 | 0.0073 | 2 |
| Cd99l2 | -0.2755 | 0.454 | 0.652 | 0.1552 | 5 |
| Sec61b | -0.2755 | 0.68 | 0.799 | 0.0223 | 0 |
| Nisch | -0.2755 | 0.836 | 0.888 | 1 | 2 |
| Fam133b | -0.2756 | 0.176 | 0.479 | 1 | 10 |
| Cntnap1 | -0.2757 | 0.013 | 0.223 | 0.0045 | 7 |
| Dmd | -0.2757 | 0.141 | 0.321 | 1 | 8 |
| Sult4a1 | -0.2757 | 0.718 | 0.839 | 1 | 7 |
| Map1a | -0.2757 | 0.236 | 0.531 | 0.0334 | 3 |
| Nbea | -0.2759 | 0.487 | 0.667 | 1 | 7 |
| Serf2 | -0.276 | 0.984 | 0.956 | 1 | 3 |
| Asph | -0.2761 | 0.205 | 0.402 | 1 | 7 |
| Arhgap36 | -0.2761 | 0.151 | 0.203 | 1 | 2 |
| Tmbim6 | -0.2761 | 0.704 | 0.809 | 1 | 8 |
| Lonrf2 | -0.2762 | 0.521 | 0.591 | 9.30E-06 | 2 |
| Fau | -0.2762 | 1 | 0.997 | 0.0026 | 0 |
| Ubxn1 | -0.2762 | 0.62 | 0.792 | 2.64E-05 | 0 |
| Tmem191 | -0.2762 | 0.596 | 0.765 | 0.0824 | 2 |
| Blcap | -0.2763 | 0.761 | 0.833 | 1 | 8 |
| Cpne5 | -0.2763 | 0.113 | 0.301 | 1 | 6 |
| Gabre | -0.2763 | 0 | 0.174 | 2.44E-06 | 3 |
| Zic1 | -0.2765 | 0.009 | 0.22 | 6.62E-06 | 4 |
| Csf2ra | -0.2766 | 0.118 | 0.41 | 1 | 10 |
| Syngr1 | -0.2766 | 0.333 | 0.531 | 1 | 7 |
| Tenm4 | -0.2766 | 0.395 | 0.549 | 1 | 1 |
| Sec61g | -0.2766 | 0.905 | 0.966 | 5.27E-05 | 0 |
| Th | -0.2767 | 0.007 | 0.124 | 1.53E-06 | 2 |
| Zfp106 | -0.2768 | 0.236 | 0.522 | 1 | 3 |
| Col25a1 | -0.277 | 0.014 | 0.213 | 0.0008 | 8 |
| Tmem126 | -0.2771 | 0.195 | 0.498 | 1 | 3 |
| Smg1 | -0.2771 | 0.155 | 0.409 | 0.9486 | 6 |
| Sst | -0.2771 | 0 | 0.022 | 1 | 3 |
| Grcc10 | -0.2772 | 0.835 | 0.905 | 0.0024 | 0 |
| Polr3h | -0.2773 | 0.089 | 0.39 | 1 | 3 |
| Pole4 | -0.2775 | 0.059 | 0.289 | 1 | 10 |
| Cntn1 | -0.2776 | 0.561 | 0.643 | 0.0005 | 3 |
| Gprasp1 | -0.2776 | 0.979 | 0.974 | 1 | 2 |
| Calr | -0.2777 | 0.76 | 0.843 | 1 | 2 |
| Stk32c | -0.2777 | 0.362 | 0.553 | 0.0018 | 1 |
| Necap1 | -0.2778 | 0.382 | 0.627 | 1 | 3 |
| Abcc5 | -0.2779 | 0.603 | 0.71 | 1 | 7 |
| Mgst3 | -0.2779 | 0.231 | 0.404 | 1 | 7 |

| | | | | | |
|---|---|---|---|---|---|
| Soga3 | -0.278 | 0.567 | 0.803 | 1 | 9 |
| Pkp4 | -0.2781 | 0.103 | 0.317 | 0.4132 | 7 |
| Usp22 | -0.2782 | 0.676 | 0.85 | 1 | 8 |
| Lingo2 | -0.2783 | 0.103 | 0.31 | 0.0001 | 5 |
| Hrh3 | -0.2783 | 0.113 | 0.289 | 0.0039 | 5 |
| Epb41l4b | -0.2784 | 0.038 | 0.247 | 0.0211 | 7 |
| Rpl13 | -0.2784 | 0.995 | 0.997 | 0.0001 | 0 |
| Dnajb6 | -0.2784 | 0.967 | 0.946 | 1 | 3 |
| Faap20 | -0.2784 | 0.118 | 0.349 | 1 | 10 |
| Polr1d | -0.2784 | 0.767 | 0.817 | 1 | 9 |
| Erlec1 | -0.2784 | 0.466 | 0.605 | 0.1483 | 2 |
| Magi3 | -0.2785 | 0.014 | 0.255 | 1.59E-05 | 8 |
| Slirp | -0.2785 | 0.341 | 0.651 | 1 | 3 |
| Cirbp | -0.2786 | 0.974 | 0.972 | 0.0961 | 1 |
| Th | -0.2786 | 0 | 0.122 | 0.0342 | 3 |
| Trappc13 | -0.2786 | 0.237 | 0.428 | 6.90E-05 | 1 |
| Emc1 | -0.2786 | 0.231 | 0.443 | 1 | 7 |
| Atp9a | -0.2786 | 0.577 | 0.752 | 1 | 7 |
| Phlda3 | -0.2787 | 0.175 | 0.348 | 0.0021 | 5 |
| Camk2d | -0.2787 | 0.254 | 0.37 | 1 | 8 |
| Ngrn | -0.2787 | 0.236 | 0.544 | 1 | 3 |
| Cnksr2 | -0.2788 | 0.217 | 0.36 | 0.0154 | 1 |
| Tmem14c | -0.2788 | 0.5 | 0.688 | 0.0983 | 7 |
| Tcf4 | -0.2789 | 0.171 | 0.321 | 1 | 3 |
| Rit2 | -0.2789 | 0.84 | 0.898 | 0.0082 | 0 |
| Ntan1 | -0.2791 | 0.333 | 0.621 | 1 | 3 |
| Lynx1 | -0.2791 | 0 | 0.183 | 0.0023 | 6 |
| Ptprt | -0.2791 | 0.093 | 0.272 | 1 | 6 |
| Plcb4 | -0.2792 | 0.466 | 0.521 | 0.2646 | 2 |
| B4gat1 | -0.2792 | 0.592 | 0.682 | 1 | 8 |
| Kcnk2 | -0.2793 | 0.171 | 0.385 | 1 | 3 |
| Naa35 | -0.2793 | 0.353 | 0.52 | 1 | 10 |
| Gabrg1 | -0.2793 | 0.463 | 0.596 | 1 | 4 |
| Mdm4 | -0.2794 | 0.2 | 0.388 | 1 | 9 |
| Pnrc1 | -0.2794 | 0.651 | 0.747 | 0.0575 | 1 |
| Rxrg | -0.2795 | 0.024 | 0.162 | 1 | 3 |
| Pcdh20 | -0.2795 | 0 | 0.188 | 0.0014 | 8 |
| Calm1 | -0.2796 | 1 | 1 | 0.4457 | 4 |
| Ppp4c | -0.2796 | 0.118 | 0.35 | 1 | 10 |
| Dmxl2 | -0.2797 | 0.179 | 0.399 | 1 | 7 |
| Gstm7 | -0.2799 | 0.195 | 0.374 | 0.1762 | 0 |
| Cplx2 | -0.2799 | 0.093 | 0.287 | 1 | 6 |

| | | | | | |
|---|---|---|---|---|---|
| Ptpro | -0.2799 | 0.165 | 0.315 | 0.0474 | 5 |
| Pdgfa | -0.2799 | 0.269 | 0.516 | 1 | 7 |
| Rasgrp1 | -0.28 | 0.13 | 0.301 | 0.1258 | 4 |
| Micu3 | -0.28 | 0.803 | 0.906 | 1 | 8 |
| Chst8 | -0.2801 | 0.165 | 0.319 | 1 | 5 |
| Suds3 | -0.2801 | 0.294 | 0.513 | 1 | 10 |
| Tial1 | -0.2802 | 0.535 | 0.687 | 1 | 8 |
| Aes | -0.2802 | 1 | 0.99 | 1 | 3 |
| Lgi1 | -0.2803 | 0.059 | 0.284 | 4.86E-07 | 1 |
| Cystm1 | -0.2804 | 0.579 | 0.719 | 0.001 | 1 |
| Eif2s2 | -0.2805 | 0.789 | 0.845 | 1 | 3 |
| Spock2 | -0.2806 | 0.732 | 0.87 | 1 | 5 |
| Ralgapa1 | -0.2807 | 0.338 | 0.524 | 1 | 8 |
| Hs2st1 | -0.2807 | 0.179 | 0.387 | 1 | 7 |
| Sap30l | -0.2807 | 0.059 | 0.314 | 1 | 10 |
| Arc | -0.2808 | 0 | 0.117 | 1 | 9 |
| Elof1 | -0.2808 | 0.537 | 0.744 | 1 | 3 |
| Erp29 | -0.2809 | 0.936 | 0.962 | 0.0599 | 7 |
| Slc25a22 | -0.2809 | 0.206 | 0.437 | 1 | 6 |
| Nkain3 | -0.2809 | 0.041 | 0.239 | 0.1759 | 3 |
| Fam173a | -0.2809 | 0.842 | 0.927 | 1 | 2 |
| Hdac11 | -0.281 | 0.731 | 0.874 | 0.1621 | 7 |
| Ip6k1 | -0.281 | 0.4 | 0.613 | 1 | 9 |
| Sulf1 | -0.2813 | 0.016 | 0.199 | 4.43E-05 | 3 |
| Rhbdd2 | -0.2814 | 0.774 | 0.847 | 1 | 2 |
| Enc1 | -0.2815 | 0.13 | 0.291 | 2.79E-12 | 2 |
| Zcchc18 | -0.2815 | 0.99 | 1 | 0.0101 | 5 |
| Acsl3 | -0.2816 | 0.465 | 0.627 | 1 | 8 |
| Agap3 | -0.2817 | 0.495 | 0.683 | 0.0207 | 5 |
| Actb | -0.2818 | 1 | 0.998 | 1 | 3 |
| Ydjc | -0.2819 | 0.141 | 0.377 | 0.2445 | 7 |
| Serinc1 | -0.2821 | 0.923 | 0.97 | 1 | 7 |
| Negr1 | -0.2821 | 0.784 | 0.841 | 0.0013 | 6 |
| Yme1l1 | -0.2822 | 0.353 | 0.552 | 1 | 10 |
| Enc1 | -0.2823 | 0.093 | 0.287 | 1 | 6 |
| Spin1 | -0.2823 | 0.333 | 0.498 | 1 | 9 |
| Gabrg2 | -0.2824 | 0.568 | 0.69 | 0.2026 | 2 |
| Nkain3 | -0.2825 | 0.064 | 0.229 | 1 | 7 |
| Ptp4a1 | -0.2825 | 0.186 | 0.415 | 1 | 6 |
| Mfsd6 | -0.2825 | 0.247 | 0.426 | 0.0015 | 5 |
| Rap1gap | -0.2826 | 0.196 | 0.436 | 1 | 6 |
| Ets2 | -0.2827 | 0.081 | 0.297 | 0.1988 | 3 |

| | | | | | |
|---|---|---|---|---|---|
| St3gal5 | -0.2827 | 0.235 | 0.414 | 1 | 10 |
| Pianp | -0.2827 | 0.833 | 0.927 | 1 | 7 |
| Lsamp | -0.2827 | 0.795 | 0.847 | 1 | 7 |
| Churc1 | -0.2828 | 0.367 | 0.624 | 1 | 9 |
| Grid1 | -0.2828 | 0.031 | 0.252 | 1.98E-07 | 5 |
| Gabrb1 | -0.2828 | 0.804 | 0.861 | 1 | 5 |
| Map4 | -0.283 | 0.26 | 0.558 | 1 | 3 |
| Dync1h1 | -0.283 | 0.486 | 0.617 | 0.0013 | 2 |
| Cyb561 | -0.283 | 0.397 | 0.543 | 1 | 7 |
| Elmo1 | -0.283 | 0.093 | 0.289 | 1 | 6 |
| Rorb | -0.283 | 0.545 | 0.697 | 0.1981 | 0 |
| Nlgn3 | -0.2831 | 0.295 | 0.453 | 1 | 7 |
| Nptn | -0.2832 | 0.756 | 0.831 | 1 | 7 |
| Rgs17 | -0.2832 | 0.585 | 0.704 | 0.4816 | 3 |
| Scn3a | -0.2833 | 0.361 | 0.524 | 0.1546 | 5 |
| Flrt2 | -0.2834 | 0.014 | 0.243 | 2.02E-05 | 8 |
| Adcy3 | -0.2834 | 0.141 | 0.353 | 0.2434 | 8 |
| Ntrk2 | -0.2834 | 0.526 | 0.675 | 1 | 5 |
| Mbd3 | -0.2836 | 0.358 | 0.653 | 1 | 3 |
| Oxr1 | -0.2837 | 0.216 | 0.386 | 0.0004 | 5 |
| Ucp2 | -0.2837 | 0.227 | 0.34 | 1 | 5 |
| Epha4 | -0.2838 | 0.155 | 0.347 | 1 | 8 |
| Tead1 | -0.2838 | 0.106 | 0.339 | 0.0266 | 3 |
| Rpl10a | -0.2839 | 0.967 | 0.973 | 1 | 3 |
| Zic1 | -0.284 | 0 | 0.202 | 1 | 10 |
| Cox7a2l | -0.2841 | 0.691 | 0.784 | 1 | 3 |
| Phpt1 | -0.2841 | 0.333 | 0.6 | 1 | 9 |
| Syp | -0.2843 | 0.907 | 0.938 | 1.43E-06 | 6 |
| Tmem128 | -0.2843 | 0.124 | 0.4 | 1 | 6 |
| Gigyf1 | -0.2843 | 0.175 | 0.383 | 1 | 6 |
| Enpp5 | -0.2844 | 0.577 | 0.709 | 1 | 7 |
| Fam73a | -0.2845 | 0.128 | 0.376 | 0.3209 | 7 |
| Mllt11 | -0.2846 | 0.926 | 0.945 | 0.3909 | 4 |
| Jun | -0.2847 | 0.569 | 0.549 | 1 | 3 |
| Myo10 | -0.2849 | 0.133 | 0.315 | 1 | 9 |
| Ssbp2 | -0.2849 | 0.641 | 0.754 | 1 | 7 |
| Pfn1 | -0.2849 | 0.715 | 0.84 | 0.0015 | 0 |
| Serinc1 | -0.285 | 0.907 | 0.973 | 7.03E-10 | 6 |
| Lingo2 | -0.285 | 0.067 | 0.298 | 1 | 9 |
| Bsg | -0.2851 | 1 | 0.998 | 1 | 7 |
| Med19 | -0.2851 | 0.118 | 0.351 | 1 | 10 |
| Sox2 | -0.2852 | 0.44 | 0.607 | 0.0017 | 0 |

| | | | | | |
|---|---|---|---|---|---|
| Lynx1 | -0.2854 | 0 | 0.188 | 1.64E-06 | 3 |
| Cacna1d | -0.2855 | 0.395 | 0.547 | 0.1189 | 1 |
| Pak3 | -0.2855 | 0.68 | 0.844 | 0.1048 | 5 |
| Mrpl11 | -0.2855 | 0.447 | 0.683 | 1 | 3 |
| Ptges3 | -0.2856 | 0.911 | 0.922 | 1 | 3 |
| Chpt1 | -0.2857 | 0.052 | 0.288 | 0.0094 | 6 |
| Fxyd7 | -0.2857 | 0.505 | 0.68 | 1 | 5 |
| Pkia | -0.2857 | 0.433 | 0.574 | 1 | 9 |
| Sar1b | -0.2857 | 0.651 | 0.756 | 1 | 2 |
| Olfm1 | -0.2857 | 0.731 | 0.857 | 0.52 | 7 |
| Slc6a17 | -0.2858 | 0.33 | 0.504 | 1 | 6 |
| l7Rn6 | -0.2858 | 0.434 | 0.595 | 0.0008 | 1 |
| Ndufb8 | -0.2858 | 0.959 | 0.975 | 1 | 3 |
| Akap9 | -0.2859 | 0.726 | 0.855 | 1 | 2 |
| Rplp2 | -0.286 | 0.965 | 0.99 | 7.72E-05 | 0 |
| Kifc2 | -0.286 | 0.205 | 0.41 | 1 | 7 |
| Aprt | -0.2861 | 0.398 | 0.659 | 1 | 3 |
| Slc9a6 | -0.2861 | 0.295 | 0.458 | 1 | 7 |
| Chd5 | -0.2861 | 0.564 | 0.79 | 0.074 | 7 |
| Golgb1 | -0.2863 | 0.733 | 0.851 | 1 | 2 |
| Tox3 | -0.2863 | 0.24 | 0.409 | 0.0182 | 0 |
| Faim | -0.2864 | 0.106 | 0.402 | 1 | 3 |
| Pfn1 | -0.2865 | 0.724 | 0.832 | 3.65E-05 | 1 |
| Atp9a | -0.2867 | 0.619 | 0.751 | 1 | 5 |
| Grcc10 | -0.2867 | 0.8 | 0.895 | 1 | 9 |
| Sgsm1 | -0.2868 | 0.308 | 0.519 | 1 | 7 |
| Elavl2 | -0.2868 | 0.165 | 0.332 | 0.0001 | 5 |
| Emc7 | -0.2868 | 0.742 | 0.859 | 0.0004 | 6 |
| Chchd7 | -0.2868 | 0.28 | 0.507 | 0.0001 | 0 |
| Astn1 | -0.2868 | 0.623 | 0.738 | 1 | 2 |
| Gas6 | -0.2869 | 0.103 | 0.306 | 0.3258 | 7 |
| Ctnnbip1 | -0.2869 | 0.098 | 0.379 | 1 | 3 |
| Fhad1 | -0.287 | 0.052 | 0.237 | 0.0001 | 5 |
| Shisa9 | -0.2871 | 0.013 | 0.241 | 0.0004 | 7 |
| Spint2 | -0.2871 | 0.885 | 0.92 | 1 | 7 |
| Hbegf | -0.2874 | 0.105 | 0.232 | 0.0511 | 1 |
| Ncam1 | -0.2874 | 0.91 | 0.955 | 1 | 7 |
| Gck | -0.2874 | 0.038 | 0.281 | 0.0003 | 7 |
| Olfm3 | -0.2877 | 0.014 | 0.228 | 0.0001 | 8 |
| Sulf1 | -0.2877 | 0 | 0.192 | 0.0004 | 7 |
| Dynlt3 | -0.2877 | 0.122 | 0.415 | 1 | 3 |
| Pdhb | -0.2878 | 0.236 | 0.567 | 1 | 3 |

| | | | | | |
|---|---|---|---|---|---|
| Elavl2 | -0.2879 | 0.122 | 0.341 | 1 | 3 |
| Tmem5 | -0.288 | 0.235 | 0.46 | 1 | 10 |
| Creld1 | -0.288 | 0.268 | 0.474 | 1 | 6 |
| Slc24a3 | -0.288 | 0.217 | 0.423 | 0.0032 | 1 |
| Cntnap5a | -0.2881 | 0.122 | 0.337 | 0.0583 | 3 |
| Apbb2 | -0.2881 | 0.13 | 0.358 | 1 | 3 |
| Lyrm4 | -0.2882 | 0.118 | 0.352 | 1 | 10 |
| Saraf | -0.2882 | 0.887 | 0.938 | 8.33E-08 | 6 |
| Edf1 | -0.2882 | 0.959 | 0.975 | 1 | 3 |
| Ptpra | -0.2882 | 0.449 | 0.647 | 0.3429 | 7 |
| Hlf | -0.2883 | 0.285 | 0.553 | 1 | 3 |
| Cbarp | -0.2884 | 0.901 | 0.969 | 1 | 8 |
| Nrp1 | -0.2885 | 0.197 | 0.369 | 1 | 8 |
| Phpt1 | -0.2886 | 0.454 | 0.614 | 0.0003 | 1 |
| Emd | -0.2888 | 0.118 | 0.345 | 1 | 10 |
| Pdia4 | -0.2888 | 0.179 | 0.399 | 1 | 7 |
| mt-Nd4 | -0.2888 | 1 | 1 | 1 | 2 |
| Fhad1 | -0.2889 | 0.055 | 0.246 | 6.87E-11 | 2 |
| Scn2a1 | -0.289 | 0.814 | 0.889 | 1 | 5 |
| Ttc14 | -0.289 | 0.564 | 0.753 | 0.1719 | 7 |
| Napb | -0.2891 | 0.361 | 0.483 | 0.0232 | 5 |
| Stk32a | -0.2892 | 0.329 | 0.423 | 0.3039 | 1 |
| Ppp1r37 | -0.2893 | 0.235 | 0.429 | 1 | 10 |
| Dnajb14 | -0.2893 | 0.603 | 0.709 | 1 | 7 |
| Tmem132 | -0.2894 | 0.11 | 0.318 | 1.02E-05 | 0 |
| Olfm3 | -0.2895 | 0 | 0.218 | 1 | 10 |
| Cdkn1c | -0.2895 | 0.015 | 0.147 | 3.95E-06 | 0 |
| Celf5 | -0.2896 | 0.732 | 0.831 | 1 | 8 |
| Lsamp | -0.2896 | 0.753 | 0.852 | 0.0002 | 6 |
| Marcksl1 | -0.2897 | 0.875 | 0.96 | 9.67E-06 | 0 |
| Rab34 | -0.2897 | 0.169 | 0.395 | 0.1829 | 8 |
| Cetn3 | -0.2897 | 0.5 | 0.67 | 1 | 9 |
| Tmx1 | -0.2898 | 0.351 | 0.527 | 1 | 6 |
| Abhd16a | -0.2898 | 0.144 | 0.388 | 0.5307 | 6 |
| Nop58 | -0.2899 | 0.195 | 0.547 | 1 | 3 |
| Hmox2 | -0.2899 | 0.407 | 0.65 | 1 | 3 |
| Sgip1 | -0.2902 | 0.637 | 0.807 | 0.0051 | 2 |
| Dnajc2 | -0.2903 | 0.122 | 0.439 | 1 | 3 |
| Serp2 | -0.2905 | 0.808 | 0.893 | 1 | 7 |
| Hpcal4 | -0.2907 | 0.436 | 0.554 | 1 | 7 |
| Gm36266 | -0.2907 | 0 | 0.122 | 2.53E-07 | 0 |
| Galnt16 | -0.2907 | 0.187 | 0.398 | 0.0218 | 3 |

| | | | | | |
|---|---|---|---|---|---|
| Fos | -0.2908 | 0.398 | 0.434 | 1 | 3 |
| Ttc19 | -0.2908 | 0.623 | 0.739 | 1 | 2 |
| Saraf | -0.291 | 0.856 | 0.941 | 1 | 5 |
| Nck1 | -0.291 | 0.059 | 0.321 | 1 | 10 |
| Arpp21 | -0.2911 | 0.194 | 0.401 | 0.0747 | 4 |
| Pvrl3 | -0.2912 | 0.237 | 0.389 | 0.0684 | 1 |
| mt-Nd2 | -0.2912 | 1 | 0.999 | 1 | 7 |
| Mcfd2 | -0.2912 | 0.628 | 0.786 | 0.053 | 7 |
| Pcdh9 | -0.2913 | 0.421 | 0.636 | 0.0216 | 1 |
| Surf1 | -0.2913 | 0.187 | 0.497 | 1 | 3 |
| Hbegf | -0.2917 | 0.113 | 0.221 | 1 | 8 |
| Igsf8 | -0.2917 | 0.718 | 0.827 | 1 | 7 |
| Lsm5 | -0.2918 | 0.059 | 0.383 | 1 | 10 |
| Ckb | -0.2919 | 0.98 | 0.993 | 0.0005 | 1 |
| Rab28 | -0.2919 | 0.421 | 0.588 | 0.0003 | 1 |
| Cdip1 | -0.292 | 0.154 | 0.501 | 1 | 3 |
| Tmeff1 | -0.2921 | 0.031 | 0.269 | 0.0716 | 6 |
| Rock2 | -0.2922 | 0.394 | 0.635 | 1 | 8 |
| Clip3 | -0.2922 | 0.195 | 0.499 | 1 | 3 |
| Mrpl28 | -0.2923 | 0.228 | 0.553 | 1 | 3 |
| Emc9 | -0.2923 | 0.295 | 0.477 | 1 | 7 |
| Fbxw7 | -0.2925 | 0.138 | 0.417 | 0.9872 | 3 |
| Tspan3 | -0.2925 | 0.966 | 0.978 | 1 | 2 |
| Aplp2 | -0.2926 | 0.641 | 0.738 | 1 | 7 |
| Rph3a | -0.2927 | 0.355 | 0.627 | 0.0002 | 1 |
| C77370 | -0.2927 | 0.167 | 0.38 | 1 | 7 |
| Syn2 | -0.2927 | 0.908 | 0.965 | 0.0004 | 1 |
| Msmo1 | -0.2929 | 0.324 | 0.483 | 1 | 8 |
| Pdzrn4 | -0.2929 | 0.089 | 0.306 | 0.0203 | 3 |
| Nrcam | -0.2929 | 0.243 | 0.445 | 0.0011 | 1 |
| B4galt6 | -0.2931 | 0.454 | 0.606 | 0.0273 | 5 |
| Slc24a2 | -0.2935 | 0.141 | 0.338 | 0.3926 | 8 |
| Crtac1 | -0.2935 | 0.096 | 0.23 | 0.0025 | 2 |
| Rpl36 | -0.2936 | 0.995 | 0.992 | 0.0001 | 0 |
| Ankrd54 | -0.2936 | 0.059 | 0.309 | 1 | 10 |
| Polr2i | -0.2936 | 0.285 | 0.602 | 1 | 3 |
| Polr3h | -0.2936 | 0.118 | 0.36 | 1 | 10 |
| Cuedc1 | -0.2937 | 0.294 | 0.448 | 1 | 10 |
| Wsb1 | -0.2938 | 0.3 | 0.505 | 1 | 9 |
| Ubqln2 | -0.294 | 0.367 | 0.61 | 1 | 9 |
| Rab3gap1 | -0.2941 | 0.351 | 0.532 | 1 | 6 |
| Gnai2 | -0.2942 | 0.867 | 0.975 | 1 | 9 |

| | | | | | |
|---|---|---|---|---|---|
| Cxx1a | -0.2942 | 0.99 | 0.992 | 0.0003 | 5 |
| Nsg2 | -0.2943 | 0.872 | 0.955 | 1 | 7 |
| Prmt2 | -0.2943 | 0.733 | 0.922 | 1 | 9 |
| Atp5g2 | -0.2944 | 0.945 | 0.978 | 0.0001 | 0 |
| Nrxn1 | -0.2944 | 0.856 | 0.929 | 1 | 5 |
| Hnrnpa3 | -0.2946 | 0.633 | 0.882 | 1 | 9 |
| Dpysl2 | -0.2946 | 0.833 | 0.968 | 1 | 9 |
| Ost4 | -0.2946 | 0.382 | 0.601 | 2.14E-06 | 1 |
| Adgrl3 | -0.2947 | 0.423 | 0.514 | 1 | 5 |
| Ctnna2 | -0.2947 | 0.139 | 0.38 | 0.0005 | 4 |
| Ldha | -0.2949 | 0.567 | 0.779 | 1 | 5 |
| Fundc1 | -0.2953 | 0.098 | 0.396 | 1 | 3 |
| Wbp1 | -0.2954 | 0.118 | 0.36 | 1 | 10 |
| Elmo1 | -0.2955 | 0.141 | 0.281 | 1 | 7 |
| Fhad1 | -0.2957 | 0.026 | 0.235 | 0.0021 | 7 |
| Adgrb2 | -0.2958 | 0.296 | 0.456 | 1 | 8 |
| Atp1b2 | -0.2958 | 0.493 | 0.585 | 1 | 8 |
| Tomm34 | -0.2959 | 0.244 | 0.56 | 1 | 3 |
| Mapk8ip3 | -0.2961 | 0.538 | 0.744 | 0.3436 | 7 |
| Gabra3 | -0.2961 | 0.103 | 0.345 | 0.258 | 7 |
| Ttc14 | -0.2962 | 0.493 | 0.757 | 1 | 8 |
| Tm9sf4 | -0.2963 | 0.466 | 0.609 | 0.0256 | 2 |
| Pianp | -0.2963 | 0.836 | 0.933 | 0.9824 | 2 |
| Fhad1 | -0.2964 | 0.033 | 0.244 | 1 | 3 |
| Nell1 | -0.2964 | 0.127 | 0.309 | 1 | 8 |
| Npm1 | -0.2966 | 0.967 | 0.981 | 1 | 3 |
| Zic1 | -0.2967 | 0 | 0.213 | 1.20E-05 | 8 |
| Tgfa | -0.2967 | 0.05 | 0.286 | 7.65E-11 | 0 |
| Nptn | -0.2968 | 0.722 | 0.836 | 0.0007 | 6 |
| B4galt6 | -0.2969 | 0.451 | 0.602 | 1 | 8 |
| Cbx3 | -0.2969 | 0.835 | 0.843 | 1 | 0 |
| Grin3a | -0.2969 | 0.179 | 0.352 | 1 | 7 |
| Abhd8 | -0.297 | 0.8 | 0.878 | 1 | 9 |
| Pdzrn4 | -0.2972 | 0.144 | 0.303 | 3.25E-08 | 2 |
| Csmd3 | -0.2973 | 0.124 | 0.327 | 3.04E-05 | 5 |
| Huwe1 | -0.2977 | 0.535 | 0.726 | 1 | 8 |
| Rmst | -0.2977 | 0.34 | 0.483 | 1 | 5 |
| Slc24a5 | -0.2978 | 0.352 | 0.579 | 1 | 8 |
| Cbln4 | -0.298 | 0.033 | 0.228 | 1 | 9 |
| Nsg2 | -0.298 | 0.897 | 0.954 | 1 | 5 |
| Txndc17 | -0.2981 | 0.504 | 0.711 | 1 | 3 |
| Camkv | -0.2982 | 0.321 | 0.496 | 1 | 7 |

| | | | | | |
|---|---|---|---|---|---|
| Ddost | -0.2983 | 0.658 | 0.729 | 1 | 2 |
| Cops2 | -0.2983 | 0.122 | 0.434 | 1 | 3 |
| Kif1a | -0.2986 | 0.907 | 0.918 | 1 | 5 |
| H3f3b | -0.2986 | 0.995 | 0.999 | 5.16E-06 | 0 |
| Sdcbp | -0.2986 | 0.267 | 0.476 | 1 | 9 |
| Tmem70 | -0.2988 | 0.176 | 0.401 | 1 | 10 |
| Tmem50a | -0.2988 | 0.641 | 0.844 | 0.002 | 7 |
| Npas4 | -0.2989 | 0.137 | 0.252 | 1 | 2 |
| Opcml | -0.2991 | 0.557 | 0.688 | 0.0046 | 6 |
| Usp34 | -0.2991 | 0.718 | 0.831 | 1 | 7 |
| Tspan5 | -0.2991 | 0.205 | 0.391 | 1 | 7 |
| Tcf4 | -0.2991 | 0.141 | 0.317 | 1 | 7 |
| Sh3kbp1 | -0.2993 | 0.178 | 0.404 | 3.18E-06 | 1 |
| Cntn5 | -0.2993 | 0.021 | 0.18 | 0.0001 | 5 |
| Odc1 | -0.2993 | 0.451 | 0.581 | 0.8952 | 8 |
| Parm1 | -0.2994 | 0.44 | 0.619 | 2.39E-05 | 0 |
| Nfe2l1 | -0.2994 | 0.549 | 0.789 | 1 | 8 |
| Ap2a2 | -0.2995 | 0.629 | 0.744 | 0.3232 | 5 |
| A230065I | -0.2996 | 0 | 0.174 | 9.84E-07 | 5 |
| Chchd10 | -0.2996 | 0.889 | 0.93 | 0.4696 | 4 |
| Fyn | -0.2997 | 0.154 | 0.393 | 0.2629 | 7 |
| Ppp3ca | -0.2998 | 0.719 | 0.805 | 0.0113 | 2 |
| Atp6ap2 | -0.2999 | 0.731 | 0.843 | 1 | 7 |
| Gabrg2 | -0.2999 | 0.546 | 0.686 | 1 | 5 |
| Ngb | -0.3 | 0.805 | 0.775 | 1 | 3 |
| Sepw1 | -0.3 | 0.995 | 0.996 | 0.0002 | 0 |
| Egr1 | -0.3002 | 0.24 | 0.344 | 1 | 0 |
| Myef2 | -0.3004 | 0.471 | 0.64 | 1 | 10 |
| Tob1 | -0.3005 | 0.233 | 0.421 | 1 | 9 |
| Ptms | -0.3006 | 0.955 | 0.975 | 0.0003 | 0 |
| Cacna1d | -0.3007 | 0.352 | 0.545 | 0.1372 | 4 |
| Bhlhe41 | -0.3007 | 0.194 | 0.337 | 1 | 4 |
| Tcf4 | -0.3007 | 0.186 | 0.316 | 0.1065 | 5 |
| Zwint | -0.3008 | 1 | 0.996 | 0.0004 | 8 |
| Sdk2 | -0.301 | 0 | 0.248 | 1 | 10 |
| Brinp2 | -0.301 | 0.115 | 0.339 | 0.3105 | 7 |
| Mrpl48 | -0.3011 | 0.333 | 0.637 | 1 | 3 |
| Slc39a10 | -0.3011 | 0.333 | 0.507 | 1 | 7 |
| Fosb | -0.3011 | 0.09 | 0.198 | 0.5036 | 0 |
| H3f3b | -0.3011 | 1 | 0.998 | 0.0063 | 1 |
| Rwdd1 | -0.3012 | 0.602 | 0.746 | 1 | 3 |
| Hdhd2 | -0.3013 | 0.176 | 0.438 | 1 | 10 |

| | | | | | |
|---|---|---|---|---|---|
| Ptma | -0.3014 | 0.985 | 0.987 | 5.12E-05 | 0 |
| Gpx3 | -0.3014 | 0.268 | 0.442 | 1 | 6 |
| Adam22 | -0.3015 | 0.41 | 0.548 | 1 | 7 |
| Kif1b | -0.3015 | 0.808 | 0.882 | 1 | 7 |
| Akap11 | -0.3015 | 0.685 | 0.793 | 0.0004 | 2 |
| Bmpr2 | -0.3017 | 0.144 | 0.394 | 1 | 6 |
| Tkt | -0.3017 | 0.462 | 0.647 | 0.1131 | 7 |
| Isca1 | -0.3017 | 0.588 | 0.666 | 1 | 10 |
| Camk2b | -0.3018 | 0.63 | 0.812 | 5.44E-05 | 2 |
| Gda | -0.3018 | 0.507 | 0.65 | 0.1341 | 1 |
| Npy1r | -0.302 | 0.033 | 0.252 | 1 | 9 |
| Faim | -0.3021 | 0.118 | 0.373 | 1 | 10 |
| Srek1ip1 | -0.3021 | 0.118 | 0.363 | 1 | 10 |
| Enc1 | -0.3021 | 0.093 | 0.287 | 1.09E-05 | 5 |
| Tm2d1 | -0.3021 | 0.412 | 0.72 | 1 | 10 |
| Ctnnb1 | -0.3021 | 0.463 | 0.735 | 0.0002 | 4 |
| Pdap1 | -0.3023 | 0.675 | 0.789 | 1 | 3 |
| Eif5a | -0.3024 | 0.992 | 0.988 | 1 | 3 |
| Fgf13 | -0.3025 | 0.059 | 0.278 | 1 | 10 |
| Gm26782 | -0.3025 | 0.059 | 0.302 | 1 | 10 |
| Camk2a | -0.3025 | 0.536 | 0.732 | 0.168 | 5 |
| Dlk1 | -0.3025 | 0.645 | 0.717 | 1 | 1 |
| Tmem9b | -0.3025 | 0.165 | 0.436 | 0.2217 | 6 |
| Rpl4 | -0.3025 | 0.927 | 0.889 | 1 | 3 |
| Cntn5 | -0.3027 | 0.013 | 0.178 | 0.0822 | 7 |
| Ubb | -0.3027 | 1 | 1 | 0.0395 | 0 |
| Rxrg | -0.3027 | 0 | 0.158 | 0.0054 | 7 |
| H2afx | -0.3028 | 0.353 | 0.541 | 1 | 10 |
| Ralyl | -0.3028 | 0.394 | 0.548 | 0.0548 | 8 |
| Adam22 | -0.3028 | 0.38 | 0.549 | 1 | 8 |
| Pgap1 | -0.3029 | 0.31 | 0.548 | 1 | 8 |
| Atp6v1b2 | -0.3032 | 0.784 | 0.869 | 0.2966 | 6 |
| Kdsr | -0.3034 | 0.133 | 0.375 | 1 | 9 |
| Susd4 | -0.3034 | 0.24 | 0.385 | 1.16E-08 | 2 |
| Rpl34 | -0.3034 | 0.935 | 0.982 | 1.94E-05 | 0 |
| Zfp467 | -0.3037 | 0.267 | 0.428 | 1 | 9 |
| Cux1 | -0.3037 | 0.25 | 0.455 | 0.0038 | 4 |
| Tmed10 | -0.3037 | 0.733 | 0.848 | 1 | 2 |
| Kdsr | -0.3038 | 0.197 | 0.38 | 0.6477 | 8 |
| Nrcam | -0.3038 | 0.225 | 0.43 | 1 | 8 |
| Ncdn | -0.304 | 0.513 | 0.635 | 0.3395 | 1 |
| Ybx1 | -0.3042 | 0.57 | 0.753 | 7.67E-06 | 0 |

| | | | | | |
|---|---|---|---|---|---|
| Cisd2 | -0.3042 | 0.479 | 0.68 | 0.9321 | 8 |
| Mef2c | -0.3042 | 0.067 | 0.234 | 1 | 9 |
| Ube2ql1 | -0.3043 | 0.191 | 0.418 | 6.43E-07 | 1 |
| Hras | -0.3043 | 0.894 | 0.929 | 1 | 3 |
| Tspan5 | -0.3044 | 0.169 | 0.392 | 0.0554 | 8 |
| Dctn1 | -0.3044 | 0.596 | 0.737 | 0.0053 | 2 |
| Xpr1 | -0.3044 | 0.352 | 0.563 | 0.0251 | 4 |
| Zdhhc20 | -0.3045 | 0.175 | 0.444 | 0.5929 | 6 |
| Ubl3 | -0.3046 | 0.163 | 0.43 | 1 | 3 |
| Nt5dc3 | -0.3047 | 0.167 | 0.339 | 1 | 9 |
| Cited1 | -0.3047 | 0.062 | 0.178 | 2.82E-05 | 2 |
| 9530068I | -0.3048 | 0.247 | 0.459 | 1 | 6 |
| Nhp2 | -0.3049 | 0.049 | 0.352 | 1 | 3 |
| Sulf1 | -0.305 | 0 | 0.206 | 4.61E-17 | 2 |
| Tctex1d2 | -0.305 | 0.059 | 0.329 | 1 | 10 |
| Vta1 | -0.3051 | 0.122 | 0.429 | 1 | 3 |
| Dek | -0.3051 | 0.235 | 0.437 | 1 | 10 |
| Abr | -0.3052 | 0.462 | 0.725 | 0.1309 | 7 |
| Slc24a5 | -0.3052 | 0.474 | 0.573 | 1 | 6 |
| Sh3gl2 | -0.3052 | 0.282 | 0.5 | 0.0404 | 8 |
| Rpl36a | -0.3053 | 0.895 | 0.937 | 0.0104 | 0 |
| Smap2 | -0.3053 | 0.22 | 0.568 | 1 | 3 |
| Tenm1 | -0.3053 | 0.329 | 0.392 | 0.0008 | 2 |
| B630019I | -0.3053 | 0.722 | 0.827 | 0.2481 | 4 |
| Hpcal4 | -0.3054 | 0.392 | 0.561 | 0.0147 | 5 |
| Brinp3 | -0.3054 | 0.296 | 0.431 | 0.6003 | 1 |
| Baiap2 | -0.3054 | 0.205 | 0.398 | 1 | 7 |
| Tcf12 | -0.3054 | 0.222 | 0.408 | 0.0508 | 4 |
| Lsm4 | -0.3056 | 0.626 | 0.771 | 1 | 3 |
| Wrb | -0.3057 | 0.529 | 0.691 | 1 | 10 |
| Spcs1 | -0.3057 | 0.936 | 0.957 | 0.0613 | 7 |
| Disp2 | -0.306 | 0.731 | 0.843 | 1 | 7 |
| Scn1b | -0.3061 | 0.479 | 0.695 | 3.97E-06 | 2 |
| Jakmip1 | -0.3061 | 0.269 | 0.487 | 0.8421 | 7 |
| Hmgb3 | -0.3062 | 0.433 | 0.597 | 1 | 9 |
| Pbx1 | -0.3064 | 0.73 | 0.77 | 0.0215 | 0 |
| Pcdh11x | -0.3065 | 0.014 | 0.184 | 0.0227 | 8 |
| 1700001I | -0.3067 | 0.175 | 0.45 | 6.42E-08 | 0 |
| Bbx | -0.3069 | 0.059 | 0.316 | 1 | 10 |
| Ldha | -0.307 | 0.797 | 0.756 | 0.1414 | 3 |
| Baiap2 | -0.307 | 0.206 | 0.401 | 0.0001 | 5 |
| Kmt2e | -0.3072 | 0.529 | 0.7 | 1 | 10 |

| | | | | | |
|---|---|---|---|---|---|
| Hs6st2 | -0.3074 | 0.296 | 0.405 | 1 | 8 |
| Fam213a | -0.3075 | 0.353 | 0.575 | 1 | 10 |
| Cpne5 | -0.3076 | 0.081 | 0.31 | 1 | 3 |
| Pnisr | -0.3077 | 0.62 | 0.8 | 1 | 8 |
| Pnmal2 | -0.3078 | 0.981 | 0.994 | 0.0003 | 4 |
| Sv2a | -0.3078 | 0.923 | 0.925 | 1 | 7 |
| B4gat1 | -0.3079 | 0.534 | 0.698 | 0.7733 | 2 |
| Ssr1 | -0.3083 | 0.654 | 0.784 | 0.6698 | 7 |
| Tsc22d1 | -0.3084 | 0.835 | 0.865 | 0.0627 | 6 |
| Rpl8 | -0.3085 | 1 | 0.998 | 1 | 3 |
| Sorcs3 | -0.3087 | 0.01 | 0.208 | 1.48E-07 | 5 |
| Tgoln1 | -0.3088 | 0.678 | 0.792 | 1 | 2 |
| Dpp10 | -0.3089 | 0.115 | 0.312 | 1 | 7 |
| Atg101 | -0.309 | 0 | 0.281 | 1 | 10 |
| Mpped2 | -0.3091 | 0.2 | 0.39 | 1 | 9 |
| Prkcsh | -0.3091 | 0.233 | 0.429 | 1 | 9 |
| Tmie | -0.3092 | 0.09 | 0.344 | 0.014 | 7 |
| Ets2 | -0.3093 | 0.072 | 0.293 | 2.16E-07 | 5 |
| Stmn4 | -0.3094 | 0.678 | 0.806 | 1 | 2 |
| Pigt | -0.3095 | 0.231 | 0.453 | 1 | 7 |
| Ppa1 | -0.3096 | 0.308 | 0.488 | 0.1236 | 7 |
| Cnpy3 | -0.3096 | 0.244 | 0.5 | 0.2762 | 7 |
| Stx1b | -0.3096 | 0.187 | 0.501 | 1 | 3 |
| Sec11c | -0.3097 | 0.732 | 0.88 | 1 | 8 |
| Ephb1 | -0.31 | 0.176 | 0.383 | 0.0136 | 4 |
| Calb1 | -0.3101 | 0.724 | 0.769 | 1 | 1 |
| Crabp1 | -0.3102 | 0.007 | 0.03 | 1 | 2 |
| Cirbp | -0.3103 | 0.935 | 0.98 | 1.51E-09 | 0 |
| Arxes2 | -0.3103 | 0.763 | 0.871 | 1 | 5 |
| AI593442 | -0.3104 | 0.141 | 0.336 | 0.1414 | 8 |
| Mgat4c | -0.3104 | 0.062 | 0.24 | 0.0004 | 5 |
| Ndufa13 | -0.3105 | 0.979 | 0.984 | 1 | 2 |
| Cx3cl1 | -0.3105 | 0.474 | 0.595 | 1 | 5 |
| Ift27 | -0.3105 | 0.041 | 0.331 | 0.5059 | 3 |
| Dhx30 | -0.3106 | 0.321 | 0.522 | 1 | 7 |
| Psme1 | -0.3106 | 0.35 | 0.629 | 1 | 3 |
| Cacna1g | -0.3107 | 0.014 | 0.276 | 5.36E-06 | 8 |
| Stmn3 | -0.3108 | 1 | 1 | 0.334 | 2 |
| Bhlhe41 | -0.3109 | 0.183 | 0.333 | 1 | 8 |
| Smarca2 | -0.3113 | 0.59 | 0.785 | 0.0385 | 7 |
| Kif5c | -0.3113 | 0.744 | 0.812 | 1 | 7 |
| Arhgap6 | -0.3115 | 0.545 | 0.619 | 0.1155 | 0 |

| | | | | | |
|---|---|---|---|---|---|
| Rrad | -0.3115 | 0.093 | 0.177 | 1 | 5 |
| Dad1 | -0.3115 | 0.911 | 0.971 | 1 | 2 |
| Polr2f | -0.3115 | 0.577 | 0.742 | 1 | 3 |
| Astn1 | -0.3117 | 0.619 | 0.733 | 1 | 5 |
| Snhg11 | -0.3117 | 1 | 0.999 | 0.045 | 6 |
| Ube3a | -0.3118 | 0.897 | 0.958 | 1 | 7 |
| Rbfox1 | -0.3119 | 0.041 | 0.226 | 8.11E-06 | 5 |
| Rps18 | -0.3119 | 0.89 | 0.977 | 4.15E-08 | 0 |
| Igsf8 | -0.312 | 0.742 | 0.827 | 1 | 5 |
| Gm15261 | -0.3121 | 0.014 | 0.24 | 0.0002 | 8 |
| mt-Nd5 | -0.3121 | 0.949 | 0.981 | 1 | 7 |
| 15000090 | -0.3121 | 0.325 | 0.621 | 1 | 3 |
| Atp6v1g2 | -0.3121 | 0.859 | 0.915 | 1 | 7 |
| Rps23 | -0.3122 | 0.925 | 0.982 | 2.93E-06 | 0 |
| Oprk1 | -0.3123 | 0.093 | 0.263 | 0.0169 | 5 |
| Ddc | -0.3123 | 0.1 | 0.239 | 1 | 9 |
| Aurkaip1 | -0.3124 | 0.772 | 0.843 | 1 | 3 |
| Pdcd5 | -0.3126 | 0.878 | 0.897 | 0.4054 | 3 |
| Tox | -0.3128 | 0.103 | 0.344 | 0.0688 | 7 |
| Cntfr | -0.3128 | 0.118 | 0.345 | 1 | 10 |
| Tmem176 | -0.3128 | 0.333 | 0.69 | 1 | 9 |
| Rpl39 | -0.3129 | 0.96 | 0.979 | 1.33E-06 | 0 |
| Rap1gap | -0.3129 | 0.171 | 0.446 | 0.0151 | 3 |
| Aatk | -0.3132 | 0.487 | 0.613 | 1 | 7 |
| Mrpl17 | -0.3133 | 0.487 | 0.689 | 6.00E-07 | 1 |
| Nrsn1 | -0.3133 | 0.904 | 0.966 | 0.0089 | 2 |
| Eif1a | -0.3134 | 0.235 | 0.57 | 1 | 10 |
| Ctxn2 | -0.3134 | 0.368 | 0.592 | 6.87E-06 | 1 |
| Ctsd | -0.3134 | 0.548 | 0.68 | 1 | 2 |
| Igf1r | -0.3136 | 0.046 | 0.332 | 1.39E-08 | 4 |
| Stoml1 | -0.3137 | 0.154 | 0.395 | 0.0228 | 7 |
| Gpm6a | -0.3138 | 0.966 | 0.966 | 1 | 2 |
| Mrpl27 | -0.3141 | 0.325 | 0.631 | 1 | 3 |
| Cntnap5a | -0.3142 | 0.171 | 0.335 | 1.09E-06 | 2 |
| Bcl7b | -0.3143 | 0.171 | 0.503 | 1 | 3 |
| Nt5dc3 | -0.3145 | 0.151 | 0.363 | 1.13E-05 | 1 |
| Tmem9 | -0.3149 | 0.485 | 0.71 | 0.0933 | 5 |
| Disp2 | -0.315 | 0.704 | 0.844 | 1 | 8 |
| Rpn1 | -0.315 | 0.521 | 0.665 | 0.0652 | 2 |
| Hook1 | -0.315 | 0.133 | 0.389 | 1 | 9 |
| Ncdn | -0.3152 | 0.465 | 0.629 | 1 | 8 |
| Syp | -0.3152 | 0.936 | 0.936 | 1 | 7 |

| | | | | | |
|---|---|---|---|---|---|
| Ociad1 | -0.3153 | 0.814 | 0.918 | 0.8148 | 5 |
| Gabarapl1 | -0.3153 | 0.967 | 0.949 | 1 | 3 |
| Elmo1 | -0.3154 | 0.137 | 0.292 | 4.11E-10 | 2 |
| Pfdn2 | -0.3154 | 0.967 | 0.973 | 0.8082 | 3 |
| Dbpht2 | -0.3155 | 0.1 | 0.328 | 0.9978 | 9 |
| Serpine2 | -0.3156 | 0.103 | 0.287 | 1 | 7 |
| Cebpzos | -0.3156 | 0.309 | 0.607 | 1 | 3 |
| Dgkg | -0.3156 | 0.041 | 0.295 | 9.88E-09 | 5 |
| Pnp | -0.3156 | 0.125 | 0.361 | 2.68E-07 | 0 |
| Phlda3 | -0.3158 | 0.098 | 0.362 | 1 | 3 |
| B3galnt1 | -0.3158 | 0.176 | 0.391 | 1 | 10 |
| Rpgrip1 | -0.3158 | 0.176 | 0.384 | 1 | 10 |
| Ucp2 | -0.3162 | 0.122 | 0.355 | 1 | 3 |
| Sez6l | -0.3162 | 0.514 | 0.619 | 0.0246 | 2 |
| Rftn1 | -0.3163 | 0.2 | 0.396 | 1 | 9 |
| Cdk14 | -0.3163 | 0.098 | 0.335 | 0.6824 | 3 |
| Tmbim6 | -0.3163 | 0.533 | 0.81 | 1 | 9 |
| Slc16a11 | -0.3164 | 0.09 | 0.295 | 0.6464 | 7 |
| Tbca | -0.3165 | 0.813 | 0.894 | 1 | 3 |
| Gabra5 | -0.3167 | 0.062 | 0.271 | 0.8418 | 6 |
| Nrxn2 | -0.3171 | 0.93 | 0.966 | 1 | 8 |
| Ap1s2 | -0.3173 | 0.426 | 0.63 | 0.1195 | 4 |
| Npy1r | -0.3174 | 0.028 | 0.26 | 0.0007 | 8 |
| Fam69b | -0.3174 | 0.623 | 0.768 | 0.483 | 2 |
| Scg5 | -0.3176 | 0.972 | 0.978 | 1 | 8 |
| Cbx3 | -0.3178 | 0.757 | 0.855 | 0.0035 | 1 |
| Fosb | -0.3178 | 0.093 | 0.187 | 1 | 6 |
| Egfl7 | -0.3179 | 0.128 | 0.357 | 0.0964 | 7 |
| Tcea2 | -0.3181 | 0.128 | 0.391 | 0.0167 | 7 |
| Tpd52l1 | -0.3181 | 0.315 | 0.487 | 0.2363 | 4 |
| Lrfn5 | -0.3184 | 0.278 | 0.494 | 0.0014 | 5 |
| Nol7 | -0.3184 | 0.398 | 0.652 | 1 | 3 |
| Arf3 | -0.3184 | 0.742 | 0.835 | 0.2854 | 5 |
| Tia1 | -0.3188 | 0.641 | 0.751 | 1 | 7 |
| Ece2 | -0.3188 | 0.186 | 0.457 | 0.038 | 6 |
| Pdia6 | -0.3188 | 0.795 | 0.837 | 1 | 7 |
| Crim1 | -0.3189 | 0 | 0.224 | 6.75E-05 | 7 |
| Celf2 | -0.3189 | 0.505 | 0.65 | 0.0782 | 5 |
| mt-Atp8 | -0.3192 | 0.887 | 0.966 | 1 | 8 |
| 2610524I | -0.3193 | 0.308 | 0.537 | 0.327 | 7 |
| Itfg1 | -0.3194 | 0.5 | 0.648 | 1 | 7 |
| Ctsl | -0.3194 | 0.664 | 0.799 | 1 | 2 |

| | | | | | |
|---|---|---|---|---|---|
| Grina | -0.3196 | 0.795 | 0.872 | 0.3758 | 7 |
| Gadd45gi | -0.3198 | 0.163 | 0.543 | 1 | 3 |
| Gabrg3 | -0.32 | 0.179 | 0.427 | 0.003 | 3 |
| Fis1 | -0.3202 | 0.915 | 0.937 | 0.0001 | 0 |
| 36324510 | -0.3203 | 0.099 | 0.317 | 7.84E-07 | 1 |
| Scg3 | -0.3203 | 0.637 | 0.789 | 1 | 2 |
| Scn1a | -0.3203 | 0.116 | 0.306 | 1.06E-12 | 2 |
| Ddost | -0.3205 | 0.577 | 0.73 | 0.0316 | 7 |
| Rpl27 | -0.3207 | 0.83 | 0.938 | 7.85E-07 | 0 |
| Pclo | -0.3207 | 0.678 | 0.781 | 0.0454 | 2 |
| Prr13 | -0.3208 | 0.431 | 0.699 | 0.0549 | 3 |
| Skp1a | -0.3208 | 0.902 | 0.904 | 1 | 3 |
| Rpn1 | -0.3208 | 0.493 | 0.656 | 1 | 8 |
| Cpne7 | -0.3209 | 0.008 | 0.218 | 2.85E-07 | 3 |
| Gria3 | -0.3209 | 0.033 | 0.224 | 0.0002 | 3 |
| Ccdc124 | -0.321 | 0.244 | 0.617 | 0.5204 | 3 |
| Cpne4 | -0.321 | 0 | 0.197 | 7.78E-05 | 7 |
| Canx | -0.3212 | 0.846 | 0.926 | 0.7012 | 7 |
| Sash1 | -0.3212 | 0.042 | 0.295 | 2.82E-05 | 8 |
| Rasl11b | -0.3214 | 0.04 | 0.199 | 5.09E-05 | 0 |
| Prune2 | -0.3214 | 0.268 | 0.476 | 1 | 6 |
| Tcf25 | -0.3214 | 0.966 | 0.986 | 2.13E-05 | 2 |
| Mesdc2 | -0.3214 | 0.616 | 0.776 | 1 | 2 |
| Gabra3 | -0.3215 | 0.134 | 0.346 | 2.14E-05 | 5 |
| B3gat2 | -0.3217 | 0.333 | 0.464 | 1 | 9 |
| Med29 | -0.3217 | 0.118 | 0.378 | 1 | 10 |
| Itpa | -0.3217 | 0.057 | 0.384 | 1 | 3 |
| Cthrc1 | -0.3218 | 0.259 | 0.371 | 1 | 4 |
| Gria3 | -0.3219 | 0.042 | 0.214 | 0.084 | 8 |
| Ptpn5 | -0.322 | 0.13 | 0.376 | 0.0062 | 3 |
| Snhg20 | -0.3221 | 0.577 | 0.847 | 1 | 8 |
| Slc24a2 | -0.3222 | 0.12 | 0.347 | 6.08E-05 | 4 |
| Kcnip4 | -0.3222 | 0.038 | 0.241 | 0.0213 | 7 |
| Dlx6 | -0.3222 | 0 | 0.27 | 1 | 10 |
| Psd3 | -0.3223 | 0.309 | 0.457 | 0.0014 | 5 |
| 9530068I | -0.3223 | 0.2 | 0.447 | 1 | 9 |
| Ywhaz | -0.3224 | 0.976 | 0.972 | 1 | 3 |
| Tagln3 | -0.3224 | 0.904 | 0.953 | 0.0036 | 2 |
| mt-Co3 | -0.3227 | 1 | 1 | 0.1885 | 7 |
| Rps2 | -0.3228 | 0.97 | 0.997 | 2.19E-12 | 0 |
| Chl1 | -0.3228 | 0.404 | 0.486 | 0.0083 | 2 |
| Rorb | -0.323 | 0.521 | 0.693 | 0.0184 | 2 |

| | | | | | |
|---|---|---|---|---|---|
| Atp8a1 | -0.323 | 0.705 | 0.802 | 0.1016 | 2 |
| Per3 | -0.323 | 0.48 | 0.603 | 0.4224 | 1 |
| Cx3cl1 | -0.3231 | 0.452 | 0.604 | 0.0006 | 2 |
| Maged1 | -0.3234 | 0.948 | 0.991 | 1 | 5 |
| H2afz | -0.3235 | 0.976 | 0.98 | 0.0749 | 3 |
| Ube2ql1 | -0.3236 | 0.169 | 0.402 | 0.0119 | 8 |
| Nxph1 | -0.3237 | 0.282 | 0.387 | 1 | 8 |
| Map1lc3a | -0.3237 | 0.901 | 0.976 | 0.0012 | 8 |
| Plxnc1 | -0.3237 | 0.254 | 0.442 | 0.754 | 8 |
| Vstm2a | -0.3237 | 0.062 | 0.287 | 2.89E-07 | 5 |
| Araf | -0.3237 | 0.933 | 0.921 | 1 | 9 |
| Isl1 | -0.3238 | 0.093 | 0.261 | 1 | 6 |
| Smim7 | -0.3238 | 0.353 | 0.527 | 1 | 10 |
| Isca2 | -0.324 | 0.294 | 0.561 | 1 | 10 |
| Atp6ap1 | -0.3241 | 0.615 | 0.805 | 0.0055 | 7 |
| Rpl3 | -0.3241 | 1 | 1 | 5.05E-07 | 0 |
| Cited1 | -0.3242 | 0.038 | 0.172 | 0.4964 | 7 |
| Gnb1 | -0.3242 | 0.917 | 0.942 | 0.1109 | 4 |
| Baiap3 | -0.3243 | 0.856 | 0.912 | 1 | 2 |
| Mrpl30 | -0.3244 | 0.472 | 0.695 | 1 | 3 |
| Lrfn5 | -0.3244 | 0.346 | 0.485 | 1 | 7 |
| Cwc15 | -0.3246 | 0.276 | 0.613 | 1 | 3 |
| Ppp3ca | -0.3248 | 0.846 | 0.787 | 0.6782 | 3 |
| Aldoa | -0.3248 | 1 | 1 | 0.0011 | 4 |
| 1500011 | -0.3248 | 0.821 | 0.963 | 2.53E-06 | 7 |
| Ppp1r9a | -0.3248 | 0.449 | 0.576 | 1 | 7 |
| Tgfa | -0.3249 | 0 | 0.251 | 0.2558 | 9 |
| Pde1a | -0.325 | 0.077 | 0.241 | 1 | 7 |
| Resp18 | -0.325 | 1 | 0.999 | 0.4283 | 6 |
| Ucp2 | -0.3251 | 0.128 | 0.345 | 0.4712 | 7 |
| Etnk1 | -0.3252 | 0.676 | 0.81 | 1 | 8 |
| Slc24a2 | -0.3252 | 0.138 | 0.355 | 2.74E-07 | 1 |
| Snrpd2 | -0.3253 | 0.846 | 0.91 | 1 | 3 |
| Necab2 | -0.3253 | 0.052 | 0.304 | 2.69E-07 | 5 |
| Adgra1 | -0.3254 | 0.26 | 0.434 | 5.47E-06 | 2 |
| Napb | -0.3256 | 0.397 | 0.484 | 1.54E-11 | 2 |
| Cbx3 | -0.3257 | 0.731 | 0.854 | 0.0595 | 4 |
| Kif1a | -0.3258 | 0.885 | 0.919 | 1 | 7 |
| Dlx1 | -0.3258 | 0.144 | 0.323 | 2.58E-11 | 2 |
| Echs1 | -0.326 | 0.235 | 0.446 | 1 | 10 |
| Cpne4 | -0.3261 | 0 | 0.201 | 4.94E-08 | 5 |
| Fkbp2 | -0.3261 | 0.876 | 0.934 | 1 | 5 |

| | | | | | |
|---|---|---|---|---|---|
| Cdc37 | -0.3261 | 0.423 | 0.68 | 1 | 3 |
| Faim2 | -0.3262 | 0.979 | 0.974 | 1 | 5 |
| Plcb1 | -0.3263 | 0.205 | 0.421 | 1 | 7 |
| Denr | -0.3264 | 0.65 | 0.794 | 1 | 3 |
| Cited1 | -0.3265 | 0.041 | 0.174 | 0.0187 | 5 |
| App | -0.3266 | 0.959 | 0.928 | 0.0004 | 6 |
| Fut9 | -0.3267 | 0.141 | 0.369 | 0.0134 | 8 |
| Dynll2 | -0.3268 | 0.919 | 0.918 | 1 | 3 |
| Atp8a1 | -0.327 | 0.692 | 0.796 | 0.618 | 7 |
| Mrpl42 | -0.3271 | 0.301 | 0.601 | 1 | 3 |
| Wrb | -0.3271 | 0.467 | 0.694 | 1 | 9 |
| B3gat2 | -0.3271 | 0.24 | 0.508 | 2.60E-07 | 0 |
| Chd3os | -0.3272 | 0.943 | 0.945 | 0.305 | 3 |
| Grm1 | -0.3272 | 0.07 | 0.289 | 0.0065 | 8 |
| Myo10 | -0.3272 | 0.099 | 0.324 | 0.0153 | 8 |
| Ptprs | -0.3273 | 0.795 | 0.853 | 1 | 7 |
| Ttc3 | -0.3273 | 1 | 1 | 0.0086 | 5 |
| Pde4b | -0.3274 | 0.039 | 0.316 | 2.11E-13 | 1 |
| Disp2 | -0.3275 | 0.767 | 0.846 | 1 | 2 |
| Tmem35 | -0.3275 | 0.296 | 0.387 | 1 | 8 |
| Terf2ip | -0.3276 | 0.294 | 0.485 | 1 | 10 |
| Ppp1r17 | -0.3277 | 0.13 | 0.347 | 6.13E-06 | 0 |
| Srsf10 | -0.3277 | 0.467 | 0.732 | 1 | 9 |
| Sox1 | -0.3278 | 0.185 | 0.39 | 0.002 | 4 |
| Mpped2 | -0.3279 | 0.224 | 0.411 | 8.38E-05 | 1 |
| Susd4 | -0.3279 | 0.186 | 0.384 | 0.0002 | 5 |
| Eno1 | -0.328 | 0.5 | 0.736 | 0.0136 | 7 |
| Hdhd2 | -0.3282 | 0.122 | 0.473 | 1 | 3 |
| Macf1 | -0.3283 | 0.692 | 0.815 | 0.9553 | 7 |
| Gt(ROSA)2 | -0.3285 | 0.333 | 0.421 | 1 | 9 |
| Acbd6 | -0.3287 | 0.171 | 0.544 | 1 | 3 |
| Fus | -0.3287 | 0.808 | 0.889 | 0.2221 | 7 |
| Cpne7 | -0.3288 | 0.007 | 0.223 | 2.25E-18 | 2 |
| Ppdpf | -0.3289 | 0.415 | 0.662 | 1 | 3 |
| Mirg | -0.329 | 0.155 | 0.395 | 0.0272 | 6 |
| Tenm2 | -0.329 | 0.171 | 0.414 | 2.39E-06 | 1 |
| Cspg5 | -0.329 | 0.773 | 0.914 | 0.0427 | 5 |
| Nr2f2 | -0.3291 | 0.295 | 0.43 | 1 | 7 |
| Lnp | -0.3292 | 0.352 | 0.536 | 1 | 8 |
| Borcs7 | -0.3292 | 0.187 | 0.537 | 1 | 3 |
| Bbip1 | -0.3295 | 0.353 | 0.577 | 1 | 10 |
| Celsr2 | -0.3298 | 0.295 | 0.518 | 1 | 7 |

| | | | | | |
|---|---|---|---|---|---|
| Dctn3 | -0.3299 | 0.886 | 0.897 | 1 | 3 |
| Tcf12 | -0.3299 | 0.167 | 0.396 | 1 | 9 |
| Adgrl1 | -0.3299 | 0.568 | 0.697 | 0.2116 | 2 |
| Oprk1 | -0.33 | 0.077 | 0.261 | 0.3738 | 7 |
| Stx16 | -0.3303 | 0.282 | 0.542 | 0.2424 | 7 |
| Map2k1 | -0.3303 | 0.41 | 0.566 | 0.9465 | 7 |
| Tgfa | -0.3304 | 0.02 | 0.279 | 5.24E-13 | 1 |
| Rasgrp1 | -0.3305 | 0.1 | 0.289 | 1 | 9 |
| Nrxn1 | -0.3305 | 0.859 | 0.927 | 0.3139 | 7 |
| Elmo1 | -0.3306 | 0.113 | 0.287 | 0.0004 | 5 |
| Gabrg3 | -0.3306 | 0.186 | 0.42 | 0.338 | 6 |
| Ssr1 | -0.3307 | 0.62 | 0.785 | 1 | 8 |
| Lysmd2 | -0.3307 | 0.358 | 0.662 | 0.6376 | 3 |
| Arl8a | -0.3307 | 0.551 | 0.737 | 0.2279 | 7 |
| Hspa5 | -0.3307 | 0.885 | 0.899 | 1 | 7 |
| Mien1 | -0.3308 | 0.407 | 0.664 | 1 | 3 |
| Adcyap1r | -0.3309 | 0.179 | 0.398 | 0.7663 | 7 |
| Inpp5f | -0.3309 | 0.474 | 0.711 | 0.0031 | 5 |
| Gabrb3 | -0.331 | 0.5 | 0.69 | 0.3424 | 7 |
| Glrx3 | -0.331 | 0.309 | 0.638 | 1 | 3 |
| Chmp4b | -0.3311 | 0.366 | 0.669 | 1 | 3 |
| Pde6d | -0.3312 | 0.13 | 0.458 | 1 | 3 |
| Rpsa | -0.3313 | 1 | 0.998 | 0.4335 | 3 |
| Ctxn1 | -0.3313 | 0.856 | 0.909 | 0.181 | 5 |
| Lrp1 | -0.3313 | 0.333 | 0.547 | 1 | 9 |
| Ptma | -0.3314 | 0.972 | 0.988 | 0.0431 | 4 |
| 36324510 | -0.3314 | 0.067 | 0.294 | 1 | 9 |
| Hnrnpab | -0.3314 | 0.301 | 0.624 | 0.4561 | 3 |
| Gabrq | -0.3315 | 0.021 | 0.231 | 1.12E-06 | 5 |
| Mrps18c | -0.3315 | 0.106 | 0.461 | 1 | 3 |
| St13 | -0.3315 | 0.633 | 0.813 | 1 | 9 |
| Ktn1 | -0.3315 | 0.63 | 0.799 | 1 | 2 |
| Tmx4 | -0.3316 | 0.962 | 0.986 | 0.1676 | 7 |
| Thap3 | -0.3316 | 0.118 | 0.39 | 1 | 10 |
| Scn3b | -0.332 | 0.299 | 0.479 | 0.0047 | 5 |
| Chga | -0.3322 | 0.763 | 0.732 | 0.0337 | 6 |
| Junb | -0.3323 | 0.245 | 0.356 | 1 | 0 |
| Dlx5 | -0.3323 | 0.056 | 0.343 | 5.25E-08 | 4 |
| Cnr1 | -0.3328 | 0.024 | 0.232 | 0.0014 | 3 |
| Abca2 | -0.3328 | 0.289 | 0.52 | 0.0004 | 5 |
| Efhd2 | -0.3329 | 0.106 | 0.391 | 0.3885 | 3 |
| Tmem255 | -0.333 | 0.436 | 0.565 | 1 | 7 |

| | | | | | |
|---|---|---|---|---|---|
| Thoc7 | -0.333 | 0.559 | 0.775 | 4.99E-08 | 1 |
| Tceal6 | -0.333 | 0.496 | 0.709 | 1 | 3 |
| Marcks | -0.3331 | 0.895 | 0.958 | 4.11E-08 | 0 |
| Plcb4 | -0.3331 | 0.346 | 0.526 | 1 | 7 |
| 2810428I | -0.3331 | 0.455 | 0.68 | 1 | 3 |
| Rftn1 | -0.3331 | 0.176 | 0.394 | 1 | 10 |
| Pcdh19 | -0.3332 | 0.37 | 0.543 | 0.1713 | 4 |
| Cpne4 | -0.3333 | 0 | 0.206 | 4.08E-06 | 3 |
| H2afv | -0.3335 | 0.625 | 0.766 | 9.64E-07 | 0 |
| Ephb1 | -0.3336 | 0.113 | 0.38 | 0.0163 | 8 |
| Sox1 | -0.3336 | 0.175 | 0.412 | 2.02E-07 | 0 |
| 50314391 | -0.3337 | 0.115 | 0.352 | 0.0522 | 7 |
| Mef2c | -0.3337 | 0.07 | 0.24 | 1 | 8 |
| Nkain3 | -0.3337 | 0.031 | 0.235 | 1.99E-06 | 5 |
| Ptprz1 | -0.3338 | 0.268 | 0.44 | 0.4909 | 6 |
| Elavl4 | -0.3339 | 0.146 | 0.442 | 1 | 3 |
| Fam3c | -0.3339 | 0.186 | 0.402 | 1.90E-05 | 5 |
| Arglu1 | -0.3342 | 0.733 | 0.855 | 1 | 9 |
| Podxl2 | -0.3342 | 0.974 | 0.987 | 0.2967 | 7 |
| Ajap1 | -0.3343 | 0.282 | 0.45 | 1 | 7 |
| Gabrq | -0.3344 | 0.013 | 0.228 | 0.0003 | 7 |
| Abhd12 | -0.3346 | 0.623 | 0.772 | 0.0007 | 2 |
| Fxyd6 | -0.3346 | 0.993 | 0.991 | 1 | 2 |
| Nr3c1 | -0.3347 | 0.116 | 0.311 | 1.01E-14 | 2 |
| Gtf2a2 | -0.3347 | 0.309 | 0.622 | 1 | 3 |
| Vamp2 | -0.3348 | 0.963 | 0.976 | 0.0005 | 4 |
| Gabrq | -0.3348 | 0.016 | 0.237 | 1.88E-06 | 3 |
| Nefl | -0.3348 | 0.052 | 0.26 | 1 | 6 |
| Atp6ap2 | -0.3351 | 0.784 | 0.841 | 0.0012 | 6 |
| Rps24 | -0.3352 | 0.995 | 0.998 | 5.73E-07 | 0 |
| Insig1 | -0.3352 | 0.141 | 0.365 | 0.0583 | 8 |
| Ict1 | -0.3353 | 0.146 | 0.522 | 1 | 3 |
| Rbbp4 | -0.3353 | 0.588 | 0.767 | 1 | 10 |
| Rap1gap | -0.3355 | 0.295 | 0.434 | 2.84E-11 | 2 |
| Hist3h2ba | -0.3356 | 0.61 | 0.713 | 0.0265 | 0 |
| Tgfa | -0.3357 | 0 | 0.26 | 7.82E-07 | 8 |
| Snap47 | -0.3357 | 0.872 | 0.979 | 0.0001 | 7 |
| Emc2 | -0.3358 | 0.098 | 0.429 | 1 | 3 |
| Ppp1r17 | -0.3358 | 0.089 | 0.335 | 1 | 3 |
| Ppfia2 | -0.3359 | 0.515 | 0.588 | 0.0128 | 0 |
| Nudc | -0.3361 | 0.821 | 0.873 | 1 | 3 |
| Minos1 | -0.3362 | 0.886 | 0.917 | 1 | 3 |

| | | | | | |
|---|---|---|---|---|---|
| Gas7 | -0.3362 | 0.151 | 0.357 | 4.60E-16 | 2 |
| Clk1 | -0.3362 | 0.467 | 0.712 | 1 | 9 |
| Zcrb1 | -0.3363 | 0.724 | 0.818 | 1 | 3 |
| Gmps | -0.3363 | 0.487 | 0.72 | 0.0057 | 7 |
| Rbfox1 | -0.3363 | 0.041 | 0.235 | 1.95E-17 | 2 |
| Pdcd6 | -0.3366 | 0.154 | 0.517 | 1 | 3 |
| 6-Mar | -0.3366 | 0.551 | 0.688 | 0.3642 | 7 |
| Ppp3r1 | -0.3366 | 0.493 | 0.709 | 0.1529 | 8 |
| Rbm4b | -0.3367 | 0.294 | 0.488 | 1 | 10 |
| Scn1b | -0.3368 | 0.535 | 0.676 | 1 | 8 |
| Dlx6 | -0.3368 | 0.009 | 0.293 | 8.19E-12 | 4 |
| Gstm5 | -0.3369 | 0.616 | 0.789 | 0.0006 | 2 |
| Nsf | -0.3371 | 0.68 | 0.85 | 0.008 | 5 |
| Emc10 | -0.3372 | 0.835 | 0.94 | 0.7563 | 5 |
| Lingo2 | -0.3372 | 0.056 | 0.308 | 0.0004 | 8 |
| Cplx2 | -0.3372 | 0.093 | 0.287 | 0.0001 | 5 |
| Scn2b | -0.3372 | 0.308 | 0.53 | 1 | 7 |
| Atp6v1d | -0.3373 | 0.984 | 0.972 | 6.99E-05 | 3 |
| Ndufv1 | -0.3374 | 0.412 | 0.568 | 1 | 10 |
| Cacna2d3 | -0.3374 | 0.254 | 0.433 | 1 | 8 |
| Anapc13 | -0.3376 | 0.228 | 0.546 | 1 | 3 |
| Disp2 | -0.3376 | 0.66 | 0.852 | 0.02 | 5 |
| Taf10 | -0.3376 | 0.447 | 0.683 | 1 | 3 |
| Chst8 | -0.3377 | 0.115 | 0.347 | 9.46E-08 | 0 |
| Calm1 | -0.3377 | 1 | 1 | 6.17E-09 | 1 |
| Ppp1cc | -0.3378 | 0.353 | 0.576 | 1 | 10 |
| Smdt1 | -0.3379 | 0.959 | 0.94 | 1 | 3 |
| Pde1a | -0.3379 | 0.075 | 0.253 | 7.49E-11 | 2 |
| Nsmce1 | -0.338 | 0.059 | 0.345 | 1 | 10 |
| Rplp0 | -0.338 | 0.86 | 0.951 | 1.15E-09 | 0 |
| Serpine2 | -0.3381 | 0.134 | 0.288 | 0.1088 | 5 |
| Atp2b4 | -0.3381 | 0.115 | 0.284 | 1 | 7 |
| Pdap1 | -0.3382 | 0.471 | 0.781 | 1 | 10 |
| Rrad | -0.3382 | 0.024 | 0.188 | 1 | 3 |
| Guk1 | -0.3384 | 0.829 | 0.867 | 1 | 3 |
| Nrsn1 | -0.3386 | 0.918 | 0.962 | 0.0945 | 5 |
| Rbfox1 | -0.3387 | 0.033 | 0.232 | 0.0317 | 3 |
| Wdr6 | -0.3388 | 0.804 | 0.946 | 0.0007 | 5 |
| Wfs1 | -0.3389 | 0.098 | 0.344 | 0.0003 | 3 |
| Wfs1 | -0.3389 | 0.155 | 0.333 | 0.0015 | 5 |
| Lrp1b | -0.3389 | 0.451 | 0.657 | 1 | 8 |
| Dmd | -0.339 | 0.103 | 0.325 | 0.0605 | 7 |

| | | | | | |
|---|---|---|---|---|---|
| Sh3gl2 | -0.3392 | 0.316 | 0.513 | 7.03E-07 | 1 |
| Unc5c | -0.3394 | 0.31 | 0.456 | 1 | 8 |
| Ndufb7 | -0.3394 | 0.821 | 0.878 | 1 | 3 |
| Igfbp5 | -0.3395 | 0.565 | 0.642 | 0.0706 | 0 |
| Pop7 | -0.3397 | 0.235 | 0.467 | 1 | 10 |
| Cplx2 | -0.3398 | 0.09 | 0.283 | 0.262 | 7 |
| Cpne4 | -0.34 | 0 | 0.211 | 5.16E-18 | 2 |
| Bhlhe41 | -0.34 | 0.2 | 0.327 | 1 | 9 |
| Iqsec3 | -0.3401 | 0.335 | 0.532 | 1.45E-06 | 0 |
| Unc5d | -0.3401 | 0.13 | 0.374 | 0.0102 | 3 |
| Ndufv3 | -0.3401 | 0.935 | 0.939 | 1 | 3 |
| Rpl30 | -0.3402 | 0.975 | 0.992 | 6.33E-06 | 0 |
| Hmgb3 | -0.3402 | 0.414 | 0.62 | 8.47E-06 | 1 |
| Ndufb2 | -0.3402 | 0.959 | 0.948 | 1 | 3 |
| Bad | -0.3403 | 0.176 | 0.432 | 1 | 10 |
| Adgrl3 | -0.3403 | 0.333 | 0.519 | 1 | 7 |
| Rabac1 | -0.3403 | 0.925 | 0.969 | 1 | 2 |
| Dtnbp1 | -0.3403 | 0.118 | 0.373 | 1 | 10 |
| Tmem132 | -0.3404 | 0.074 | 0.303 | 0.0002 | 4 |
| Cyp46a1 | -0.3404 | 0.141 | 0.384 | 0.0672 | 7 |
| Rps10 | -0.3404 | 0.97 | 0.987 | 1.33E-07 | 0 |
| C1ql3 | -0.3404 | 0.052 | 0.177 | 1 | 6 |
| Gabra2 | -0.3405 | 0.443 | 0.619 | 0.0706 | 5 |
| Gria3 | -0.3405 | 0.031 | 0.219 | 1.23E-05 | 5 |
| Izumo4 | -0.3408 | 0.433 | 0.618 | 0.0368 | 5 |
| Dusp1 | -0.3409 | 0.225 | 0.303 | 1 | 0 |
| Myl12b | -0.341 | 0.943 | 0.959 | 1 | 3 |
| Mrpl54 | -0.3411 | 0.472 | 0.709 | 1 | 3 |
| Gng3 | -0.3411 | 0.98 | 0.989 | 0.0005 | 0 |
| Akr1a1 | -0.3412 | 0.951 | 0.937 | 1 | 3 |
| Pop7 | -0.3413 | 0.163 | 0.501 | 1 | 3 |
| N6amt2 | -0.3413 | 0.098 | 0.424 | 1 | 3 |
| Cxx1b | -0.3415 | 0.854 | 0.896 | 0.1036 | 3 |
| Slc12a5 | -0.3416 | 0.808 | 0.913 | 1 | 7 |
| Ndufv2 | -0.3417 | 0.927 | 0.94 | 0.4185 | 3 |
| Nefl | -0.3418 | 0.067 | 0.247 | 1 | 9 |
| Scoc | -0.3421 | 0.943 | 0.935 | 1 | 3 |
| Ywhaz | -0.3423 | 0.908 | 0.982 | 8.35E-11 | 1 |
| Efna5 | -0.3425 | 0.072 | 0.33 | 3.45E-08 | 5 |
| Cdkn2d | -0.3425 | 0.276 | 0.606 | 1 | 3 |
| Ap2a2 | -0.3427 | 0.513 | 0.75 | 0.0313 | 7 |
| Abca2 | -0.3428 | 0.295 | 0.515 | 0.6986 | 7 |

| | | | | | |
|---|---|---|---|---|---|
| Pkib | -0.343 | 0.355 | 0.493 | 0.0509 | 0 |
| Ddx39b | -0.343 | 0.533 | 0.801 | 1 | 9 |
| Srp9 | -0.3431 | 0.902 | 0.928 | 1 | 3 |
| Itm2c | -0.3433 | 1 | 0.998 | 0.0577 | 7 |
| Tmem9 | -0.3433 | 0.526 | 0.703 | 0.0071 | 7 |
| Thsd7b | -0.3435 | 0.15 | 0.406 | 3.78E-09 | 0 |
| Ly6e | -0.3435 | 0.308 | 0.476 | 1 | 7 |
| Scd2 | -0.3435 | 0.69 | 0.813 | 1 | 8 |
| Dnajb4 | -0.3437 | 0.4 | 0.532 | 1 | 9 |
| Unc80 | -0.3437 | 0.487 | 0.678 | 0.2189 | 7 |
| 1110059I | -0.3438 | 0.118 | 0.402 | 0.9645 | 10 |
| Csrnp3 | -0.3439 | 0.128 | 0.367 | 0.1762 | 7 |
| Amd1 | -0.3442 | 0.282 | 0.501 | 0.0229 | 8 |
| Gm3811Z | -0.3446 | 0.211 | 0.397 | 1 | 8 |
| Magel2 | -0.3446 | 0.233 | 0.439 | 1 | 9 |
| Enpp5 | -0.3447 | 0.521 | 0.712 | 1 | 8 |
| Morf4l1 | -0.3447 | 1 | 0.998 | 1 | 3 |
| Pnck | -0.3451 | 0.866 | 0.932 | 0.0093 | 5 |
| Plcxd3 | -0.3452 | 0.197 | 0.427 | 0.0268 | 8 |
| Lyrm2 | -0.3452 | 0.118 | 0.435 | 1 | 10 |
| Fam213b | -0.3453 | 0.211 | 0.568 | 1 | 3 |
| Epb41l3 | -0.3453 | 0.338 | 0.562 | 0.0737 | 8 |
| Fosb | -0.3454 | 0.082 | 0.193 | 1 | 2 |
| Cd83 | -0.3457 | 0.179 | 0.407 | 0.9554 | 7 |
| 1700001I | -0.3457 | 0.191 | 0.434 | 3.75E-09 | 1 |
| Zbtb20 | -0.3458 | 0.805 | 0.731 | 0.0026 | 3 |
| Kcnip4 | -0.3458 | 0.021 | 0.247 | 6.19E-09 | 5 |
| Tmsb10 | -0.346 | 1 | 0.997 | 0.0007 | 4 |
| Tmx2 | -0.346 | 0.679 | 0.905 | 9.85E-05 | 7 |
| Rpp25l | -0.3461 | 0.118 | 0.404 | 1 | 10 |
| Camk2d | -0.3461 | 0.226 | 0.383 | 6.88E-14 | 2 |
| Rhoa | -0.3464 | 0.4 | 0.682 | 1 | 9 |
| Glrx5 | -0.3464 | 0.667 | 0.794 | 1 | 3 |
| Gabra1 | -0.3466 | 0.195 | 0.387 | 0.1336 | 3 |
| Npas4 | -0.3466 | 0.145 | 0.257 | 1 | 0 |
| Sugt1 | -0.3466 | 0.154 | 0.503 | 1 | 3 |
| Ddc | -0.3469 | 0.038 | 0.25 | 0.0166 | 7 |
| Gap43 | -0.347 | 0.969 | 0.993 | 0.0132 | 5 |
| Tuba1a | -0.3471 | 0.97 | 0.983 | 2.66E-06 | 0 |
| Lonrf2 | -0.3472 | 0.423 | 0.594 | 1 | 7 |
| Spock3 | -0.3472 | 0.306 | 0.576 | 0.0044 | 4 |
| Pde1a | -0.3475 | 0.062 | 0.246 | 0.7115 | 6 |

| | | | | | |
|---|---|---|---|---|---|
| mt-Nd3 | -0.3475 | 0.856 | 0.934 | 1 | 5 |
| Igsf8 | -0.3477 | 0.69 | 0.828 | 1 | 8 |
| Rpl7 | -0.348 | 0.894 | 0.922 | 1 | 3 |
| Gaa | -0.348 | 0.987 | 0.973 | 1 | 7 |
| Enc1 | -0.3481 | 0.051 | 0.286 | 0.009 | 7 |
| 4930447( | -0.3482 | 0 | 0.269 | 0.7854 | 10 |
| Pgm2l1 | -0.3483 | 0.31 | 0.487 | 0.9547 | 8 |
| Pmvk | -0.3483 | 0.146 | 0.498 | 1 | 3 |
| Txndc15 | -0.3485 | 0.753 | 0.875 | 1 | 2 |
| Snhg11 | -0.3485 | 1 | 0.999 | 1 | 9 |
| Rps5 | -0.3485 | 1 | 0.989 | 0.6854 | 3 |
| Ildr2 | -0.3485 | 0.137 | 0.347 | 1.49E-12 | 2 |
| Gm16105 | -0.3487 | 0.353 | 0.467 | 1 | 10 |
| Stmn1 | -0.3487 | 0.986 | 1 | 1 | 8 |
| Ndufa6 | -0.3488 | 0.878 | 0.89 | 1 | 3 |
| Snrpd3 | -0.3492 | 0.585 | 0.747 | 1 | 3 |
| Tmem199 | -0.3493 | 0.412 | 0.656 | 1 | 10 |
| Vamp2 | -0.3493 | 0.93 | 0.978 | 0.0004 | 8 |
| Timm9 | -0.3494 | 0.171 | 0.535 | 1 | 3 |
| Dlx1 | -0.3496 | 0.052 | 0.323 | 0.0781 | 6 |
| Clu | -0.3497 | 0.577 | 0.606 | 2.04E-05 | 3 |
| Pde10a | -0.3498 | 0.093 | 0.32 | 0.5698 | 6 |
| Gm15261 | -0.3498 | 0 | 0.261 | 4.87E-17 | 1 |
| Tcf12 | -0.3498 | 0.191 | 0.421 | 1.59E-06 | 1 |
| Erdr1 | -0.35 | 0.948 | 0.902 | 0.1075 | 6 |
| Tcf4 | -0.3501 | 0.118 | 0.308 | 1 | 10 |
| Rcn1 | -0.3503 | 0.141 | 0.342 | 0.0442 | 7 |
| Kif1b | -0.3503 | 0.773 | 0.886 | 0.0059 | 5 |
| Wfs1 | -0.3505 | 0.144 | 0.343 | 6.13E-10 | 2 |
| Slc39a6 | -0.3506 | 0.294 | 0.48 | 1 | 10 |
| Rac3 | -0.3506 | 0.588 | 0.749 | 0.0159 | 5 |
| Gpm6a | -0.3507 | 0.887 | 0.971 | 1 | 8 |
| Btg1 | -0.3507 | 0.6 | 0.782 | 1 | 9 |
| Romo1 | -0.3509 | 0.93 | 0.968 | 2.61E-07 | 0 |
| Gm13889 | -0.3509 | 0.25 | 0.393 | 1 | 0 |
| Arl6ip5 | -0.351 | 0.788 | 0.873 | 0.587 | 2 |
| Gabbr1 | -0.3511 | 0.911 | 0.96 | 0.0065 | 2 |
| Uba52 | -0.3512 | 0.767 | 0.877 | 1 | 9 |
| Dynlrb1 | -0.3512 | 0.927 | 0.95 | 0.5876 | 3 |
| Dbpht2 | -0.3514 | 0.073 | 0.352 | 1 | 3 |
| Tmem191 | -0.3514 | 0.538 | 0.758 | 0.0051 | 7 |
| 4930447( | -0.3517 | 0 | 0.272 | 0.0985 | 9 |

| | | | | | |
|---|---|---|---|---|---|
| Vdac1 | -0.3518 | 0.901 | 0.957 | 1.65E-08 | 1 |
| Dlx1 | -0.3518 | 0.059 | 0.303 | 1 | 10 |
| 2010107I | -0.3519 | 0.902 | 0.933 | 1 | 3 |
| Cnksr2 | -0.352 | 0.113 | 0.356 | 0.0102 | 8 |
| Eif1b | -0.352 | 0.992 | 0.975 | 0.4546 | 3 |
| Ywhab | -0.3521 | 0.984 | 0.984 | 1 | 3 |
| Slc24a5 | -0.3521 | 0.333 | 0.571 | 1 | 9 |
| Eno2 | -0.3522 | 0.565 | 0.733 | 0.0105 | 4 |
| Lypd1 | -0.3523 | 0.089 | 0.232 | 5.69E-08 | 2 |
| Tspan13 | -0.3523 | 0.719 | 0.855 | 0.0019 | 2 |
| Aip | -0.3523 | 0.211 | 0.569 | 1 | 3 |
| Tm2d3 | -0.3523 | 0.699 | 0.819 | 1 | 2 |
| Ociad1 | -0.3525 | 0.788 | 0.927 | 0.0004 | 2 |
| Hmgb3 | -0.3527 | 0.46 | 0.621 | 0.0001 | 0 |
| Sirpa | -0.3528 | 0.333 | 0.568 | 0.4706 | 7 |
| Tef | -0.353 | 0.479 | 0.654 | 1 | 8 |
| N6amt2 | -0.353 | 0.118 | 0.392 | 1 | 10 |
| Gabrq | -0.3531 | 0.014 | 0.243 | 1.03E-16 | 2 |
| Rps29 | -0.3532 | 1 | 0.997 | 8.53E-06 | 0 |
| Arl6 | -0.3534 | 0.176 | 0.421 | 1 | 10 |
| Bad | -0.3534 | 0.122 | 0.466 | 1 | 3 |
| Rsrp1 | -0.3535 | 0.824 | 0.977 | 1 | 10 |
| Brinp3 | -0.3535 | 0.254 | 0.424 | 1 | 8 |
| Gria3 | -0.3539 | 0.013 | 0.217 | 0.0026 | 7 |
| Cpne5 | -0.3539 | 0.123 | 0.309 | 8.87E-11 | 2 |
| Ptpn5 | -0.3539 | 0.164 | 0.376 | 2.76E-12 | 2 |
| Unc50 | -0.3539 | 0.118 | 0.452 | 1 | 10 |
| Cyfip2 | -0.354 | 0.5 | 0.67 | 1 | 7 |
| Rplp1 | -0.354 | 0.992 | 0.992 | 1 | 3 |
| Ogfrl1 | -0.3541 | 0.465 | 0.654 | 1 | 8 |
| Plppr3 | -0.3541 | 0.41 | 0.617 | 0.0886 | 7 |
| Ptges3 | -0.3542 | 0.765 | 0.923 | 1 | 10 |
| Synj1 | -0.3545 | 0.462 | 0.671 | 0.0946 | 7 |
| mt-Co1 | -0.3546 | 1 | 1 | 1.82E-08 | 2 |
| Zcchc12 | -0.3546 | 0.907 | 0.967 | 0.0454 | 5 |
| Fam136a | -0.3547 | 0.203 | 0.553 | 1 | 3 |
| Fam174a | -0.3548 | 0.767 | 0.86 | 1 | 9 |
| Cnr1 | -0.3549 | 0.021 | 0.237 | 2.07E-19 | 2 |
| Kcnq1ot1 | -0.3553 | 0.62 | 0.849 | 0.8433 | 8 |
| Fis1 | -0.3554 | 0.927 | 0.934 | 1 | 3 |
| Snf8 | -0.3555 | 0.146 | 0.51 | 1 | 3 |
| Btg2 | -0.3556 | 0.196 | 0.299 | 1 | 6 |

| | | | | | |
|---|---|---|---|---|---|
| Ppa1 | -0.3556 | 0.187 | 0.511 | 1 | 3 |
| Ssbp3 | -0.3556 | 0.633 | 0.826 | 1 | 9 |
| Shfm1 | -0.3557 | 0.935 | 0.943 | 1 | 3 |
| Hbegf | -0.3557 | 0.07 | 0.246 | 7.47E-06 | 0 |
| Pde10a | -0.3557 | 0.178 | 0.319 | 7.71E-06 | 2 |
| Rsrp1 | -0.3558 | 0.93 | 0.978 | 0.1548 | 8 |
| Plcl1 | -0.3558 | 0.066 | 0.329 | 1.43E-12 | 1 |
| Nsg2 | -0.3558 | 0.921 | 0.953 | 0.0042 | 1 |
| Rcn2 | -0.3558 | 0.467 | 0.649 | 1 | 9 |
| Gad1 | -0.356 | 0.664 | 0.775 | 0.0079 | 1 |
| Sgcz | -0.3561 | 0.093 | 0.341 | 2.39E-05 | 4 |
| Kif5c | -0.3563 | 0.639 | 0.823 | 0.0174 | 5 |
| Ntrk2 | -0.3564 | 0.462 | 0.677 | 0.0285 | 7 |
| Sdc2 | -0.3564 | 0.25 | 0.511 | 4.69E-07 | 1 |
| Rps7 | -0.3565 | 0.95 | 0.987 | 5.37E-08 | 0 |
| Fuom | -0.357 | 0.268 | 0.579 | 1 | 3 |
| Lrrc4b | -0.357 | 0.62 | 0.761 | 1 | 8 |
| Ddc | -0.3571 | 0.052 | 0.252 | 0.2485 | 6 |
| Gm6483 | -0.3575 | 0.507 | 0.729 | 1 | 8 |
| Galnt16 | -0.3575 | 0.247 | 0.394 | 5.70E-08 | 2 |
| Rpl41 | -0.3576 | 1 | 1 | 1.29E-10 | 0 |
| Cfdp1 | -0.3576 | 0.911 | 0.926 | 1 | 3 |
| Ptms | -0.3579 | 0.967 | 0.972 | 0.0002 | 1 |
| Ppfia2 | -0.3579 | 0.417 | 0.591 | 0.1826 | 4 |
| Wfs1 | -0.358 | 0.103 | 0.333 | 0.0356 | 7 |
| Gnas | -0.3581 | 1 | 1 | 9.24E-06 | 5 |
| Dnajc12 | -0.3581 | 0.118 | 0.405 | 1 | 10 |
| Sdf2 | -0.3581 | 0.633 | 0.797 | 1 | 9 |
| Rasgrf1 | -0.3581 | 0.62 | 0.828 | 0.0002 | 4 |
| Chst1 | -0.3582 | 0.763 | 0.905 | 2.19E-06 | 1 |
| Serpine2 | -0.3584 | 0.127 | 0.284 | 1 | 8 |
| Hmgn2 | -0.3584 | 0.8 | 0.907 | 1 | 9 |
| 7-Sep | -0.3586 | 0.824 | 0.916 | 1 | 10 |
| Anks1b | -0.3587 | 0.179 | 0.428 | 0.1028 | 7 |
| Gria3 | -0.3587 | 0.021 | 0.23 | 3.09E-15 | 2 |
| Hnrnpk | -0.3589 | 0.882 | 0.958 | 1 | 10 |
| Per3 | -0.359 | 0.495 | 0.606 | 0.0001 | 0 |
| Tmem91 | -0.359 | 0.505 | 0.766 | 0.0014 | 5 |
| Pnoc | -0.359 | 0.008 | 0.172 | 0.3185 | 3 |
| Polb | -0.3591 | 0.059 | 0.361 | 1 | 10 |
| Cdh13 | -0.3592 | 0.185 | 0.336 | 4.83E-08 | 2 |
| Tenm4 | -0.3595 | 0.397 | 0.538 | 1 | 7 |

| | | | | | |
|---|---|---|---|---|---|
| Nptxr | -0.3595 | 0.39 | 0.6 | 0.0001 | 2 |
| PISD | -0.3596 | 0.704 | 0.843 | 1 | 8 |
| Oaz1 | -0.3598 | 1 | 0.998 | 0.1091 | 3 |
| Rps3a1 | -0.3599 | 0.94 | 0.988 | 5.80E-08 | 0 |
| Dtnbp1 | -0.36 | 0.067 | 0.377 | 0.0738 | 9 |
| Dbi | -0.36 | 0.366 | 0.552 | 0.8221 | 8 |
| Timp2 | -0.3604 | 0.831 | 0.898 | 1 | 8 |
| Cpe | -0.3605 | 0.966 | 0.968 | 1 | 2 |
| Slc24a5 | -0.3605 | 0.471 | 0.566 | 1 | 10 |
| Rasgrf2 | -0.3608 | 0.521 | 0.703 | 1 | 8 |
| Dnm1 | -0.3608 | 0.366 | 0.693 | 0.0828 | 8 |
| Lamtor2 | -0.3609 | 0.837 | 0.898 | 1 | 3 |
| Minpp1 | -0.3609 | 0.294 | 0.597 | 1 | 10 |
| Dpysl2 | -0.361 | 0.887 | 0.969 | 0.0034 | 8 |
| Sez6l | -0.361 | 0.433 | 0.621 | 0.0413 | 5 |
| Chst8 | -0.361 | 0.112 | 0.336 | 2.38E-05 | 1 |
| Etnk1 | -0.3611 | 0.654 | 0.813 | 0.0958 | 7 |
| Rps3 | -0.3611 | 0.97 | 0.984 | 1.18E-07 | 0 |
| Tra2a | -0.3613 | 0.529 | 0.793 | 1 | 10 |
| Gm21092 | -0.3614 | 0.437 | 0.65 | 1 | 8 |
| Sssca1 | -0.3614 | 0.235 | 0.469 | 1 | 10 |
| Hlf | -0.3616 | 0.333 | 0.529 | 1 | 9 |
| Scn3a | -0.3618 | 0.321 | 0.524 | 0.5675 | 7 |
| Syp | -0.3621 | 0.887 | 0.94 | 0.2926 | 5 |
| Krt1 | -0.3622 | 0.016 | 0.319 | 0.0057 | 3 |
| Prlr | -0.3624 | 0.052 | 0.234 | 1 | 6 |
| Pdzrn4 | -0.3626 | 0.077 | 0.298 | 0.0239 | 7 |
| Vgf | -0.3627 | 0.093 | 0.298 | 1 | 6 |
| Atp6ap1 | -0.3627 | 0.644 | 0.814 | 0.2982 | 2 |
| Hmgn2 | -0.3627 | 0.824 | 0.913 | 0.0063 | 4 |
| Ddt | -0.3627 | 0.081 | 0.434 | 1 | 3 |
| Nr3c1 | -0.3633 | 0.082 | 0.305 | 2.77E-07 | 5 |
| Tpm1 | -0.3634 | 0.252 | 0.582 | 1 | 3 |
| Nt5m | -0.3635 | 0.235 | 0.471 | 1 | 10 |
| Anxa5 | -0.3636 | 0.008 | 0.27 | 0.0039 | 3 |
| Atp6v1f | -0.3637 | 0.984 | 0.97 | 1 | 3 |
| Ptpn5 | -0.3638 | 0.103 | 0.372 | 0.0151 | 6 |
| Dnajb1 | -0.3639 | 0.308 | 0.333 | 1 | 2 |
| Olfm2 | -0.3645 | 0.059 | 0.359 | 1 | 10 |
| Cpne2 | -0.3645 | 0.231 | 0.477 | 6.02E-05 | 4 |
| Nt5dc3 | -0.3645 | 0.085 | 0.351 | 0.0019 | 8 |
| Sbds | -0.3646 | 0.089 | 0.457 | 1 | 3 |

| | | | | | |
|---|---|---|---|---|---|
| Rcn1 | -0.3646 | 0.113 | 0.348 | 0.0766 | 6 |
| Arpp21 | -0.3646 | 0.171 | 0.414 | 6.95E-07 | 1 |
| Rnf7 | -0.3652 | 0.81 | 0.916 | 9.38E-10 | 0 |
| Nefl | -0.3655 | 0.052 | 0.26 | 1.14E-07 | 5 |
| Cntnap2 | -0.3656 | 0.25 | 0.518 | 3.50E-05 | 4 |
| Cacna2d2 | -0.3657 | 0.269 | 0.579 | 1.06E-05 | 4 |
| Tox3 | -0.3659 | 0.194 | 0.399 | 0.0056 | 4 |
| Epdr1 | -0.3659 | 0.134 | 0.447 | 0.0032 | 6 |
| Peg10 | -0.366 | 0.216 | 0.336 | 0.1478 | 5 |
| Mycbp2 | -0.366 | 0.767 | 0.911 | 0.5864 | 2 |
| Rcn1 | -0.366 | 0.118 | 0.331 | 1 | 10 |
| Sf3b5 | -0.3661 | 0.317 | 0.63 | 1 | 3 |
| Mrps14 | -0.3661 | 0.647 | 0.906 | 1 | 10 |
| Cntnap5a | -0.3663 | 0.077 | 0.331 | 0.011 | 7 |
| Gnas | -0.3663 | 1 | 1 | 0.0045 | 8 |
| B3galnt1 | -0.3664 | 0.082 | 0.417 | 0.0039 | 6 |
| Ecel1 | -0.3664 | 0.157 | 0.237 | 1 | 4 |
| Grin3a | -0.3668 | 0.165 | 0.356 | 0.003 | 5 |
| Akr7a5 | -0.3669 | 0.059 | 0.374 | 1 | 10 |
| Npas4 | -0.367 | 0.155 | 0.245 | 1 | 6 |
| Ppil3 | -0.3672 | 0 | 0.332 | 0.11 | 10 |
| Nsf | -0.3673 | 0.733 | 0.851 | 2.48E-07 | 2 |
| Nsg1 | -0.3673 | 1 | 0.997 | 0.002 | 2 |
| Gemin7 | -0.3675 | 0.059 | 0.391 | 1 | 10 |
| Iqsec3 | -0.3676 | 0.259 | 0.522 | 0.0002 | 4 |
| Ociad1 | -0.3676 | 0.744 | 0.921 | 1.67E-05 | 7 |
| Nme1 | -0.3677 | 0.967 | 0.975 | 1 | 3 |
| Gnai1 | -0.368 | 0.588 | 0.745 | 1 | 10 |
| Mtfp1 | -0.368 | 0.167 | 0.41 | 0.019 | 7 |
| Ache | -0.368 | 0.299 | 0.575 | 0.0004 | 5 |
| Pcbd2 | -0.368 | 0.089 | 0.423 | 1 | 3 |
| Avpi1 | -0.3681 | 0.057 | 0.351 | 1 | 3 |
| Gng5 | -0.3683 | 0.065 | 0.382 | 1 | 3 |
| Sec61b | -0.369 | 0.675 | 0.79 | 1 | 3 |
| Ssbp2 | -0.3692 | 0.533 | 0.752 | 1 | 9 |
| Psmb7 | -0.3692 | 0.211 | 0.579 | 1 | 3 |
| Gabra5 | -0.3694 | 0.055 | 0.283 | 3.48E-16 | 2 |
| Ephb1 | -0.3698 | 0.151 | 0.396 | 4.20E-08 | 1 |
| Tial1 | -0.3699 | 0.467 | 0.683 | 1 | 9 |
| Pcnp | -0.3703 | 0.5 | 0.713 | 1 | 9 |
| Tpi1 | -0.3704 | 0.976 | 0.981 | 0.0255 | 3 |
| Snca | -0.3704 | 0.141 | 0.337 | 0.244 | 7 |

| | | | | | |
|---|---|---|---|---|---|
| Cnr1 | -0.3706 | 0 | 0.225 | 7.67E-05 | 7 |
| Kcnd2 | -0.3706 | 0.165 | 0.418 | 2.43E-07 | 5 |
| Nrxn1 | -0.3707 | 0.918 | 0.923 | 0.9889 | 2 |
| mt-Co3 | -0.3707 | 1 | 1 | 0.0101 | 5 |
| Zfp560 | -0.371 | 0.059 | 0.369 | 1 | 10 |
| Rora | -0.3711 | 0.38 | 0.465 | 0.2187 | 0 |
| Ift22 | -0.3713 | 0.39 | 0.674 | 1 | 3 |
| Hmgcs1 | -0.3716 | 0.402 | 0.624 | 0.0022 | 5 |
| Slc25a5 | -0.3716 | 0.231 | 0.47 | 0.0008 | 4 |
| Fth1 | -0.3718 | 1 | 1 | 0.1003 | 3 |
| Gas7 | -0.3718 | 0.089 | 0.359 | 0.3091 | 3 |
| Arglu1 | -0.3719 | 0.588 | 0.856 | 1 | 10 |
| Alcam | -0.3722 | 0.765 | 0.806 | 1.88E-05 | 0 |
| Cnksr2 | -0.3724 | 0.1 | 0.347 | 1 | 9 |
| Cspg5 | -0.3726 | 0.815 | 0.915 | 0.0291 | 2 |
| Bex1 | -0.3726 | 0.976 | 0.978 | 1 | 3 |
| Dusp26 | -0.3727 | 0.471 | 0.792 | 1 | 10 |
| Tmem50a | -0.373 | 0.726 | 0.846 | 0.5539 | 2 |
| Pmepa1 | -0.3732 | 0.176 | 0.397 | 1 | 10 |
| Gria1 | -0.3733 | 0.39 | 0.593 | 5.29E-08 | 3 |
| Meg3 | -0.3733 | 1 | 0.999 | 0.001 | 6 |
| Lonrf2 | -0.3733 | 0.423 | 0.593 | 0.7508 | 8 |
| Elmo1 | -0.3734 | 0.057 | 0.298 | 1 | 3 |
| Commd1 | -0.3739 | 0.118 | 0.407 | 1 | 10 |
| Sdhaf1 | -0.374 | 0.059 | 0.377 | 1 | 10 |
| Rala | -0.374 | 0.114 | 0.491 | 0.1734 | 3 |
| Rps27 | -0.3741 | 0.97 | 0.998 | 2.86E-08 | 0 |
| Mpc1 | -0.3743 | 0.707 | 0.822 | 0.4895 | 3 |
| Kif5a | -0.3743 | 0.551 | 0.732 | 0.0759 | 7 |
| Trp53i11 | -0.3744 | 0.615 | 0.748 | 2.94E-07 | 0 |
| Jund | -0.3744 | 0.98 | 0.989 | 6.80E-06 | 0 |
| Rpl12 | -0.3745 | 0.846 | 0.893 | 1 | 3 |
| Rab28 | -0.3745 | 0.367 | 0.571 | 1 | 9 |
| Nnat | -0.3745 | 0.796 | 0.905 | 0.0024 | 1 |
| Zrsr2 | -0.3747 | 0.301 | 0.624 | 0.5827 | 3 |
| Rcan2 | -0.3747 | 0.13 | 0.484 | 1 | 3 |
| Crtac1 | -0.3748 | 0.01 | 0.232 | 5.32E-07 | 5 |
| Gabarap | -0.375 | 0.959 | 0.953 | 1 | 3 |
| Bcl11a | -0.375 | 0.167 | 0.448 | 1 | 9 |
| 2700060I | -0.375 | 0.764 | 0.842 | 1 | 3 |
| Gas7 | -0.3751 | 0.103 | 0.347 | 0.0064 | 7 |
| Edil3 | -0.3751 | 0.454 | 0.676 | 0.0437 | 5 |

| | | | | | |
|---|---|---|---|---|---|
| Dner | -0.3752 | 0.697 | 0.855 | 5.34E-05 | 1 |
| Morn2 | -0.3752 | 0.106 | 0.432 | 1 | 3 |
| Psap | -0.3753 | 0.936 | 0.966 | 0.2048 | 7 |
| Six6 | -0.3755 | 0.285 | 0.448 | 0.0199 | 0 |
| Baiap3 | -0.3757 | 0.825 | 0.912 | 0.0091 | 5 |
| Hsp90aa1 | -0.3757 | 1 | 1 | 1.22E-08 | 2 |
| Psma1 | -0.3757 | 0.412 | 0.799 | 1 | 10 |
| Snrnp40 | -0.3758 | 0.059 | 0.39 | 1 | 10 |
| Grm1 | -0.3758 | 0.039 | 0.312 | 3.37E-13 | 1 |
| Atox1 | -0.376 | 0.439 | 0.672 | 1 | 3 |
| Mpped2 | -0.376 | 0.148 | 0.41 | 1.98E-05 | 4 |
| Rpl37a | -0.3761 | 1 | 0.998 | 1.86E-07 | 0 |
| Trappc6b | -0.3762 | 0.647 | 0.879 | 1 | 10 |
| H3f3a | -0.3763 | 0.917 | 0.955 | 0.0012 | 4 |
| Ik | -0.3765 | 0.235 | 0.537 | 1 | 10 |
| Clta | -0.377 | 0.943 | 0.962 | 0.391 | 3 |
| Rps20 | -0.3772 | 0.935 | 0.988 | 2.48E-10 | 0 |
| Prkacb | -0.3775 | 0.859 | 0.957 | 0.0104 | 7 |
| Acot13 | -0.3776 | 0.163 | 0.513 | 1 | 3 |
| Btf3 | -0.3778 | 0.984 | 0.969 | 1 | 3 |
| Cux1 | -0.3779 | 0.2 | 0.442 | 1 | 9 |
| Nsf | -0.378 | 0.676 | 0.846 | 1 | 8 |
| mt-Nd1 | -0.378 | 1 | 1 | 1 | 5 |
| Fosb | -0.3782 | 0.052 | 0.191 | 0.685 | 5 |
| Dst | -0.3782 | 0.436 | 0.654 | 0.0971 | 7 |
| B630019I | -0.3784 | 0.645 | 0.844 | 1.80E-08 | 1 |
| Rpl11 | -0.3784 | 0.99 | 0.996 | 7.23E-12 | 0 |
| Ttc33 | -0.3785 | 0.353 | 0.561 | 1 | 10 |
| Zfhx3 | -0.3788 | 0.665 | 0.745 | 3.20E-05 | 0 |
| Ndufb10 | -0.3789 | 0.935 | 0.926 | 1 | 3 |
| Rmst | -0.379 | 0.254 | 0.486 | 1 | 8 |
| Srsf1 | -0.3792 | 0.433 | 0.703 | 1 | 9 |
| Atp6v0e2 | -0.3794 | 0.815 | 0.943 | 1.68E-05 | 2 |
| Dgcr6 | -0.3794 | 0.276 | 0.612 | 1 | 3 |
| Nptn | -0.3796 | 0.676 | 0.836 | 0.1356 | 8 |
| Ptpro | -0.3797 | 0.041 | 0.334 | 6.43E-07 | 3 |
| Grin2b | -0.3799 | 0.515 | 0.689 | 0.0169 | 5 |
| Atp1b1 | -0.38 | 1 | 0.998 | 3.17E-10 | 6 |
| Npdc1 | -0.3802 | 0.918 | 0.955 | 0.5646 | 5 |
| Magee1 | -0.3802 | 0.367 | 0.605 | 1 | 9 |
| Alcam | -0.3803 | 0.645 | 0.823 | 0.0003 | 1 |
| L1cam | -0.3804 | 0.267 | 0.513 | 1 | 9 |

| | | | | | |
|---|---|---|---|---|---|
| Selk | -0.3804 | 0.882 | 0.993 | 1 | 10 |
| Zrsr2 | -0.3805 | 0.353 | 0.593 | 1 | 10 |
| Llph | -0.3807 | 0.146 | 0.508 | 1 | 3 |
| Tmem256 | -0.3808 | 0.472 | 0.698 | 1 | 3 |
| Mast4 | -0.3809 | 0.3 | 0.471 | 1 | 9 |
| Rps15a | -0.3812 | 0.96 | 0.987 | 5.55E-11 | 0 |
| Arhgdig | -0.3812 | 0.733 | 0.84 | 0.0037 | 2 |
| Rplp2 | -0.3818 | 0.992 | 0.985 | 1 | 3 |
| Ntm | -0.3819 | 0.658 | 0.834 | 0.0037 | 2 |
| Ptprt | -0.382 | 0.039 | 0.291 | 5.95E-11 | 1 |
| Camk2d | -0.3821 | 0.124 | 0.386 | 0.6918 | 6 |
| Stmn3 | -0.3828 | 1 | 1 | 1.56E-05 | 5 |
| Ildr2 | -0.383 | 0.082 | 0.342 | 3.11E-08 | 5 |
| Mrps33 | -0.383 | 0.894 | 0.914 | 1 | 3 |
| Sparcl1 | -0.383 | 0.026 | 0.183 | 0.2437 | 7 |
| Pnrc1 | -0.3831 | 0.6 | 0.763 | 1.47E-08 | 0 |
| Lypd1 | -0.3832 | 0.051 | 0.226 | 0.2974 | 7 |
| Sox5 | -0.3833 | 0.18 | 0.462 | 9.02E-12 | 0 |
| Sparcl1 | -0.3835 | 0.021 | 0.186 | 0.0003 | 5 |
| Rps28 | -0.3836 | 0.965 | 0.995 | 1.59E-10 | 0 |
| Cadm3 | -0.3838 | 0.425 | 0.597 | 8.23E-07 | 2 |
| Eef1d | -0.3838 | 0.176 | 0.446 | 1 | 10 |
| Camk2g | -0.3838 | 0.308 | 0.535 | 0.0223 | 7 |
| Astn1 | -0.3839 | 0.538 | 0.737 | 0.0044 | 7 |
| Rcn1 | -0.384 | 0.113 | 0.348 | 7.99E-06 | 5 |
| Cst3 | -0.3841 | 1 | 0.988 | 0.684 | 7 |
| Ppp2r2b | -0.3844 | 0.256 | 0.515 | 0.0024 | 7 |
| Tpm1 | -0.3845 | 0.294 | 0.55 | 1 | 10 |
| Sox11 | -0.3846 | 0.3 | 0.495 | 1 | 9 |
| Stmn1 | -0.3847 | 1 | 0.999 | 2.20E-05 | 1 |
| Zfp771 | -0.3847 | 0.118 | 0.503 | 1 | 10 |
| 0610009I | -0.3847 | 0.176 | 0.505 | 1 | 10 |
| Scn1a | -0.3848 | 0.051 | 0.299 | 0.0019 | 7 |
| Ptprs | -0.3849 | 0.711 | 0.862 | 0.0739 | 5 |
| Chst1 | -0.3849 | 0.775 | 0.893 | 0.9551 | 8 |
| Ccdc32 | -0.3849 | 0.081 | 0.445 | 1 | 3 |
| Tmx2 | -0.3849 | 0.801 | 0.902 | 0.0872 | 2 |
| Tesc | -0.3855 | 0.033 | 0.335 | 0.4217 | 3 |
| Rps21 | -0.3855 | 0.965 | 0.99 | 1.23E-08 | 0 |
| Pnoc | -0.3856 | 0.026 | 0.163 | 0.3903 | 7 |
| PISD | -0.3856 | 0.856 | 0.832 | 3.96E-05 | 6 |
| Cdh13 | -0.3859 | 0.128 | 0.33 | 0.6148 | 7 |

| | | | | | |
|---|---|---|---|---|---|
| Pnoc | -0.386 | 0.01 | 0.167 | 0.0025 | 5 |
| Tenm2 | -0.3862 | 0.133 | 0.388 | 1 | 9 |
| Pcdh17 | -0.3864 | 0.707 | 0.636 | 0.0001 | 3 |
| Ptprt | -0.3864 | 0.038 | 0.273 | 0.0022 | 7 |
| Tmsb10 | -0.3865 | 0.99 | 0.999 | 4.64E-07 | 0 |
| Grin1 | -0.3866 | 0.513 | 0.696 | 0.0321 | 7 |
| Ncdn | -0.3868 | 0.433 | 0.636 | 0.0017 | 5 |
| Hist3h2ba | -0.3868 | 0.546 | 0.71 | 0.0179 | 4 |
| Nefl | -0.3869 | 0.024 | 0.269 | 0.0055 | 3 |
| Cdk14 | -0.3873 | 0.077 | 0.327 | 0.0034 | 7 |
| Txnl4a | -0.3874 | 0.163 | 0.515 | 1 | 3 |
| Wbp5 | -0.3874 | 0.943 | 0.945 | 0.0014 | 3 |
| Trnau1ap | -0.3875 | 0.059 | 0.393 | 1 | 10 |
| Fabp5 | -0.3876 | 0.886 | 0.861 | 1 | 3 |
| Slc16a11 | -0.3878 | 0.028 | 0.298 | 2.56E-05 | 8 |
| Rpl18a | -0.3881 | 0.992 | 0.99 | 1 | 3 |
| Rps13 | -0.3882 | 0.992 | 0.977 | 1 | 3 |
| Tsnax | -0.3883 | 0.3 | 0.632 | 0.1309 | 9 |
| Adh5 | -0.3883 | 0.118 | 0.371 | 1 | 10 |
| Gde1 | -0.3884 | 0.704 | 0.885 | 0.0152 | 8 |
| 1700086I | -0.3885 | 0.186 | 0.389 | 0.0004 | 5 |
| Ube3a | -0.3885 | 0.887 | 0.958 | 0.3027 | 8 |
| Gm21092 | -0.3886 | 0.4 | 0.643 | 1 | 9 |
| Ptprs | -0.3886 | 0.801 | 0.856 | 1 | 2 |
| Rpl17 | -0.3887 | 0.911 | 0.921 | 1 | 3 |
| Ptpro | -0.3888 | 0.096 | 0.333 | 1.21E-16 | 2 |
| Tcaf1 | -0.3888 | 0.718 | 0.896 | 0.1558 | 8 |
| Gpx3 | -0.3889 | 0.212 | 0.459 | 2.35E-12 | 2 |
| Gda | -0.3891 | 0.426 | 0.653 | 0.0025 | 4 |
| Pcdh9 | -0.3893 | 0.306 | 0.639 | 1.33E-06 | 4 |
| Dlx5 | -0.3896 | 0 | 0.32 | 1 | 10 |
| Tshz2 | -0.3896 | 0.233 | 0.527 | 1 | 9 |
| Gt(ROSA)2 | -0.3897 | 0.225 | 0.432 | 0.0147 | 8 |
| 1110004I | -0.3898 | 0.967 | 0.963 | 1 | 9 |
| Fxyd7 | -0.3898 | 0.461 | 0.697 | 9.03E-06 | 1 |
| Tubb3 | -0.3898 | 0.87 | 0.959 | 5.66E-08 | 2 |
| Negr1 | -0.3898 | 0.742 | 0.845 | 0.1471 | 5 |
| Sorcs1 | -0.3899 | 0.031 | 0.317 | 4.31E-11 | 5 |
| Irs4 | -0.3903 | 0.093 | 0.289 | 1 | 6 |
| Kcnk2 | -0.3904 | 0.218 | 0.372 | 1 | 7 |
| Erh | -0.3905 | 0.935 | 0.94 | 1 | 3 |
| Ngb | -0.3913 | 0.705 | 0.794 | 0.0028 | 0 |

| | | | | | |
|---|---|---|---|---|---|
| Nenf | -0.3915 | 0.986 | 0.992 | 0.0948 | 2 |
| Adcy2 | -0.3917 | 0.028 | 0.325 | 1.94E-06 | 8 |
| Cnbp | -0.3918 | 0.943 | 0.95 | 0.1426 | 3 |
| Tle4 | -0.3918 | 0.525 | 0.702 | 2.63E-08 | 0 |
| Tmed2 | -0.3919 | 0.471 | 0.878 | 1 | 10 |
| Sod1 | -0.3919 | 0.26 | 0.601 | 0.4572 | 3 |
| Gadd45g | -0.3921 | 0.465 | 0.524 | 1 | 0 |
| Akap8l | -0.3923 | 0.367 | 0.657 | 1 | 9 |
| Gpx1 | -0.3925 | 0.309 | 0.598 | 1 | 3 |
| Ildr2 | -0.3925 | 0.051 | 0.34 | 0.0002 | 7 |
| Gabarapl2 | -0.3926 | 0.992 | 0.987 | 1 | 3 |
| Gm13889 | -0.3927 | 0.268 | 0.377 | 1 | 5 |
| Sparcl1 | -0.3928 | 0.014 | 0.182 | 0.0173 | 8 |
| Calb2 | -0.3933 | 0.495 | 0.587 | 1 | 5 |
| Pdzrn4 | -0.3936 | 0.041 | 0.305 | 2.69E-09 | 5 |
| Edil3 | -0.3936 | 0.447 | 0.69 | 3.61E-06 | 1 |
| Ogt | -0.3938 | 0.676 | 0.892 | 0.0915 | 8 |
| Abr | -0.3938 | 0.495 | 0.727 | 2.64E-05 | 5 |
| Tceb2 | -0.394 | 1 | 0.992 | 1 | 3 |
| Hmgn1 | -0.3941 | 0.835 | 0.941 | 3.80E-13 | 0 |
| Fabp5 | -0.3941 | 0.732 | 0.877 | 0.0028 | 5 |
| Atp6ap2 | -0.3943 | 0.732 | 0.843 | 1 | 8 |
| Unc5c | -0.3945 | 0.222 | 0.471 | 9.47E-05 | 4 |
| Hnrnpd | -0.3947 | 0.588 | 0.811 | 1 | 10 |
| Flrt3 | -0.3951 | 0.405 | 0.58 | 5.12E-07 | 0 |
| Tmem132 | -0.3952 | 0.066 | 0.314 | 5.81E-10 | 1 |
| Ncam1 | -0.3954 | 0.887 | 0.956 | 1 | 8 |
| Tenm3 | -0.3958 | 0.365 | 0.576 | 2.07E-09 | 0 |
| Faim2 | -0.3959 | 0.962 | 0.975 | 0.0188 | 7 |
| Glod4 | -0.396 | 0.118 | 0.426 | 1 | 10 |
| Hmgcs1 | -0.396 | 0.433 | 0.61 | 1 | 9 |
| Susd4 | -0.3961 | 0.077 | 0.388 | 0.0009 | 7 |
| Nefl | -0.3963 | 0.026 | 0.258 | 0.0006 | 7 |
| Grpel1 | -0.3967 | 0.26 | 0.626 | 0.1576 | 3 |
| Pde10a | -0.3967 | 0.124 | 0.317 | 0.0008 | 5 |
| Hmgcr | -0.3967 | 0.474 | 0.716 | 0.0089 | 5 |
| Serf1 | -0.3968 | 0.106 | 0.466 | 1 | 3 |
| Clstn3 | -0.3968 | 0.648 | 0.837 | 1 | 8 |
| Ranbp1 | -0.397 | 0.87 | 0.911 | 0.7772 | 3 |
| Sgcz | -0.397 | 0.125 | 0.347 | 3.83E-08 | 1 |
| Rtn4 | -0.3971 | 0.887 | 0.954 | 0.205 | 5 |
| AI593442 | -0.3973 | 0.059 | 0.328 | 0.241 | 10 |

| | | | | | |
|---|---|---|---|---|---|
| Atp2a2 | -0.3974 | 0.634 | 0.806 | 1 | 8 |
| Plcb4 | -0.3975 | 0.34 | 0.53 | 0.0016 | 5 |
| Pin4 | -0.3976 | 0.171 | 0.533 | 1 | 3 |
| Pnmal2 | -0.3976 | 0.993 | 0.993 | 4.37E-11 | 1 |
| Nell2 | -0.3978 | 0.268 | 0.484 | 0.0015 | 5 |
| Fam204a | -0.3979 | 0.059 | 0.379 | 1 | 10 |
| Eid1 | -0.3983 | 0.951 | 0.964 | 0.1091 | 3 |
| Hprt | -0.3983 | 0.706 | 0.863 | 1 | 10 |
| Vstm2b | -0.3985 | 0.113 | 0.383 | 4.15E-05 | 8 |
| Srsf5 | -0.3987 | 0.824 | 0.909 | 1 | 10 |
| Podxl2 | -0.3987 | 0.969 | 0.987 | 0.0012 | 5 |
| Ddc | -0.3989 | 0.031 | 0.254 | 3.27E-07 | 5 |
| Pcsk2 | -0.3989 | 0.735 | 0.862 | 1.71E-09 | 0 |
| Use1 | -0.3989 | 0.176 | 0.479 | 1 | 10 |
| Sncb | -0.3991 | 0.927 | 0.917 | 0.9565 | 3 |
| Maged1 | -0.3993 | 0.986 | 0.988 | 0.3183 | 2 |
| Atp5o | -0.3995 | 0.87 | 0.873 | 1 | 3 |
| Cnksr2 | -0.3996 | 0.111 | 0.365 | 1.68E-06 | 4 |
| Rps6 | -0.3999 | 0.927 | 0.942 | 1 | 3 |
| Pde1a | -0.3999 | 0.01 | 0.25 | 2.12E-09 | 5 |
| Atp1a3 | -0.4001 | 0.904 | 0.952 | 0.0562 | 2 |
| Gcsh | -0.4005 | 0.13 | 0.509 | 0.9815 | 3 |
| Scn1a | -0.4009 | 0.041 | 0.304 | 1.58E-10 | 5 |
| Rpl36 | -0.4013 | 0.992 | 0.993 | 1 | 3 |
| Mrpl20 | -0.4015 | 0.813 | 0.887 | 1 | 3 |
| Rps26 | -0.4016 | 0.951 | 0.939 | 1 | 3 |
| Rtn4 | -0.4016 | 0.897 | 0.952 | 0.0006 | 7 |
| Chst8 | -0.4016 | 0.067 | 0.312 | 1 | 9 |
| Pcbd1 | -0.4017 | 0.642 | 0.731 | 1 | 3 |
| Tmed9 | -0.4017 | 0.718 | 0.844 | 0.0002 | 7 |
| Eef1a1 | -0.4018 | 1 | 1 | 1.09E-08 | 0 |
| Hddc2 | -0.4018 | 0 | 0.355 | 1 | 10 |
| Tmem132 | -0.4019 | 0.033 | 0.287 | 1 | 9 |
| Tmem258 | -0.4019 | 0.412 | 0.742 | 1 | 10 |
| Tmem91 | -0.4024 | 0.5 | 0.762 | 4.46E-05 | 7 |
| Thsd7b | -0.4024 | 0.141 | 0.375 | 0.0233 | 8 |
| Matk | -0.4025 | 0.538 | 0.762 | 0.0011 | 7 |
| Nudt19 | -0.4025 | 0.176 | 0.456 | 1 | 10 |
| Dlx1 | -0.4026 | 0.065 | 0.328 | 0.0537 | 3 |
| 2700029 | -0.4026 | 0.294 | 0.54 | 1 | 10 |
| Cbx3 | -0.403 | 0.676 | 0.853 | 0.004 | 8 |
| 1110004 | -0.4032 | 0.928 | 0.969 | 1.83E-07 | 1 |

| | | | | | |
|---|---|---|---|---|---|
| Ccdc136 | -0.4032 | 0.098 | 0.458 | 0.2993 | 3 |
| 1700086I | -0.4033 | 0.106 | 0.405 | 1 | 3 |
| Ap1s2 | -0.4034 | 0.463 | 0.629 | 1 | 3 |
| Cyfip2 | -0.4034 | 0.521 | 0.678 | 9.49E-12 | 2 |
| Gng3 | -0.4034 | 0.954 | 0.993 | 3.18E-09 | 1 |
| Rab3b | -0.4036 | 0.059 | 0.324 | 1 | 10 |
| Cpne5 | -0.4039 | 0.051 | 0.303 | 0.0003 | 7 |
| Lypd1 | -0.4048 | 0.033 | 0.236 | 0.0037 | 3 |
| Fxyd7 | -0.4049 | 0.436 | 0.682 | 0.0075 | 7 |
| Arhgap36 | -0.405 | 0.052 | 0.21 | 1 | 6 |
| Spint2 | -0.405 | 0.733 | 0.923 | 1 | 9 |
| Rprm | -0.4055 | 0.296 | 0.431 | 0.0944 | 1 |
| Mrap2 | -0.4058 | 0.118 | 0.378 | 1 | 10 |
| Cadm3 | -0.4058 | 0.397 | 0.588 | 0.2077 | 7 |
| Clstn3 | -0.4058 | 0.733 | 0.839 | 0.0008 | 2 |
| Gabrb1 | -0.4059 | 0.769 | 0.863 | 0.1819 | 7 |
| Cntnap5a | -0.4059 | 0.082 | 0.336 | 3.21E-07 | 5 |
| Slc25a5 | -0.406 | 0.195 | 0.478 | 1 | 3 |
| Scn9a | -0.406 | 0.3 | 0.603 | 1 | 9 |
| Gabrg2 | -0.406 | 0.538 | 0.684 | 0.0437 | 7 |
| Atp2a2 | -0.4061 | 0.705 | 0.809 | 0.0014 | 2 |
| Snca | -0.4064 | 0.103 | 0.344 | 1.13E-08 | 5 |
| Arl6ip1 | -0.4069 | 0.973 | 0.997 | 4.99E-05 | 2 |
| Dlgap1 | -0.4069 | 0.155 | 0.425 | 2.12E-07 | 5 |
| Mrpl57 | -0.407 | 0.447 | 0.684 | 1 | 3 |
| Spcs1 | -0.4071 | 0.925 | 0.96 | 0.1098 | 2 |
| Nr3c1 | -0.4073 | 0.041 | 0.316 | 4.44E-05 | 3 |
| Cbln2 | -0.4074 | 0.102 | 0.289 | 0.0198 | 4 |
| Gabrg3 | -0.4075 | 0.212 | 0.428 | 7.16E-13 | 2 |
| Cd200 | -0.4075 | 0.756 | 0.876 | 0.0012 | 7 |
| Ckmt1 | -0.4075 | 0.449 | 0.738 | 1.53E-06 | 7 |
| Cers4 | -0.4079 | 0.367 | 0.716 | 1 | 9 |
| Parm1 | -0.408 | 0.394 | 0.6 | 1 | 8 |
| Dlx1 | -0.4081 | 0.077 | 0.316 | 0.0011 | 7 |
| Syt10 | -0.4086 | 0.611 | 0.649 | 1 | 4 |
| Polr2l | -0.4086 | 0.294 | 0.626 | 1 | 10 |
| Stk32a | -0.4086 | 0.211 | 0.424 | 0.016 | 8 |
| Rpl13a | -0.4088 | 0.967 | 0.945 | 0.0917 | 3 |
| Aplp2 | -0.4089 | 0.603 | 0.75 | 0.0021 | 2 |
| Atp2b2 | -0.4091 | 0.472 | 0.755 | 6.94E-07 | 4 |
| Cdh2 | -0.4093 | 0.632 | 0.798 | 3.56E-06 | 1 |
| Ddx24 | -0.4095 | 0.842 | 0.954 | 5.02E-07 | 2 |

| | | | | | |
|---|---|---|---|---|---|
| Uqcr10 | -0.4097 | 0.976 | 0.96 | 1 | 3 |
| Scd2 | -0.4098 | 0.691 | 0.816 | 0.0048 | 5 |
| Rwdd2a | -0.41 | 0.218 | 0.498 | 0.0002 | 7 |
| Hist3h2ba | -0.4102 | 0.526 | 0.721 | 1.01E-05 | 1 |
| Sparcl1 | -0.4105 | 0.014 | 0.195 | 6.93E-13 | 2 |
| Snhg20 | -0.4108 | 0.654 | 0.843 | 0.0004 | 7 |
| Gng3 | -0.4108 | 1 | 0.986 | 0.0043 | 4 |
| Tmem59 | -0.411 | 0.588 | 0.929 | 1 | 10 |
| Rpl14 | -0.4112 | 0.992 | 0.982 | 0.1608 | 3 |
| Pcdh17 | -0.4112 | 0.526 | 0.655 | 0.0052 | 6 |
| Rps4x | -0.4113 | 0.965 | 0.986 | 4.67E-10 | 0 |
| Nr3c1 | -0.4113 | 0.038 | 0.305 | 5.65E-05 | 7 |
| Gabra5 | -0.4113 | 0.008 | 0.283 | 6.15E-09 | 3 |
| 1110008I | -0.4118 | 0.176 | 0.493 | 1 | 10 |
| Rgs17 | -0.4119 | 0.449 | 0.709 | 0.0005 | 7 |
| Thsd7b | -0.4122 | 0.102 | 0.388 | 1.43E-07 | 4 |
| Gm6483 | -0.4123 | 0.449 | 0.735 | 0.0006 | 7 |
| Rpl10 | -0.4127 | 0.935 | 0.94 | 1 | 3 |
| Scd2 | -0.4128 | 0.615 | 0.819 | 0.0004 | 7 |
| Grina | -0.4128 | 0.767 | 0.882 | 0.0261 | 2 |
| Lrpap1 | -0.4129 | 0.782 | 0.893 | 0.0057 | 7 |
| Syngr3 | -0.413 | 0.814 | 0.917 | 0.0024 | 5 |
| Ddt | -0.4133 | 0.059 | 0.4 | 1 | 10 |
| Cript | -0.4136 | 0.211 | 0.596 | 0.3253 | 3 |
| Ptn | -0.4138 | 0.432 | 0.522 | 1 | 2 |
| Cox7b | -0.4138 | 0.919 | 0.915 | 1 | 3 |
| 1110004I | -0.4139 | 0.925 | 0.972 | 8.12E-12 | 0 |
| Celf6 | -0.414 | 0.471 | 0.729 | 1 | 10 |
| Spock2 | -0.414 | 0.753 | 0.874 | 0.0007 | 2 |
| Gpx3 | -0.4143 | 0.171 | 0.459 | 0.0056 | 3 |
| Smim11 | -0.4144 | 0.059 | 0.414 | 1 | 10 |
| Ctsb | -0.4147 | 0.833 | 0.93 | 0.0004 | 7 |
| Malat1 | -0.4151 | 0.993 | 1 | 0.0232 | 1 |
| Sarnp | -0.4155 | 0.294 | 0.601 | 1 | 10 |
| Nptxr | -0.4156 | 0.34 | 0.595 | 0.0003 | 5 |
| Rab3b | -0.4156 | 0.057 | 0.352 | 0.1843 | 3 |
| Matk | -0.4161 | 0.485 | 0.771 | 5.78E-06 | 5 |
| Leng8 | -0.4164 | 0.465 | 0.71 | 0.2021 | 8 |
| Tmem191 | -0.4167 | 0.557 | 0.76 | 0.0002 | 5 |
| Atp9a | -0.4172 | 0.555 | 0.768 | 7.13E-05 | 2 |
| Ttc3 | -0.4172 | 1 | 1 | 1.26E-06 | 7 |
| Grin3a | -0.4174 | 0.141 | 0.353 | 0.0862 | 8 |

| | | | | | |
|---|---|---|---|---|---|
| Gpx4 | -0.4177 | 0.992 | 0.987 | 1 | 3 |
| Rph3a | -0.4178 | 0.294 | 0.594 | 1 | 10 |
| Sez6l2 | -0.4186 | 0.753 | 0.857 | 0.2704 | 2 |
| 1810022I | -0.4187 | 0.471 | 0.812 | 1 | 10 |
| Hmgn2 | -0.4188 | 0.82 | 0.923 | 2.00E-10 | 0 |
| Dbp | -0.4188 | 0.515 | 0.681 | 7.35E-05 | 0 |
| Vdac1 | -0.4195 | 0.803 | 0.959 | 1.33E-06 | 8 |
| Hsbp1 | -0.4198 | 0.951 | 0.966 | 0.0501 | 3 |
| Flrt3 | -0.4199 | 0.38 | 0.56 | 1 | 8 |
| Serp2 | -0.4202 | 0.76 | 0.906 | 3.02E-12 | 2 |
| Dpp6 | -0.4205 | 0.756 | 0.851 | 0.0124 | 7 |
| C1ql3 | -0.4208 | 0.052 | 0.177 | 0.1309 | 5 |
| Tac2 | -0.421 | 0 | 0.043 | 1 | 5 |
| Fam136a | -0.421 | 0.235 | 0.519 | 1 | 10 |
| Stmn4 | -0.4212 | 0.546 | 0.812 | 2.45E-05 | 5 |
| Myef2 | -0.4218 | 0.38 | 0.655 | 0.0051 | 8 |
| Stmn1 | -0.4222 | 1 | 0.999 | 6.80E-07 | 0 |
| Tomm5 | -0.4222 | 0.146 | 0.538 | 1 | 3 |
| Cd81 | -0.4222 | 0.949 | 0.973 | 4.89E-05 | 7 |
| Ppp1r14c | -0.4223 | 0 | 0.349 | 0.5131 | 10 |
| Ndufa7 | -0.4223 | 0.984 | 0.95 | 1 | 3 |
| Mea1 | -0.4224 | 0.122 | 0.507 | 1 | 3 |
| Srp14 | -0.4227 | 0.976 | 0.975 | 0.692 | 3 |
| Lypd1 | -0.4228 | 0.031 | 0.231 | 1.84E-06 | 5 |
| Ddc | -0.4229 | 0.008 | 0.263 | 1.14E-06 | 3 |
| Gria2 | -0.4229 | 0.897 | 0.959 | 0.0056 | 7 |
| Cacna2d1 | -0.4231 | 0.206 | 0.39 | 0.0111 | 5 |
| Rps29 | -0.4234 | 0.992 | 0.998 | 1 | 3 |
| Acyp2 | -0.4235 | 0 | 0.36 | 0.4309 | 10 |
| Rasgrp1 | -0.4235 | 0 | 0.289 | 1 | 10 |
| Marcksl1 | -0.4236 | 0.959 | 0.943 | 0.1572 | 3 |
| Fam171b | -0.4238 | 0.641 | 0.826 | 0.0058 | 7 |
| Ptpro | -0.4241 | 0.026 | 0.323 | 5.35E-06 | 7 |
| Bola3 | -0.4242 | 0.081 | 0.478 | 1 | 3 |
| Nefl | -0.4244 | 0.021 | 0.275 | 1.47E-27 | 2 |
| Coa3 | -0.4247 | 0.943 | 0.933 | 0.7849 | 3 |
| Spon1 | -0.4247 | 0.155 | 0.465 | 1.43E-11 | 0 |
| Sparcl1 | -0.4247 | 0 | 0.193 | 8.67E-08 | 3 |
| Rbfox3 | -0.4248 | 0.103 | 0.429 | 1.81E-05 | 7 |
| Atp2a2 | -0.425 | 0.619 | 0.812 | 0.0007 | 5 |
| Tmem59l | -0.425 | 0.846 | 0.927 | 0.0012 | 7 |
| Kcnk2 | -0.4255 | 0.165 | 0.38 | 0.2443 | 6 |

| | | | | | |
|---|---|---|---|---|---|
| Atpif1 | -0.4262 | 0.984 | 0.994 | 0.199 | 3 |
| Napb | -0.4264 | 0.256 | 0.489 | 0.0444 | 7 |
| Ddc | -0.4264 | 0.021 | 0.267 | 5.34E-18 | 2 |
| Park7 | -0.4267 | 0.943 | 0.946 | 1 | 3 |
| Dynll1 | -0.4269 | 0.976 | 0.98 | 1 | 3 |
| Ubc | -0.427 | 0.7 | 0.917 | 1 | 9 |
| Mrpl18 | -0.4271 | 0.203 | 0.585 | 0.0713 | 3 |
| Itm2c | -0.4273 | 0.972 | 1 | 0.0022 | 8 |
| Pabpn1 | -0.4273 | 0.732 | 0.883 | 0.6599 | 8 |
| Reep1 | -0.4274 | 0.167 | 0.486 | 0.0004 | 7 |
| Nagk | -0.4275 | 0.353 | 0.671 | 1 | 10 |
| C1ql3 | -0.4275 | 0.048 | 0.184 | 3.53E-05 | 2 |
| Gabra1 | -0.4276 | 0.186 | 0.383 | 1 | 6 |
| Pdcl3 | -0.4276 | 0 | 0.393 | 0.1342 | 10 |
| Fkbp2 | -0.4282 | 0.859 | 0.935 | 0.0012 | 7 |
| Atp6v1g1 | -0.4282 | 0.943 | 0.943 | 1 | 3 |
| Fam19a1 | -0.4283 | 0.256 | 0.438 | 1 | 7 |
| Vgf | -0.4284 | 0.197 | 0.286 | 1 | 8 |
| Cplx2 | -0.4284 | 0.041 | 0.304 | 1.33E-23 | 2 |
| H3f3a | -0.4285 | 0.89 | 0.965 | 5.10E-15 | 0 |
| Hnrnpk | -0.4285 | 0.867 | 0.96 | 0.1419 | 9 |
| Saraf | -0.4288 | 0.87 | 0.943 | 0.1363 | 2 |
| Spock1 | -0.4288 | 0.225 | 0.411 | 0.2629 | 8 |
| Slc6a1 | -0.4289 | 0.353 | 0.653 | 1 | 10 |
| Rps9 | -0.4291 | 0.94 | 0.982 | 1.92E-13 | 0 |
| Psap | -0.4292 | 0.948 | 0.966 | 0.1751 | 5 |
| Pkib | -0.4292 | 0.324 | 0.484 | 0.2486 | 4 |
| Grina | -0.4293 | 0.732 | 0.88 | 0.0019 | 5 |
| Tmem179 | -0.4295 | 0.742 | 0.876 | 0.0004 | 5 |
| Mif | -0.4295 | 0.967 | 0.971 | 1 | 3 |
| Tmem179 | -0.4296 | 0.731 | 0.874 | 0.0003 | 7 |
| Pnrc1 | -0.4297 | 0.529 | 0.737 | 1 | 10 |
| Set | -0.4298 | 0.941 | 0.983 | 1 | 10 |
| Dlx1 | -0.4301 | 0.065 | 0.324 | 2.57E-08 | 4 |
| Ly6e | -0.4305 | 0.26 | 0.495 | 1.44E-12 | 2 |
| Lrrtm3 | -0.4306 | 0.25 | 0.543 | 2.57E-10 | 1 |
| 1700037 | -0.4307 | 0.13 | 0.501 | 0.4845 | 3 |
| Cdk14 | -0.4312 | 0.041 | 0.335 | 2.64E-12 | 5 |
| Gria2 | -0.4313 | 0.945 | 0.956 | 0.1903 | 2 |
| Mycbp2 | -0.4313 | 0.784 | 0.902 | 0.0012 | 5 |
| Ndfip2 | -0.4318 | 0.3 | 0.579 | 0.462 | 9 |
| Pnck | -0.4319 | 0.795 | 0.936 | 2.24E-07 | 7 |

| | | | | | |
|---|---|---|---|---|---|
| Rpl36al | -0.4324 | 0.951 | 0.952 | 1 | 3 |
| Mpped2 | -0.4324 | 0.118 | 0.389 | 1 | 10 |
| Ywhah | -0.4325 | 1 | 0.999 | 0.2483 | 3 |
| Txnl1 | -0.4329 | 0.647 | 0.862 | 1 | 10 |
| Myeov2 | -0.4334 | 0.854 | 0.903 | 1 | 3 |
| Dnajb1 | -0.4334 | 0.106 | 0.357 | 1 | 3 |
| Slc17a6 | -0.4335 | 0 | 0.128 | 1 | 9 |
| Rpl35a | -0.4343 | 0.99 | 0.992 | 3.00E-08 | 0 |
| Snca | -0.4345 | 0.073 | 0.354 | 1 | 3 |
| 5330434 | -0.4345 | 0.485 | 0.754 | 0.0007 | 5 |
| Ccdc92 | -0.4347 | 0.146 | 0.569 | 0.0002 | 3 |
| Atp5g1 | -0.435 | 0.992 | 0.989 | 0.7441 | 3 |
| Tcf12 | -0.4353 | 0.099 | 0.409 | 2.67E-05 | 8 |
| Pnoc | -0.4355 | 0 | 0.177 | 6.03E-13 | 2 |
| Ajap1 | -0.4357 | 0.25 | 0.458 | 0.0005 | 4 |
| Tmem30a | -0.4359 | 0.705 | 0.881 | 9.77E-05 | 7 |
| Mir124-2 | -0.4359 | 0.046 | 0.355 | 3.42E-17 | 1 |
| Spock3 | -0.4359 | 0.316 | 0.586 | 1.49E-07 | 1 |
| Dusp1 | -0.4361 | 0.067 | 0.295 | 1 | 9 |
| Slc17a6 | -0.4363 | 0.013 | 0.133 | 1 | 7 |
| Tmem158 | -0.4364 | 0.608 | 0.723 | 8.57E-05 | 6 |
| Camk2n1 | -0.4364 | 0.533 | 0.794 | 9.05E-11 | 1 |
| Rps25 | -0.4366 | 0.935 | 0.925 | 1 | 3 |
| Prlr | -0.4366 | 0.033 | 0.241 | 4.94E-05 | 3 |
| Snrpf | -0.4367 | 0.529 | 0.83 | 1 | 10 |
| Stk32c | -0.437 | 0.239 | 0.547 | 3.34E-05 | 8 |
| App | -0.437 | 0.789 | 0.94 | 1 | 8 |
| Phyh | -0.4373 | 0.235 | 0.495 | 1 | 10 |
| Upf3b | -0.4373 | 0.235 | 0.529 | 1 | 10 |
| Hmgb3 | -0.4377 | 0.358 | 0.621 | 1 | 3 |
| Chl1 | -0.4378 | 0.351 | 0.487 | 0.0059 | 5 |
| Gabra2 | -0.4383 | 0.3 | 0.612 | 1 | 9 |
| Slc12a5 | -0.4386 | 0.887 | 0.907 | 0.3802 | 5 |
| Chga | -0.4388 | 0.608 | 0.747 | 1 | 5 |
| Olfm1 | -0.4391 | 0.68 | 0.864 | 5.08E-05 | 5 |
| Zcchc12 | -0.4396 | 0.882 | 0.963 | 1 | 10 |
| Rps27a | -0.4397 | 0.99 | 0.996 | 3.10E-11 | 0 |
| Bub3 | -0.4398 | 0.5 | 0.732 | 1 | 9 |
| Kit | -0.4402 | 0.092 | 0.346 | 2.33E-10 | 1 |
| Grin1 | -0.4403 | 0.5 | 0.711 | 1.87E-06 | 2 |
| Rpl11 | -0.4406 | 1 | 0.994 | 1 | 3 |
| Ube2n | -0.4408 | 0.412 | 0.799 | 1 | 10 |

| | | | | | |
|---|---|---|---|---|---|
| Tceal3 | -0.4409 | 0.943 | 0.945 | 0.058 | 3 |
| Nrp2 | -0.4417 | 0.282 | 0.501 | 0.0934 | 8 |
| Aldoa | -0.4418 | 1 | 1 | 5.32E-19 | 1 |
| Rac3 | -0.442 | 0.474 | 0.754 | 5.34E-07 | 7 |
| Parm1 | -0.4422 | 0.349 | 0.625 | 1.28E-07 | 1 |
| Tmem255 | -0.4422 | 0.3 | 0.563 | 1 | 9 |
| Serinc1 | -0.4425 | 0.915 | 0.97 | 0.3358 | 8 |
| Rgs4 | -0.4428 | 0.051 | 0.224 | 0.0451 | 7 |
| Zfhx3 | -0.4436 | 0.645 | 0.745 | 0.0006 | 1 |
| Ly6e | -0.4437 | 0.216 | 0.488 | 3.26E-07 | 5 |
| Cpne5 | -0.4437 | 0.031 | 0.309 | 7.78E-11 | 5 |
| Dpp6 | -0.4438 | 0.722 | 0.856 | 0.0137 | 5 |
| Cox5a | -0.4438 | 0.976 | 0.978 | 1 | 3 |
| Gm561 | -0.4438 | 0.412 | 0.74 | 1 | 10 |
| Commd7 | -0.444 | 0.118 | 0.475 | 1 | 10 |
| Atp6v0a1 | -0.4443 | 0.705 | 0.865 | 3.13E-05 | 2 |
| Btg1 | -0.4452 | 0.593 | 0.797 | 4.27E-05 | 4 |
| Dpp10 | -0.4455 | 0.031 | 0.324 | 6.51E-13 | 5 |
| Tsc22d1 | -0.4463 | 0.773 | 0.871 | 0.0372 | 5 |
| C1ql3 | -0.4464 | 0 | 0.169 | 1 | 10 |
| Timm8b | -0.4466 | 0.943 | 0.941 | 1 | 3 |
| Cebpzos | -0.4466 | 0.294 | 0.579 | 1 | 10 |
| Pomp | -0.4466 | 0.854 | 0.913 | 0.756 | 3 |
| Rgs10 | -0.4467 | 0.089 | 0.329 | 1 | 3 |
| Sox5 | -0.4467 | 0.141 | 0.43 | 3.60E-05 | 8 |
| Slc17a6 | -0.4472 | 0 | 0.133 | 0.093 | 8 |
| Ecel1 | -0.4473 | 0.062 | 0.246 | 0.9314 | 6 |
| Cd24a | -0.4473 | 0.454 | 0.491 | 1 | 5 |
| Tvp23b | -0.4475 | 0 | 0.4 | 0.0249 | 10 |
| Celf6 | -0.4476 | 0.467 | 0.732 | 0.2939 | 9 |
| Mir124-2 | -0.4477 | 0 | 0.321 | 0.0028 | 9 |
| Basp1 | -0.4479 | 1 | 0.999 | 0.0002 | 3 |
| Prnp | -0.4483 | 0.979 | 0.998 | 0.0004 | 5 |
| Lhfpl5 | -0.4484 | 0.059 | 0.381 | 1 | 10 |
| Slc17a6 | -0.4485 | 0.008 | 0.139 | 0.0002 | 3 |
| Tenm3 | -0.4487 | 0.289 | 0.577 | 1.16E-08 | 1 |
| Tenm1 | -0.4487 | 0.179 | 0.399 | 0.1257 | 7 |
| Hspe1 | -0.4488 | 0.951 | 0.963 | 0.0206 | 3 |
| Arl1 | -0.4488 | 0.647 | 0.868 | 1 | 10 |
| Sez6l2 | -0.449 | 0.667 | 0.848 | 1 | 9 |
| Rgs4 | -0.4491 | 0.041 | 0.228 | 3.38E-05 | 5 |
| Lamtor1 | -0.4493 | 0.294 | 0.585 | 1 | 10 |

| | | | | | |
|---|---|---|---|---|---|
| Celf2 | -0.4494 | 0.436 | 0.652 | 0.0088 | 7 |
| C1ql3 | -0.4495 | 0.008 | 0.186 | 0.036 | 3 |
| Fam162a | -0.4497 | 0.203 | 0.555 | 1 | 3 |
| Shtn1 | -0.4499 | 0.303 | 0.574 | 7.22E-10 | 1 |
| Macf1 | -0.4501 | 0.658 | 0.828 | 3.43E-05 | 2 |
| Rraga | -0.4503 | 0.647 | 0.887 | 1 | 10 |
| Emc10 | -0.4504 | 0.863 | 0.941 | 0.0195 | 2 |
| Odc1 | -0.4507 | 0.235 | 0.578 | 1 | 10 |
| Cacna2d2 | -0.4508 | 0.267 | 0.556 | 1 | 9 |
| Cox6c | -0.4509 | 0.984 | 0.996 | 0.2817 | 3 |
| Rps14 | -0.4509 | 0.919 | 0.931 | 0.0178 | 3 |
| Ndn | -0.4511 | 0.603 | 0.91 | 0.0323 | 2 |
| Eif4e | -0.4511 | 0.471 | 0.761 | 1 | 10 |
| Higd1a | -0.4514 | 0.62 | 0.8 | 1.62E-06 | 4 |
| Myt1l | -0.4514 | 0.493 | 0.707 | 0.0038 | 8 |
| Cpe | -0.4518 | 0.958 | 0.969 | 0.0146 | 8 |
| Cacna1d | -0.452 | 0.254 | 0.545 | 0.0034 | 8 |
| Timp2 | -0.4522 | 0.815 | 0.905 | 0.0001 | 2 |
| Ephb1 | -0.4523 | 0.067 | 0.371 | 0.1882 | 9 |
| Faim2 | -0.4526 | 0.918 | 0.983 | 2.70E-05 | 2 |
| Rps9 | -0.4527 | 0.984 | 0.973 | 1 | 3 |
| Trpc5 | -0.4527 | 0 | 0.305 | 1.31E-08 | 8 |
| Nrsn2 | -0.4527 | 0.979 | 0.998 | 1.80E-07 | 2 |
| Cplx2 | -0.4535 | 0.008 | 0.302 | 2.10E-08 | 3 |
| Spon1 | -0.4536 | 0.2 | 0.415 | 1 | 9 |
| Epha5 | -0.4538 | 0.732 | 0.858 | 1 | 8 |
| Vat1l | -0.4541 | 0.367 | 0.602 | 1 | 9 |
| Scg5 | -0.4542 | 0.923 | 0.982 | 2.72E-05 | 7 |
| Caly | -0.4542 | 1 | 0.999 | 0.0339 | 5 |
| Stoml2 | -0.4543 | 0.118 | 0.483 | 1 | 10 |
| Irs4 | -0.4544 | 0.064 | 0.287 | 0.0104 | 7 |
| Tpt1 | -0.4546 | 0.992 | 0.987 | 0.0447 | 3 |
| Slc17a6 | -0.4555 | 0.013 | 0.142 | 8.99E-05 | 1 |
| Cck | -0.4557 | 0 | 0.041 | 1 | 6 |
| Gabrg1 | -0.4559 | 0.267 | 0.592 | 1 | 9 |
| Timp2 | -0.4563 | 0.731 | 0.906 | 0.0001 | 7 |
| Slc17a6 | -0.4563 | 0 | 0.136 | 0.0002 | 5 |
| Camk2n2 | -0.4564 | 0.882 | 0.949 | 1 | 10 |
| Atp6v0b | -0.4567 | 1 | 1 | 0.0086 | 2 |
| Erp29 | -0.457 | 0.932 | 0.964 | 0.1936 | 2 |
| Pnrc1 | -0.4575 | 0.467 | 0.741 | 1 | 9 |
| Ppp1r1a | -0.4577 | 0.325 | 0.425 | 1 | 0 |

| | | | | | |
|---|---|---|---|---|---|
| Car10 | -0.4577 | 0.028 | 0.329 | 3.53E-07 | 8 |
| Mpc1 | -0.4579 | 0.471 | 0.815 | 1 | 10 |
| Syf2 | -0.4585 | 0.118 | 0.488 | 0.2416 | 10 |
| Dst | -0.4586 | 0.432 | 0.67 | 1.74E-11 | 2 |
| Irs4 | -0.4599 | 0.103 | 0.297 | 3.19E-14 | 2 |
| Cd83 | -0.4601 | 0.103 | 0.419 | 5.78E-11 | 5 |
| Ppp1r17 | -0.4602 | 0 | 0.313 | 1 | 10 |
| PISD | -0.4603 | 0.667 | 0.846 | 0.0007 | 7 |
| Actb | -0.4603 | 1 | 0.998 | 1 | 10 |
| Caly | -0.4604 | 1 | 0.999 | 0.0002 | 7 |
| Tmem59 | -0.4605 | 0.8 | 0.927 | 1 | 9 |
| Matk | -0.4607 | 0.549 | 0.76 | 0.1491 | 8 |
| Pkig | -0.4609 | 0.294 | 0.634 | 1 | 10 |
| Ube2v2 | -0.4614 | 0.471 | 0.834 | 1 | 10 |
| Deb1 | -0.4616 | 0.176 | 0.525 | 1 | 10 |
| Mir124-2 | -0.4617 | 0 | 0.334 | 1.25E-10 | 8 |
| Snrpb | -0.4623 | 0.353 | 0.75 | 1 | 10 |
| Nebl | -0.4624 | 0.233 | 0.575 | 0.8325 | 9 |
| Pcdh10 | -0.4624 | 0.487 | 0.745 | 0.0007 | 7 |
| Rpl9 | -0.4626 | 0.965 | 0.992 | 3.36E-16 | 0 |
| Clu | -0.4629 | 0.454 | 0.617 | 3.76E-06 | 6 |
| Irs4 | -0.4631 | 0.057 | 0.298 | 0.0018 | 3 |
| Ncald | -0.4633 | 0.618 | 0.687 | 0.0009 | 3 |
| App | -0.4634 | 0.897 | 0.933 | 0.2456 | 5 |
| Ndrg4 | -0.4638 | 0.753 | 0.898 | 8.10E-06 | 6 |
| Ubl5 | -0.4639 | 0.976 | 0.974 | 1 | 3 |
| Ctxn2 | -0.4639 | 0.235 | 0.566 | 1 | 10 |
| Rps3 | -0.4644 | 0.984 | 0.981 | 1 | 3 |
| Gabra1 | -0.4644 | 0.167 | 0.38 | 0.696 | 7 |
| Tro | -0.4651 | 0.801 | 0.94 | 0.0018 | 2 |
| Hypk | -0.4652 | 0.588 | 0.902 | 1 | 10 |
| Sv2a | -0.4655 | 0.789 | 0.934 | 0.4197 | 8 |
| Gap43 | -0.4657 | 0.992 | 0.991 | 0.0002 | 3 |
| Fstl5 | -0.4657 | 0.082 | 0.285 | 0.2489 | 6 |
| Prlr | -0.4659 | 0.013 | 0.233 | 0.0004 | 7 |
| Btg1 | -0.4659 | 0.612 | 0.804 | 1.65E-07 | 1 |
| Pde10a | -0.4664 | 0.057 | 0.33 | 1.59E-05 | 3 |
| Cck | -0.4665 | 0.007 | 0.042 | 1 | 2 |
| Cthrc1 | -0.4666 | 0.12 | 0.412 | 7.06E-12 | 0 |
| Btg1 | -0.4668 | 0.655 | 0.804 | 1.12E-09 | 0 |
| 5330434 | -0.4671 | 0.541 | 0.76 | 0.0001 | 2 |
| Atp5j | -0.4673 | 1 | 0.979 | 0.0315 | 3 |

| | | | | | |
|---|---|---|---|---|---|
| Zbtb20 | -0.4676 | 0.515 | 0.76 | 0.013 | 5 |
| Atp2b1 | -0.4677 | 0.67 | 0.844 | 0.0039 | 5 |
| Impact | -0.4677 | 0.958 | 0.988 | 8.31E-07 | 8 |
| Clstn1 | -0.4682 | 0.731 | 0.858 | 0.0009 | 7 |
| Ptprz1 | -0.4685 | 0.274 | 0.448 | 1.71E-05 | 2 |
| Shtn1 | -0.4686 | 0.267 | 0.545 | 1 | 9 |
| Aldoc | -0.4688 | 0.065 | 0.33 | 0.001 | 3 |
| Ywhae | -0.469 | 1 | 1 | 0.0426 | 3 |
| Ly6h | -0.469 | 1 | 1 | 1 | 10 |
| Galnt16 | -0.4696 | 0.103 | 0.395 | 0.001 | 7 |
| Pebp1 | -0.4697 | 0.992 | 0.999 | 0.0064 | 3 |
| Epha4 | -0.4698 | 0 | 0.344 | 0.0035 | 9 |
| Rgs10 | -0.4698 | 0.144 | 0.327 | 1.49E-13 | 2 |
| Pvrl3 | -0.4701 | 0.115 | 0.423 | 4.39E-15 | 0 |
| Epha5 | -0.4703 | 0.671 | 0.878 | 2.27E-09 | 1 |
| Ppp3ca | -0.4704 | 0.577 | 0.814 | 1.99E-05 | 5 |
| Calm3 | -0.4705 | 0.984 | 0.989 | 1.34E-06 | 3 |
| Stk32a | -0.4705 | 0.13 | 0.471 | 7.22E-16 | 0 |
| Atp1a1 | -0.4707 | 0.377 | 0.629 | 3.04E-07 | 2 |
| Eif3h | -0.4709 | 0.471 | 0.851 | 1 | 10 |
| Marcks | -0.471 | 0.882 | 0.947 | 1 | 10 |
| Tmem35 | -0.471 | 0.205 | 0.395 | 0.2857 | 7 |
| Rap1gap | -0.4713 | 0.128 | 0.437 | 6.18E-05 | 7 |
| Oxct1 | -0.4714 | 0.351 | 0.693 | 2.66E-10 | 5 |
| Chga | -0.4715 | 0.563 | 0.746 | 1 | 8 |
| Tmem179 | -0.4716 | 0.685 | 0.891 | 5.71E-10 | 2 |
| Ostc | -0.4721 | 0.588 | 0.837 | 1 | 10 |
| Chchd7 | -0.4722 | 0.118 | 0.472 | 1 | 10 |
| Ost4 | -0.4723 | 0.235 | 0.576 | 1 | 10 |
| Fam162a | -0.4727 | 0.176 | 0.522 | 1 | 10 |
| Pfdn5 | -0.4729 | 0.992 | 0.959 | 1 | 3 |
| Sv2a | -0.4729 | 0.863 | 0.934 | 8.11E-06 | 2 |
| Pou2f2 | -0.4729 | 0.366 | 0.617 | 0.108 | 8 |
| Lingo1 | -0.473 | 0.183 | 0.556 | 5.22E-06 | 8 |
| Olfm1 | -0.4733 | 0.647 | 0.851 | 1 | 10 |
| Cystm1 | -0.4735 | 0.412 | 0.704 | 1 | 10 |
| Camkv | -0.4737 | 0.247 | 0.506 | 6.78E-08 | 5 |
| Dst | -0.4738 | 0.412 | 0.66 | 2.43E-06 | 5 |
| Ncam1 | -0.474 | 0.856 | 0.961 | 2.18E-05 | 5 |
| Txn2 | -0.4741 | 0.471 | 0.806 | 1 | 10 |
| Cpne2 | -0.4742 | 0.118 | 0.458 | 0.3811 | 10 |
| Slc17a6 | -0.4746 | 0 | 0.143 | 1.23E-11 | 2 |

| | | | | | |
|---|---|---|---|---|---|
| Nxph1 | -0.4749 | 0.102 | 0.41 | 2.03E-07 | 4 |
| Scrn1 | -0.475 | 0.38 | 0.731 | 1.81E-06 | 8 |
| Camk2d | -0.4751 | 0.073 | 0.399 | 1.29E-05 | 3 |
| Parm1 | -0.4754 | 0.278 | 0.62 | 4.57E-08 | 4 |
| Htr2c | -0.4755 | 0.021 | 0.319 | 1.94E-10 | 5 |
| Rpl26 | -0.4758 | 0.967 | 0.972 | 1 | 3 |
| Ppfia2 | -0.4761 | 0.4 | 0.579 | 1 | 9 |
| Ndufa2 | -0.4762 | 0.951 | 0.938 | 0.8358 | 3 |
| Unc5d | -0.4767 | 0.072 | 0.374 | 8.97E-12 | 5 |
| Pld3 | -0.4771 | 0.667 | 0.815 | 1 | 9 |
| Syt11 | -0.4775 | 0.821 | 0.951 | 9.13E-06 | 7 |
| Rpl24 | -0.4776 | 0.992 | 0.999 | 0.0002 | 3 |
| Scn2a1 | -0.4776 | 0.676 | 0.897 | 0.0046 | 8 |
| Gstm7 | -0.4778 | 0.065 | 0.377 | 0.0426 | 3 |
| Atp5k | -0.478 | 0.976 | 0.964 | 1 | 3 |
| Id4 | -0.4785 | 0.423 | 0.658 | 0.0158 | 8 |
| Tmem59l | -0.4785 | 0.842 | 0.933 | 0.4606 | 2 |
| Polr2j | -0.4786 | 0.294 | 0.646 | 1 | 10 |
| Opcml | -0.4787 | 0.41 | 0.696 | 0.0003 | 7 |
| Ndufs5 | -0.4788 | 0.984 | 0.976 | 1 | 3 |
| Chgb | -0.4788 | 0.722 | 0.866 | 0.0405 | 5 |
| Trp53i11 | -0.4789 | 0.593 | 0.738 | 0.0001 | 4 |
| Mettl9 | -0.479 | 0.412 | 0.751 | 1 | 10 |
| mt-Nd3 | -0.4791 | 0.87 | 0.936 | 2.39E-05 | 2 |
| Msi2 | -0.4796 | 0.435 | 0.68 | 1.86E-05 | 4 |
| Rpl39 | -0.4796 | 0.976 | 0.976 | 1 | 3 |
| Smim14 | -0.4797 | 0.353 | 0.763 | 1 | 10 |
| Ebpl | -0.4798 | 0.118 | 0.492 | 1 | 10 |
| Kif1b | -0.48 | 0.726 | 0.899 | 1.01E-11 | 2 |
| Gm1007( | -0.48 | 0.642 | 0.722 | 0.0002 | 3 |
| Romo1 | -0.4802 | 0.959 | 0.962 | 1 | 3 |
| Aplp2 | -0.4804 | 0.526 | 0.75 | 0.0001 | 5 |
| Hspa1a | -0.4806 | 0.067 | 0.195 | 1 | 9 |
| 6330403I | -0.4822 | 1 | 0.999 | 0.0005 | 3 |
| Cfl2 | -0.4823 | 0.471 | 0.819 | 1 | 10 |
| Rgs17 | -0.4825 | 0.381 | 0.72 | 1.57E-09 | 5 |
| Epha5 | -0.4837 | 0.713 | 0.864 | 4.61E-07 | 4 |
| Egr1 | -0.4837 | 0.165 | 0.341 | 0.1617 | 5 |
| Hlf | -0.4845 | 0.235 | 0.528 | 1 | 10 |
| Cox8a | -0.4849 | 1 | 0.998 | 1 | 3 |
| Rps8 | -0.4849 | 1 | 0.998 | 0.0007 | 3 |
| Uqcrb | -0.4852 | 0.967 | 0.978 | 0.2398 | 3 |

| | | | | | |
|---|---|---|---|---|---|
| Galnt16 | -0.4853 | 0.134 | 0.397 | 1.34E-08 | 5 |
| Dpm1 | -0.4861 | 0.353 | 0.72 | 1 | 10 |
| Prlr | -0.4863 | 0 | 0.239 | 2.32E-10 | 5 |
| Gabra1 | -0.4867 | 0.212 | 0.388 | 9.08E-13 | 2 |
| Gabra2 | -0.4869 | 0.296 | 0.625 | 0.0004 | 8 |
| Rorb | -0.4871 | 0.588 | 0.678 | 0.0111 | 6 |
| Ppp1r1a | -0.4875 | 0.274 | 0.428 | 2.44E-05 | 2 |
| Rpl38 | -0.4882 | 0.992 | 0.998 | 1 | 3 |
| Msi2 | -0.4883 | 0.51 | 0.688 | 1.96E-10 | 0 |
| Htr2c | -0.4884 | 0 | 0.298 | 0.5729 | 10 |
| Impact | -0.4885 | 0.882 | 0.987 | 0.0061 | 10 |
| Rps19 | -0.4889 | 0.985 | 0.99 | 1.79E-12 | 0 |
| Gad2 | -0.4895 | 0.742 | 0.826 | 0.1229 | 5 |
| Hmgn3 | -0.4896 | 0.667 | 0.925 | 0.2284 | 9 |
| Tubb5 | -0.4899 | 0.933 | 0.979 | 0.0171 | 9 |
| Rasgrf1 | -0.4899 | 0.592 | 0.823 | 0.0086 | 8 |
| Pcp4 | -0.4903 | 0.679 | 0.895 | 0.0004 | 7 |
| Atp1a1 | -0.4905 | 0.351 | 0.619 | 0.0002 | 5 |
| Gria1 | -0.4905 | 0.381 | 0.589 | 0.0004 | 5 |
| Prlr | -0.4906 | 0.014 | 0.249 | 1.23E-17 | 2 |
| Swi5 | -0.4909 | 0.927 | 0.941 | 1 | 3 |
| Ppp2cb | -0.4909 | 0.353 | 0.659 | 0.7195 | 10 |
| Tenm1 | -0.4913 | 0.144 | 0.406 | 5.84E-09 | 5 |
| Cacna2d1 | -0.4918 | 0.169 | 0.388 | 0.1031 | 8 |
| Sgcz | -0.4919 | 0.014 | 0.338 | 1.58E-08 | 8 |
| Arl6ip5 | -0.4921 | 0.654 | 0.877 | 3.22E-08 | 7 |
| Aplp1 | -0.4928 | 0.973 | 0.985 | 0.016 | 2 |
| Frrs1l | -0.4928 | 0.295 | 0.631 | 9.49E-06 | 7 |
| Tox3 | -0.4928 | 0.125 | 0.419 | 9.24E-14 | 1 |
| Tmem176 | -0.4928 | 0.059 | 0.397 | 1 | 10 |
| Flrt3 | -0.4936 | 0.296 | 0.576 | 2.43E-06 | 4 |
| Fam19a1 | -0.4938 | 0.195 | 0.476 | 9.03E-13 | 0 |
| Ahi1 | -0.4946 | 1 | 1 | 2.69E-21 | 2 |
| Dgkb | -0.4949 | 0.041 | 0.319 | 0.002 | 6 |
| Syp | -0.4951 | 0.761 | 0.948 | 0.0007 | 8 |
| Bsg | -0.4952 | 0.993 | 0.999 | 0.0397 | 2 |
| Thy1 | -0.4952 | 0.299 | 0.432 | 1 | 6 |
| Kit | -0.4953 | 0 | 0.32 | 0.0236 | 9 |
| Rgs10 | -0.4955 | 0.103 | 0.322 | 6.92E-07 | 5 |
| Phpt1 | -0.4956 | 0.235 | 0.598 | 1 | 10 |
| Atp8a1 | -0.4956 | 0.577 | 0.809 | 1.31E-06 | 5 |
| Kit | -0.4959 | 0.014 | 0.332 | 1.50E-07 | 8 |

| | | | | | |
|---|---|---|---|---|---|
| Tspan7 | -0.497 | 0.746 | 0.855 | 1 | 8 |
| Ndufb8 | -0.4972 | 0.647 | 0.978 | 1 | 10 |
| Ap1s2 | -0.4972 | 0.329 | 0.655 | 2.51E-14 | 1 |
| Ptprn | -0.4974 | 0.712 | 0.923 | 3.17E-05 | 2 |
| Brinp1 | -0.4975 | 0.333 | 0.614 | 1 | 9 |
| Alyref | -0.4975 | 0.059 | 0.456 | 1 | 10 |
| Mrpl27 | -0.4976 | 0.176 | 0.603 | 1 | 10 |
| Ier3ip1 | -0.4984 | 0.471 | 0.871 | 1 | 10 |
| Rab18 | -0.4985 | 0.294 | 0.73 | 0.5578 | 10 |
| Ngfrap1 | -0.4988 | 0.992 | 0.993 | 0.0059 | 3 |
| Irs4 | -0.499 | 0.052 | 0.293 | 8.58E-09 | 5 |
| Junb | -0.4992 | 0.216 | 0.347 | 1 | 5 |
| Chst8 | -0.4993 | 0.014 | 0.325 | 5.38E-07 | 8 |
| Cd83 | -0.4994 | 0.033 | 0.401 | 0.0157 | 9 |
| Ube2e2 | -0.5001 | 0.412 | 0.887 | 1 | 10 |
| Pam | -0.5002 | 0.495 | 0.705 | 0.0084 | 5 |
| Cox5b | -0.5003 | 0.976 | 0.991 | 0.0315 | 3 |
| Atp8a1 | -0.5005 | 0.667 | 0.792 | 1 | 9 |
| Fau | -0.5006 | 1 | 0.997 | 1 | 3 |
| Rps10 | -0.5011 | 0.976 | 0.985 | 0.1352 | 3 |
| Lrp1b | -0.5013 | 0.381 | 0.669 | 6.82E-05 | 5 |
| Ucp2 | -0.5015 | 0.033 | 0.338 | 0.0201 | 9 |
| Ttc3 | -0.5017 | 1 | 1 | 5.85E-15 | 2 |
| Rps7 | -0.502 | 0.984 | 0.98 | 1 | 3 |
| Atp2b1 | -0.5024 | 0.69 | 0.839 | 0.7212 | 8 |
| Cfl1 | -0.503 | 1 | 0.99 | 0.0002 | 3 |
| Thsd7b | -0.5031 | 0.033 | 0.369 | 0.0481 | 9 |
| Malat1 | -0.5031 | 1 | 0.999 | 2.12E-10 | 0 |
| Ajap1 | -0.5034 | 0.2 | 0.444 | 1 | 9 |
| Rpl34 | -0.5041 | 0.976 | 0.973 | 1 | 3 |
| Clstn1 | -0.5042 | 0.74 | 0.865 | 1.43E-05 | 2 |
| Nrp1 | -0.5045 | 0 | 0.364 | 0.6187 | 10 |
| Tmem158 | -0.5048 | 0.548 | 0.738 | 3.78E-10 | 2 |
| Tsc22d1 | -0.5049 | 0.854 | 0.863 | 6.48E-12 | 3 |
| Map1b | -0.5052 | 0.945 | 0.984 | 7.32E-11 | 2 |
| Kcnk2 | -0.5053 | 0.158 | 0.392 | 1.34E-13 | 2 |
| Kit | -0.5054 | 0.019 | 0.343 | 1.87E-13 | 4 |
| Spock1 | -0.5063 | 0.128 | 0.42 | 0.0008 | 7 |
| Rps15a | -0.5065 | 0.984 | 0.982 | 0.3949 | 3 |
| Rps20 | -0.507 | 0.984 | 0.978 | 0.2559 | 3 |
| Cartpt | -0.5071 | 0.085 | 0.136 | 1 | 0 |
| Tspan31 | -0.5075 | 0.118 | 0.525 | 1 | 10 |

| | | | | | |
|---|---|---|---|---|---|
| Mrpl30 | -0.508 | 0.235 | 0.677 | 1 | 10 |
| Peg10 | -0.5081 | 0.1 | 0.331 | 1 | 9 |
| Rsrc2 | -0.5081 | 0.529 | 0.823 | 1 | 10 |
| Serinc1 | -0.509 | 0.959 | 0.968 | 1.42E-06 | 2 |
| Mrpl33 | -0.509 | 0.353 | 0.722 | 1 | 10 |
| Chchd10 | -0.5094 | 0.951 | 0.923 | 0.1955 | 3 |
| Ndufc2 | -0.5094 | 1 | 0.995 | 1 | 10 |
| mt-Co3 | -0.5094 | 1 | 1 | 2.27E-12 | 2 |
| Rpl37 | -0.5099 | 1 | 0.996 | 1 | 3 |
| Ldha | -0.5099 | 0.526 | 0.778 | 2.20E-07 | 7 |
| Tmx4 | -0.5104 | 0.966 | 0.987 | 2.29E-07 | 2 |
| Aldoc | -0.5104 | 0.038 | 0.321 | 1.46E-06 | 7 |
| Rpl6 | -0.511 | 1 | 0.998 | 0.012 | 3 |
| Rpl22l1 | -0.5122 | 1 | 0.997 | 0.7414 | 3 |
| Smim19 | -0.5123 | 0.412 | 0.73 | 1 | 10 |
| Rgs4 | -0.5126 | 0 | 0.238 | 4.51E-06 | 3 |
| Iqsec3 | -0.5133 | 0.169 | 0.519 | 4.10E-06 | 8 |
| Cntn1 | -0.5134 | 0.438 | 0.663 | 8.26E-14 | 2 |
| Tecr | -0.5135 | 0.959 | 0.997 | 2.14E-05 | 5 |
| Tmem35 | -0.514 | 0.093 | 0.412 | 2.81E-08 | 4 |
| Syngr3 | -0.5142 | 0.801 | 0.924 | 2.99E-06 | 2 |
| Rgs4 | -0.5146 | 0.007 | 0.243 | 1.67E-20 | 2 |
| Uqcrq | -0.5148 | 0.984 | 0.968 | 1 | 3 |
| Churc1 | -0.5151 | 0.118 | 0.624 | 1 | 10 |
| Arl4a | -0.5153 | 0.235 | 0.56 | 1 | 10 |
| Ctsb | -0.5154 | 0.815 | 0.939 | 3.09E-06 | 2 |
| Gabrg1 | -0.5156 | 0.336 | 0.623 | 3.48E-11 | 1 |
| Atp5e | -0.5157 | 0.967 | 0.97 | 1 | 3 |
| Grin2b | -0.516 | 0.452 | 0.707 | 7.18E-14 | 2 |
| Syt4 | -0.5161 | 0.691 | 0.905 | 2.20E-12 | 1 |
| Morf4l1 | -0.5162 | 1 | 0.998 | 1 | 10 |
| Prkce | -0.5163 | 0.31 | 0.592 | 7.67E-05 | 8 |
| Rpl27 | -0.5178 | 0.706 | 0.922 | 1 | 10 |
| Kif1a | -0.5183 | 0.781 | 0.937 | 3.04E-09 | 2 |
| Syt1 | -0.519 | 0.732 | 0.808 | 0.0001 | 6 |
| Cox7a2 | -0.5194 | 0.984 | 0.98 | 0.2239 | 3 |
| Pvrl3 | -0.5196 | 0.07 | 0.388 | 3.23E-06 | 8 |
| Unc5c | -0.5197 | 0.178 | 0.489 | 1.13E-14 | 1 |
| Tram1l1 | -0.52 | 0.412 | 0.701 | 1 | 10 |
| Vstm2l | -0.5201 | 0.697 | 0.868 | 1.29E-08 | 1 |
| Ptprz1 | -0.5203 | 0.206 | 0.446 | 6.25E-05 | 5 |
| Rpl37a | -0.5204 | 1 | 0.998 | 1 | 3 |

| | | | | | |
|---|---|---|---|---|---|
| Pgrmc1 | -0.5216 | 1 | 0.999 | 2.70E-20 | 1 |
| Ict1 | -0.5217 | 0.059 | 0.487 | 0.1789 | 10 |
| Rpl41 | -0.5219 | 1 | 1 | 0.1529 | 3 |
| 1500009l | -0.5222 | 0.235 | 0.596 | 1 | 10 |
| Atp2b2 | -0.5232 | 0.408 | 0.749 | 0.0004 | 8 |
| Rps16 | -0.5234 | 0.984 | 0.985 | 0.1401 | 3 |
| Pcdh19 | -0.5237 | 0.224 | 0.574 | 8.93E-13 | 1 |
| Negr1 | -0.5242 | 0.699 | 0.857 | 1.55E-09 | 2 |
| Dpp6 | -0.5243 | 0.692 | 0.867 | 2.92E-07 | 2 |
| Mrpl17 | -0.5245 | 0.294 | 0.667 | 1 | 10 |
| Cntn1 | -0.5254 | 0.372 | 0.653 | 3.22E-05 | 7 |
| Aldoc | -0.5255 | 0.031 | 0.327 | 6.57E-13 | 5 |
| Mrpl34 | -0.5262 | 0.235 | 0.598 | 1 | 10 |
| Rps11 | -0.5262 | 0.992 | 0.995 | 0.0001 | 3 |
| Ecel1 | -0.5267 | 0.041 | 0.253 | 0.0004 | 3 |
| Prnp | -0.5269 | 1 | 0.996 | 8.21E-08 | 7 |
| Rer1 | -0.5269 | 0.353 | 0.729 | 1 | 10 |
| Drap1 | -0.5272 | 0.529 | 0.903 | 1 | 10 |
| Nap1l5 | -0.5273 | 0.992 | 1 | 0.1038 | 3 |
| Araf | -0.5277 | 0.529 | 0.927 | 0.0594 | 10 |
| Id4 | -0.5279 | 0.433 | 0.649 | 1 | 9 |
| Fam19a1 | -0.5296 | 0.169 | 0.443 | 0.0069 | 8 |
| Sez6l | -0.5297 | 0.372 | 0.622 | 2.76E-05 | 7 |
| Rps4x | -0.53 | 0.992 | 0.981 | 1 | 3 |
| Npdc1 | -0.5302 | 0.877 | 0.963 | 7.41E-07 | 2 |
| Calm1 | -0.5305 | 1 | 1 | 0.0575 | 3 |
| Ran | -0.5315 | 0.941 | 0.987 | 1 | 10 |
| Rpl27a | -0.5319 | 0.992 | 0.995 | 0.062 | 3 |
| Rheb | -0.532 | 0.824 | 0.912 | 1 | 10 |
| Tsg101 | -0.5321 | 0.412 | 0.767 | 1 | 10 |
| Tmem256 | -0.5322 | 0.294 | 0.679 | 1 | 10 |
| Rtn3 | -0.5323 | 0.901 | 0.984 | 1.05E-05 | 8 |
| Naca | -0.5329 | 0.992 | 0.985 | 0.2966 | 3 |
| Fos | -0.5331 | 0.385 | 0.44 | 1 | 0 |
| Kif5c | -0.5334 | 0.582 | 0.841 | 8.88E-12 | 2 |
| Serp2 | -0.5336 | 0.829 | 0.895 | 1.57E-08 | 3 |
| Cdh2 | -0.5339 | 0.433 | 0.785 | 1 | 9 |
| Npas4 | -0.5346 | 0.082 | 0.251 | 0.0984 | 5 |
| Chchd4 | -0.5347 | 0.412 | 0.764 | 1 | 10 |
| Cbarp | -0.5355 | 0.8 | 0.969 | 0.3209 | 9 |
| Cst3 | -0.5355 | 1 | 0.988 | 1 | 2 |
| Cthrc1 | -0.5368 | 0.125 | 0.397 | 2.57E-11 | 1 |

| | | | | | |
|---|---|---|---|---|---|
| Iscu | -0.537 | 0.471 | 0.862 | 1 | 10 |
| Egr1 | -0.537 | 0.185 | 0.346 | 0.0014 | 2 |
| Vdac2 | -0.5372 | 0.706 | 0.944 | 1 | 10 |
| Atp5g3 | -0.5372 | 0.984 | 0.977 | 0.0014 | 3 |
| Cops6 | -0.5375 | 0.294 | 0.725 | 1 | 10 |
| Pou2f2 | -0.5375 | 0.375 | 0.637 | 8.34E-10 | 1 |
| Chmp2a | -0.5375 | 0.353 | 0.77 | 1 | 10 |
| Chgb | -0.5376 | 0.733 | 0.857 | 1 | 9 |
| Cuta | -0.5394 | 0.353 | 0.84 | 1 | 10 |
| Flrt3 | -0.5398 | 0.353 | 0.552 | 1 | 10 |
| Vbp1 | -0.5415 | 0.176 | 0.575 | 0.1425 | 10 |
| Atp1b1 | -0.5416 | 0.993 | 0.999 | 2.72E-13 | 2 |
| Cacna1d | -0.5423 | 0.233 | 0.534 | 0.5962 | 9 |
| Aldoc | -0.5424 | 0.034 | 0.341 | 1.41E-28 | 2 |
| Rps23 | -0.5431 | 0.967 | 0.972 | 0.874 | 3 |
| Ajap1 | -0.5432 | 0.145 | 0.484 | 1.08E-14 | 1 |
| Rprm | -0.5439 | 0.21 | 0.457 | 5.33E-09 | 0 |
| Tmem255 | -0.5446 | 0.216 | 0.588 | 6.03E-11 | 5 |
| Cycs | -0.5461 | 0.902 | 0.93 | 0.0018 | 3 |
| Fam195b | -0.5461 | 0.294 | 0.796 | 1 | 10 |
| Lin7a | -0.5463 | 0.505 | 0.725 | 1.46E-14 | 0 |
| Hsp90b1 | -0.5464 | 0.959 | 0.985 | 0.0015 | 2 |
| Dusp1 | -0.5469 | 0.155 | 0.301 | 1 | 5 |
| Cbln2 | -0.5469 | 0.052 | 0.292 | 7.85E-07 | 5 |
| Timm13 | -0.5471 | 0.529 | 0.915 | 1 | 10 |
| Ldha | -0.5471 | 0.527 | 0.795 | 1.95E-12 | 2 |
| Gnb1 | -0.5477 | 0.833 | 0.942 | 0.0543 | 9 |
| Ntrk3 | -0.5478 | 0.169 | 0.549 | 1.15E-06 | 8 |
| Cd63 | -0.5487 | 0.118 | 0.469 | 1 | 10 |
| Vstm2l | -0.5492 | 0.796 | 0.85 | 5.13E-05 | 4 |
| Prnp | -0.55 | 0.993 | 0.997 | 3.81E-06 | 2 |
| Per3 | -0.5502 | 0.353 | 0.59 | 1 | 10 |
| Snrpg | -0.5505 | 0.471 | 0.814 | 1 | 10 |
| Tmed4 | -0.5509 | 0.412 | 0.777 | 1 | 10 |
| Vimp | -0.5515 | 0.176 | 0.613 | 1 | 10 |
| Cd47 | -0.552 | 0.7 | 0.879 | 1 | 9 |
| Commd3 | -0.5525 | 0.294 | 0.672 | 1 | 10 |
| Uqcrh | -0.5526 | 0.967 | 0.947 | 0.0116 | 3 |
| Cd47 | -0.5527 | 0.62 | 0.891 | 7.01E-06 | 8 |
| Cthrc1 | -0.5528 | 0.1 | 0.367 | 0.9565 | 9 |
| Cetn2 | -0.5532 | 0.412 | 0.759 | 1 | 10 |
| Pvrl3 | -0.5534 | 0.033 | 0.377 | 0.0233 | 9 |

| | | | | | |
|---|---|---|---|---|---|
| Rprm | -0.5537 | 0.222 | 0.433 | 0.0008 | 4 |
| Ntm | -0.5537 | 0.628 | 0.824 | 9.94E-06 | 7 |
| Nrp2 | -0.5539 | 0.118 | 0.493 | 1 | 10 |
| Fkbp2 | -0.5541 | 0.822 | 0.946 | 5.36E-10 | 2 |
| Rps21 | -0.5542 | 0.992 | 0.985 | 1 | 3 |
| Trmt112 | -0.5542 | 0.647 | 0.928 | 1 | 10 |
| Fkbp3 | -0.5542 | 0.976 | 0.968 | 0.0001 | 3 |
| Ccdc85b | -0.5545 | 0.235 | 0.637 | 1 | 10 |
| Btg2 | -0.5557 | 0.144 | 0.312 | 0.0393 | 2 |
| Ndrg4 | -0.5559 | 0.676 | 0.9 | 3.14E-07 | 8 |
| Pbx1 | -0.5564 | 0.667 | 0.766 | 1 | 9 |
| Erdr1 | -0.5566 | 0.7 | 0.912 | 1 | 9 |
| Rpl22 | -0.5572 | 0.294 | 0.695 | 1 | 10 |
| Cthrc1 | -0.5573 | 0.099 | 0.378 | 4.11E-05 | 8 |
| Timp2 | -0.5584 | 0.763 | 0.906 | 4.31E-08 | 5 |
| Unc5c | -0.5595 | 0.067 | 0.457 | 0.0187 | 9 |
| Rtn4 | -0.5595 | 0.877 | 0.959 | 4.13E-07 | 2 |
| Erdr1 | -0.56 | 0.761 | 0.916 | 0.1437 | 8 |
| App | -0.5602 | 0.859 | 0.936 | 0.0001 | 7 |
| Eif1 | -0.5605 | 1 | 0.998 | 0.0031 | 3 |
| Gabra1 | -0.5605 | 0.124 | 0.388 | 4.69E-09 | 5 |
| Rpl21 | -0.5607 | 0.992 | 0.998 | 0.1342 | 3 |
| Mrps24 | -0.5611 | 0.353 | 0.711 | 1 | 10 |
| Rps27 | -0.5615 | 1 | 0.992 | 1 | 3 |
| Scg5 | -0.5617 | 0.938 | 0.984 | 0.0208 | 2 |
| Rsrp1 | -0.5619 | 0.867 | 0.978 | 0.007 | 9 |
| Fstl5 | -0.5621 | 0.041 | 0.295 | 1.85E-08 | 3 |
| Slc12a5 | -0.5636 | 0.761 | 0.915 | 0.028 | 8 |
| Trp53i11 | -0.5645 | 0.487 | 0.761 | 1.77E-12 | 1 |
| Bod1 | -0.5649 | 0.412 | 0.78 | 1 | 10 |
| Rprm | -0.5655 | 0.206 | 0.432 | 0.0002 | 5 |
| Cspg5 | -0.5666 | 0.692 | 0.917 | 3.96E-11 | 7 |
| Arhgap36 | -0.5666 | 0 | 0.215 | 1.34E-08 | 5 |
| Cbln2 | -0.5666 | 0 | 0.275 | 1 | 10 |
| Rpl19 | -0.5667 | 1 | 0.999 | 0.0394 | 3 |
| Dgkb | -0.5671 | 0.021 | 0.321 | 2.16E-13 | 5 |
| Rpl29 | -0.5691 | 0.647 | 0.945 | 1 | 10 |
| Tm2d2 | -0.5694 | 0.353 | 0.835 | 1 | 10 |
| Zbtb20 | -0.5694 | 0.61 | 0.758 | 8.30E-06 | 2 |
| Mt3 | -0.5696 | 0.825 | 0.977 | 7.73E-13 | 5 |
| Ntng1 | -0.5705 | 0.175 | 0.362 | 0.1358 | 6 |
| Rab3c | -0.5711 | 0.664 | 0.912 | 1.83E-19 | 1 |

| | | | | | |
|---|---|---|---|---|---|
| Prkacb | -0.5713 | 0.835 | 0.961 | 3.25E-16 | 5 |
| Syt11 | -0.5714 | 0.866 | 0.949 | 1.69E-08 | 5 |
| Stk32a | -0.5715 | 0.067 | 0.42 | 0.0092 | 9 |
| mt-Nd1 | -0.5717 | 1 | 1 | 1.69E-07 | 2 |
| Cbln2 | -0.5717 | 0 | 0.278 | 0.223 | 9 |
| Cntn1 | -0.5721 | 0.371 | 0.659 | 5.90E-09 | 5 |
| Txn1 | -0.5726 | 0.412 | 0.823 | 1 | 10 |
| Nono | -0.5733 | 0.353 | 0.71 | 1 | 10 |
| Calm2 | -0.5737 | 1 | 1 | 0.0063 | 3 |
| Nrsn2 | -0.5738 | 1 | 0.995 | 1 | 10 |
| Fam174a | -0.5738 | 0.412 | 0.865 | 0.5115 | 10 |
| Ncald | -0.5747 | 0.486 | 0.708 | 1.08E-12 | 2 |
| Kcnk2 | -0.5755 | 0.062 | 0.389 | 1.64E-11 | 5 |
| Fstl5 | -0.5757 | 0.062 | 0.298 | 5.57E-16 | 2 |
| Psenen | -0.5757 | 0.412 | 0.848 | 1 | 10 |
| Vti1b | -0.576 | 0.529 | 0.856 | 1 | 10 |
| Isl1 | -0.5761 | 0 | 0.265 | 1.68E-07 | 7 |
| Ndufc1 | -0.5762 | 0.959 | 0.94 | 0.0011 | 3 |
| Denr | -0.5764 | 0.353 | 0.785 | 1 | 10 |
| Msi2 | -0.5777 | 0.447 | 0.689 | 2.49E-11 | 1 |
| Rit2 | -0.5778 | 0.684 | 0.919 | 2.14E-22 | 1 |
| Pou2f2 | -0.5779 | 0.395 | 0.646 | 1.18E-16 | 0 |
| Cadps2 | -0.579 | 0.38 | 0.59 | 1.37E-12 | 0 |
| Ar | -0.5792 | 0.028 | 0.437 | 1.85E-12 | 8 |
| Nol7 | -0.5795 | 0.235 | 0.63 | 1 | 10 |
| Rpl13 | -0.5798 | 1 | 0.996 | 2.52E-06 | 3 |
| Mdh1 | -0.5801 | 0.824 | 0.987 | 1 | 10 |
| Ptn | -0.5803 | 0.34 | 0.526 | 0.015 | 5 |
| Dgkb | -0.5804 | 0 | 0.317 | 1.76E-09 | 7 |
| Tspan7 | -0.5817 | 0.782 | 0.853 | 0.0006 | 7 |
| Bub3 | -0.5819 | 0.294 | 0.732 | 1 | 10 |
| Atp5l | -0.5824 | 0.984 | 0.974 | 0.1062 | 3 |
| 2900011 | -0.5827 | 0.471 | 0.897 | 1 | 10 |
| Id4 | -0.5829 | 0.398 | 0.67 | 1.26E-07 | 4 |
| Id4 | -0.5841 | 0.375 | 0.686 | 4.17E-13 | 1 |
| Isl1 | -0.5842 | 0 | 0.27 | 6.84E-12 | 5 |
| Rgs10 | -0.5845 | 0 | 0.308 | 0.0741 | 10 |
| Ngb | -0.5846 | 0.564 | 0.794 | 3.77E-06 | 7 |
| Grin2b | -0.5854 | 0.385 | 0.695 | 1.25E-08 | 7 |
| Pnmal2 | -0.5855 | 0.944 | 0.996 | 1.62E-09 | 8 |
| Slc6a1 | -0.5858 | 0.296 | 0.673 | 6.94E-08 | 8 |
| Timm17a | -0.5859 | 0.471 | 0.826 | 1 | 10 |

| | | | | | |
|---|---|---|---|---|---|
| Trp53i11 | -0.5861 | 0.465 | 0.741 | 0.0002 | 8 |
| Ajap1 | -0.587 | 0.099 | 0.461 | 8.23E-07 | 8 |
| Isl1 | -0.587 | 0.008 | 0.276 | 9.01E-10 | 3 |
| Uqcr11 | -0.587 | 0.943 | 0.974 | 3.24E-05 | 3 |
| Vsnl1 | -0.5871 | 0.612 | 0.869 | 4.23E-21 | 1 |
| Hint1 | -0.5873 | 0.992 | 0.987 | 0.0133 | 3 |
| Cbln2 | -0.5886 | 0 | 0.289 | 3.80E-07 | 8 |
| Dgkb | -0.5887 | 0.021 | 0.336 | 1.40E-28 | 2 |
| Fam171b | -0.589 | 0.616 | 0.843 | 2.13E-12 | 2 |
| H3f3a | -0.5905 | 0.706 | 0.956 | 1 | 10 |
| Pcdh19 | -0.591 | 0.155 | 0.552 | 1.00E-06 | 8 |
| Dgkb | -0.5914 | 0.008 | 0.33 | 1.53E-11 | 3 |
| Syt13 | -0.5925 | 0.338 | 0.693 | 2.88E-07 | 8 |
| Lamtor2 | -0.5929 | 0.588 | 0.896 | 1 | 10 |
| Cox7c | -0.5941 | 1 | 0.99 | 0.6029 | 3 |
| Pcdh7 | -0.5951 | 0.141 | 0.502 | 1.02E-06 | 7 |
| Shtn1 | -0.5951 | 0.141 | 0.564 | 2.54E-09 | 8 |
| Timm10b | -0.5952 | 0.176 | 0.636 | 1 | 10 |
| Psap | -0.5953 | 0.945 | 0.967 | 3.17E-06 | 2 |
| Fstl5 | -0.5958 | 0.013 | 0.286 | 4.29E-06 | 7 |
| Caly | -0.5959 | 1 | 0.999 | 5.38E-08 | 2 |
| Pkib | -0.5963 | 0.233 | 0.475 | 1 | 9 |
| Cbln2 | -0.597 | 0.02 | 0.31 | 2.06E-15 | 1 |
| Clu | -0.5971 | 0.381 | 0.624 | 6.38E-05 | 5 |
| Gaa | -0.5972 | 0.966 | 0.975 | 2.66E-05 | 2 |
| Cd81 | -0.5972 | 0.925 | 0.978 | 3.57E-05 | 2 |
| Auts2 | -0.5973 | 0.463 | 0.768 | 1.40E-12 | 4 |
| Rps19 | -0.5987 | 1 | 0.988 | 1 | 3 |
| Gria1 | -0.5987 | 0.244 | 0.596 | 3.62E-07 | 7 |
| Nrgn | -0.5988 | 0.103 | 0.246 | 1 | 7 |
| Ppp1r2 | -0.599 | 0.235 | 0.727 | 0.2133 | 10 |
| Prdx2 | -0.599 | 0.824 | 0.989 | 1 | 10 |
| Ndufab1 | -0.5994 | 0.824 | 0.965 | 1 | 10 |
| Tmem13( | -0.5996 | 0.925 | 0.983 | 2.89E-11 | 2 |
| Tagln3 | -0.5997 | 0.927 | 0.949 | 5.40E-10 | 3 |
| 1810037I | -0.5997 | 0.353 | 0.779 | 0.7729 | 10 |
| Penk | -0.6002 | 0.067 | 0.342 | 1 | 9 |
| Rpl3 | -0.6002 | 1 | 1 | 0.0047 | 3 |
| Pbx1 | -0.6008 | 0.574 | 0.783 | 3.48E-09 | 4 |
| Scg2 | -0.601 | 0.925 | 0.97 | 2.57E-18 | 0 |
| Fos | -0.6014 | 0.536 | 0.42 | 1 | 6 |
| Atp6ap2 | -0.602 | 0.633 | 0.841 | 1 | 9 |

| | | | | | |
|---|---|---|---|---|---|
| Ube2e1 | -0.6022 | 0.412 | 0.769 | 1 | 10 |
| Fam171b | -0.6025 | 0.567 | 0.837 | 4.24E-10 | 5 |
| Cxxc4 | -0.6029 | 0.267 | 0.691 | 0.0014 | 9 |
| Tenm3 | -0.603 | 0.183 | 0.562 | 3.95E-07 | 8 |
| Slc22a17 | -0.6034 | 0.887 | 0.974 | 3.09E-09 | 5 |
| Flrt3 | -0.6052 | 0.243 | 0.597 | 4.80E-16 | 1 |
| Ptprz1 | -0.6052 | 0.154 | 0.446 | 3.24E-05 | 7 |
| Emc6 | -0.6054 | 0.353 | 0.789 | 1 | 10 |
| Isl1 | -0.6062 | 0 | 0.284 | 3.11E-24 | 2 |
| Fstl5 | -0.6067 | 0.01 | 0.292 | 4.79E-12 | 5 |
| Lsm4 | -0.6068 | 0.353 | 0.761 | 1 | 10 |
| Six3 | -0.6076 | 0.63 | 0.694 | 8.00E-06 | 0 |
| Synpr | -0.6077 | 0.711 | 0.717 | 1 | 5 |
| Rpl23 | -0.6087 | 1 | 0.985 | 5.20E-06 | 3 |
| Chchd2 | -0.6088 | 1 | 0.998 | 0.0098 | 3 |
| Msi2 | -0.6117 | 0.352 | 0.677 | 2.77E-05 | 8 |
| Arhgdig | -0.6125 | 0.495 | 0.857 | 2.05E-13 | 5 |
| Rprm | -0.6125 | 0.183 | 0.428 | 0.0046 | 8 |
| Ssu72 | -0.6128 | 0.353 | 0.774 | 1 | 10 |
| Ntng1 | -0.6141 | 0.187 | 0.365 | 0.0006 | 3 |
| Sox2 | -0.6145 | 0.183 | 0.604 | 6.25E-10 | 8 |
| Minos1 | -0.6166 | 0.529 | 0.919 | 1 | 10 |
| Coa3 | -0.6175 | 0.529 | 0.94 | 1 | 10 |
| Rab3c | -0.6176 | 0.633 | 0.885 | 0.031 | 9 |
| Ptms | -0.6184 | 0.898 | 0.979 | 1.38E-15 | 4 |
| Snap25 | -0.6186 | 0.9 | 0.923 | 1 | 9 |
| mt-Nd5 | -0.6188 | 0.952 | 0.983 | 7.74E-09 | 2 |
| Cdh2 | -0.619 | 0.465 | 0.797 | 2.62E-07 | 8 |
| Syt1 | -0.6193 | 0.943 | 0.784 | 4.87E-08 | 3 |
| Sumo2 | -0.6223 | 0.941 | 0.994 | 1 | 10 |
| Cst3 | -0.6226 | 0.933 | 0.991 | 1 | 9 |
| Hpcal1 | -0.6228 | 0.34 | 0.444 | 1 | 6 |
| Rpl36al | -0.6235 | 0.706 | 0.956 | 1 | 10 |
| Rpl9 | -0.6243 | 0.992 | 0.987 | 0.8191 | 3 |
| Thy1 | -0.6244 | 0.195 | 0.449 | 7.66E-09 | 3 |
| Jun | -0.6265 | 0.485 | 0.566 | 0.0233 | 0 |
| Lsm6 | -0.6265 | 0.294 | 0.753 | 1 | 10 |
| Bnip3l | -0.6267 | 0.176 | 0.737 | 0.2941 | 10 |
| Lin7a | -0.6269 | 0.417 | 0.714 | 1.22E-10 | 4 |
| Bex2 | -0.6271 | 1 | 1 | 0.0003 | 3 |
| D8Ertd73 | -0.6282 | 0.412 | 0.887 | 1 | 10 |
| Resp18 | -0.6289 | 1 | 0.999 | 0.0034 | 7 |

| | | | | | |
|---|---|---|---|---|---|
| Tle4 | -0.6291 | 0.353 | 0.675 | 1 | 10 |
| Opcml | -0.6293 | 0.418 | 0.715 | 1.27E-15 | 2 |
| Pcdh19 | -0.6297 | 0.133 | 0.537 | 0.0435 | 9 |
| 2700094[illegible] | -0.6297 | 0.235 | 0.684 | 1 | 10 |
| Mrfap1 | -0.63 | 0.984 | 0.995 | 7.46E-09 | 3 |
| Stk32a | -0.6314 | 0 | 0.417 | 0.7711 | 10 |
| Bsg | -0.6315 | 0.941 | 0.999 | 1 | 10 |
| Rorb | -0.6316 | 0.593 | 0.68 | 4.19E-09 | 3 |
| Cacybp | -0.6328 | 0.706 | 0.932 | 1 | 10 |
| Rora | -0.6333 | 0.167 | 0.48 | 6.55E-09 | 4 |
| Pcsk1n | -0.6337 | 1 | 1 | 2.72E-19 | 5 |
| B3gat2 | -0.6338 | 0 | 0.467 | 0.025 | 10 |
| Ecel1 | -0.6343 | 0.01 | 0.25 | 2.07E-08 | 5 |
| Bri3 | -0.6352 | 0.353 | 0.831 | 1 | 10 |
| Tpt1 | -0.6356 | 0.882 | 0.989 | 1 | 10 |
| Tenm3 | -0.6356 | 0.133 | 0.549 | 0.0307 | 9 |
| Arhgap6 | -0.6358 | 0.287 | 0.64 | 9.74E-12 | 4 |
| Hspe1 | -0.6361 | 0.765 | 0.965 | 1 | 10 |
| Ecel1 | -0.6363 | 0 | 0.247 | 6.77E-07 | 7 |
| Ppp3ca | -0.6367 | 0.526 | 0.814 | 1.21E-08 | 7 |
| Sec61b | -0.6369 | 0.235 | 0.786 | 1 | 10 |
| Spon1 | -0.6376 | 0.059 | 0.464 | 4.23E-23 | 1 |
| App | -0.6381 | 0.87 | 0.939 | 2.45E-09 | 2 |
| St13 | -0.6386 | 0.353 | 0.815 | 0.0221 | 10 |
| BC03118[illegible] | -0.6387 | 0.706 | 0.975 | 1 | 10 |
| Cthrc1 | -0.6439 | 0 | 0.366 | 0.2789 | 10 |
| Rpl30 | -0.6439 | 1 | 0.988 | 1.52E-05 | 3 |
| Pmm1 | -0.644 | 0.412 | 0.879 | 1 | 10 |
| Pcbd1 | -0.6443 | 0.33 | 0.758 | 1.09E-15 | 5 |
| Mrpl54 | -0.6444 | 0.235 | 0.69 | 1 | 10 |
| Sepw1 | -0.6449 | 0.992 | 0.996 | 6.81E-05 | 3 |
| Tenm4 | -0.6452 | 0.183 | 0.552 | 4.68E-07 | 8 |
| Map1b | -0.6465 | 0.928 | 0.983 | 1.68E-16 | 5 |
| 2010107[illegible] | -0.6467 | 0.647 | 0.934 | 1 | 10 |
| Nop10 | -0.6482 | 0.647 | 0.956 | 1 | 10 |
| Ssb | -0.65 | 0.353 | 0.897 | 1 | 10 |
| Prdx1 | -0.65 | 0.647 | 0.966 | 1 | 10 |
| Btg2 | -0.6501 | 0.082 | 0.31 | 0.0006 | 5 |
| Npm1 | -0.6506 | 0.824 | 0.982 | 1 | 10 |
| Alcam | -0.6509 | 0.471 | 0.804 | 1 | 10 |
| Fos | -0.6516 | 0.133 | 0.438 | 1 | 9 |
| Syt11 | -0.6528 | 0.877 | 0.952 | 1.29E-13 | 2 |

| | | | | | |
|---|---|---|---|---|---|
| Polr2k | -0.653 | 0.412 | 0.799 | 0.3879 | 10 |
| Ptprn | -0.653 | 0.533 | 0.905 | 0.0985 | 9 |
| Ecel1 | -0.654 | 0.014 | 0.262 | 4.56E-16 | 2 |
| Tmco1 | -0.657 | 0.353 | 0.776 | 0.2625 | 10 |
| Rpl18a | -0.6573 | 0.882 | 0.992 | 1 | 10 |
| Uqcrfs1 | -0.6587 | 0.471 | 0.844 | 1 | 10 |
| Tle4 | -0.6614 | 0.375 | 0.717 | 6.68E-19 | 1 |
| Rora | -0.6616 | 0.167 | 0.457 | 0.5893 | 9 |
| Pomp | -0.6626 | 0.471 | 0.913 | 1 | 10 |
| Pou2f2 | -0.6629 | 0.269 | 0.637 | 6.98E-13 | 4 |
| Edil3 | -0.6631 | 0.3 | 0.667 | 0.0999 | 9 |
| Pbx1 | -0.6631 | 0.521 | 0.78 | 4.96E-05 | 8 |
| Arhgap6 | -0.6638 | 0.282 | 0.628 | 2.38E-06 | 8 |
| Gnai2 | -0.6641 | 0.824 | 0.975 | 1 | 10 |
| Id4 | -0.6641 | 0.415 | 0.693 | 3.42E-20 | 0 |
| Msi2 | -0.6643 | 0.294 | 0.662 | 1 | 10 |
| Trappc2l | -0.6643 | 0.471 | 0.877 | 1 | 10 |
| Per3 | -0.6653 | 0.167 | 0.598 | 0.043 | 9 |
| Tecr | -0.6664 | 0.993 | 0.994 | 1.43E-11 | 2 |
| Coprs | -0.6676 | 0.176 | 0.716 | 1 | 10 |
| Ntan1 | -0.6681 | 0.059 | 0.598 | 0.0262 | 10 |
| Gabrg1 | -0.6683 | 0.183 | 0.611 | 3.20E-09 | 8 |
| Rpl15 | -0.6683 | 0.647 | 0.976 | 0.5061 | 10 |
| Actg1 | -0.6683 | 1 | 0.997 | 1 | 10 |
| Gria1 | -0.6686 | 0.288 | 0.614 | 2.09E-20 | 2 |
| Scn9a | -0.6694 | 0.176 | 0.602 | 1 | 10 |
| Syt10 | -0.6696 | 0.455 | 0.687 | 6.54E-16 | 0 |
| Auts2 | -0.6705 | 0.461 | 0.782 | 2.04E-19 | 1 |
| Tac1 | -0.6737 | 0.026 | 0.102 | 1 | 1 |
| 15-Sep | -0.6738 | 0.647 | 0.952 | 1 | 10 |
| Negr1 | -0.6756 | 0.513 | 0.861 | 4.00E-13 | 7 |
| Spcs2 | -0.676 | 0.765 | 0.963 | 1 | 10 |
| Cetn3 | -0.678 | 0.176 | 0.673 | 0.2701 | 10 |
| Chgb | -0.6784 | 0.664 | 0.882 | 3.54E-07 | 2 |
| Gpx4 | -0.6784 | 0.941 | 0.988 | 1 | 10 |
| Msi2 | -0.6785 | 0.367 | 0.664 | 0.3903 | 9 |
| Stmn1 | -0.6791 | 1 | 0.999 | 1 | 3 |
| Rps16 | -0.6797 | 0.941 | 0.985 | 1 | 10 |
| Chl1 | -0.6808 | 0.103 | 0.503 | 1.26E-09 | 7 |
| Rgs16 | -0.6814 | 0.241 | 0.37 | 0.1977 | 4 |
| Spock3 | -0.6819 | 0.169 | 0.575 | 1.72E-09 | 8 |
| Rps8 | -0.6819 | 0.941 | 0.999 | 1 | 10 |

| | | | | | |
|---|---|---|---|---|---|
| Fau | -0.6822 | 1 | 0.997 | 1 | 10 |
| Parm1 | -0.6822 | 0.176 | 0.593 | 1 | 10 |
| Pou2f2 | -0.6824 | 0.333 | 0.609 | 1 | 9 |
| Ppfia2 | -0.6835 | 0.27 | 0.623 | 1.86E-18 | 1 |
| Usmg5 | -0.6854 | 0.992 | 0.99 | 0.0233 | 3 |
| Tmed3 | -0.6857 | 0.118 | 0.637 | 0.0996 | 10 |
| Sap18 | -0.6858 | 0.235 | 0.801 | 0.1884 | 10 |
| Cfdp1 | -0.686 | 0.529 | 0.93 | 1 | 10 |
| Bcas2 | -0.6881 | 0.706 | 0.938 | 1 | 10 |
| Rpl36 | -0.6888 | 1 | 0.993 | 1 | 10 |
| Lsamp | -0.6893 | 0.658 | 0.872 | 1.01E-14 | 2 |
| Gm1673 | -0.6897 | 0.941 | 0.991 | 1 | 10 |
| Chchd1 | -0.6903 | 0.176 | 0.741 | 1 | 10 |
| Rgs16 | -0.6913 | 0.21 | 0.39 | 7.04E-05 | 0 |
| Rps24 | -0.6922 | 1 | 0.997 | 0.0046 | 3 |
| Tspan7 | -0.6933 | 0.705 | 0.869 | 8.24E-12 | 2 |
| Igfbp5 | -0.6933 | 0.3 | 0.637 | 1 | 9 |
| Uqcr10 | -0.6943 | 0.647 | 0.966 | 1 | 10 |
| Ppp1r11 | -0.6944 | 0.353 | 0.862 | 1 | 10 |
| Rpl27a | -0.695 | 0.882 | 0.996 | 1 | 10 |
| Eef1b2 | -0.6951 | 0.471 | 0.939 | 1 | 10 |
| Rps27a | -0.6971 | 1 | 0.994 | 0.0112 | 3 |
| Ptn | -0.6983 | 0.169 | 0.533 | 3.41E-06 | 8 |
| Rpl10a | -0.6989 | 0.647 | 0.977 | 1 | 10 |
| Pdcd5 | -0.6998 | 0.529 | 0.9 | 1 | 10 |
| Rps2 | -0.7 | 0.824 | 0.995 | 1 | 10 |
| Ptprd | -0.7003 | 0.579 | 0.88 | 2.07E-22 | 1 |
| Txndc17 | -0.7005 | 0.118 | 0.697 | 0.4323 | 10 |
| Ppp1r1a | -0.7023 | 0.114 | 0.444 | 0.3705 | 3 |
| Fis1 | -0.7061 | 0.471 | 0.94 | 1 | 10 |
| Tsc22d1 | -0.7069 | 0.603 | 0.901 | 2.73E-24 | 2 |
| Mrpl52 | -0.7075 | 0.412 | 0.872 | 1 | 10 |
| Lhx1os | -0.7075 | 0.385 | 0.578 | 3.51E-11 | 0 |
| Rpl34 | -0.7075 | 0.765 | 0.976 | 1 | 10 |
| Tle4 | -0.7089 | 0.233 | 0.682 | 0.0126 | 9 |
| Igfbp5 | -0.7099 | 0.398 | 0.653 | 8.47E-08 | 4 |
| Lrp1b | -0.7109 | 0.269 | 0.672 | 1.02E-11 | 7 |
| Nrgn | -0.7135 | 0.103 | 0.249 | 0.0001 | 5 |
| Resp18 | -0.7143 | 1 | 0.999 | 1 | 9 |
| Mrps21 | -0.7144 | 0.235 | 0.78 | 1 | 10 |
| Cox8a | -0.7146 | 0.941 | 0.999 | 1 | 10 |
| Srsf3 | -0.7146 | 0.706 | 0.967 | 1 | 10 |

| | | | | | |
|---|---|---|---|---|---|
| Ndufa2 | -0.7152 | 0.706 | 0.943 | 1 | 10 |
| Scand1 | -0.7165 | 0.353 | 0.826 | 1 | 10 |
| Atp5g2 | -0.7176 | 0.765 | 0.975 | 1 | 10 |
| mt-Nd2 | -0.718 | 0.993 | 1 | 5.65E-19 | 2 |
| Ppfia2 | -0.7233 | 0.169 | 0.602 | 2.54E-09 | 8 |
| Rpl35a | -0.7271 | 0.984 | 0.993 | 0.0016 | 3 |
| Rps3 | -0.728 | 0.882 | 0.983 | 1 | 10 |
| Nxph1 | -0.728 | 0.007 | 0.44 | 7.59E-31 | 1 |
| 23100360 | -0.7287 | 0.294 | 0.846 | 1 | 10 |
| Grm5 | -0.729 | 0.141 | 0.599 | 5.24E-11 | 8 |
| Ubl5 | -0.7294 | 0.882 | 0.975 | 1 | 10 |
| Rpl8 | -0.7296 | 0.941 | 0.999 | 1 | 10 |
| Nr2f2 | -0.7311 | 0 | 0.449 | 4.34E-17 | 8 |
| Zbtb20 | -0.732 | 0.449 | 0.761 | 4.37E-11 | 7 |
| Ap1s2 | -0.7321 | 0.176 | 0.617 | 0.4824 | 10 |
| Nrgn | -0.7346 | 0.066 | 0.263 | 2.12E-08 | 1 |
| Alcam | -0.7362 | 0.467 | 0.808 | 0.3876 | 9 |
| Rorb | -0.7369 | 0.235 | 0.677 | 1 | 10 |
| Gad2 | -0.7389 | 0.481 | 0.855 | 3.02E-19 | 4 |
| Sec61g | -0.7401 | 0.647 | 0.96 | 1 | 10 |
| Uqcc2 | -0.7423 | 0.588 | 0.928 | 1 | 10 |
| Fam19a1 | -0.7433 | 0 | 0.437 | 0.0001 | 9 |
| Rps3a1 | -0.7435 | 0.976 | 0.98 | 6.53E-05 | 3 |
| Prmt2 | -0.7451 | 0.588 | 0.922 | 0.795 | 10 |
| Gm10076 | -0.746 | 0.493 | 0.746 | 5.43E-08 | 2 |
| Cox7a2 | -0.7467 | 0.824 | 0.983 | 1 | 10 |
| Rora | -0.7474 | 0.086 | 0.507 | 7.90E-23 | 1 |
| Ngb | -0.7477 | 0.485 | 0.806 | 4.27E-11 | 5 |
| Rpl19 | -0.7497 | 1 | 0.999 | 1 | 10 |
| Cartpt | -0.7501 | 0.021 | 0.137 | 1 | 6 |
| Gabrb1 | -0.7511 | 0.563 | 0.876 | 2.32E-09 | 8 |
| Flrt3 | -0.7547 | 0.067 | 0.562 | 0.0004 | 9 |
| Chchd2 | -0.7561 | 0.941 | 0.999 | 1 | 10 |
| Ranbp1 | -0.7577 | 0.471 | 0.913 | 1 | 10 |
| Cox7b | -0.7624 | 0.588 | 0.92 | 1 | 10 |
| Rpl24 | -0.7642 | 1 | 0.998 | 1 | 10 |
| Atp5g3 | -0.7658 | 0.882 | 0.979 | 1 | 10 |
| Tusc3 | -0.7671 | 0.765 | 0.972 | 1 | 10 |
| Thy1 | -0.7673 | 0.113 | 0.442 | 5.29E-05 | 8 |
| Pkib | -0.7688 | 0.118 | 0.474 | 1 | 10 |
| Zfhx3 | -0.7688 | 0.366 | 0.756 | 8.95E-11 | 8 |
| Mrfap1 | -0.7703 | 0.941 | 0.995 | 1 | 10 |

| | | | | | |
|---|---|---|---|---|---|
| Eef1a1 | -0.7715 | 1 | 1 | 2.69E-11 | 3 |
| Eef1a1 | -0.7717 | 1 | 1 | 1 | 10 |
| Rpsa | -0.7724 | 0.941 | 0.999 | 0.4167 | 10 |
| Cartpt | -0.7735 | 0.016 | 0.141 | 1 | 3 |
| Nap1l5 | -0.7738 | 1 | 0.999 | 0.0138 | 10 |
| Pbx1 | -0.7761 | 0.441 | 0.814 | 2.88E-26 | 1 |
| Higd1a | -0.7778 | 0.268 | 0.818 | 3.54E-22 | 8 |
| Btf3 | -0.7789 | 0.706 | 0.975 | 1 | 10 |
| Fam19a1 | -0.7795 | 0.026 | 0.488 | 9.22E-32 | 1 |
| Cisd1 | -0.7807 | 0.647 | 0.968 | 1 | 10 |
| Oaz1 | -0.7816 | 1 | 0.998 | 0.557 | 10 |
| Cd24a | -0.784 | 0.305 | 0.528 | 2.40E-15 | 0 |
| Map1lc3a | -0.7853 | 0.882 | 0.973 | 1 | 10 |
| Lin7a | -0.7866 | 0.176 | 0.693 | 1 | 10 |
| Tpbg | -0.7869 | 0.325 | 0.545 | 3.06E-14 | 0 |
| Slc32a1 | -0.7869 | 0.454 | 0.885 | 2.48E-30 | 4 |
| Snap25 | -0.7885 | 0.718 | 0.936 | 2.30E-10 | 8 |
| Bex2 | -0.7897 | 1 | 1 | 1 | 10 |
| Zcrb1 | -0.7918 | 0.176 | 0.818 | 0.3606 | 10 |
| Trappc4 | -0.7939 | 0.235 | 0.836 | 1 | 10 |
| Grcc10 | -0.7949 | 0.412 | 0.9 | 1 | 10 |
| Thy1 | -0.7973 | 0.155 | 0.446 | 7.51E-11 | 5 |
| Sf3b6 | -0.7981 | 0.412 | 0.892 | 0.2852 | 10 |
| Sepw1 | -0.7997 | 1 | 0.995 | 1 | 10 |
| Arhgap6 | -0.8 | 0.167 | 0.618 | 0.0047 | 9 |
| Myeov2 | -0.8016 | 0.412 | 0.905 | 1 | 10 |
| Six6 | -0.8028 | 0.059 | 0.425 | 1 | 10 |
| Rit2 | -0.8074 | 0.412 | 0.895 | 1 | 10 |
| Tmem158 | -0.8103 | 0.308 | 0.744 | 1.10E-18 | 7 |
| Rora | -0.8108 | 0.028 | 0.478 | 8.30E-14 | 8 |
| Eif1b | -0.8118 | 0.824 | 0.979 | 1 | 10 |
| Snap25 | -0.8119 | 0.796 | 0.936 | 3.96E-17 | 4 |
| Rps5 | -0.8126 | 0.882 | 0.992 | 1 | 10 |
| Camk2n1 | -0.813 | 0.233 | 0.773 | 5.14E-07 | 9 |
| Atp5d | -0.816 | 0.706 | 0.966 | 1 | 10 |
| Zfhx3 | -0.816 | 0.352 | 0.772 | 1.41E-22 | 4 |
| Zfhx3 | -0.8173 | 0.235 | 0.739 | 1 | 10 |
| Tspan7 | -0.8182 | 0.546 | 0.877 | 3.24E-16 | 5 |
| Malat1 | -0.8201 | 1 | 0.999 | 0.0045 | 10 |
| Rpl18 | -0.8207 | 0.765 | 0.985 | 1 | 10 |
| Rplp2 | -0.8214 | 0.941 | 0.986 | 1 | 10 |
| Naca | -0.824 | 1 | 0.985 | 1 | 10 |

| | | | | | |
|---|---|---|---|---|---|
| Lin7a | -0.8257 | 0.239 | 0.716 | 2.80E-13 | 8 |
| Sub1 | -0.8267 | 0.882 | 0.99 | 0.2898 | 10 |
| Psmd7 | -0.8272 | 0.412 | 0.881 | 1 | 10 |
| Atp6v1g1 | -0.8273 | 0.588 | 0.948 | 1 | 10 |
| Rpl23 | -0.8273 | 0.941 | 0.987 | 1 | 10 |
| Ndufa7 | -0.8283 | 0.588 | 0.959 | 1 | 10 |
| Jun | -0.8323 | 0.247 | 0.58 | 0.002 | 5 |
| Lin7a | -0.8335 | 0.267 | 0.697 | 0.0016 | 9 |
| Ntng1 | -0.8342 | 0.116 | 0.38 | 2.14E-14 | 2 |
| Atp5e | -0.8348 | 0.765 | 0.973 | 1 | 10 |
| Gabarap | -0.8353 | 0.647 | 0.958 | 1 | 10 |
| Clu | -0.8381 | 0.205 | 0.633 | 9.96E-16 | 7 |
| mt-Nd4l | -0.8392 | 1 | 1 | 3.02E-19 | 2 |
| Slc25a4 | -0.8412 | 1 | 0.999 | 0.1172 | 10 |
| Lsamp | -0.8423 | 0.526 | 0.874 | 7.61E-19 | 5 |
| Igfbp5 | -0.845 | 0.296 | 0.651 | 7.44E-07 | 8 |
| Pcdh17 | -0.8465 | 0.346 | 0.666 | 1.99E-11 | 7 |
| Pkib | -0.8466 | 0.092 | 0.527 | 7.18E-27 | 1 |
| Rgs16 | -0.8473 | 0.1 | 0.365 | 0.6893 | 9 |
| Fos | -0.8476 | 0.124 | 0.459 | 0.0021 | 5 |
| Rpl21 | -0.8493 | 0.941 | 0.998 | 1 | 10 |
| Tmsb4x | -0.8499 | 0.984 | 0.996 | 2.24E-07 | 3 |
| Marcksl1 | -0.8504 | 0.588 | 0.95 | 0.891 | 10 |
| 2700060I | -0.8511 | 0.235 | 0.843 | 1 | 10 |
| H2afz | -0.8515 | 0.824 | 0.982 | 0.1854 | 10 |
| Uqcrh | -0.8522 | 0.471 | 0.956 | 0.3462 | 10 |
| Pkib | -0.8542 | 0.056 | 0.496 | 9.65E-13 | 8 |
| Romo1 | -0.8555 | 0.706 | 0.966 | 1 | 10 |
| Selm | -0.8558 | 0.706 | 0.982 | 0.0286 | 10 |
| C1d | -0.8563 | 0.412 | 0.947 | 0.3268 | 10 |
| Six6 | -0.8637 | 0.028 | 0.461 | 1.66E-19 | 4 |
| Pcdh17 | -0.8642 | 0.315 | 0.693 | 3.72E-23 | 2 |
| Resp18 | -0.867 | 0.941 | 1 | 1 | 10 |
| Six6 | -0.8672 | 0 | 0.431 | 7.61E-05 | 9 |
| Pcbd1 | -0.8711 | 0.176 | 0.73 | 0.5017 | 10 |
| Ubb | -0.873 | 1 | 1 | 0.1218 | 3 |
| Cd24a | -0.8757 | 0.176 | 0.521 | 1.40E-11 | 4 |
| Swi5 | -0.878 | 0.647 | 0.944 | 1 | 10 |
| Rgs16 | -0.884 | 0.056 | 0.378 | 2.55E-07 | 8 |
| Pcsk2 | -0.8848 | 0.433 | 0.85 | 0.0127 | 9 |
| Srp14 | -0.8879 | 0.824 | 0.977 | 0.7536 | 10 |
| Six6 | -0.8896 | 0 | 0.448 | 1.48E-15 | 8 |

| | | | | | |
|---|---|---|---|---|---|
| Calb2 | -0.8903 | 0.268 | 0.6 | 1.08E-05 | 8 |
| Tceal3 | -0.8935 | 0.647 | 0.949 | 0.7368 | 10 |
| Thy1 | -0.8938 | 0.116 | 0.467 | 1.68E-30 | 2 |
| Fos | -0.8942 | 0.199 | 0.465 | 0.0094 | 2 |
| Ntng1 | -0.8942 | 0.052 | 0.374 | 5.41E-12 | 5 |
| Snap25 | -0.8945 | 0.789 | 0.943 | 6.53E-30 | 1 |
| Shfm1 | -0.8991 | 0.471 | 0.949 | 0.0236 | 10 |
| Cadps2 | -0.9017 | 0.111 | 0.599 | 6.58E-23 | 4 |
| Resp18 | -0.9022 | 1 | 0.999 | 1.91E-12 | 5 |
| Ftl1 | -0.9023 | 0.647 | 0.958 | 1 | 10 |
| Lhx1 | -0.9058 | 0.465 | 0.643 | 4.99E-18 | 0 |
| Rps28 | -0.9072 | 1 | 0.989 | 1 | 10 |
| Sumo1 | -0.9075 | 0.471 | 0.954 | 0.5357 | 10 |
| Cartpt | -0.9079 | 0.026 | 0.134 | 0.3453 | 7 |
| Ntng1 | -0.9091 | 0.026 | 0.37 | 1.32E-09 | 7 |
| Six6 | -0.9102 | 0.026 | 0.481 | 3.57E-32 | 1 |
| Clu | -0.9149 | 0.151 | 0.671 | 9.03E-34 | 2 |
| Trp53i11 | -0.9163 | 0.167 | 0.739 | 2.53E-06 | 9 |
| Rps10 | -0.9217 | 0.941 | 0.985 | 1 | 10 |
| Hpcal1 | -0.9228 | 0.282 | 0.446 | 0.0002 | 8 |
| Six3 | -0.9231 | 0.509 | 0.701 | 4.40E-09 | 4 |
| Mrps33 | -0.9247 | 0.412 | 0.919 | 1 | 10 |
| Id4 | -0.9256 | 0.118 | 0.652 | 0.6373 | 10 |
| Syt10 | -0.9275 | 0.267 | 0.656 | 0.0287 | 9 |
| Cadps2 | -0.9279 | 0.125 | 0.619 | 1.26E-32 | 1 |
| Arhgap6 | -0.9283 | 0.145 | 0.678 | 9.08E-39 | 1 |
| Syt10 | -0.9286 | 0.296 | 0.7 | 5.61E-25 | 1 |
| Nrxn3 | -0.93 | 0.556 | 0.948 | 2.43E-39 | 4 |
| Rps19 | -0.9315 | 0.882 | 0.991 | 1 | 10 |
| Hmgn1 | -0.933 | 0.529 | 0.928 | 1 | 10 |
| Auts2 | -0.9366 | 0.118 | 0.748 | 0.178 | 10 |
| Cartpt | -0.9452 | 0 | 0.139 | 0.0018 | 5 |
| Tac1 | -0.9475 | 0 | 0.099 | 1 | 7 |
| Tac1 | -0.9482 | 0.01 | 0.1 | 0.2956 | 5 |
| Tac1 | -0.9482 | 0.01 | 0.1 | 1 | 6 |
| Scg2 | -0.9494 | 0.873 | 0.968 | 4.84E-08 | 8 |
| Thy1 | -0.9544 | 0.026 | 0.451 | 2.02E-13 | 7 |
| Tac1 | -0.9579 | 0.008 | 0.102 | 1 | 3 |
| Rpl6 | -0.9598 | 1 | 0.998 | 1 | 10 |
| Tac1 | -0.9608 | 0.027 | 0.102 | 0.0013 | 2 |
| Rnf7 | -0.9628 | 0.235 | 0.907 | 0.009 | 10 |
| Synpr | -0.9629 | 0.52 | 0.76 | 7.78E-27 | 0 |

| | | | | | |
|---|---|---|---|---|---|
| Rpl36a | -0.9641 | 0.471 | 0.936 | 1 | 10 |
| Rps23 | -0.9644 | 0.824 | 0.974 | 1 | 10 |
| Pfdn5 | -0.9683 | 0.706 | 0.966 | 1 | 10 |
| Pcdh17 | -0.9684 | 0.216 | 0.684 | 5.53E-19 | 5 |
| Syt10 | -0.9691 | 0.254 | 0.672 | 1.10E-11 | 8 |
| Rps13 | -0.9736 | 0.647 | 0.984 | 0.0817 | 10 |
| Cartpt | -0.9755 | 0 | 0.146 | 6.72E-08 | 2 |
| Rpl11 | -0.9802 | 0.882 | 0.996 | 1 | 10 |
| Ncald | -0.9806 | 0.124 | 0.732 | 5.65E-30 | 5 |
| Ptma | -0.983 | 0.824 | 0.989 | 1 | 10 |
| Synpr | -0.9875 | 0.487 | 0.753 | 6.10E-17 | 1 |
| Cadps2 | -0.9881 | 0.042 | 0.587 | 3.25E-18 | 8 |
| Rps7 | -0.9904 | 0.824 | 0.983 | 1 | 10 |
| Itm2b | -0.9939 | 1 | 0.999 | 1 | 10 |
| Cd24a | -0.999 | 0.1 | 0.499 | 0.0155 | 9 |
| Rps9 | -1.0003 | 0.588 | 0.98 | 1 | 10 |
| Cd24a | -1.0037 | 0.145 | 0.542 | 3.89E-23 | 1 |
| Nrxn3 | -1.0069 | 0.706 | 0.913 | 0.6607 | 10 |
| Tmem176 | -1.0085 | 0 | 0.691 | 0.0038 | 10 |
| Rpl37a | -1.0085 | 1 | 0.998 | 0.0557 | 10 |
| Hpcal1 | -1.0103 | 0.203 | 0.464 | 7.02E-14 | 3 |
| Igfbp5 | -1.0117 | 0.217 | 0.693 | 6.87E-32 | 1 |
| Uba52 | -1.0136 | 0.353 | 0.882 | 0.007 | 10 |
| Hmgn2 | -1.018 | 0.294 | 0.914 | 0.9582 | 10 |
| Rpl3 | -1.0204 | 1 | 1 | 0.4187 | 10 |
| Gad1 | -1.021 | 0.059 | 0.77 | 1.97E-06 | 10 |
| Rpl35a | -1.0218 | 0.882 | 0.994 | 1 | 10 |
| Bex1 | -1.0249 | 0.706 | 0.982 | 0.7662 | 10 |
| Cd24a | -1.0299 | 0.059 | 0.495 | 1 | 10 |
| Alcam | -1.0339 | 0.38 | 0.827 | 1.28E-16 | 8 |
| Penk | -1.0353 | 0 | 0.357 | 1.11E-10 | 8 |
| Rps21 | -1.0398 | 0.824 | 0.988 | 1 | 10 |
| Rps29 | -1.0465 | 1 | 0.997 | 1 | 10 |
| Cd24a | -1.0513 | 0.07 | 0.516 | 2.34E-12 | 8 |
| Jund | -1.0701 | 0.941 | 0.988 | 1 | 10 |
| Tpbg | -1.0748 | 0.102 | 0.549 | 1.00E-20 | 4 |
| Zfhx3 | -1.0762 | 0.067 | 0.749 | 9.66E-10 | 9 |
| Spcs1 | -1.0863 | 0.588 | 0.961 | 0.1616 | 10 |
| Ubb | -1.0933 | 1 | 1 | 0.0145 | 10 |
| Hint1 | -1.0967 | 0.706 | 0.992 | 0.0029 | 10 |
| Tpbg | -1.0979 | 0.042 | 0.537 | 3.93E-14 | 8 |
| Lhx1os | -1.1 | 0.13 | 0.588 | 7.57E-23 | 4 |

| | | | | | |
|---|---|---|---|---|---|
| Hpcal1 | -1.1045 | 0.072 | 0.47 | 3.18E-14 | 5 |
| Hpcal1 | -1.1047 | 0.09 | 0.461 | 2.62E-13 | 7 |
| Tpbg | -1.1081 | 0.033 | 0.519 | 0.0002 | 9 |
| Rpl39 | -1.1087 | 0.647 | 0.981 | 0.1529 | 10 |
| Rpl22l1 | -1.1101 | 0.941 | 0.998 | 0.0361 | 10 |
| Rorb | -1.1144 | 0.1 | 0.686 | 3.90E-08 | 9 |
| Tpbg | -1.1375 | 0.086 | 0.572 | 1.41E-33 | 1 |
| Fxyd6 | -1.14 | 0.647 | 0.996 | 2.66E-07 | 10 |
| Calb2 | -1.1414 | 0.191 | 0.64 | 6.59E-29 | 1 |
| Gabarapl2 | -1.1483 | 0.647 | 0.993 | 0.0007 | 10 |
| Cst3 | -1.1512 | 0.765 | 0.993 | 0.0815 | 10 |
| 1110004 | -1.1536 | 0.471 | 0.971 | 0.0255 | 10 |
| Gal | -1.155 | 0.32 | 0.352 | 1 | 6 |
| Btg1 | -1.1576 | 0.059 | 0.789 | 0.001 | 10 |
| Eif1 | -1.1586 | 0.941 | 0.999 | 0.0025 | 10 |
| Gal | -1.1617 | 0.345 | 0.35 | 1 | 0 |
| Rpl9 | -1.1647 | 0.765 | 0.991 | 0.5919 | 10 |
| Lhx1os | -1.1668 | 0.042 | 0.577 | 1.99E-18 | 8 |
| Lhx1os | -1.1676 | 0.099 | 0.613 | 2.13E-38 | 1 |
| Lhx1os | -1.174 | 0 | 0.552 | 1 | 10 |
| Hpcal1 | -1.1777 | 0.062 | 0.491 | 6.57E-31 | 2 |
| Lhx1os | -1.1823 | 0 | 0.558 | 4.76E-07 | 9 |
| Scg2 | -1.1881 | 0.824 | 0.964 | 1 | 10 |
| Rpl41 | -1.1899 | 1 | 1 | 0.0149 | 10 |
| Gad2 | -1.1934 | 0.118 | 0.829 | 0.001 | 10 |
| Slc32a1 | -1.1973 | 0.118 | 0.855 | 0.0001 | 10 |
| Syt10 | -1.2096 | 0 | 0.655 | 0.0144 | 10 |
| H3f3b | -1.2184 | 0.941 | 0.999 | 3.00E-05 | 10 |
| Rps20 | -1.222 | 0.882 | 0.98 | 0.9257 | 10 |
| Rpl13 | -1.2232 | 0.882 | 0.998 | 0.001 | 10 |
| Cirbp | -1.2292 | 0.529 | 0.979 | 0.0005 | 10 |
| Six3 | -1.2449 | 0.329 | 0.738 | 4.52E-33 | 1 |
| Syt1 | -1.2554 | 0.564 | 0.819 | 1.43E-13 | 7 |
| Dlk1 | -1.2601 | 0.235 | 0.714 | 1 | 10 |
| Rps24 | -1.2617 | 1 | 0.997 | 1 | 10 |
| Rps4x | -1.2837 | 0.765 | 0.985 | 0.099 | 10 |
| Lhx1 | -1.2915 | 0.059 | 0.62 | 1 | 10 |
| Six3 | -1.299 | 0.059 | 0.692 | 1 | 10 |
| Rps3a1 | -1.3011 | 0.765 | 0.983 | 0.4561 | 10 |
| Calb1 | -1.3017 | 0.155 | 0.804 | 2.38E-29 | 8 |
| Lhx1 | -1.3155 | 0.222 | 0.653 | 1.14E-25 | 4 |
| Rpl30 | -1.3508 | 0.824 | 0.992 | 0.0098 | 10 |

| | | | | | |
|---|---|---|---|---|---|
| Lhx1 | -1.3721 | 0.127 | 0.644 | 5.91E-18 | 8 |
| Lhx1 | -1.3869 | 0.178 | 0.679 | 1.09E-41 | 1 |
| Six3 | -1.3919 | 0.155 | 0.719 | 3.92E-21 | 8 |
| Synpr | -1.395 | 0.235 | 0.724 | 1 | 10 |
| Rps27a | -1.3986 | 0.824 | 0.997 | 0.002 | 10 |
| Syt1 | -1.4135 | 0.493 | 0.848 | 1.79E-29 | 2 |
| Synpr | -1.4182 | 0.225 | 0.75 | 5.55E-20 | 8 |
| Six3 | -1.4245 | 0.1 | 0.699 | 1.35E-07 | 9 |
| Lhx1 | -1.4308 | 0.033 | 0.627 | 2.16E-07 | 9 |
| Vstm2l | -1.4436 | 0.183 | 0.889 | 1.99E-43 | 8 |
| Rps27 | -1.455 | 0.941 | 0.994 | 5.61E-05 | 10 |
| Gal | -1.4713 | 0.282 | 0.354 | 1 | 8 |
| Dlk1 | -1.5233 | 0.133 | 0.723 | 8.14E-07 | 9 |
| Pcp4 | -1.5762 | 0.454 | 0.921 | 2.28E-33 | 5 |
| Grp | -1.7671 | 0.245 | 0.366 | 1.07E-10 | 0 |
| Grp | -2.1096 | 0.093 | 0.368 | 1.36E-06 | 5 |
| Grp | -2.1374 | 0.083 | 0.372 | 2.03E-11 | 4 |
| Grp | -2.142 | 0 | 0.349 | 1 | 10 |
| Grp | -2.1525 | 0 | 0.354 | 0.0664 | 9 |
| Gal | -2.1547 | 0.317 | 0.353 | 4.93E-10 | 3 |
| Grp | -2.1585 | 0.028 | 0.365 | 1.46E-07 | 8 |
| Grp | -2.1624 | 0.092 | 0.384 | 2.47E-15 | 1 |
| Gal | -2.3215 | 0.154 | 0.364 | 1.39E-07 | 7 |
| Gal | -2.4331 | 0.052 | 0.378 | 1.13E-08 | 5 |
| Gal | -2.4914 | 0.048 | 0.395 | 2.09E-15 | 2 |
| Vip | -2.552 | 0.25 | 0.46 | 3.15E-10 | 4 |
| Vip | -2.5734 | 0.295 | 0.471 | 6.39E-22 | 0 |
| Vip | -2.6276 | 0.211 | 0.476 | 6.62E-17 | 1 |
| Vip | -2.6307 | 0.127 | 0.461 | 8.75E-08 | 8 |
| Vip | -2.6476 | 0.103 | 0.472 | 1.48E-09 | 5 |
| Vip | -2.6819 | 0.033 | 0.451 | 0.012 | 9 |
